# A Refined Phylochronology of the Second Plague Pandemic in Western Eurasia

**DOI:** 10.1101/2023.07.18.549544

**Authors:** Marcel Keller, Meriam Guellil, Philip Slavin, Lehti Saag, Kadri Irdt, Helja Niinemäe, Anu Solnik, Martin Malve, Heiki Valk, Aivar Kriiska, Craig Cessford, Sarah A. Inskip, John E. Robb, Christine Cooper, Conradin von Planta, Mathias Seifert, Thomas Reitmaier, Willem A. Baetsen, Don Walker, Sandra Lösch, Sönke Szidat, Mait Metspalu, Toomas Kivisild, Kristiina Tambets, Christiana L. Scheib

## Abstract

Although dozens of ancient *Yersinia pestis* genomes and a vast corpus of documentary data are available, the origin and spread of consecutive outbreaks of the Second Plague Pandemic in Europe (14th–18th c.) are still poorly understood. For the majority of ancient genomes, only radiocarbon dates spanning several decades are available, hampering an association with historically recorded plague outbreaks. Here, we present new genomic evidence of the Second Pandemic from 11 sites in England, Estonia, the Netherlands, Russia, and Switzerland yielding 11 *Y. pestis* genomes with >4-fold mean coverage dating to between 1349 and 1710. In addition, we present a novel approach for integrating the chronological information retrieved from phylogenetic analysis with their respective radiocarbon dates, based on a novel methodology offering more precise dating intervals. Together with a fine-grained analysis of documentarily recorded plague outbreaks, this allows us to tentatively associate all available *Y. pestis* genomes of the Second Pandemic with historically documented plague outbreaks. Through these combined multidisciplinary analytical efforts, our newly sequenced genomes can be attributed to the Black Death in Cambridge (England), the *pestis tertia* or *pestis quarta* in the late 14th century (Estonia), previously unknown branches emerging in the 15th century (Estonia, the Netherlands and England), and a widespread pandemic in Eastern Europe around 1500 (western Russia), which all seem to have originated from one or multiple reservoirs located in Central Europe. While the latter continued to harbour a major *Y. pestis* lineage at least until the 1630s, represented by new genomes of the Thirty Years’ War plague (Switzerland), another lineage consecutively spread into Europe between the 17th and 18th century from the Ottoman Empire, as evidenced by a genome associated with the Great Northern War plague (Estonia). By combining phylogenetic analysis with a systematic historical reconstruction based on textual sources and an innovative phylogenetically informed radiocarbon modelling (PhIRM), we offer a new groundbreaking interdisciplinary approach that solves several fundamental methodological challenges associated with phylogenetic and spatio-temporal reconstruction of historical pandemics.

## Introduction

Since the publication of the first *Yersinia pestis* genome reconstructed from ancient DNA^1^, palaeogenetics has increasingly become an integral part of interdisciplinary plague studies. Analyses of ancient genomes have directly confirmed the involvement of *Y. pestis* in plague pandemics, settled a number of previous controversies and offered some novel insights for the study of the history of the Second Plague Pandemic. Recently published genomes from Kara-Djigach, Kyrgyzstan, confirm the emergence of the Black Death and all known Second Pandemic strains in earlier fourteenth-century Central Asia^2^. The retrieval of identical genomes from eight different sites in England, Germany, Italy, France, Norway and Spain dating to the mid-14th century showed the rapid spread of a single *Y. pestis* lineage during the Black Death in Europe (1347–1353) followed by a split into two branches, Branch 1B, associated with the *pestis secunda* of 1356–1366^3^ and later giving rise to all modern Branch 1 genomes, including those associated with the Third Pandemic, and Branch 1A, comprising exclusively of Second Pandemic genomes retrieved, as of 2023, from different spatio-temporal contexts from Europe and the Caucasus from the late 14th to the late 18th century. Whereas genomes of the early Second Pandemic and Branch 1A2, with the youngest genomes dating to the Thirty Years’ War^4–6^, show no fundamental alteration in their genomic makeup that would lead to a higher virulence or attenuation, the 17th–18th-century genomes of Branch 1A1 show a large genomic deletion with currently unknown consequences for the bacterium coinciding with a higher substitution rate^4^. Beyond the reconstruction of the evolutionary history of *Y. pestis* during the Second Pandemic, the retrieval of ancient *Y. pestis* DNA can offer crucial information for research questions of other related disciplines. For instance, the analysis of a broad diversity of burial customs during plague outbreaks^7^, or the refinement of models for zoonotic spillover events in the context of specific climatic fluctuations and changing ecological contexts can improve our understanding of past pandemics.

However, the insights offered by palaeogenetics of the Second Pandemic are currently hampered by the imprecise dating of the vast majority of reconstructed *Y. pestis* genomes. Apart from exceptionally precisely dated contexts such as the Kara-Djigach genomes (1338– 1339)^2^, the Black Death genomes of East-Smithfield (1348–1350)^1^ or the genomes associated with the Great Plague of Marseille (1720–1722)^8^, we mostly rely on radiocarbon dating of the deceased individuals, often yielding time spans of 100–150 years. Not only does this temporal imprecision heavily affect molecular dating approaches – leading to broad timeframes for the age of internal nodes – but it also obscures spatiotemporal patterns, which otherwise could inform us about the timing and location of the emergence of new lineages. Without precise temporal information, attributions of events in pathogen evolution to anthropogenic or climatic factors, as often attempted for the well-dated onsets of the First and Second Pandemic, remain purely conjectural.

Here, we present newly reconstructed *Y. pestis* genomes for 11 samples (4.4–24.2-fold mean coverage) and genomic data for another 11 samples (1.0–4.4-fold mean coverage) from 11 sites in Estonia, the Netherlands, western Russia, Switzerland and England, spanning the late 14th century to the early 18th century. In addition, we present and apply a novel methodological approach – “phylogenetically informed radiocarbon modelling” (PhIRM) – for a refined dating of plague burials through the integration of phylogenetic information into Bayesian radiocarbon modelling. The refined dating is then juxtaposed against the documentary record, to associate the sequenced and analysed genomes with a series of documented plague outbreaks. This approach is applied both to the new genomes presented in this study as well as previously published genomes.

## Results

For this study, we screened 117 individuals (one sample per individual) from 11 sites for the presence of *Y. pestis*. The sites Domat/Ems (Switzerland), Tallinn Pärnu mnt 59B, Lehmja-Pildiküla, Otepää (Estonia) and Cambridge Bene’t Street (England) were identified as potential emergency burials linked to plague outbreaks through their archaeological or documentary context, while other sites such as Arnhem (the Netherlands) were sampled and screened for pathogens without previous suspicion of a plague context. After pathogen screening with KrakenUniq^9^, calculation of e-values, mappings to the chromosome and plasmids of *Y. pestis* as described in Material and Methods, we classified samples as negative, positive or tentatively positive. Of these, 26 were selected for a *Yersinia pestis/pseudotuberculosis-*specific enrichment with RNA baits. For sites with multiple hits and uniform temporal and archaeological context, only the best samples were selected. Except for three samples from the site of Mäletjärve, we built new UDG-treated libraries for all samples prior to enrichment to remove deamination damage prior to enrichment.

Using the newly sequenced samples MAL003/MAL004 and previously published genomes derived from non-UDG data with mean coverages of at least 3.5-fold, we tested different in-silico treatments for their ability to reduce false-positive SNPs due to damage and/or lenient mapping parameters while maintaining a correct phylogenetic placement. As shown in Supplementary Section 4, a lenient mapping followed by a rescaling of damaged sites with mapdamage2^10^ and a stricter re-mapping was found to be superior to all other tested methods in reducing false-positive SNPs without impacting the phylogenetic placement, e.g., by introducing a reference bias.

For the phylogenetic analyses, we initially considered 236 modern *Y. pestis* genomes. To exclude potentially problematic genomes, we filtered our modern reference panel based on the detection of homoplastic mutations, which are associated with excessively long terminal branches, and excluded genomes with 44 or more homoplastic mutations. In addition, we excluded two genomes due to excessive multiallelic sites, identified two cases of identical genomes and one case of identically labelled but genetically distinct genomes (see Supplementary Section 3, Table S2).

### Newly reconstructed *Y. pestis* genomes

Following target enrichment, between 3,989 and 2,084,944 unique reads mapping to the *Y. pestis* chromosome reference sequence (minimum mapping quality of 37) were retrieved for individual libraries (Table 1). We used eleven genomes with coverages between 4.4- and 24.2-fold for the reconstruction of a maximum likelihood tree (Table 1, Fig. 2B). Furthermore, we created a schematic tree (Fig. 2A; see Supplementary Section 4) by manually checking phylogenetically informative positions and filtering for false-positive SNPs for the same set of genomes (Supplementary Section 3, Table S5 and S6). Another phylogenetic tree including 208 modern and 67 ancient genomes with depths of coverages ranging from 1–4.4-fold was reconstructed to check for their phylogenetic placement (Fig. S37) and relevant SNPs were investigated in IGV (Fig. S34–42); see Supplementary Section 7 for a detailed discussion.

**Fig. 1:**
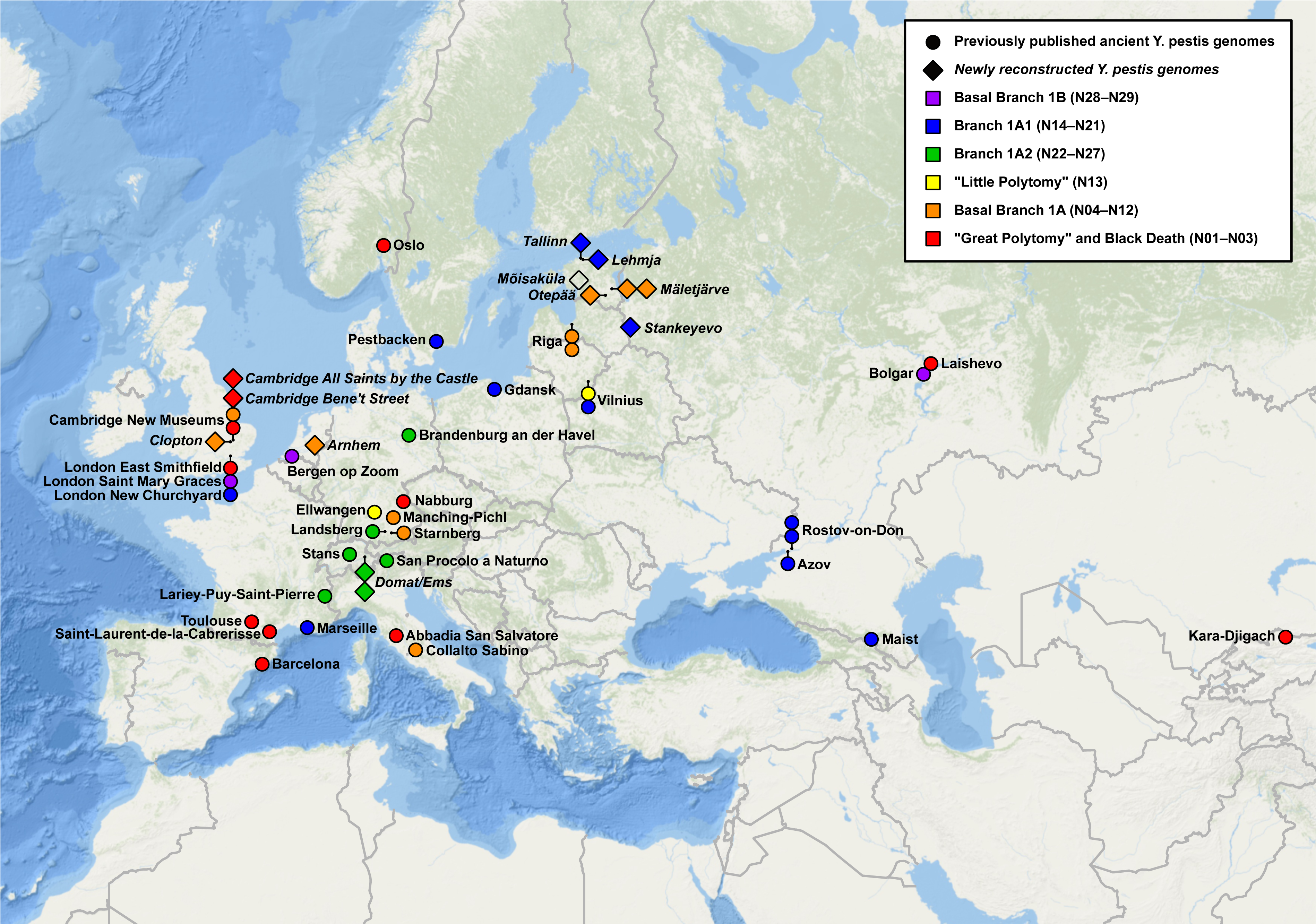
Map of all sites with palaeogenomic evidence for *Y. pestis* associated with the Second Pandemic from previous publications and this study. For an overview of all ancient *Y. pestis* genomes of the Second Pandemic, see Table S3.

**Fig. 2:**
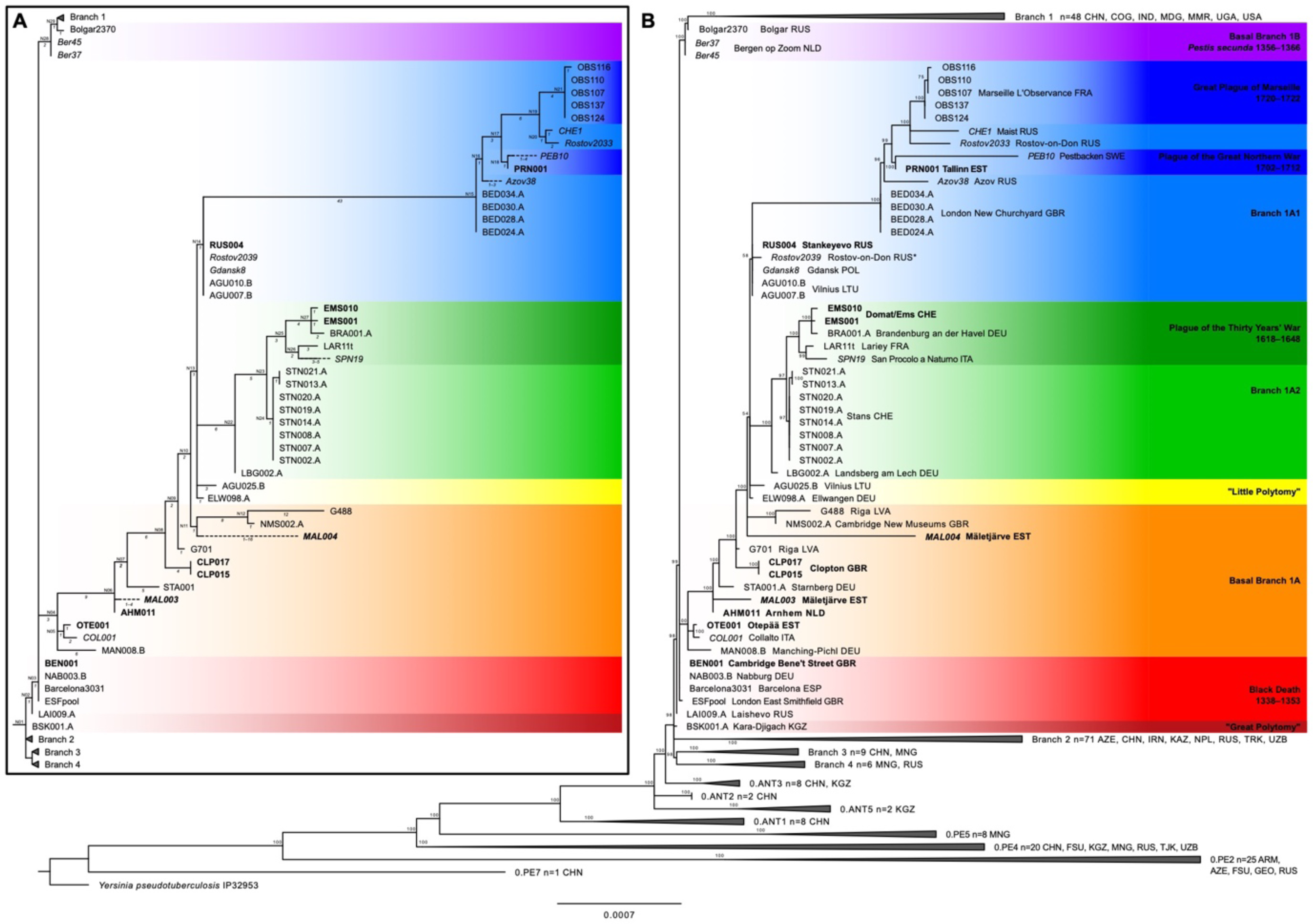
A: Schematic tree of *Y. pestis* genomes of the Second Pandemic. Newly sequenced genomes are highlighted in bold. Ancient genomes reconstructed from non-UDG data only are highlighted in italics. Branch lengths are shown in italics and correspond to Table S6; internal nodes are numbered from N01-N29. B: Maximum likelihood tree generated with IQTree with 1000 bootstraps based on a 95% partial deletion SNP alignment (3524 positions) of 56 ancient *Y. pestis* genomes, 208 modern *Y. pestis* genomes, and *Y. pseudotuberculosis* IP32953 as outgroup. For the corresponding SNP table see Table S5; for an expanded tree Fig. S19.

**Table 1:** Sequencing statistics after target enrichment, calibrated as well as modelled radiocarbon dates, and potential associated plague outbreak for all newly sequenced plague genomes.

| Site | Sample | Mapped reads | Mean read length | Mean coverage | Coverage $\geq 3X$ | Tree Fig. 2 | Tree Fig. S37 | RC date [cal CE 2σ] | RC date [modelled CE 2σ] | Potential plague outbreak |
| --- | --- | --- | --- | --- | --- | --- | --- | --- | --- | --- |
| St. Eusebius' churchyard, Arnhem, NLD | AHM001 | 175 990 | 52.78 | 2.0X | 31.30% | no | YES | 1307–1416 | 1385–1418 | 1398/1410–1412/1421 |
| St. Eusebius' churchyard, Arnhem, NLD | AHM011 | 855 496 | 52.46 | 9.6X | 79.90% | YES | YES | 1300–1399 | 1377–1416 | 1398/1410–1412/1421 |
| Bene't Street/Corpus Christi, Cambridge, GBR | BEN001 | 842 524 | 50.63 | 9.2X | 81.50% | YES | YES | N/A | N/A | 1349 |
| Bene't Street/Corpus Christi, Cambridge, GBR | BEN002 | 385 473 | 52.96 | 4.4X | 64.30% | no | YES | N/A | N/A | 1349 |
| St. Mary's, Clopton, Cambridgeshire, GBR | CLP015 | 633 008 | 59.22 | 8.1X | 84.40% | YES | YES | 1405–1443 | 1413–1458 | 1441–1443/1456–1459/1463–1465 |
| St. Mary's, Clopton, Cambridgeshire, GBR | CLP017 | 769 858 | 58.85 | 9.7X | 90.20% | YES | YES | 1317–1425 | 1406–1444 | 1441–1443/1456–1459/1463–1465 |
| All Saints by the Castle, Cambridge, GBR | COM042 | 242 669 | 50.25 | 2.6X | 41.80% | no | YES | N/A | N/A | 1349 |
| Sogn Pieder, Domat/Ems, CHE | EMS001 | 1 588 293 | 53.44 | 18.2X | 90.30% | YES | YES | 1433–1620 | 1491–1633 | 1629–1631 |
| Sogn Pieder, Domat/Ems, CHE | EMS002 | 49 238 | 52.19 | 0.6X | 3.90% | no | no | N/A | N/A | 1629–1631 |
| Sogn Pieder, Domat/Ems, CHE | EMS003 | 204 928 | 51.59 | 2.3X | 35.90% | no | YES | N/A | N/A | 1629–1631 |
| Sogn Pieder, Domat/Ems, CHE | EMS010 | 1 559 516 | 55.41 | 18.6X | 91.50% | YES | YES | 1431–1490 | 1495–1636 | 1629–1631 |
| Lehmja-Pildiküla, EST | LHP001 | 148 432 | 57.29 | 1.8X | 28.70% | no | YES | 1530–... | N/A | 1710 |
| Mäletjärve cemetery, EST | MAL003 | 466 083 | 59.06 | 5.9X | 75.80% | YES | YES | 1425–1472 | 1429–1622 | 1404/1420–1421/1424–1425 |
| Mäletjärve cemetery, EST | MAL004 | 369 376 | 54.94 | 4.4X | 60.00% | YES | YES | 1456–1633 | 1476–1641 | 1566–1567/1568–1570/1570–1571 |
| Mäletjärve cemetery, EST | MAL007 | 112 526 | 55.22 | 1.3X | 18.80% | no | YES | 1461–1635 | 1482–1645 | 1566–1567/1568–1570/1570–1571 |
| Mõisaküla village cemetery, EST | MOI001 | 15 233 | 46.53 | 0.2X | 1.50% | no | no | 1640–... | N/A | N/A |
| Otepää, EST | OTE001 | 735 679 | 50.56 | 8.0X | 76.10% | YES | YES | 1269–1380 | 1363–1394 | 1378–1379/1389–1390 |
| Pärnu mnt. 59, Tallinn, EST | PRN001 | 500 639 | 64.16 | 6.9X | 77.90% | YES | YES | 1670–... | N/A | 1710 |
| Pärnu mnt. 59, Tallinn, EST | PRN003 | 3 989 | 47.12 | 0.0X | 0.10% | no | no | 1693–1918 | N/A | 1710 |
| Pärnu mnt. 59, Tallinn, EST | PRN004 | 114 580 | 53.56 | 1.3X | 18.50% | no | YES | 1650–... | N/A | 1710 |
| Pärnu mnt. 59, Tallinn, EST | PRN005 | 98 688 | 44.92 | 1.0X | 11.30% | no | YES | 1694–1917 | N/A | 1710 |
| Pärnu mnt. 59, Tallinn, EST | PRN008 | 248 523 | 64.75 | 3.5X | 58.50% | no | YES | 1666–... | N/A | 1710 |
| Pärnu mnt. 59, Tallinn, EST | PRN009 | 205 867 | 67.81 | 3.0X | 50.40% | no | YES | 1643–... | N/A | 1710 |
| Pärnu mnt. 59, Tallinn, EST | PRN010 | 70 686 | 46.63 | 0.7X | 6.40% | no | no | 1660–... | N/A | 1710 |
| Stankeyevo village cemetery, RUS | RUS003 | 307 828 | 50.41 | 3.3X | 49.60% | no | YES | 1447–1631 | 1455–1572 | c.1474–1475/c.1482–1484/c.1495–1496/c.1505–1506 |
| Stankeyevo village cemetery, RUS | RUS004 | 2 084 944 | 54.11 | 24.2X | 91.00% | YES | YES | 1450–1631 | 1456–1575 | c.1474–1475/c.1482–1484/c.1495–1496/c.1505–1506 |

The high-coverage genome from Cambridge Bene’t Street, BEN001, is identical with seven other genomes associated with the Black Death in Europe 1347–1353 – Abbadia San Salvatore (Italy), Barcelona (Spain), London East-Smithfield (England), Nabburg (Germany), Oslo (Norway), Saint-Laurent-de-la-Cabrerisse and Toulouse (both in France). As far as can be reconstructed from the low-coverage genomic data (Supplementary Section 7), another genome from the same site, BEN002, and COM042, retrieved from Cambridge All Saints by the Castle, are identical with the aforementioned genomes as well. Together with the previously published genome from Cambridge Augustinian Friary, NMS003, we now have evidence for three funerary spaces within the medieval city of Cambridge utilised to bury victims of the Black Death, including single burials in regular cemeteries (Augustinian Friary, All Saints by the Castle) and a mass burial (Bene’t Street). All other newly reconstructed genomes presented in this study occupy positions on Branch 1A, which, so far, is only represented by ancient genomes of the Second Pandemic from Europe and the Caucasus (14th–18th centuries).

The genome of Otepää (Estonia), OTE001, represents the oldest Second Pandemic genome retrieved from the Baltic region, dating to the second half of the 14th century. It shares a short branch with the genome COL001 (Collalto Sabino, Italy), emerging from a trifurcation (node N04, according to the numbering established here; see Fig. 2, Supplementary Section 5) together with MAN008 (Manching-Pichl, Germany) and all other Branch 1A genomes, after the separation of the *pestis secunda* of 1356–1366 falling on basal Branch 1B. The MRCA of all other available Branch 1A genomes except MAN008, OTE001 and COL001 is represented by the genome of Arnhem (the Netherlands), AHM011, falling on node N06, and presumably the low-coverage genome AHM001 from the same site, likely situated on the same node. Another newly reported genome, MAL003, retrieved from Mäletjärve, Estonia, forms a terminal branch derived from AHM011. The two identical genomes from Clopton, a deserted medieval village in Cambridgeshire, derive from a trifurcation with the genome G701 from a multiple burial (“burial pit” with 15 individuals) from Riga St. Gertrude’s Church cemetery (Latvia). The second genome from Mäletjärve (Estonia), MAL004, forms a clade with NMS002.A, retrieved from a burial in the chapter house of the Augustinian Friary in Cambridge (England), and G488, from a different mass grave on St. Gertrude’s Church cemetery in Riga (Latvia). The short, shared branch of these three genomes is not visible in the Maximum Likelihood tree (Fig. 2B) due to low coverage (<3X) of the defining SNP in MAL004, but has been added to the schematic tree (Fig. 2A) after visual inspection. The clustering together of these genomes in a common clade is further supported by a deletion of the *inv* gene which has been previously reported by Spyrou et al. 2019 for NMS002.A^4^ and is also found in G488 and MAL004 (see Supplementary Section 8).

From the site of Domat/Ems (Switzerland), two distinct new genomes reconstructed here (EMS001 and EMS010) form a clade with the Brandenburg an der Havel genome (BRA001.A) emerging with short terminal branches defined by a single or two SNPs from a trifurcation. Together with the genomes from San Procolo a Naturno (SPN19) and Lariey-Puy-Saint-Pierre (Lar11t), they form a subclade of Branch 1A2, which is associated with the Thirty Years’ War plague (c.1628–1632)^5, 6^.

The genome of Stankeyevo (Pskov Oblast, Russia), RUS004, and presumably the low coverage genome of RUS003 from the same site are identical to previously published genomes of Gdańsk (Gdansk8), Vilnius (AGU007.B, AGU010.B) and the genome Rostov2039, associated in the original publication with Rostov-on-Don (but see Supplementary Section 11.10 for a discussion of its provenance). Situated one SNP derived from the polytomy N13, they represent a direct ancestor of all other Branch 1A1 genomes, but do not show the large genomic deletion observed in those (see Table S13, Fig. S43). The genome of Tallinn (Estonia), PRN001, is associated with a plague outbreak in the city, in the context of the plague of the Great Northern War in 1710. A previously published genome from Pestbacken (Sweden), PEB10, associated with the plague of the Great Northern War as well, is either identical with or directly derived from PRN001.

### Treatment of genomic data retrieved from non-UDG libraries

While the majority (n=53) of ancient *Y. pestis* genomes of the Second Pandemic are derived from UDG-treated or half-UDG-treated libraries, a smaller set is derived from non-UDG-treated libraries (n=20). Among the newly sequenced genomes presented here, two are reconstructed from non-UDG data. A common problem of non-UDG derived genomes is deamination patterns since they can interfere with the mapping of metagenomic ancient DNA data. The presence of deaminated sites is most commonly accommodated with allowance of a higher mismatch rate; a low mismatch rate tolerance threshold is thought to introduce a reference bias, to give preference to reads originating from modern contaminants (more relevant in context of human DNA) and reduce the genomic coverage. A higher mismatch rate, on the other hand, allows more reads from non-target organisms to map to the reference, causing an increased number of false-positive SNPs and ‘heterozygous’ positions. We, therefore, tested different in-silico non-UDG data treatment strategies and compared them on their outcome on private SNPs, phylogenetically informative positions and ultimately on phylogenetic analyses (see Supplementary Section 4).

Across all tested non-UDG samples, a combination of damage rescaling (mapdamage2^10^) after mapping with low stringency (bwa-aln -n 0.01^11^) and remapping with higher stringency (bwa-aln -n 0.1), showed the strongest reduction of excessive private SNPs and ‘heterozygous’ positions. Effects on phylogenetically informative positions were variable depending on the sample, but are proportional for C>T and G>A SNPs compared to other positions. Regarding the phylogenetic analyses, none of the tested treatments caused a reference bias strong enough to affect the tree topology significantly (see Supplementary Section 4). However, both rescaling alone as well as rescaling and remapping show the strongest effect on shortening terminal branches, while rescaling and remapping shows more accurate phylogenetic placements of the samples Rostov2039 and MAL004 based on the visual inspection of phylogenetically relevant SNPs (see Supplementary section 5).

### Phylogenetically informed Radiocarbon Modelling

To increase the resolution of dating intervals for plague genomes without precise documentary dating of buried individuals, we developed a novel approach using OxCal4.4.4^12^ and IntCal20^13^ by implementing the relative chronology embedded in the phylogenetic tree of *Y. pestis* in the form of nested sequences and phases within a common model. For dates of the same archaeological context or dates of identical genomes, we utilised Kernel Density Plots to summarise the probability distributions^14^, since this function, e.g., in contrast to the function R_Combine, allows for a variety of offsets such as reservoir effects, human bone collagen offset (HBCO, i.e. the difference between the year of death and the apparent age of the bone)^15, 16^ or laboratory specific biases.

Using radiocarbon dates without correction yielded some results, but gave poor agreement values of 34 (Amodel) and 36.6 (Aoverall) with five radiocarbon dates not passing the threshold of 60 (Code S5, Table S11, Fig. S46). This was triggered by individual radiocarbon dates that show significantly older prior than posterior probability distributions. Such a discrepancy may have been caused by the marine reservoir effect^17^, freshwater reservoir effect^18^ and/or an HBCO which all result in a bias towards older ages compared to the age-at-death. While the HBCO ranges between years and a few decades^15, 16^, marine and freshwater reservoir effects can cause offsets of several centuries in individuals with substantial marine or limnic dietary components^17, 18^. The individual samples may have been affected by these offsets to different degrees depending on the place of residence, diet, and age-at-death of the dated individual, as well as on the dated skeletal element. However, the majority of previously published radiocarbon dates provide neither carbon nor nitrogen isotope ratios to reconstruct proportions of marine or freshwater resources in diet, and the anthropological age estimation and the dated skeletal element were not always reported. In addition to the lack of data, there are also methodological challenges since the interpolation of dietary components is always an approximation and dependent on local reference data. Furthermore, the HBCO has not been examined systematically with large cohorts and a variety of skeletal elements. We therefore refrained from determining or estimating offsets (ΔR) on individual radiocarbon dates, but tested two approaches of applying the same offset to all radiocarbon dates in the model. Since we included samples from juvenile individuals and sites distant from marine or limnic shores unlikely to have any offset, we chose a moderate offset with a broad distribution: a normally distributed ΔR with median of 30 radiocarbon years and a standard deviation of 20 [ΔR (30, 20)], and a uniform distribution between 0 and 50 radiocarbon years [ΔR U(0, 50)], which can only be applied to MCMC models. Due to the inclusion of the offsets into the Bayesian modelling, the posterior distributions of the ΔR can be examined as well.

Within the sequence model, we used the Black Death with a calendar date of 1350±2 as absolute *terminus post quem* for all Branch 1A genomes and the Great Northern War plague with a calendar date of 1710±1 as absolute *terminus ante quem* for the direct ancestors such as the basal branch 1A1 cluster and AHM001/AHM011. In addition, radiocarbon dates serve as relative *termini ante quem* within the sequence model for direct ancestors and in some cases as *termini post quem* for direct descendants. For genomes of different sites, we neglect a possible coexistence of direct ancestor/descendant pairs based on the low phylodynamic threshold (e.g., AHM001 and MAL003 with a genetic distance of 1–4 SNPs). For genomes of the same site with a genetic distance of maximum 1 SNPs to the MRCA, such as the genomes of Stans (STN) and Domat/Ems (EMS), contemporaneity was assumed based on the archaeological context.

Both models passed the threshold of 60 for the model agreement value [ΔR (30, 20): Amodel 97.7, Aoverall 87.4; ΔR U(0, 50): Amodel 78.5, Aoverall 76.8] with two (uniform distribution) or one (normal distribution) radiocarbon date(s) with low agreement values. Since the normally distributed generic offset is a better representation of reality and allows for a better estimate of the posterior distribution, we chose the sequence model with a normally distributed ΔR (30, 20) as the most realistic and reliable. To determine the effect of the ΔR and kernel density plot function on individual dates, we also applied them to radiocarbon dates outside of the sequence model, and applied the kernel density model function separately, since it cannot be combined with the sequence function due to confounding effects. The used Oxcal codes can be found under Code S1–S11 with corresponding plots in Figures S45–48 and dates in Table S11. For easier comparison of different models and functions, 2σ intervals were plotted for each radiocarbon sample in Figures S49–57.

The sequence model with ΔR (30, 20) has the strongest effect on radiocarbon dates corresponding to genomes that have temporally close direct descendants or ancestors or both, since this allows for their direct integration into the sequence (Code S3, Table S11, Fig. 3 and S46). The intervals for MAN008 and OTE001 as direct descendants of the European Black Death strain (set here as prior with 1350 ± 2 years) are pushed towards the late 14th century, also suggesting a strong offset for OTE001. The dates for Arnhem (AHM001, AHM010) are constrained both by the date of European Black Death genomes and the dates of their direct descendants, such as CLP015 and CLP017 (Clopton), to 2σ intervals of less than 40 years. *Vice versa*, the dates of Arnhem narrow down the 2σ interval of CLP017 and shift the intervals for both Clopton dates to the early/mid-15th century. As a direct ancestor of the Stans cluster and the Thirty Years’ War cluster, the dates of Landsberg (LBG002, LBG005, LBG007) are confined drastically to an interval between 1460 and 1520. Here, the offset appears to be negligible and does not cause a significant change in the lower boundary of the 2σ intervals. Whereas the genomes Gdansk8, AGU007.B, AGU010.B and RUS004 appear identical, the oldest and youngest calibrated radiocarbon 2σ intervals without offset do not overlap. While solely adding an offset of ΔR (30, 20) without any further modelling causes all 2σ intervals to overlap, interval lengths also increase to 153–185 years mostly because of an additional factor of uncertainty. The KDE_Plot and KDE_Model functions outside of the sequence model are not able to narrow down the intervals significantly. The sequence model with ΔR (30, 20) generates intervals between 75 and 119 years with an overlap of all intervals between 1473 and 1510 (KDE_Plot median at 1493).

**Fig. 3:**
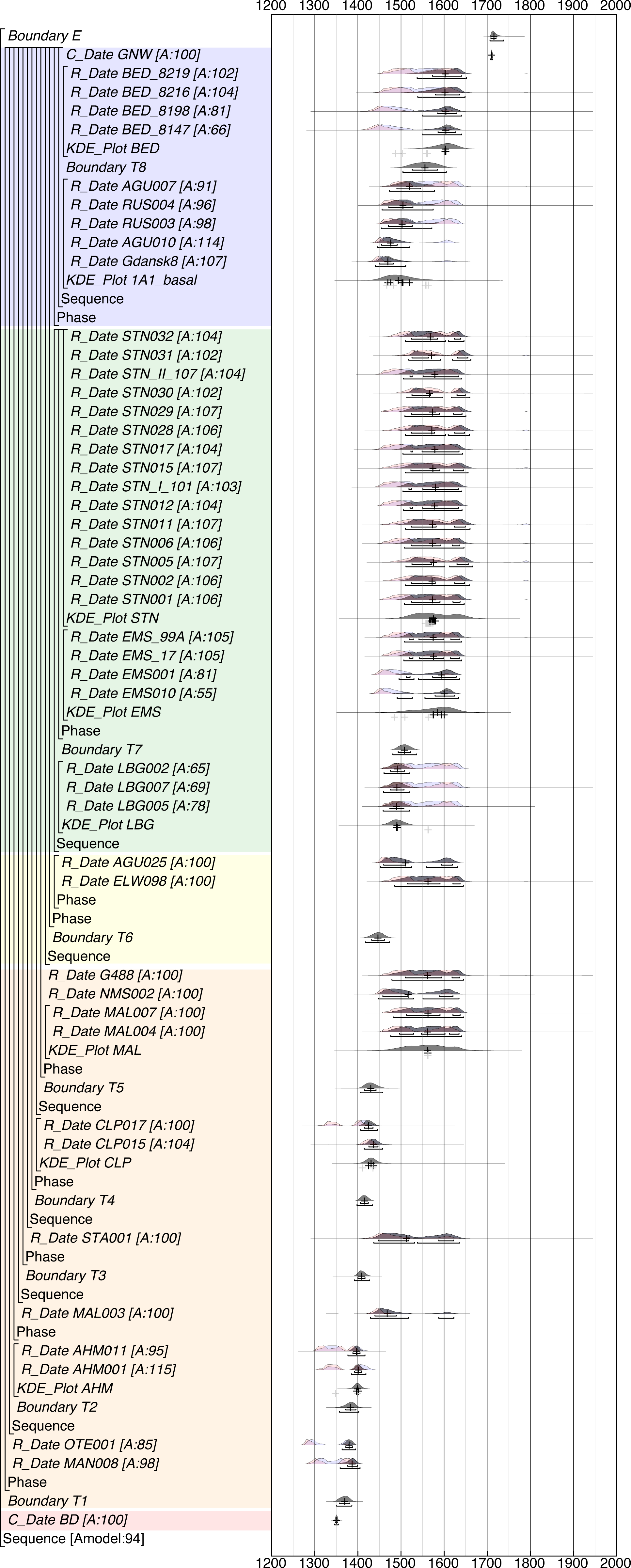
Modelled radiocarbon dates of skeletal remains of 45 individuals associated with the Second Pandemic through palaeogenomic evidence. Probability distributions without ΔR are colored in red; prior probability distributions of the model with ΔR (30, 20) are colored in blue; posterior probability distributions are shown in dark grey. Brackets correspond to 1σ (upper) and 2σ (lower) intervals; “+” indicate the medians of the respective distributions. The OxCal code can be found under Code S2; the unmodified OxCal plots in Fig. S45 (individually calibrated dates) and Fig. S46 (model); the corresponding table can be found under Table S11.

The London New Churchyard (BED) cluster gets a more precise date through the use of the aforementioned basal Branch 1A1 cluster as the ancestor in the sequence model. The 2σ intervals of up to 177 to 217 years, caused by a plateau in the calibration curve, get narrowed down to 90–115 years. Surprisingly, the sequence model with ΔR (30, 20) yields more narrow intervals than the sequence model without offset, although another factor of uncertainty is added.

### Treponemal coinfection

In addition to *Y. pestis*, we identified the individual PRN008 (Tallinn Pärnu mnt 59B) to be also infected with *Treponema pallidum*. Due to low coverage, a further characterization of the strain as *T. pallidum pallidum* (venereal syphilis) or *T. pallidum pertenue* (yaws) was not attempted, but misidentification of *T. denticola* could be excluded (see Supplementary Section 9). We present here the second known case of coinfection of plague and treponematosis, identified through ancient DNA. Giffin et al. 2020 reported a coinfection with *T. pallidum pertenue* in an individual from Vilnius (Lithuania), AGU007, dating to the second half of the 15th century^19^. *T. pallidum ssp.* is known to be notoriously difficult to detect, even in patients in the latency stage^20^. Most ancient genomes so far have been retrieved from bones with characteristic lesions, often identified as cases of congenital treponematosis. Therefore, the identification of this pathogen in the teeth of adult individuals – suspected to be due to proliferation in the bloodstream – is remarkable. *Y. pestis* might be able to create a permissive environment for other bacteria to proliferate in the host and therefore facilitate their detection in metagenomic screenings, as previously discussed for a case of a coinfection with *Y. pestis* and *Haemophilus influenzae* in an Anglo-Saxon child dating to the First Pandemic^21^.

## Discussion

The Second Plague Pandemic in Europe is the historical pandemic with the densest spatiotemporal sampling of ancient genomic data. In combination with the plethora of documentary and archaeological data on plague outbreaks as well as modern biomedical research, *Yersinia pestis* has the potential to serve as a “model organism” for historical epidemics and pandemics caused by zoonotic diseases. However, the fact that for the vast majority of ancient *Y. pestis* genomes only broad radiocarbon date intervals of more than 100 years are available, greatly reduces their usability for more advanced analyses, such as molecular dating or phylodynamics. Because of the lack of high temporal resolution, transdisciplinary approaches, combining aDNA, historical, bioarchaeological and paleoclimatological data and methods to identify potential climatic factors for the spread or decline of plague in Europe, are seriously constrained. In addition to reporting newly reconstructed *Y. pestis* genomes, we propose here a novel approach for a more precise dating of ancient plague genomes through a combination of phylogenetics, Bayesian modelling of radiocarbon dates, and documentary evidence. Since *Y. pesti*s evolves in a strictly clonal manner, its phylogeny reflects a precise relative chronological order. This was already implicitly utilised in the past, e.g., by associating (or verifying the association of) poorly dated plague burials with the European Black Death (1347–1353) through their genetic identity with the genome recovered from well-dated burials of London East-Smithfield^4, 22^. In this study, however, we apply this concept consequently to a whole set of cases over more than 300 years in form of Bayesian modelling.

So far, retrieving temporal information on the *Y. pestis* phylogeny has been done primarily by applying molecular dating, although the applicability has been questioned because of a lack of a temporal signal in some studies^4, 23, 24^. In a recent study, Eaton et al. (2023) demonstrated that the temporal signal across the entire species is unstable but increases significantly when considering only subclades^25^. However, the study was also able to show that *Y. pestis* has a high phylodynamic threshold (estimated to one substitution per 1.1 to 14.1 years). This is one key factor for the poor performance of phylogeographic analyses to trace the spread of *Y. pestis*, especially during the Second Pandemic. Molecular dating approaches are further complicated by three methodological problems:

(1) A commonly used tool for molecular dating is BEAST^26^ or BEAST2^27^, which forcefully resolves true polytomies into bifurcations while building Maximum Clade Credibility trees. This results in the creation of an artificial phylogenetic structure with ultra-short branches that are poorly supported, not only for true polytomies such as the “Big Bang” giving rise to Branch 1–4, but also for cases where a basal strain has been sampled multiple times, as in the case of the identical Black Death genomes, as discussed also by Spyrou et al. 2022^2^.

(2) While sampling dates for modern *Y. pestis* genomes have a yearly resolution, sampling dates for ancient genomes have to be inferred through documentary evidence or radiocarbon dating, resulting in broad intervals of up to 150 years. Usually, the mean date is used as a tip date for molecular dating^2, 4, 25^ instead of uniform distributions of the 2-σ intervals^28^ or the full probability distribution^29^. This results in an ‘artificial’ precision, causing too narrow (and erroneous) intervals for dates of most recent common ancestors (MRCA)^25^, which could be one explanation for discrepancies between molecular dating analyses and historical analyses regarding the emergence of pandemics^25^.

(3) Both modern and ancient *Y. pestis* genomes can suffer from false-positive SNPs, resulting in artificially long branch lengths which will interfere with molecular dating approaches. For modern genomes, the cause is poorly understood and can hardly be investigated when raw sequencing data is not made available (see Supplementary Section 3). For ancient genomes, the problem of contamination of alignments with closely related environmental bacteria is well known and can be mitigated with rigorous filtering in the case of full-UDG data^2, 30^. However, for non-UDG data, no standards have been established so far, although the respective genomes suffer more from contamination when deamination damage is accounted for through lenient mapping parameters.

In this study, we propose new standards for data curation to mitigate several of the aforementioned problems with respect to the selection of modern genomes, ancient genome reconstruction from non-UDG data, and the evaluation of phylogenetically informative SNPs.

### Phylogenetically informed radiocarbon modelling

With our novel method, we take a radically different approach to refine the temporal information embedded in the phylogeny of the Second Pandemic. By implementing the relative chronological data extracted from the phylogeny into a sequence model as commonly used, e.g., for stratigraphic archaeological data, we set our focus on more precise tip dates, instead of estimating tMRCAs and substitution rates. This approach is, however, extremely sensitive to small changes in the tree topology, which makes it necessary to filter all private SNPs in the phylogeny of the Second Pandemic and assess ambiguous positions visually (see Supplementary Section 5).

The improvement of tip dates, in turn, allows for a better historical contextualization of the data hand in hand with a phylogeographic reconstruction, as shown here in Supplementary Section 11. This approach mitigates the poor performance of phylogeographic analyses based solely on genomic data^25^. Since we used only relative chronological data, raw radiocarbon dates, and a few calendar dates for precisely dated plague genomes in our sequence model, we were able to cross-validate the results on several occasions with additional chronological information and, therefore, showcase the reliability of this method.

The poor performance of the sequence model using radiocarbon dates without offset showed that reservoir effects or HBCOs are a common problem in the context of fine-scale temporal analyses, as performed here on the Second Pandemic genomes, and that several ancient *Y. pestis* genomes might be significantly younger than previously thought. Reservoir effects have been previously discussed for the site of London New Churchyard^4^ and Cambridge Augustinian Friary^7^ and in a recent publication, Andrades Valtueña et al. were able to find strong evidence for a reservoir effect for the Neolithic plague genome I5884 of Dereivka I (Ukraine) through molecular dating^28^. For the site of Domat/Ems, dated 1629–1631, we discuss the discrepancy of individual radiocarbon dates with the historical date in Supplementary Section 11.9.

Due to the partial lack of stable isotope data for the sampled individuals and their possible dietary sources, we were unable to calculate reservoir effects on all included radiocarbon dates – neither did we have sufficient data to estimate HBCOs. Therefore, we chose to apply a moderate generic offset to all radiocarbon dates in our model with a uniform or normal distribution. Applying a normally distributed generic offset of 30 RC years (± 20), we were able to improve the performance of the model beyond the critical threshold and to reflect different possible offsets more realistically. As part of the Bayesian modelling, the broad normal distribution is flexible enough to allow for stronger or weaker offsets (posterior median ΔR between 15.5 and 43.5, see Table S11). This underlines the importance of full reporting of raw radiocarbon dates and the need for more comprehensive bioarchaeological analyses of plague burials, even as for primarily evolutionary genetic studies.

Our method performs especially well for genomes that have direct descendants and direct ancestors in temporal proximity, for example Arnhem (AHM001/AHM011) and Landsberg am Lech (LBG002.A). Therefore, further sampling of new plague genomes will also improve the precision of modelled dating intervals for previously published genomes. In other cases, the uncertainty introduced by a broad, normally distributed offset causes an expansion of the date intervals for genomes sitting on terminal branches, for which an archaeological-historical contextualization is, therefore, of paramount importance.

In the future, other chronological information could be fed into PhIRM as well, such as archaeological dates, e.g., from coinage, or contextual documentary information, such as the periods that cemeteries are known to have been in use. For this study, we refrained from this for the aforementioned cross-validation. Also, the phylodynamic threshold (recently estimated to be between 1.1 and 14.1 years per SNP)^25^ could be considered in the future to apply boundaries in such models. However, due to the possible confounding factors in molecular dating and PhIRM and common problems in molecular dating discussed above, a rigid framework needs to be established before combining molecular dating and PhIRM to avoid biases.

### Matching phylochronological and historical data

The number of currently available genomes of the Second Pandemic surpasses a hundred (Table S3, S4, ^31^), but only a minority of them can be attributed to a specific epidemic event. In parallel, more existing datasets of documentarily reported plague outbreaks have been digitized^32–34^ and historical reconstructions of plague outbreaks and waves, connecting them to hypothetical reservoirs in different regions of Eurasia, have been published by historians^3, 35–41^. However, despite these advances across fields of research, any attempts to associate ancient plague genomes or clades with documentary data remains significantly impaired by uncertainties pertaining to both palaeogenetics and history. While the first palaeogenetic studies concentrated on known plague cemeteries^1, 8^, more recent studies often included poorly dated multiple and mass burials from cemeteries with long periods of use^4, 6, 19, 22, 42, 43^ or even applied broad population-wide screenings^31^, often providing radiocarbon dates with intervals up to 150 years as the only chronological information. Published and digitized datasets of historical plague outbreaks, both regional and trans-national, often do not meet modern academic standards and suffer from uncertain retrospective diagnoses^44^. Moreover, comprehensive and detailed historical studies frequently tend to be focused on single outbreaks or waves, such as the Black Death^35, 36^, the *pestis secunda*^3^, the Great Northern War plague^45, 46^, or the Great Plague of Marseille^47–49^. Attempts to locate putative reservoirs for the Second Pandemic varied in their approaches ranging from spatial data analysis combined with climatic modelling^32^, over statistical analysis of environmental data^50^, to identifications of putative reservoir species and ecology, combined with historical data^2, 3, 37, 40^ (see also Supplementary Section 12).

In this study, we provide a fine-scale, comprehensive historical contextualization of our newly sequenced as well as previously published plague genomes (see Supplementary Section 11), building upon the improved dating of individual genomes through PhIRM and an SNP-level phylogenetic analysis (see Supplementary Section 5). A thorough analysis of documentarily reported outbreaks reconstructed from original historical sources further allows us to hypothesize about locations and timeframes for the emergence of plague lineages from a local reservoir(s). We were able to associate three genomes of the late 14th century (MAN008, COL001, OTE001) with the *pestis tertia* (1362–1379) or *pestis quarta* (1378/1379–1389) – so far the oldest post-Black Death genomes on Branch 1A. We were able to track the *pestis secunda* (1356–1366) and later outbreaks in Europe until the mid-15th century back to a putative sylvatic reservoir in the territory in modern-day Germany, which appears to have potentially moved in a gradual fashion from Hesse^3^ towards Southwest – possibly into a Frankonian or Swabian territory (Supplementary Section 11.3).

Along the basal Branch 1A, we identified with the newly reconstructed genome of AHM011 (Arnhem, the Netherlands, see Fig. 2) a common ancestor of all known Second Pandemic strains postdating the 14th century, including the genome MAL003 (Mäletjärve, Estonia) presented here (Supplementary Section 11.4). Through PhIRM and historical contextualization, we were able to narrow down the potentially associated outbreaks to 1398, 1410–1412 and 1420–1421 in the Low Countries, commencing, respectively, in 1388, 1405 and 1418–1419 in southwest Germany. Furthermore, we identified a previously unknown terminal branch along basal Branch 1A, formed by two identical genomes (CLP015 and CLP017) from Clopton, dating to the mid-15th century. For the previously published genome G701 from Riga, Latvia, we were able to show that it is most likely associated with an outbreak in the mid-15th century, and not 1550–1650 as proposed in the original publication^51^ (Supplementary Section 11.5).

Within the 15th century, we reconstructed the emergence of 3 major clades within a short period time: Branch 1A1 and Branch 1A2, emerging from the “Little Polytomy” (N13) along with two terminal branches represented by single genomes, and Branch 1A3, emerging only one SNP ancestral from node N10 and therefore shortly before the “Little Polytomy”, roughly between 1440 and 1460. At present, the phylogeography and history of Branch 1A3 are difficult to reconstruct due to the low number of available genomes (Supplementary Section 11.6): the newly reconstructed genome MAL004 (Mäletjärve, Estonia) dating to the late 16th century, NMS002 (Cambridge, England) dating to the second half of the 15th or first third of the 16th century, and G488 (Riga, Latvia), presumably associated with a local outbreak in 1602, as proposed in the original paper^51^.

The exact timeframe for the “Little Polytomy” remains unclear but can be narrowed down to c.1450 – c.1500, coinciding with the ‘Great Renaissance Drought’, a severe climatic crisis in Central Europe (Supplementary Section 11.7). Climatic anomalies have been previously discussed as contributing factors for plague pandemics^52–55^, and specifically the Justinianic Plague^56^ as well as the Black Death^37^, and consecutive outbreaks of the Second Pandemic ^6, 32^. The exact relationship between the ‘Great Renaissance Drought’ and the ‘Little Polytomy’, however, remains to be studied.

The chronological and geographic contours of Branch 1A2 and its corresponding plague outbreaks are easier to reconstruct due to the well-dated genomes of the Thirty Years’ War plague. By combining PhIRM and contextual data, we were able to narrow down the oldest representative of this branch, the genome from Landsberg am Lech, to an outbreak between 1507–1520, probably originating in a reservoir in Central Europe (Supplementary Section 11.8). This seems to have given rise to two distinct lineages circulating during the Thirty Years’ War (1618–1648), which show a remarkable microdiversity for *Y. pestis*. Their phylogeographical overlap in the Alpine region could be explained by the movement of troops importing plague from an ‘Austrian’ (Brandenburg an der Havel and Domat/Ems) and ‘French’ (Lariey and Naturno) proximate origin (Supplementary Section 11.9).

Contrary to the Thirty Years’ War plague genomes, we find a cluster of identical genomes positioned basal of Branch 1A1 – just one SNP derived from the ‘Little Polytomy’ – in Vilnius (Lithuania), Gdansk (Poland) and Stankeyevo (Russia). This is the first observation of identical genomes in different geographical locations after the Black Death, presumably indicating a fast-spreading epidemic event in the last third of the 15th century or the first decade of the 16th century. While the palaeogenomic evidence is so far limited to the Baltic region, the historical context suggests that the respective genomes are related to a widespread epidemic also affecting other parts of Europe, although the provenance of another identical genome (Rostov2039) is questionable (Supplementary Section 11.10).

Before its next known occurrence in London (New Churchyard) between c. 1550–1650 (Supplementary Section 11.12), Branch 1A1 seems to have retracted from Europe, establishing a putative reservoir in the Ottoman Empire^41^, from where it spread repeatedly at least until the 18th century (Supplementary Section 11.11). This was concurrent with a large genomic deletion and an accelerated substitution rate ^4^, possibly caused by the introduction of Branch 1A1 to a new ecological niche, although a possible connection between these three observations remains purely hypothetical for now. Younger genomes of this branch, originating in a putative Ottoman reservoir, are either imported directly from the Ottoman territory, via the Black Sea or Caucasus (Azov, Rostov-on-Don, Maist, Supplementary Section 11.12, 11.14), via the Mediterranean (the Great Plague of Marseille genome, Supplementary Section 11.13, 11.14), or via South-Eastern and Eastern Europe into the Baltic (the Great Northern War plague genomes from Pestbacken and Tallinn, Supplementary Section 11.13, 11.14), as has been discussed previously ^6^.

## Conclusion

Through the reconstruction of 11 new high-quality *Y. pestis* genomes (>4.4-fold mean coverage), we were able to further elucidate the history of the Second Pandemic in Europe from the 14th to 18th century and identify so-far unknown lineages emerging along basal Branch 1A (AHM011–MAL003 and CLP015/CLP017). We found additional evidence for a remarkable diversity of *Y. pestis* during the Thirty Years’ War plague and could identify a widespread epidemic at some point between c.1470 and c.1510 through identical genomes from around the Baltic region that most likely affected other parts of Europe as well.

In addition, we were able to show that the integration of phylogenetic information into radiocarbon modelling results in more accurate and more precise date intervals for ancient plague genomes, which could be equally applicable to other clonally evolving organisms. Together with a fine-scale historical contextualization, this offers new avenues for the reconstruction of the phylochronology of historical plague pandemics that, so far, have not been accessible with phylodynamic methods. However, both reservoir effects and HBCO were taken into account only as generic offsets, currently limiting the power of this new methodological approach. Therefore, future palaeogenomic plague studies should not only include comprehensive reports of raw radiocarbon data but should ideally be performed in a broader bioarchaeological framework including the analysis of stable isotope data for diet reconstruction. While the models presented in this study only included radiocarbon dates, selected calendar dates and relative chronological information extracted from the phylogeny of the Second Pandemic as a proof of concept, future studies could refine the models by including contextual data, such as the occupation periods of associated cemeteries or dates of potentially associated plague outbreaks known from historical sources. Due to the threat of confounding effects, we refrained here from a combination of modelled posterior distributions for radiocarbon dates with molecular dating – a problem that could be potentially solved by integrating both into a single Bayesian model.

## Material and Methods

### Laboratory work

In total, 117 individuals from 11 sites were sampled for this study with one tooth sample per individual. All lab work was performed in the ancient DNA facility of the Institute of Genomics, University of Tartu, following the general guidelines^57^. Samples were processed according to the “Sampling of tooth roots for ancient DNA” protocol^58^, followed by “Decontamination of tooth roots/petrous bone cores for ancient DNA extraction”^59^, “Ancient DNA extraction (chunk samples/high volume)”^60^, and “Ancient DNA extract purification (chunk samples/high volume)”^61^. For pathogen screening we prepared non-UDG treated double-stranded DNA libraries^62, 63^. For capture, we prepared UDG-treated libraries of the same extracts as the screening libraries^63, 64^, with the exception of MAL003, MAL004 and MAL007, for which the original non-UDG-treated library was used. Non-UDG-treated libraries for screening were sequenced on a NextSeq500 (75 bp, single-end) to around 20 M reads. Captured libraries were sequenced with 150 bp paired-end kits with facultative resequencing depending on coverage and library complexity.

We enriched UDG-treated and non-UDG-treated libraries using a custom *Y. pestis/Y. pseudotuberculosis* myBaits target enrichment kit from Daicel Arbor Biosciences (v4). The capture design covers the *Y. pestis* CO92 reference genome (including all plasmids) and the *Y. pseudotuberculosis* reference genome (NC_006155.1). We followed the myBaits v4 protocol with one major exception: 2.75 µl hybridization baits were used for each reaction instead of 5.5 µl. Capture products were amplified using 2X KAPA HiFi HotStart ReadyMix DNA Polymerase and primers IS5 and IS6 ^65^. A second round of capture was performed for samples AHM001, LHP001, MOI001, PRN003 and PRN009 to reach higher concentrations of target DNA for efficient sequencing.

### Pathogen screening

For the pathogen screening, the raw sequencing data were quality-checked using FastQC and data points across the analysis were compared using MultiQC. Datasets were trimmed and quality filtered using cutadapt ^66^(-m 30 --nextseq-trim=20 --times 3 -e 0.2 --trim-n) and deduplicated using clumpify.sh ^67^. We computed the metagenomic profiles for our sample using KrakenUniq ^9^ following the workflow and database described in Guellil et al. 2022^21^.

### Modern reference data

Modern *Y. pestis* genomes were retrieved as raw sequencing data (FASTQ) if possible. For genomes only available as a FASTA file, artificial reads were created with a bedtools2 (2.27.1) with a length of 100 bases and a tiling of 2 bases. Original sequencing reads and artificial reads were processed using nf-core/eager (v.2.2.2). Reads were mapped with bwa-aln (v0.7.17, -n 0.1, -l 32), mapped reads were filtered for mapping quality 37, duplicates were removed with MarkDuplicates (v2.22.9) and GATK Unified Genotyper (v3.5.0 using EMIT_ALL_SITES) was used for genotyping. The selection of modern reference sequences is described under [selection of modern genomes].

### Ancient genomic data

We retrieved previously published ancient *Y. pestis* genomes as raw sequencing data (FASTQ) and processed them using nf-core/eager (v2.3.0). After adapter removal with standard settings (AdapterRemoval v2.3.1), we mapped reads of UDG-treated libraries with bwa-aln (v0.7.17, -n 0.1, -l 32). After filtering for mapping quality 37 and deduplication using MarkDuplicates, genotyping was performed with GATK Unified Genotyper (v3.5.0 using EMIT_ALL_SITES).

For libraries without UDG treatment, we tested several, previously published or suggested strategies, since non-UDG libraries regularly show artificially long terminal branches in phylogenetic analyses dues to DNA damage and/or mapping parameters that allow for a higher edit distance to accommodate for DNA damage, causing reads originating from closely related environmental bacteria to contaminate the mapping. We compared both the SNP statistics and the phylogenetic positioning of genomes using the following treatments using bwa-aln (v0.7.17):

1. Stringent mapping with -n 0.1, -l 32
2. Standard mapping with -n 0.04, -l 1000 (seed disabled)
3. Loose mapping with -n 0.01, -l 16
4. Clipping 3 bases from both sides on FASTQ files prior to with stringent mapping with -n 0.1, -l 32
5. Soft clipping 3 bases from both sides on BAM files after loose mapping with -n 0.01, -l 16
6. Loose mapping with -n 0.01, -l 16 followed by damage rescaling using MapDamage (v2.2.1)
7. Loose mapping with -n 0.01, -l 16 followed by damage rescaling using MapDamage (v2.2.1) and stringent remapping with -n 0.1, -l 32

As shown in Supplementary Section 4, the last method performed best in removing false-positive SNPs causing artificially long terminal branches, and was therefore used for all following analyses.

For the genome BSK001.A, reconstructed both from partially UDG-treated double-stranded libraries and a single-stranded library without UDG treatment, the publicly available BAM file was used for downstream analyses after remapping with bwa-aln (v0.7.17, -n 0.01, -l 16) und genotyping as described above using nf-core/eager (v2.3.0).

### Phylogenetic analyses

The selection of modern genomes as a reference set is described in Supplementary Section 1. In total, we used 208 modern *Y. pestis* genomes and the *Y. pseudotuberculosis* reference genome as well as 44 previously published ancient genomes (Table S3).

MultiVCFAnalyzer (v0.85.2) was used to produce SNP alignments using a minimum genotyping quality of 30, a minimal coverage of 3-fold and a minimal allele frequency of 0.9 for homozygous calls. The genome of Y. pseudotuberculosis IP32953 was treated as an outgroup. Regions previously identified as non-core regions, containing repetitive elements or coding for tRNAs, rRNAs or tmRNAs were excluded. Maximum likelihood trees were calculated with IQTree v2.1.2 with bootstrapping (1000 replications) and GTR+F+ASC+R2 as a substitution model.

### Radiocarbon modeling

For the radiocarbon modeling, we collected all available information from previous publications on Second Pandemic *Y. pestis* genomes and were able to retrieve previously unpublished raw data (see Table S10). This includes quality criteria, sampled skeletal element and stable isotope ratios (δ^13^C, δ^15^N), if available, and archaeological/historical datings included in the publications. In addition, we produced 21 new radiocarbon dates for sites included in this study, including stable isotope ratios (δ^13^C, δ^15^N).

All calibrations and models on the radiocarbon dates were performed with OxCal 4.4 and the IntCal 20 calibration curve. To make use of the relative temporal information embedded in the rooted phylogenetic tree of Second Pandemic *Y. pestis* genomes for the recalibration of radiocarbon dates, we developed a novel approach we term “phylogenetically informed radiocarbon modeling” (PhIRM). PhIRM integrates phylogenetic information as relative temporal information in the form of a cascade of nested phases and sequences with boundaries. PhIRM makes use of Markov chain Monte Carlo (MCMC) analysis as part of the multi-parameter Bayesian analysis performed with OxCal 4.4.

Every clade is treated as a phase within a sequence with a lower boundary. The lower boundary represents the earliest possible emergence of the respective clade based on the oldest radiocarbon date within the phase, without considering its branch length. This sequence is then nested within another sequence by integrating a genome or clade that shares the next most recent common ancestor with the first clade. Only genomes that represent common ancestors, i.e., occupy positions directly on nodes in the phylogenetic tree, are directly integrated into sequences: descending genomes must date younger, genomes that are directly ancestral must be older. Genomes that directly represent common ancestors are the ‘Black Death cluster’ genomes NAB003.B, ESFpool, Barcelona and BEN001; AHM011/AHM001; LBG002.A; the ‘Eastern Europe cluster’ genomes Rostov2039, Gdansk8, AGU007.B, AGU010.B, RUS004; London New Churchyard genomes BED024.A, BED028.A, BED030.A, BED034.A. The genome PRN001 representing the Great Northern War plague was integrated as terminal point on Branch 1A1 as a direct descendant of the London New Churchyard genomes. The dates corresponding to these genomes are the only fixed constraints in the model. For two contexts, we included absolute calendar dates into the model instead of radiocarbon dates: the ‘Black Death cluster’ was assigned the calendar date 1350 (± 2 years) and the Great Northern War Plague (PRN001) was assigned 1710 (± 1 year). For identical genomes we assume that they represent the same epidemic event, however without offering a constraint of its duration. We refrained from using R_Combine as done previously^19^, since this function should only be used when the radiocarbon dates come from the same source, i.e. bone or tooth, and does not allow for offsets. This is problematic because it neglects differences in the marine or freshwater reservoir effect of the dated individuals, and differences in the human bone collagen offset. In addition, for the ‘Eastern Europe cluster’, samples were dated in different radiocarbon laboratories, adding the possibility of laboratory-specific offsets. We therefore use Kernel Density Estimation (i.e., the function KDE_Plot) to combine dates corresponding to identical genomes^14^.

Since the necessary data to estimate the exact HBCO and reservoir effects were not available for the majority of radiocarbon dates, we tested three approaches: First, the model was run without any correction; second, we assumed an offset (ΔR) of 30 radiocarbon years and uncertainty of 20 radiocarbon years for all samples; third, we allowed the MCMC analysis to take an offset (ΔR) between 0 and 50 radiocarbon years (U(0,50)).

## Contributions

Designed the study: MK, MG, PS, HV, AK, JER, KT, CLS

Performed laboratory work: MK, MG, LS, KI, HN, AS, CLS

Performed data analyses: MK, MG

Provided archaeological, anthropological and historical analyses and data: PS, MM, HV, AK,

CCessford, SAI, JER,CCooper, CvP, MS, TR, WAB

Advised radiocarbon modelling: CCessford, SS

Provided raw radiocarbon dates: DW, SL

Provided funding: JER, TR, MM, TK, KT

Wrote the paper with contributions of all authors: MK, PS

## Acknowledgements

We thank Dr. Trish Biers (The Duckworth Collection, Cambridge, England) for providing samples of individuals of All Saints by the Castle (Cambridge, England) and Clopton (Cambridgeshire, England) as well as Aleksandr Mikhailov and Viktoria Derkach for providing samples of individuals of Stankeyevo (Russia). We acknowledge the excavators of All Saints by the Castle (Cambridge, England) Paul Craddock and Vince Gregory as well as the Arnhem municipality of RAAP for their support on the research of the site of Arnhem. We thank Johannes Krause for providing unpublished raw radiocarbon dates.

This study was supported by the Wellcome Trust Collaborative Grant 200368/Z/15/Z (CCessford, SAI, JR, TK, CLS), the Estonian Research Council grants PRG1027 (MK, MG, LS, AK, KT) and PRG1931 (HV); the European Union through the European Regional Development Fund project no. 2014-2020.4.01.16-0030 (MK, MG, MM, CLS) and the European Regional Development Fund project no. 2014-2020.4.01.15-0012 (MM).

## Competing interests

The authors declare no competing interests.

## Data availability

The sequencing data will be made available through the European Nucleotide Archive under the project accession number PRJEB64346.

## Notes

### Competing Interest Statement

The authors have declared no competing interest.

## References

1. Bos, K.I., Schuenemann, V.J., Golding, G.B., Burbano, H.A., Waglechner, N., Coombes, B.K., McPhee, J.B., DeWitte, S.N., Meyer, M., Schmedes, S., et al. (2011). A draft genome of Yersinia pestis from victims of the Black Death. Nature 478, 506–510. 10.1038/nature10549.

2. Spyrou, M.A., Musralina, L., Gnecchi Ruscone, G.A., Kocher, A., Borbone, P.-G., Khartanovich, V.I., Buzhilova, A., Djansugurova, L., Bos, K.I., Kühnert, D., et al. (2022). The source of the Black Death in fourteenth-century central Eurasia. Nature, 1–7. 10.1038/s41586-022-04800-3.

3. Slavin, P. (2021). Out of the West: Formation of a Permanent Plague Reservoir in South-Central Germany (1349–1356) and its Implications. Past Present 252, 3–51. 10.1093/pastj/gtaa028.

4. Spyrou, M.A., Keller, M., Tukhbatova, R.I., Scheib, C.L., Nelson, E.A., Andrades Valtueña, A., Neumann, G.U., Walker, D., Alterauge, A., Carty, N., et al. (2019). Phylogeography of the second plague pandemic revealed through analysis of historical Yersinia pestis genomes. Nat. Commun. 10, 4470. 10.1038/s41467-019-12154-0.

5. Seguin-Orlando, A., Costedoat, C., Der Sarkissian, C., Tzortzis, S., Kamel, C., Telmon, N., Dalén, L., Thèves, C., Signoli, M., and Orlando, L. (2021). No particular genomic features underpin the dramatic economic consequences of 17th century plague epidemics in Italy. iScience 24, 102383. 10.1016/j.isci.2021.102383.

6. Guellil, M., Kersten, O., Namouchi, A., Luciani, S., Marota, I., Arcini, C.A., Iregren, E., Lindemann, R.A., Warfvinge, G., Bakanidze, L., et al. (2020). A genomic and historical synthesis of plague in 18th century Eurasia. Proc. Natl. Acad. Sci. U. S. A. 10.1073/pnas.2009677117.

7. Cessford, C., Scheib, C.L., Guellil, M., Keller, M., Alexander, C., Inskip, S.A., and Robb, J.E. (2021). Beyond Plague Pits: Using Genetics to Identify Responses to Plague in Medieval Cambridgeshire. European Journal of Archaeology, 1–23. 10.1017/eaa.2021.19.

8. Bos, K.I., Herbig, A., Sahl, J., Waglechner, N., Fourment, M., Forrest, S.A., Klunk, J., Schuenemann, V.J., Poinar, D., Kuch, M., et al. (2016). Eighteenth century Yersinia pestis genomes reveal the long-term persistence of an historical plague focus. Elife 5, e12994. 10.7554/eLife.12994.

9. Breitwieser, F.P., Baker, D.N., and Salzberg, S.L. (2018). KrakenUniq: confident and fast metagenomics classification using unique k-mer counts. Genome Biol. 19, 198. 10.1186/s13059-018-1568-0.

10. Jónsson, H., Ginolhac, A., Schubert, M., Johnson, P.L.F., and Orlando, L. (2013). mapDamage2.0: fast approximate Bayesian estimates of ancient DNA damage parameters. Bioinformatics 29, 1682–1684. 10.1093/bioinformatics/btt193.

11. Li, H., and Durbin, R. (2010). Fast and accurate long-read alignment with Burrows-Wheeler transform. Bioinformatics 26, 589–595. 10.1093/bioinformatics/btp698.

12. Bronk Ramsey, C. (2009). Bayesian Analysis of Radiocarbon Dates. Radiocarbon 51, 337–360. 10.1017/S0033822200033865.

13. Reimer, P.J., Austin, W.E.N., Bard, E., Bayliss, A., Blackwell, P.G., Ramsey, C.B., Butzin, M., Cheng, H., Lawrence Edwards, R., Friedrich, M., et al. (2020). The IntCal20 Northern Hemisphere Radiocarbon Age Calibration Curve (0–55 cal kBP). Radiocarbon 62, 725– 757. 10.1017/RDC.2020.41.

14. Ramsey, C.B. (2017). Methods for Summarizing Radiocarbon Datasets. Radiocarbon 59, 1809–1833. 10.1017/RDC.2017.108.

15. Barta, P., and Štolc, S. (2007). HBCO Correction: Its Impact on Archaeological Absolute Dating. Radiocarbon 49, 465–472. 10.1017/s0033822200042399.

16. Ubelaker, D.H., Plens, C.R., Soriano, E.P., Diniz, M.V., de Almeida Junior, E., Junior, E.D., Júnior, L.F., and Machado, C.E.P. (2022). Lag time of modern bomb-pulse radiocarbon in human bone tissues: New data from Brazil. Forensic Sci. Int. 331, 111143. 10.1016/j.forsciint.2021.111143.

17. Ascough, P., Cook, G., and Dugmore, A. (2005). Methodological approaches to determining the marine radiocarbon reservoir effect. Progress in Physical Geography: Earth and Environment 29, 532–547. 10.1191/0309133305pp461ra.

18. Philippsen, B. (2013). The freshwater reservoir effect in radiocarbon dating. Heritage Science 1, 1–19. 10.1186/2050-7445-1-24.

19. Giffin, K., Lankapalli, A.K., Sabin, S., Spyrou, M.A., Posth, C., Kozakaitė, J., Friedrich, R., Miliauskienė, Ž., Jankauskas, R., Herbig, A., et al. (2020). A treponemal genome from an historic plague victim supports a recent emergence of yaws and its presence in 15th century Europe. Sci. Rep. 10, 9499. 10.1038/s41598-020-66012-x.

20. Radolf, J.D., Deka, R.K., Anand, A., Šmajs, D., Norgard, M.V., and Yang, X.F. (2016). Treponema pallidum, the syphilis spirochete: making a living as a stealth pathogen. Nat. Rev. Microbiol. 14, 744–759. 10.1038/nrmicro.2016.141.

21. Guellil, M., Keller, M., Dittmar, J.M., Inskip, S.A., Cessford, C., Solnik, A., Kivisild, T., Metspalu, M., Robb, J.E., and Scheib, C.L. (2022). An invasive Haemophilus influenzae serotype b infection in an Anglo-Saxon plague victim. Genome Biol. 23, 22. 10.1186/s13059-021-02580-z.

22. Namouchi, A., Guellil, M., Kersten, O., Hänsch, S., Ottoni, C., Schmid, B.V., Pacciani, E., Quaglia, L., Vermunt, M., Bauer, E.L., et al. (2018). Integrative approach using *Yersinia pestis* genomes to revisit the historical landscape of plague during the Medieval Period. Proc. Natl. Acad. Sci. U. S. A. 115, E11790–E11797. 10.1073/pnas.1812865115.

23. Wagner, D.M., Klunk, J., Harbeck, M., Devault, A., Waglechner, N., Sahl, J.W., Enk, J., Birdsell, D.N., Kuch, M., Lumibao, C., et al. (2014). Yersinia pestis and the plague of Justinian 541-543 AD: a genomic analysis. Lancet Infect. Dis. 14, 319–326. 10.1016/S1473-3099(13)70323-2.

24. Cui, Y., Yu, C., Yan, Y., Li, D., Li, Y., Jombart, T., Weinert, L.A., Wang, Z., Guo, Z., Xu, L., et al. (2013). Historical variations in mutation rate in an epidemic pathogen, Yersinia pestis. Proc. Natl. Acad. Sci. U. S. A. 110, 577–582. 10.1073/pnas.1205750110.

25. Eaton, K., Featherstone, L., Duchene, S., Carmichael, A.G., Varlık, N., Golding, G.B., Holmes, E.C., and Poinar, H.N. (2023). Plagued by a cryptic clock: insight and issues from the global phylogeny of Yersinia pestis. Commun Biol 6, 23. 10.1038/s42003-022-04394-6.

26. Drummond, A.J., and Rambaut, A. (2007). BEAST: Bayesian evolutionary analysis by sampling trees. BMC Evol. Biol. 8. 10.1186/1471-2148-7-214.

27. Bouckaert, R., Heled, J., Kühnert, D., Vaughan, T., Wu, C.-H., Xie, D., Suchard, M.A., Rambaut, A., and Drummond, A.J. (2014). BEAST 2: A Software Platform for Bayesian Evolutionary Analysis. PLoS Comput. Biol. 10, e1003537. 10.1371/journal.pcbi.1003537.

28. Andrades Valtueña, A., Neumann, G.U., Spyrou, M.A., Musralina, L., Aron, F., Beisenov, A., Belinskiy, A.B., Bos, K.I., Buzhilova, A., Conrad, M., et al. (2022). Stone Age Yersinia pestis genomes shed light on the early evolution, diversity, and ecology of plague. Proc. Natl. Acad. Sci. U. S. A. 119, e2116722119. 10.1073/pnas.2116722119.

29. Molak, M., Suchard, M.A., Ho, S.Y.W., Beilman, D.W., and Shapiro, B. (2015). Empirical calibrated radiocarbon sampler: a tool for incorporating radiocarbon-date and calibration error into Bayesian phylogenetic analyses of ancient DNA. Mol. Ecol. Resour. 15, 81–86. 10.1111/1755-0998.12295.

30. Keller, M., Spyrou, M.A., Scheib, C.L., Neumann, G.U., Kröpelin, A., Haas-Gebhard, B., Päffgen, B., Haberstroh, J., Ribera I Lacomba, A., Raynaud, C., et al. (2019). Ancient Yersinia pestis genomes from across Western Europe reveal early diversification during the First Pandemic (541-750). Proc. Natl. Acad. Sci. U. S. A. 116, 12363–12372. 10.1073/pnas.1820447116.

31. Eaton, K., Sidhu, R.K., Klunk, J., Gamble, J.A., Boldsen, J.L., Carmichael, A.G., Varlık, N., Duchene, S., Featherstone, L., Grimes, V., et al. (2023). Emergence, continuity, and evolution of Yersinia pestis throughout medieval and early modern Denmark. Curr. Biol. 0. 10.1016/j.cub.2023.01.064.

32. Schmid, B.V., Büntgen, U., Easterday, W.R., Ginzler, C., Walløe, L., Bramanti, B., and Stenseth, N.C. (2015). Climate-driven introduction of the Black Death and successive plague reintroductions into Europe. Proc. Natl. Acad. Sci. U. S. A. 112, 3020–3025. 10.1073/pnas.1412887112.

33. Krauer, F., and Schmid, B.V. (2021). Mapping the plague through natural language processing. medRxiv, 2021.04.27.21256212. 10.1101/2021.04.27.21256212.

34. Büntgen, U., Ginzler, C., Jan, E., Tegel, W., and McMichael, A.J. (2012). Digitizing historical plague. Clin. Infect. Dis. 55, 1586–1588. 10.1093/cid/cis723.

35. Slavin, P. (2019). Death by the Lake: Mortality Crisis in Early Fourteenth-Century Central Asia. J. Interdiscip. Hist. 50, 59–90. 10.1162/jinh_a_01376.

36. Green, M.H. (2020). The Four Black Deaths. Am. Hist. Rev. 125, 1601–1631. 10.1093/ahr/rhaa511.

37. Slavin, P. (2023). The Birth of the Black Death: Biology, Climate, Environment, and the Beginnings of the Second Plague Pandemic in Early Fourteenth-Century Central Asia. Environ. Hist. Durh. N. C., 000–000. 10.1086/723955.

38. Varlık, N. (2022). The Rise and Fall of a Historical Plague Reservoir. The Case of Ottoman Anatolia. In Disease and the Environment in the Medieval and Early Modern Worlds, L. Jones, ed. (Routledge), pp. 159–183.

39. Varlik, N. (2015). Plague and Empire in the Early Modern Mediterranean World: The Ottoman Experience, 1347–1600 (Cambridge University Press) 10.1017/CBO9781139004046.

40. Carmichael, A.G. (2014). Plague Persistence in Western Europe: A Hypothesis. In, pp. 157–192.

41. Slavin, P. (2022). Reply: Out of the West — and neither East, nor North, nor South. Past Present 256, 325–360. 10.1093/pastj/gtac026.

42. Spyrou, M.A., Tukhbatova, R.I., Feldman, M., Drath, J., Kacki, S., Beltrán de Heredia, J., Arnold, S., Sitdikov, A.G., Castex, D., Wahl, J., et al. (2016). Historical Y. pestis Genomes Reveal the European Black Death as the Source of Ancient and Modern Plague Pandemics. Cell Host Microbe 19, 874–881. 10.1016/j.chom.2016.05.012.

43. Morozova, I., Kasianov, A., Bruskin, S., Neukamm, J., Molak, M., Batieva, E., Pudło, A., Rühli, F.J., and Schuenemann, V.J. (2020). New ancient Eastern European Yersinia pestis genomes illuminate the dispersal of plague in Europe. Philos. Trans. R. Soc. Lond. B Biol. Sci. 375, 20190569. 10.1098/rstb.2019.0569.

44. Roosen, J., and Curtis, D.R. (2018). Dangers of noncritical use of historical plague data. Emerg. Infect. Dis. 24, 103–110. 10.3201/eid2401.170477.

45. Zapnik, J. (2007). Pest und Krieg im Ostseeraum: Der “Schwarze Tod” in Stralsund während des Großen Nordischen Krieges (1700–1721) (Verlag Dr. Kovač).

46. Frandsen, K.-E. (2010). The Last Plague in the Baltic Region, 1709–1713 (Museum Tusculanum Press).

47. Ermus, C. (2022). The Great Plague Scare of 1720: Disaster and Diplomacy in the Eighteenth-Century Atlantic World (Cambridge University Press) 10.1017/9781108784733.

48. Devaux, C.A. (2013). Small oversights that led to the Great Plague of Marseille (1720– 1723): Lessons from the past. Infect. Genet. Evol. 14, 169–185.

49. Signoli, M. (2022). History of the plague of 1720-1722, in Marseille. La Presse Médicale 51, 104138. 10.1016/j.lpm.2022.104138.

50. Stenseth, N.C., Tao, Y., Zhang, C., Bramanti, B., Büntgen, U., Cong, X., Cui, Y., Zhou, H., Dawson, L.A., Mooney, S.J., et al. (2022). No evidence for persistent natural plague reservoirs in historical and modern Europe. Proc. Natl. Acad. Sci. U. S. A. 119, e2209816119. 10.1073/pnas.2209816119.

51. Susat, J., Bonczarowska, J.H., Pētersone-Gordina, E., Immel, A., Nebel, A., Gerhards, G., and Krause-Kyora, B. (2020). Yersinia pestis strains from Latvia show depletion of the pla virulence gene at the end of the second plague pandemic. Sci. Rep. 10, 14628. 10.1038/s41598-020-71530-9.

52. Campbell, B.M.S. (2016). The Great Transition: Climate, Disease and Society in the Late-Medieval World (Cambridge University Press) 10.1017/CBO9781139031110.

53. Green, M.H. (2018). Climate and Disease in Medieval Eurasia. In Oxford Research Encyclopedia of Asian History (Oxford University Press). 10.1093/acrefore/9780190277727.013.6.

54. Fell, H.G., Baldini, J.U.L., Dodds, B., and Sharples, G.J. (2020). Volcanism and global plague pandemics: Towards an interdisciplinary synthesis. J. Hist. Geogr. 70, 36–46. 10.1016/j.jhg.2020.10.001.

55. Luterbacher, J., Newfield, T.P., Xoplaki, E., Nowatzki, E., Luther, N., Zhang, M., and Khelifi, N. (2020). Past pandemics and climate variability across the Mediterranean. Euro-Mediterranean Journal for Environmental Integration 5, 46. 10.1007/s41207-020-00197-5.

56. Büntgen, U., Myglan, V.S., Ljungqvist, F.C., McCormick, M., Di Cosmo, N., Sigl, M., Jungclaus, J., Wagner, S., Krusic, P.J., Esper, J., et al. (2016). Cooling and societal change during the Late Antique Little Ice Age from 536 to around 660 AD. Nat. Geosci. 9, 231–236. 10.1038/ngeo2652.

57. Keller, M., and Scheib, C.L. (2023). Ancient DNA protocol collection – University of Tartu, Institute of Genomics v1. 10.17504/protocols.io.n92ldzbd8v5b/v1.

58. Keller, M., and Scheib, C.L. (2023). Sampling of tooth roots for ancient DNA v1. 10.17504/protocols.io.6qpvrdr83gmk/v1.

59. Keller, M., and Scheib, C.L. (2023). Decontamination of tooth roots/petrous bone cores for ancient DNA extraction v1. 10.17504/protocols.io.eq2lynp4qvx9/v1.

60. Keller, M., and Scheib, C.L. (2023). Ancient DNA extraction (chunk samples/high volume) v1. 10.17504/protocols.io.n92ldzbpxv5b/v1.

61. Keller, M., and Scheib, C.L. (2023). Ancient DNA extract purification (chunk samples/high volume) v1. 10.17504/protocols.io.j8nlkwje6l5r/v1.

62. Keller, M., and Scheib, C.L. (2023). Library preparation (dsDNA single indexing, non-UDG, no split) v1. 10.17504/protocols.io.n92ldpjexl5b/v1.

63. Keller, M., Scheib, C.L., and Bonucci, B. (2023). Indexing PCR and purification of dsDNA libraries v1. 10.17504/protocols.io.rm7vzboy4vx1/v1.

64. Keller, M., and Scheib, C.L. (2023). Library preparation (dsDNA single indexing, full-UDG, no split) v1. 10.17504/protocols.io.yxmvm2qj6g3p/v1.

65. Meyer, M., and Kircher, M. (2010). Illumina sequencing library preparation for highly multiplexed target capture and sequencing. Cold Spring Harb. Protoc. 2010, db.prot5448. 10.1101/pdb.prot5448.

66. Martin, M. (2011). Cutadapt removes adapter sequences from high-throughput sequencing reads. EMBnet.journal 17, 10–12. 10.14806/ej.17.1.200.

67. Bushnell, B. (2014). BBMap: A fast, accurate, splice-aware aligner (Lawrence Berkeley National Lab. (LBNL), Berkeley, CA (United States)).

