## Supplementary Text for "A Refined Phylochronology of the Second Plague Pandemic in Western Eurasia"

Fig. S27: IGV Screenshots of 'N' positions manually checked for the schematic tree Figs. 2A and S11 (1). 43
Fig. S28: IGV Screenshots of 'N' positions manually checked for the schematic tree Figs. 2A and S11 (2). 44
Fig. S29: IGV Screenshots of 'N' positions manually checked for the schematic tree Figs. 2A and S11 (3). 45
Fig. S30: IGV Screenshots of 'N' positions manually checked for the schematic tree Figs. 2A and S11 (4). 46

**6 Heterozygosity plots 47**

Fig. S31: Heterozygosity plots for MAL003 and MAL004. 47
Fig. S32: Heterozygosity plots for Azov38, Ber37, Ber45, and CHE1. 47
Fig. S33: Heterozygosity plots for COL001, Gdansk8, PEB10, Rostov2033, Rostov2039, and SPN19. 48
Fig. S34: Heterozygosity plots for AHM011, BEN001, CLP015, CLP017, EMS001, EMS010, OTE001, PRN001, and RUS004. 49
Fig. S35: Heterozygosity plots for AGU007.B, AGU010.B, AGU025.B, Barcelona, BED024.A, BED028.A, BED030.A, BED034.A, Bolgar, BRA001.A, BSK001.A, ELW098.A, and ESFpool, G488, and G70. 50
Fig. S36: Heterozygosity plots for LAI009.A, LAR11t, LBG002.A, MAN008.B, NAB003.B, NMS002.A, OBS017, OBS110, OBS116, OBS124, OBS137, STA001.A, STN002.A, STN007.A, STN008.A, STN013.A, STN014.A, STN019.A, STN020.A, and STN021.A. 51

**7 Newly sequenced low coverage genomes 52**

Fig. S37: Phylogenetic tree including newly sequenced low coverage genomes (>1x) 54
Fig. S38: IGV screenshots of MAL004 and MAL007 for positions leading from N06–N11 (1) 55
Fig. S39: IGV screenshots of MAL004 and MAL007 for positions leading from N06–N11 (2) 56
Fig. S40: IGV screenshot of RUS003 and RUS004 for the SNP defining the basal Branch 1A1 cluster 57
Fig. S41: IGV screenshot of EMS001, EMS002, EMS003 and EMS010 for the private SNPs of EMS001 and EMS010. 57
Fig. S42: IGV screenshot of PRN001, PRN003, PRN004, PRN005, PRN008, PRN009 and PRN010 for two positions exclusively shared among low coverage PRN genomes. 58

**8 Deletion analysis and functional analysis of newly identified SNPs 59**

Fig. S43: Heatmap showing breadth of coverage over virulence-associated genes on the chromosome and the plasmids pCD1, pMT1 and pPCP1 (yellow: no coverage; dark blue: gene fully covered). 60

***9* Coinfection with *Yersinia pestis* and *Treponema pallidum* 61**

Fig. S44: Mapping plots of PRN008 for *T. pallidum pallidum, T. pallidum pertenue* and *T. denticola.* 62

**10 Radiocarbon Modelling 63**

Code S1: Oxcal code for individually calibrated dates 63
Code S2: Oxcal code for individually calibrated dates with ΔR (30, 20) 64
Code S3: Oxcal sequence model using KDE_Plots and ΔR (30, 20) 65
Code S4: Oxcal sequence model using KDE_Plots and ΔR U(0, 50) 68
Code S5: Oxcal sequence model using KDE_Plots without ΔR 71
Code S6: Oxcal code for KDE_Plots using ΔR (30, 20) 74
Code S7: Oxcal code for KDE_Plots using ΔR U(0, 50) 75
Code S8: Oxcal code for KDE_Plots without ΔR 77
Code S9: Oxcal code for KDE_Models using ΔR (30, 20) 78
Code S10: Oxcal code for KDE_Models using ΔR U(0, 50) 80
Code S11: Oxcal code for KDE_Models without ΔR 82

Fig. S49: Plots showing the 2σ intervals for calibrated radiocarbon dates of the individuals MAN008, OTE001, AHM001, AHM011, MAL003, and STA001 with different modeling approaches. ... 88

Fig. S65: probability distributions (brackets indicating 2-sigma intervals) of ELW098 and AGU025 without offset (left) and after integrating into the sequence model with a generic offset of 30 ± 20 RC years (right). 124
**11.8 Branch 1A2, LBG002.A and the STN genomes** 125
Fig. S66: probability distribution (brackets indicating 2-sigma intervals) of Boundary T7 as part of the sequence model with a generic offset of 30 ± 20 RC years. 126
Fig. S67: Probability distributions for KDE_Models (top row), separate KDE_Plots (middle row) and KDE_Plots within the sequence model (bottom row) without (left) and with (right) a generic offset of 30 ± 20 RC years. 128
**11.9 Genomes of the Plague of the Thirty Years' War – SPN19, LAR11t, BRA001.A and the EMS genomes** 128
Fig. S68: Probability distributions of radiocarbon dates (row 1-4, left), the respective offsets (row 1-4, right) as well as the KDE_Plot (row 5) for Domat/Ems within the sequence model with generic offset of 30 ± 20 RC years. 133
**11.10 Basal Branch 1A1 – AGU007.B, AGU010.B, Gdansk8, the RUS genomes – and the provenance of Rostov2039** 134
Fig. S69: probability distributions (brackets indicating 2-sigma intervals) of AGU007 and AGU010 without offset (left) and after integrating into the sequence model with a generic offset of 30 ± 20 RC years (right). 134
Fig. S70: Probability distributions for KDE_Models (top row), separate KDE_Plots (middle row) and KDE_Plots within the sequence model (bottom row) without (left) and with (right) a generic offset of 30 ± 20 RC years. 136
Fig. S71: Probability distribution for the calibrated radiocarbon dates (left) and the offsets (right) within the sequence model and a generic offset of 30 ± 20 RC years. 137
**11.11 The Ottoman Reservoir of Branch 1A1** 141
Fig. S72: Probability distribution for Boundary T8 as part of the sequence model with a generic offset of 30 ± 20 RC years. 142
**11.12 The London New Churchyard genomes (BED) and Azov38** 143
Fig. S73: Probability distribution for the calibrated radiocarbon dates (row 1-4, left) and the respective offsets (right) as well as the KDE_Plot (row 5 left) within the sequence model and a generic offset of 30 ± 20 RC years. 145
Fig. S74: Probability distributions for KDE_Models (top row), separate KDE_Plots (middle row) and KDE_Plots within the sequence model (bottom row) without (left) and with (right) a generic offset of 30 ± 20 RC years. 146
**11.13 Genomes of the Plague of the Great Northern War – PEB10, the PRN genomes, LHM001 and potentially MOI001** 150
Fig. S75: probability distributions (brackets indicating 2-sigma intervals) of MOI001 without offset (left) and with a generic offset of 30 ± 20 RC years (right). 151
**11.14 The genomes of the Great Plague of Marseille (OBS); CHE1 and Rostov2033** 153
**12 Historical plague reservoir theories** 157
**13 Archival sources** 160
**14 References** 160

### 1 Nomenclature of *Yersinia pestis* lineages of the Second Pandemic

*Marcel Keller*

To facilitate the description and discussion of *Yersinia pestis* lineages in this paper and beyond, we propose a universal nomenclature of lineages, following the initial numbering system of main branches – established by Achtman et al. 2004 (Branches 0–2) and extended by Cui et al. 2013 (Branches 3–4) – and the designation of the sub-branches 1B (modern Branch 1 genomes and basal post-Black Death genomes from Bergen op Zoom, Bolgar and London St. Mary Graces) and 1A (all other ancient post-Black Death genomes sequenced so far), established by Green 2018. The hierarchical structure of sublineages is indicated by alternatingly appended numbers or letters to every clade with more than two distinct genomes (i.e., not counting direct ancestors representing nodes) that forms a separate lineage diverging from a main lineage. Because two similarly prominent clades emerged out of the polytomy N13, , we denote these as Branch 1A1 and Branch 1A2. The genomes G488 (Riga), NMS002 (Cambridge) and MAL004 (Mäletjärve) form the third lineage – Branch 1A3. Following this nomenclature, the genomes EMS001, EMS010 (Domat/Ems) and BRA001 (Brandenburg an der Havel), form the lineage 1A2A as a sublineage of 1A2.

### 2 Site descriptions

#### 2.1 Pärnu mnt 59, Tallinn (PRN), Harjumaa, Estonia

*Martin Malve*

During the rescue excavations carried out in 2018–2022 (Martin Malve and Tvauri 2022) 117 skeletons were unearthed at Pärnu maantee (mnt) 59 in Tallinn, the skeletons analysed as part of this study were all found in 2018 (Fig. S1). The burial site was located outside the medieval city walls, on the edge of Tõnismäe suburb on the sand dunes situated east from Pärnu mnt. So far, human remains and graves have been discovered from an area of approximately 15,000 $m^2$. The graves were situated irregularly on a large area and most of the deceased had been interred in multiple burials over a very short time period. Numerous artifacts dating to the second half of the 17th and early 18th century were found in the burials, including coinage, jewellery, garment closures and personal objects such as a clay pipe. In total, 34 Swedish coins were found in the graves, with the most recent – a 5-öre silver coin – being minted in 1710. Furthermore, at least one individual could be identified as a Russian soldier based on an Eastern Orthodox cross pendant and a rapier. Therefore, the burial site can be securely associated with a local plague outbreak in 1710 coinciding with the siege of Tallinn/Reval during the Great Northern War. Based on artifacts collected from the burials, both urban and suburban commoners, peasants, as well as Swedish and Russian soldiers and potentially their family members were interred at this plague cemetery. The deceased had been interred in a single layer and the graves were situated irregularly over a larger area. The graveyard was not organised into discernible segments, with the graves being dug individually or as smaller clusters. This layout is likely a result of the constant and continuous on-site burying of plague victims.

There were 29 single burials (the location of one of the burial 24 is unknown, as the bones were collected from the cut and fill of a later garbage pit), a remarkable 14 double burials**,** 7 triple burials and 6 mass graves. In total, the mass graves contained the remains of 29 individuals. There are no regular patterns in the mass graves, with the numbers of individuals interred together ranging from four to seven. Samples were taken from single burials 12 (PRN002; female adult; tentatively plague-positive), 53 (PRN004; female, more than 40 years old; plague-positive), 49 (PRN005; male, 40–45 years old; plague-positive); individuals 21 (PRN008; female, 20–24 years old; plague-positive), 82/6 (PRN009; female, 18–25 years old; plague-positive) and 18 (PRN010; female, 18–21 years old; plague-positive) from double burials; individual 81 (PRN006; male, 30–39 years old; plague-negative) from a triple burial; individual 58 (PRN007; female, 20–21 years old; plague-positive) from a quintuple burial; and individuals 68 (PRN001; male, 18–25 years old; plague-positive) and 66 (PRN003; female, 30–40 years old; plague-positive) from a septuple burial.

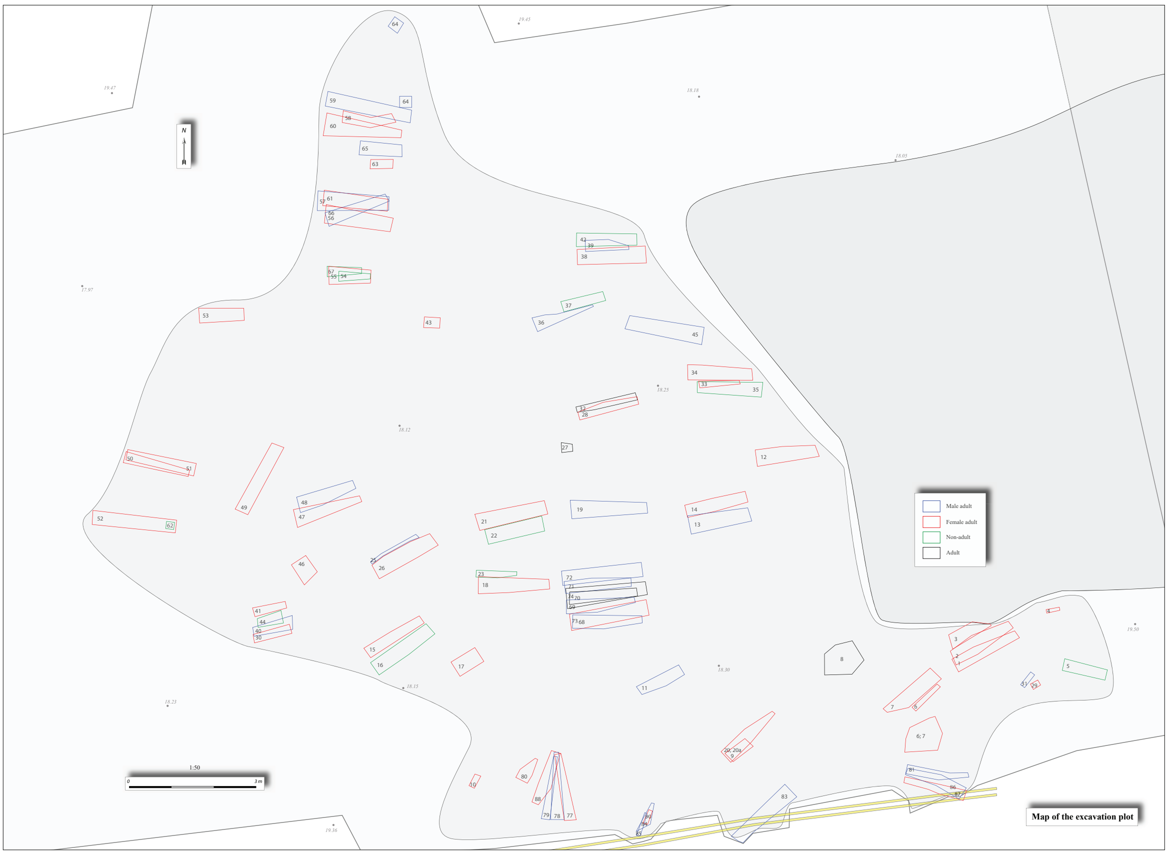

**Fig. S1: Pärnu mnt 59, Tallinn, excavation plan.**
Drawing by Raido Roog.

#### 2.2 Lehmja-Pildiküla (LHP), Harjumaa, Estonia

*Aivar Kriiska*

The burial site of Lehmja-Pildiküla, located about 10 km southeast of Tallinn, was discovered in 1990, while digging a drainage ditch (Niinre 1990). During the work, the skeletons of two people were found and partially disturbed (burial I and II based on numbering of the later excavations, see Fig. S2). An iron knife, a bronze button, a small bronze tobacco box, and an iron spur were found with the human bones. In the same year, small-scale rescue excavations (the size of the open area was 7.1 m$^2$) were carried out there (Kriiska 1990).

The burial site did not stand out in the landscape as a cemetery, the graves were dug into the cultural layer of the Lehmja settlement site. According to the finds, this part of the village was probably inhabited in the 17th century (Kriiska 2017). In total, three burials were found about half a meter below the modern ground (Fig. S2).

From burial I (LHP001, plague-positive), the lower part of the left leg, a right tibia and a right foot were recovered *in situ*. Iron spurs were preserved around both feet and a bronze chain was found near the right tibia. The skull, pelvis, femora and some ribs found during the digging of the drainage ditch also belong to the same burial. The individual was identified as 172±2 cm tall male, aged 30–35 at death, with an antemortem (~3-4 years) blunt force trauma to the skull (anthropological studies by Leiu Heapost and Galina Sarap; Kriiska 1990).

Burial II, south of burial I, contained a skull, claviculae, right humerus and ulna, right rib cage and the upper part of the spine of a 170±2 cm tall male individual aged around 25–30 years at the time of death, lying in a supine position with the right arm positioned at the side. Associated objects included a spur around the left foot and four bronze buttons. Another spur found in the soil thrown out of the ditch is probably also connected with this burial.

Burial III, located north of Burial I, was found in a supine position with arms crossed on the left shoulder. This individual was identified as a 174±2 cm tall male, aged 25–30 years at the time of death. The burial contained two bronze mounts and two bronze buttons.

All burials were oriented with their heads to the West, no traces of coffins or burial pits were found. The skeletons were positioned adjacent to each other and interred roughly on the same level, implying that they were contemporary with each other. Further findings potentially associated with Burials I and II (found near skeletons or in soil removed from a ditch) are a whetstone, two knives, a Dutch smoking pipe, some bronze buttons, an orthodox cross, and three coins. The coins were identified as an oval Russian silver *kopek* minted in the 1690s, a Russian denga minted in 1705 and a Russian *kopek* minted in 1708 – all from the reign of Peter the Great.

The findings of spurs, an Eastern Orthodox cross and pieces of baize on buttons (determined by Jüri Peets; Kriiska 1994) in burials are exceptional compared to contemporary burial grounds of the native (Lutheran) Estonian population. Due to the similar age and the sex of the burials as well as finds including buttons, spurs, and baize referring to uniforms, the three individuals may be identified as troopers of a Russian dragoon regiment, stationed in the hinterland of Tallinn/Reval during the siege of the city in 1710 (Kriiska 1994). At the same time, there was a plague outbreak in the region, as it is well documented in historical sources (Hartmann 1973, 69–72; Parts 2010).

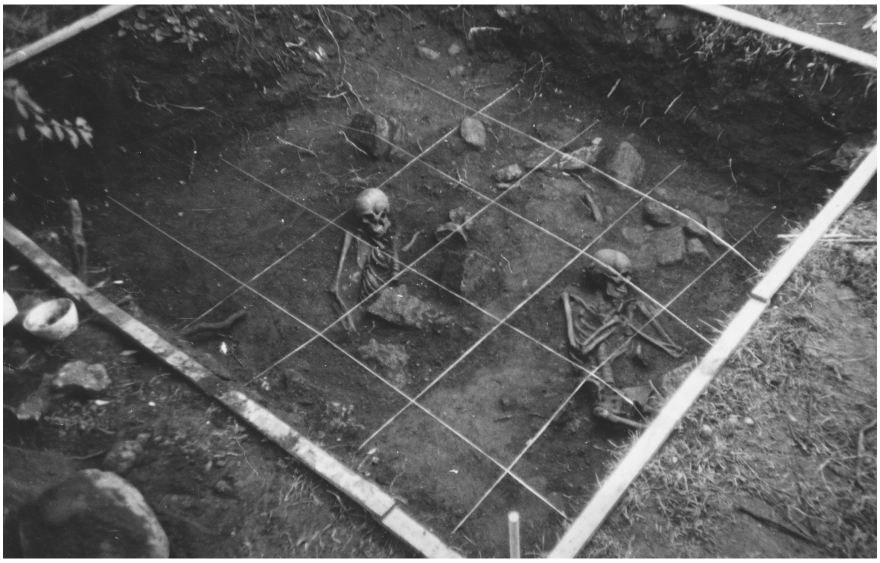

**Fig. S2: Lehmja-Pildiküla burial site.**
Burials II (left) and III (right), burial I was located between them. Photo: Aivar Kriiska.

#### 2.3 Otepää (OTE), Valgamaa, Estonia

*Heiki Valk*

Otepää, a small town in south-eastern Estonia was in medieval times an important centre of Roman Catholic Principality-Bishopric of Tartu/Dorpat – a centre consisting of a strong castle, attached borough and parish church. The cemetery at the crossing of Tartu and Piiri streets is one of the three medieval cemeteries of Otepää. Rescue excavations took place there in 1928, 1929 and 1938. During the Soviet period, a large part of the cemetery was destroyed, with new rescue excavations taking place there in 1996 and 2020. In total, 150 burials have been unearthed from the cemetery (Valk 1997; Malve and Valk 2021).

The general pattern of the cemetery at the crossing of Tartu and Piiri streets differs greatly from that of medieval village cemeteries. The site is of short time use and almost no graves were disturbed by later burials. A specific feature of the site is that burials are arranged in rows, often in groups of 2–4 individuals, and often buried together in the same grave. This feature is most atypical for village cemeteries in the region, making it possible to associate these burials with a mortality crisis, such as an epidemic. The assemblage of findings – consisting of jewellery (brooches, rings, necklaces), metal accessories of belt and costume, and knives – is characteristic of medieval Estonian village cemeteries, indicating the native origin of the buried individuals. The cemetery was probably attached to the medieval borough of Otepää. The assemblage of finds is characteristic of the time span between ca. 1250 and 1450, but the lack of disturbed graves indicates a limited period of use. A more definite date is provided by the contents of a purse where the latest coin was minted between ca. 1370–1375 (?), the others dating from the second or third quarter of the 14$^{th}$ century.

As part of a pilot study, only one individual, burial 1 (OTE011; male, 25–35 years old; plague-positive; Fig. S3) was sampled for ancient DNA.

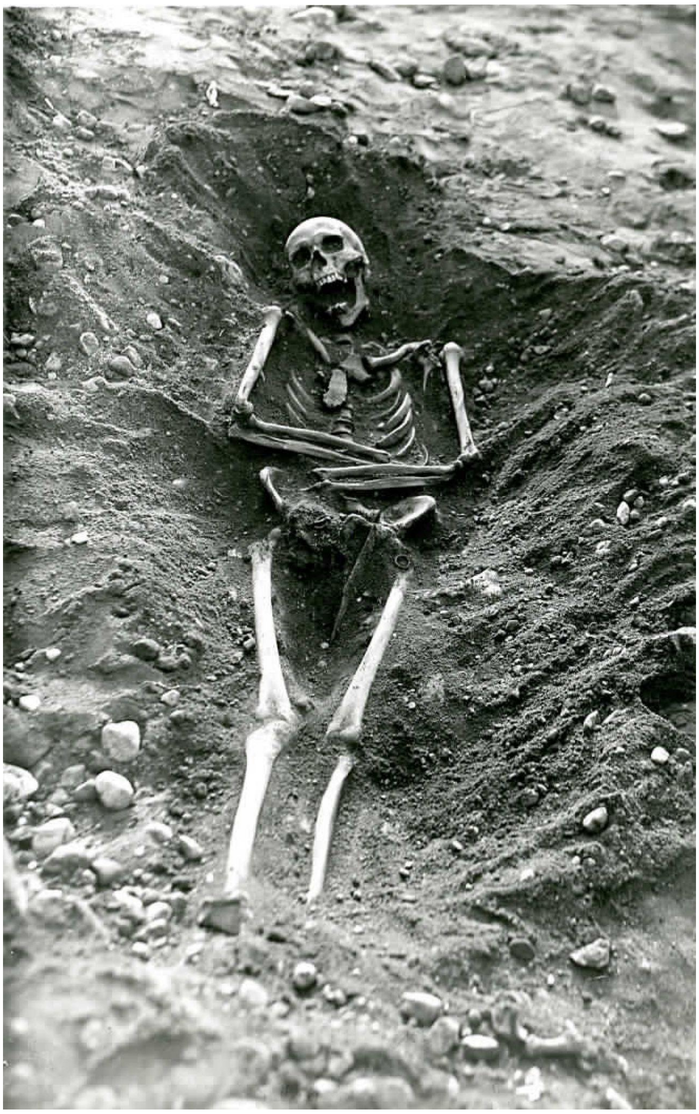

**Fig. S3: Photograph from 1938 of the excavation of Otepää burial 1.**
Photo: Osvald Saadre.

#### 2.4 Mäletjärve village cemetery (MAL), Tartumaa, Estonia

*Heiki Valk, Marcel Keller*

A specific feature of medieval Estonian funerary practices is a double system of burial grounds: church graveyards, founded at the time of Northern crusades and Christianization ($12^{th}$–13th centuries), coexisted with the pre-Christian tradition of local village burial sites. The main features of burial rites at these local cemeteries mostly conform to Christian burial practices (inhumation, head orientation towards the West). In southern Estonia, it was also a commonplace practice to bury the dead with jewellery, coins, small tools and utensils. The village cemeteries, evidently legal in Catholic period but prohibited by the Lutheran church, remained in use until the 1720s or 1730s.

The cemetery of Mäletjärve, located in south-eastern Estonia, ca. 20 km south-east of Tartu, is one of such village cemeteries (Fig. S4). It was founded beside a 3rd–5th-century cemetery with cremations and stone settings. During the 1984 excavations, 50 skeletons, mainly from the 16th century, were unearthed from a trench measuring 60 $m^2$ (Valk 1985).

The dead were buried in shallow graves (ca. 60 cm deep), often furnished with brooches, rings, coins and knives, sometimes with needles. Most of the interred individuals were oriented with the head towards the south-west. In the outline of the cemetery, vague grave rows can be observed.

In total, seven individuals were sampled for ancient DNA: burial numbers 38 (MAL003) and 39 (MAL004) corresponding, respectively, to an adult woman (aged 30–53 years; 165 cm) and a 9.5–10.5-year-old girl (130 cm). With the distance between humerus bones being 8 cm and with both individuals buried found on the same horizon, the burial was interpreted as simultaneous double burial (Fig. S5). A coin from the filling of their grave, a so-called “penny of Damerow” (or bracteate, Fig. S6) was minted between 1379 and 1420. A brooch and two rings were found in the woman’s grave (Burial 38) and a brooch and a necklace with 2 lead medallions, one of which depicting the image of Virgin Mary and Jesus the Child, were found in the girl’s grave (Burial 39). These burials are, judging by their finds, the oldest in the excavated part of the cemetery and their orientation (NW) also differs from those of the others, buried with the head mainly towards south-west.
As shown in Fig. 2, the two *Y. pestis* genomes reconstructed for 38/MAL003 and 39/MAL004 occupy vastly different positions in the phylogenetic tree, and also radiocarbon dating yielded divergent intervals. For a detailed discussion, see Supplementary Section 11.6.
Grave 28 (MAL007) is a typical grave of the cemetery, and belongs to an adult female individual, buried with a ring, a knife and two coins – one from 1537 and the other from 1561–1568.

Further sampled were the burials 49 (MAL001, male, 30–40 years old), 18 (MAL002, male of unknown age), 15 (MAL005, female, 45+ years old) and 26 (MAL006, male, 12–18 years old), which were all tested negative for plague.

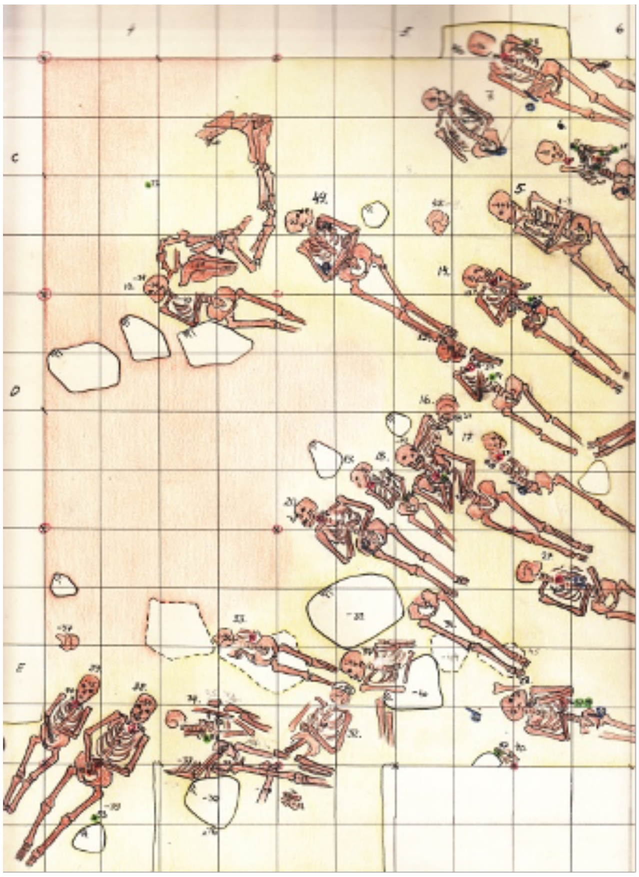

**Fig. S4: Mäletjärve cemetery, abstract from the excavation plan with burials 38, 39 and 28.**

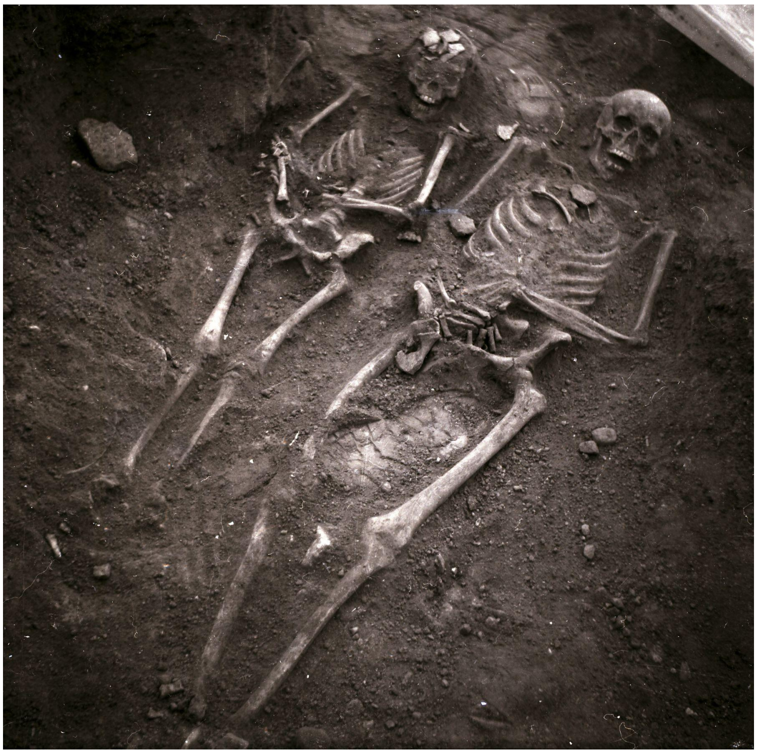

**Fig. S5: Mäletjärve cemetery, burials 38 (taller) and 39 (shorter, from the south-east).**
Photo: Heiki Valk.

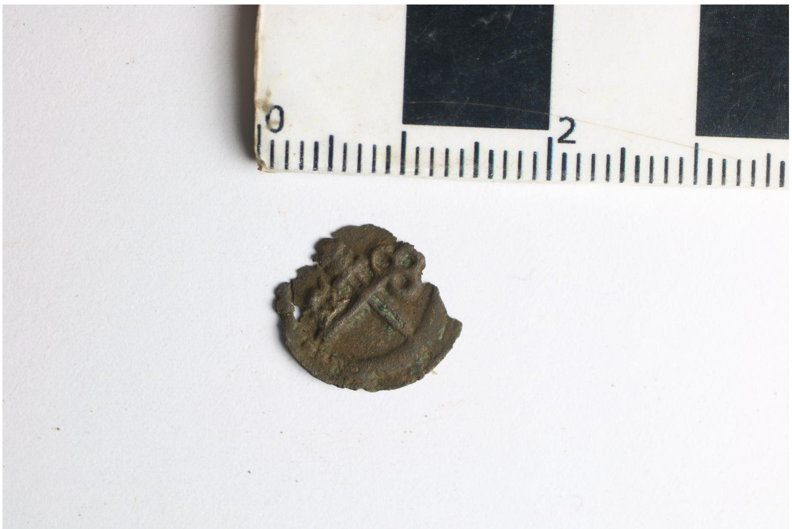

**Fig. S6: Mäletjärve cemetery. "Penny of Damerow".**
Minted between 1379 and 1420, found in the grave fill of the burials 38/39. Photo: Heiki Valk.

#### 2.5 Mõisaküla Margu cemetery (MOI), Pärnumaa, Estonia

*Heiki Valk*

The cemetery of Margu in Mõisaküla village, Pärnumaa County, Estonia, was studied in conjunction with rescue excavations in 1986 (Pärn 1987). The burial site, damaged by a gravel pit, was located beside an Iron Age cemetery with cremation graves from the 11th–13th centuries. A total of 49 burials, partly damaged by later graves or gravel digging, and representing both sexes and different age groups, were found in the core area of the cemetery (c.60 $m^2$).

The skeletons lay in the depth of 0.3–0.5 m and were oriented with the head towards NW, W and SW, sometimes towards N, NE, and E. In some cases, coffin remains were found. About half of the graves were furnished with grave goods – mainly coins, rings and knives dating to the 16th and 17th centuries. Since Swedish schillings minted in Riga in 1630s–1660s remained in circulation all the way to the first quarter of the 18th century, the latest graves may belong also to that period. Burial 4 (MOI001, unknown sex and age, plague-positive) was the only individual sampled from this site and radiocarbon dated to 1640 or younger (see also Supplementary Section 10.13 and Fig. S75).

#### 2.6 Stankeyevo village cemetery (RUS), Pskov Oblast, Russia

*Heiki Valk*

The cemetery of Stankeyevo is located in north-western Russia, Pskov oblast, c.110 km south of Pskov, 1 km north-west of Krasnogorodsk (district centre), and 600 m south-west of Stankeyevo village. The site lies on a sandy hillock 1.5 km north-west of medieval Krasnyi stronghold, along the Gavry-Krasnogorodsk road. The cemetery was discovered in 2016 during road construction, and rescue excavations were carried out by the Pskov Oblast Archaeology Centre, under the supervision of Viktoria Derkach and Svetlana Shun'gina (Derkach and Shun'gina 2018).

Two intact unfurnished burials were found in a trench (measuring 23.75 $m^2$) dug in the central part of the hillock, with each measuring at about 40–50 cm in depth and 170 cm in length. The skeletons were in single graves, located beside each other, oriented with their heads towards south-west and having no grave goods, as characteristic of medieval cemeteries in that region. Burial 1 (RUS004) was identified as an adult female and burial 2 (RUS004) as an adult male, interred in a wooden coffin, preserved in a decayed state. Overall, the cemetery was preliminarily dated to the 14th–16th centuries but it may also be younger.

#### 2.7 Bene't Street/Corpus Christi (BEN), Cambridge, Cambridgeshire, United Kingdom

*Craig Cessford, Marcel Keller*

St Bene’t’s (St Benedict’s) was established in the first half of the 11th century as a parish church in Cambridge, with the adjacent burial ground remaining in use until the 1850s. At some point between 1352 and 1377, part of the churchyard west of the church was transferred to the Corpus Christi College and turned into an entrance to the college from Bene’t Street. This strip of land was excavated in 2006 by the Cambridge Archaeological Unit and revealed a section of mass burial (F.1522), whose full dimensions are unclear, since the borders could not be reached. Only five individuals could be excavated within the small area (1.0 x 0.4 m), but the stacking in at least three layers suggests that the total number of buried individuals was as a minimum at least two to three times higher, and there could even by 100–120 burials assuming a maximum potential extent.

The mass burial, so far the only one known in Cambridge, must have been established before 1352-1377, when Corpus Christi College was built and this part of the cemetery was closed.

Plague outbreaks in Cambridge and East Anglia during this timeframe are recorded for 1349 (Black Death), 1361 (*pestis secunda*), 1369 and 1375. A more extensive discussion of the site was published by Cessford et al. 2021.

Teeth of four individuals were sampled for ancient DNA: 1608 (BEN001, 18–25-year-old female, plague-positive), 1609 (BEN002, 16–25-year-old female, plague-positive), 1610 (BEN003, adult male), and 1612 (BEN004, at least 60-year-old male).

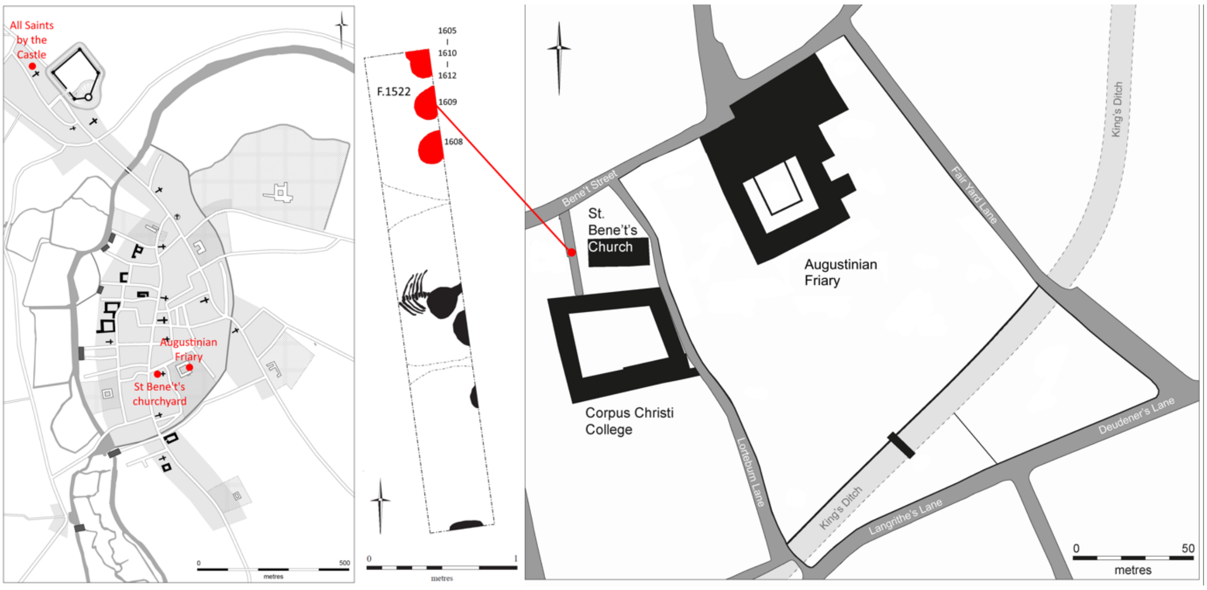

**Fig. S7: Overview of Cambridge sites with palaeogenetic evidence for *Y. pestis;* plan of Bene't Street.** Left: Plan of Cambridge c.1350 showing locations of sites where skeletons have tested positive for *Y. pestis*. Right: Part of Cambridge showing St. Bene't's Church, Corpus Christi College, the Augustinian friary with the location of the mass burial shown as a red dot. Centre: detail of trench excavated, with individuals in mass burial shown in red. Based on original figures produced by Vicki Herring for the After the Plague project and by the Cambridge Archaeological Unit.

#### 2.8 All Saints by the Castle (COM), Cambridge, Cambridgeshire, United Kingdom

*Craig Cessford, Marcel Keller*

The site of All Saints by the Castle, also known as "Comet Place", was principally excavated in 1973 by Paul Craddock and Vince Gregory. In total, over 200 skeletons have been uncovered from this cemetery. The parish church was founded between 940–1150 and was in use until 1365/1366 when it was merged with close-by St Giles, which might have been a consequence of population loss due to the Black Death (1349) and the *pestis secunda* (1361). By 1365/1366, the cemetery was abandoned, therefore offering a *terminus ante quem* for all burials, which is supported by the available archaeological evidence for the cemetery including radiocarbon dating.

In total, 48 individuals were sampled for ancient DNA (see Table S1), with all dating to between the 10th and mid-14th centuries. The skeleton 14 was accidentally sampled twice (COM009, COM044) as revealed later through later analysis. Only one of the individuals was positive for *Y. pestis*: skeleton 82 (COM042) was identified as a 46–59-years-old male who was buried in a single burial typical for parish cemeteries of this period (Fig. S8). Interestingly, the latest grave in the local stratigraphic sequence was a double burial (143 and 156) in prone position, but due to later truncation, the skulls were missing and thus could not be sampled for ancient DNA. A more extensive discussion of the site was published by Cessford et al. 2021.

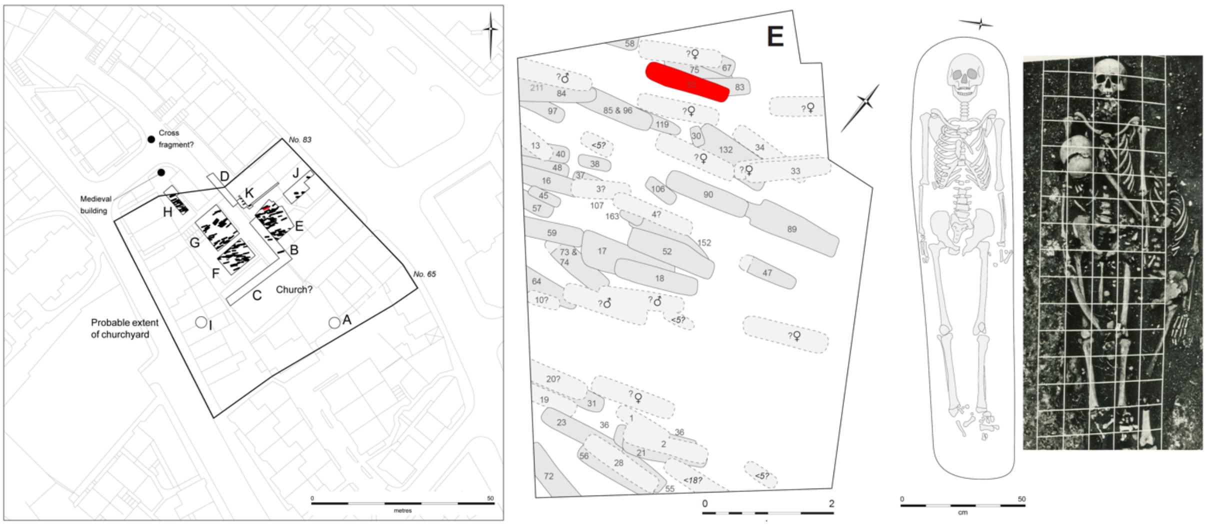

**Fig. S8: Plan of All Saints by the Castle; plan and photograph of burial 82.**
Plan of the cemetery of All Saints by the Castle, showing all known archaeological interventions (left), plan of Interventions E with *Y. pestis* positive skeleton shown in red (centre) and plan and photograph of *Y. pestis* positive skeleton 82 (right). Plans based on original figures produced by Vicki Herring for the After the Plague project, photograph courtesy of Vince Gregory.

#### 2.9 St Mary's, Clopton (CLP), Cambridgeshire, United Kingdom

*Craig Cessford, Marcel Keller*

The deserted village of Clopton was located around 19 km southwest of Cambridge. The age of the parish church St. Mary's Clopton is unknown; it appears first in written records in 1254. Between 1960 and 1964, John Alexander excavated the site with a focus in 1964 on the church building and the cemetery to the south of it (Fig. S9). Alexander identified that the southern part of the cemetery was established on an artificial terrace covering a 12th-century ditch and dates to the 13th–15th centuries, based on pottery finds – although this dating could not be re-confirmed. Due to the shrinking population size, the parish was merged with the adjacent parish of Croydon in 1561, proposing a *terminus ante quem* for all the burials. This dating scheme is compatible with recent radiocarbon dating of the cemetery.
In total, teeth of 17 individuals were sampled for ancient DNA, see Table S1. Three individuals were positive for *Y. pestis*: S9.24 (CLP006, 13–17-year-old female), S6.3I (CLP015, 36–45-year-old male) and S4.4 (CLP017, 25–44-year-old male). All of them were identified as single burials that appear typical for the cemetery. A more extensive discussion of the site was published by Cessford et al. 2021.

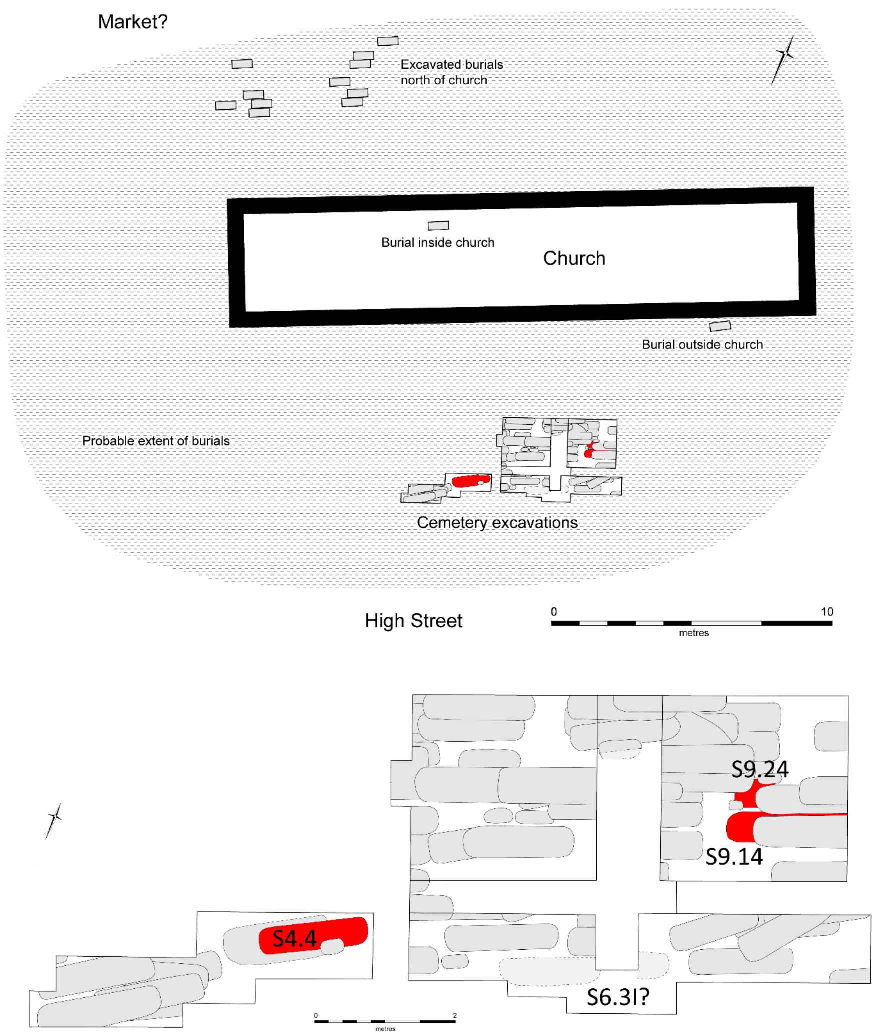

**Fig. S9: Plan of St Mary's Clopton church and cemetery (upper) plus detail of excavations with *Y. pestis* positive/tentative skeletons shown in red (lower).**
Based on original figure produced by Vicki Herring for the After the Plague project.

#### 2.10 Sogn Pieder, Domat/Ems (EMS), Graubünden, Switzerland

*Marcel Keller*

The town of Domat (Romansh) or Ems (German) is located in the Alpine Rhine valley close to Chur in the Swiss canton of Graubünden. Sogn Pieder (Romansh for St. Peter) is the oldest church of Domat/Ems. The church dates back to the Carolingian period, but was built on the foundations of an even older *curtis* (homestead) dating to the 7th century. While the adjacent agricultural and residential building structures of the Carolingian complex were demolished in the 13th century, the church itself survived until today, making it one of only a few Carolingian churches in Switzerland that are still in use.

These exceptional findings were revealed during excavations between 1975 and 1979 by the archaeological service (Archäologischer Dienst) Graubünden and the “Büro für Archäologie des Mittelalters und Bauforschung” of Hans Rudolf Sennhauser. The excavations elucidated not only the long history of the church building and its secular progenitor, but also uncovered a medieval cemetery South and East of the church and an Early Modern cemetery in the North and West (Fig. S10). The results of the excavations, the anthropological examinations of the skeletons, done by Christine Cooper, and preliminary palaeogenetic results were published in Archäologischer Dienst Graubünden and Burkhardt 2020, herein especially Seifert et al. 2020.

The early modern cemetery seems to be closely connected with the so-called “Bündner Wirren” (1618–1639), geostrategic and confessional conflicts between the Spanish-Austrian and Franco-Venetian coalitions during the Thirty Years’ War (1618–1648). In total, 68 individuals were buried primarily in the North of the church in an area of around 150 $m^2$. Three burials with five individuals are located in the West; the dating of two burials found under the foundations of the later-built church tower on the Western side is not secure. In contrast to the medieval cemetery in the South, this cemetery is characterized by its regular outline, the abundance of multiple burials with two to four individuals, age and sex bias as well as the grave goods. The positions of the burials must have been known during the usage time of the cemetery, although no indications for gravestones or post holes were found, suggesting that the cemetery was in use for only a short period of time. Radiocarbon analysis confirmed a dating between 1433 and 1632 (widest span; see Table S11 and Supplementary Section 11.9). Grave goods, including pilgrimage coins (“Wallfahrtspfennige”) and a relic medallion offer a *terminus post quem* of 1610. Military tools found in Burial 23, which is part of the quadruple burial, allow the individual’s identification as a soldier. Several rosaries found in the cemetery reveal the Catholic confession of the buried individuals.

Two single burials were dug at 160 cm below ground level and contained coffins nails. All other burials, including all multiple burials, were only 40–50 cm deep and did not show any evidence for the use of coffins.

14 out of 46 burials were identified as multiple burials: one quadruple burial (23/24/25/26), six triple burials (2/3/5, 8/11/12, 27/28/29, 89A/89B/89C, 91/92/97, 101/102/103) and seven double burials (14/15, 86/87, 94A/94B, 96A/96B, 98A/98B, 99A/99B, 109/110). In several cases, individuals were added to a single or double burial in opposite orientation (94A/94B,

98A/98B, 99A/99B, 2/3/5, 8/11/12, 27/28/29, 101/102/103) without disturbance of the burial below, therefore a simultaneous interment can be assumed for all multiple burials (Fig. S11). All individuals were buried in supine position with varying arm postures.

An in-situ age estimation revealed a demographic pattern contradicting a regular, “attritional” cemetery. Of all individuals, only four individuals were children (6%, in contrast to 46.5% for the medieval cemetery). Anthropological examinations performed by Christine Cooper on the 15 individuals of 6 multiple burials further confirmed the peculiar demography. She identified a 20–30-year-old female (94B) and an infant (89C), the remaining 13 individuals were identified as male with an age-at-death between 17 and 40. None of the individuals showed signs of perimortal trauma, therefore an epidemic seemed the most likely explanation for the high frequency of multiple burials.

Regarding the dating to the 17th century, the identification of military tools and the demography, the individuals buried in the multiple burials are most likely soldiers of the Austrian-Spanish troops that were stationed in Chur, Felsberg and Domat/Ems during the “Bündner Wirren”. This would also explain the installation of this separate cemetery for the ‘foreigners’, since the parish church in the 17th century was Sogn Gion Battista (St. John Baptist, established latest in the 12th century) with a surrounding parish cemetery for the local community (von Planta 2020).

Based on this evidence, the burials can be dated more narrowly to 1629–1631. During these years, two and a half companies of the regiment of Alwig von Sulz were stationed in Domat/Ems, left behind by the imperial army of commander Ramboldo XIII, Count of Collalto, on the way to Northern Italy. Major sources are the “damage accounts” sent from Ems to the Three Leagues between 1620 and 1632, pleading for financial adjustment for the expenses caused by the stationing of foreign troops in the town (von Planta 2020).
This coincides with plague outbreaks in the Three Leagues in 1628–1632, the last plague wave of the Second Pandemic in Graubünden. After the Black Death in Raetia (1348) and the *pestis secunda* in Chur (1361), plague reappears in the chronicles of Graubünden not before 1550. The chronicles report severe outbreaks of bubonic plague in Chur in 1550, 1556, 1560, 1566 and 1574, in Thusis in 1581, and in Graubünden in general between 1584 and 1595 (von Sprecher 1942; Lorenz 1868–1869). Accepting 1610 as *terminus post quem* (see above), these outbreaks can be excluded for the plague burials of Domat/Ems. In the following years until 1628, the chronicles report only on an outbreak of the “Hungarian disease” in 1622–1623, which can clearly be distinguished from plague (von Sprecher 1942). Later outbreaks in Germany, Italy and the Old Swiss Confederacy between 1665 and 1668 could be held back by strict border restrictions (Maissen 1971).

Unfortunately, local parish registers as well as other records of Domat/Ems were destroyed in the town fire of 1776, hampering the identification of the individuals – and possibly their cause of death – buried at Sogn Pieder. However, a reconstructed parish register mentions that the priest of Domat/Ems died of plague in 1631.

In total, twelve individuals were sampled for ancient DNA (see Table S1): individuals 24 (EMS001, 25–35-year-old male, plague-positive), 25 (EMS002, 25–30-year-old male, plague-positive) and 26 (EMS003, 20–25-year-old male, plague-positive) of the quadruple burial 23/24/25/26; all individuals of the triple burial 89A (EMS004, 17–20-year-old male),

89B (EMS005, 17–20-year-old male) and 89C (EMS006, 6–12 months old); the double burial 94A (EMS007, 17–20-year-old male) and 94B (EMS008, 20–30-year-old female, plague-positive); the double burial 98A (EMS009, 6–12-months-old male) and 98B (EMS0010, 20–30-year-old male, plague-positive); and the double burial 99A (EMS0011, 18–22-year-old male) and 99B (EMS0012, 17–20-year-old male).

The phylogenetic clustering of the Domat/Ems genomes with genomes from Brandenburg an der Havel (1618–1648), Naturno/Naturns (1636) and Lariey, (1629–1630) further support a dating to the time of the Thirty Years' War (Spyrou et al. 2019; Guellil et al. 2020; Seguin-Orlando 2021). Taking together the archaeological findings and dating, radiocarbon dating, demographic data, and phylogenetic analyses, the plague burials can securely be dated to the years 1629–1631. For a more detailed discussion of the 1629–1631 outbreak in the region and its likely associated with the Ems specimens, see Supplementary Section 11.9.

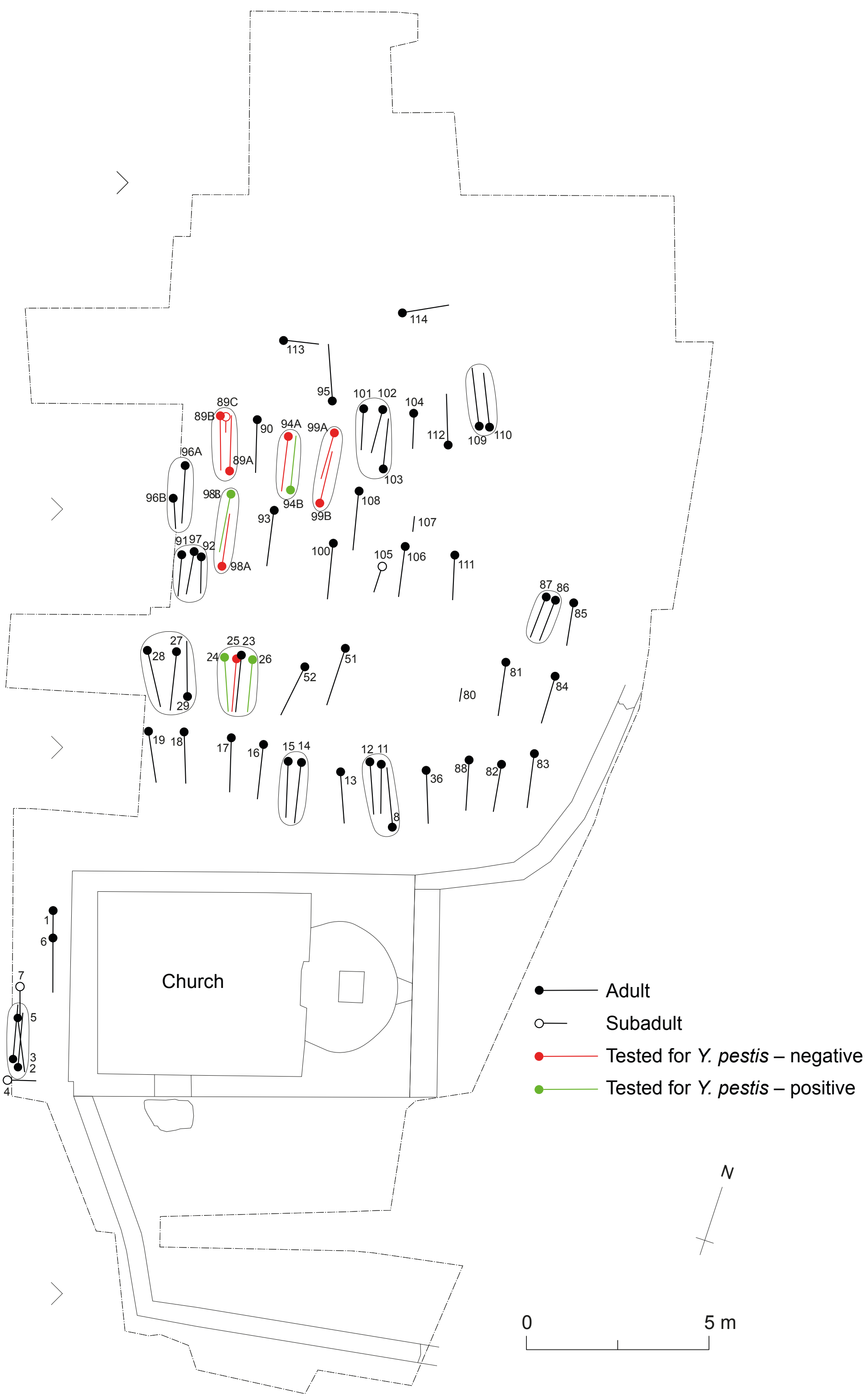

**Fig. S10: Plan of the early modern cemetery of Sogn Pieder, Domat/Ems.**

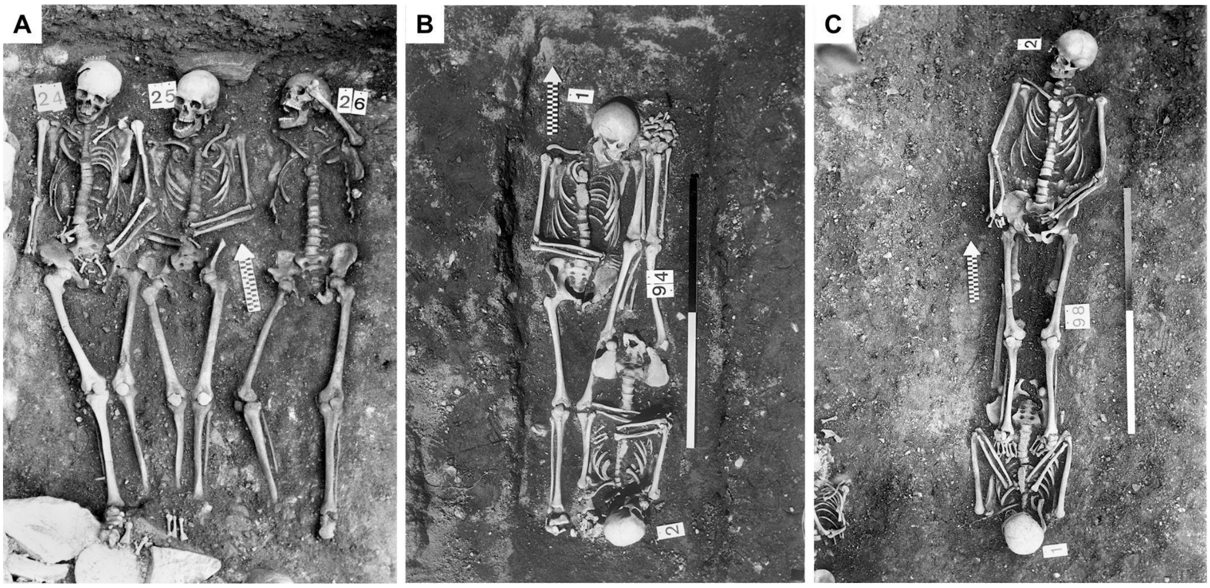

**Fig. S11: Multiple burials of Domat/Ems positive for *Y. pestis* DNA.**
A: Quadruple burial 23/24/25/26 – burial 23 was situated above burials 25 and 26 and is already removed here; B: double burial 94A/94B; C: double burial 98A/98B. Photo: Archaeological Service of the Canton of Grisons.

#### 2.11 St. Eusebius' churchyard, Arnhem (AHM), The Netherlands

*Willem A. Baetsen*

During a major part of 2017, the city centre of Arnhem (the Netherlands) was the scene of construction works that were commissioned for the purpose of a large urban renewal project. A vital component of this project was the reintegration of 'St. John's Brook' (Dutch: St. Jansbeek) into the cityscape: a stream that in the past had played an essential role in the establishment of Arnhem and had been the lifeblood of the city. For centuries, the brook had been part of the daily lives of its inhabitants, until it was filled in by the end of the 19$^{th}$ century (Vierdag 2020). At present, the stream is expected to be reinstated by public initiative and with municipal support (Wolthuis 2020).

However, because of the urban developments in the past century – not in the least due to damage during World War II – the city plan looks quite different now than it did when the St. Jansbeek was last an open stream. This means that, in order for the project to succeed without tearing down many of Arnhem's buildings, an alternative route through the city centre had to be conceived. This new course would flow through areas of significant archaeological interest, such as the former monastery of the Friars Minor (Dutch: Minderbroedersklooster) and sections of the historical brook. Of main interest for this paper, the brook would also pass along the northern side of St. Eusebius' Church, the largest church in Arnhem, past and present. The same church also boasted, at least until 1829, the largest churchyard in the city (Fig. S12). Significantly, the construction plans envisioned an open stream at this location, with a slope leading up to street level. Since the works necessary to execute these plans would likely cause major damage to the underlying cemetery, archaeological excavations were conducted.

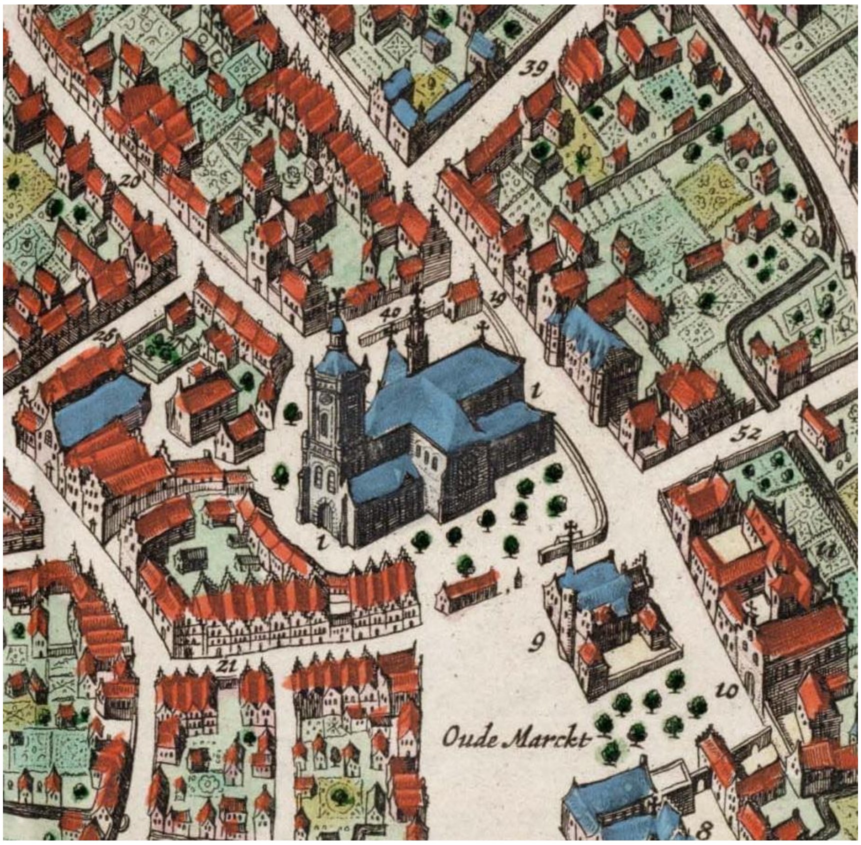

**Fig. S12: Detail of a map of Arnhem by Blaeu, showing St. Eusebius' Church with surrounding churchyard.**
Image source: ("GA 1551, Inv.nr. 4187-0004 (Gelders Archief 1551 Topografisch-Historische Atlas Gelderland, Inventarisnummer 4187-0004). Aernhem, 1649. Plattegrond van Arnhem Op Een Schaal van 1:3000 Door Joan Blaeu" n.d.)

The excavation along most of the course of the new St. Jansbeek and subsequent post-excavation work were conducted by RAAP Archeologisch Adviesbureau B.V. Trenches 8 and 10, to the north of St. Eusebius' Church, traversed the former churchyard of the extant church and its predecessor, St. Martin's Church (Dutch: Maartenskerk). Finds recovered from between the graves, in conjunction with radiocarbon dates from human bone collagen, date the active period of the churchyard between at least the late 14th century and 1829 (Zielman and Baetsen 2020, 94–104). During the medieval and post-medieval periods, many of the citizens of Arnhem were buried around these churches (Zielman and Baetsen 2020, 83–94). Afterwards, the churchyard was completely covered over and the resulting square had several functions over past 200 years, including a fish market, and a poultry market (Zielman and Baetsen 2020, 116–117).

At this location, a total of 659 primary graves in over ten burial layers were meticulously uncovered, documented and lifted, along with over thirty charnel pits and numerous disarticulated bones (Fig. S13; Baetsen and Baetsen 2020, 382). The majority of the primary individuals were buried in a supine position, with their feet roughly positioned towards the east. Significant grave disturbances were noted, mainly in the western part of trench 10, as a result of digging new graves in an already densely populated churchyard in the past. Consequently, many of the individuals were recovered in an incomplete state. Nevertheless, for 47.8% of primary individuals more than half of the skeleton, including cranium and pelvis, were available for assessment (Baetsen and Baetsen 2020, 386–392).

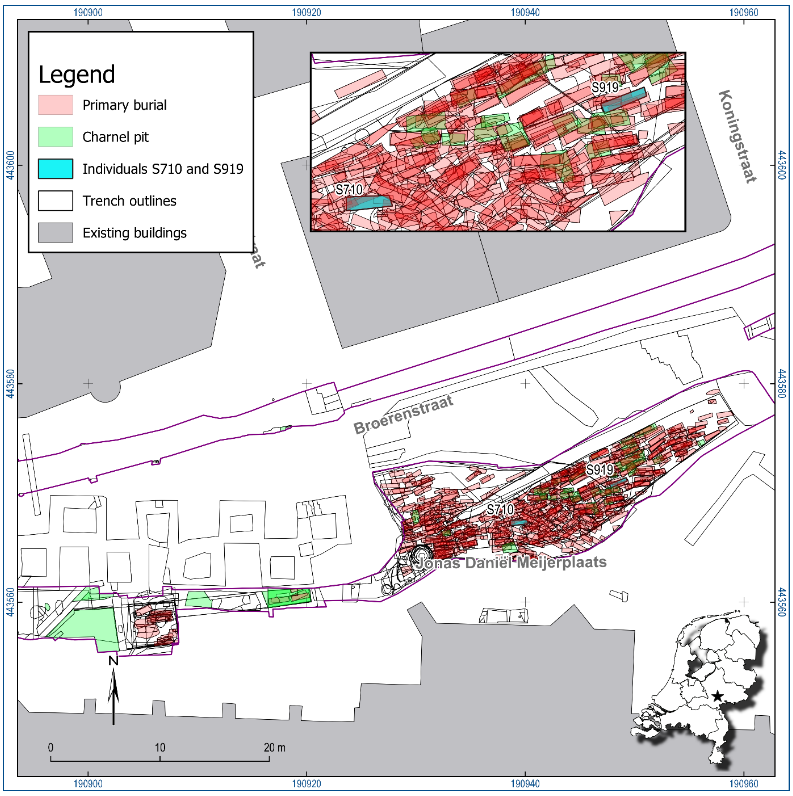

**Fig. S13: Results of the excavations north of St. Eusebius' Church (bottom of the plan, in grey) in Arnhem, showing the primary burials and charnel pits.**
Individuals S710/V1401 and S919/V1883, referred to in this paper, are highlighted. Top inset: Detailed view of the burial positions within the churchyard. Bottom inset: Location of Arnhem in the Netherlands. Image by Willem A. Baetsen, generated using QGIS, version 3.10.3-A Coruña.

Over the course of 2018 and 2019, all primary individuals were macroscopically analysed. The results of all archaeological investigations of the St. Jansbeek project were reported near the end of 2020 (Zielman and Baetsen 2020; Baetsen and Zielman 2020).

Two of the primary burials are of main importance to this paper, namely individuals S710/V1401 (AHM011, Fig. S14) and S919/V1883 (AHM001, Fig. S15). Both are adult individuals who were included in an initial DNA screening sample as part of the kinship analyses of the Jansbeek project. Although no familial relationships between these individuals and the other cases were identified, metagenomic screening did detect DNA of *Y. pestis* (Baetsen and Scheib 2020).

Individual S710/V1401 (AHM011, plague-positive) was first dated stratigraphically to the early phase of the churchyard, namely between 1350 and 1650, and most likely in the earliest period, judging by the burial location and level. Stratigraphical dates were derived from typochronological dates of finds throughout the burial grounds, and several strategically placed radiocarbon dates of bone collagen belonging to primary burials scattered across the churchyard. For the purpose of the current paper, radiocarbon analysis of this individual specifically was undertaken, confirming stratigraphical dating and providing a dating between 1300 and 1399 calAD (2-sigma).

Osteological assessment of sex estimated this individual to be female, which was confirmed by human aDNA analysis. Macroscopic analysis further estimates that this individual died

between the ages of 36 and 49 years old, and must have stood about 162 cm tall. Macroscopic analysis of palaeopathology only yielded slight signs of rotator cuff disease (RCD) around the tuberculum minor on both humeri. Her skeleton was unearthed together with the remains of a young child between the ages of 3 and 6 years old, positioned on her lower legs (individual S715/V1413; Fig. S14). As previously mentioned, human aDNA analysis (using the petrous bone) showed no familial relationship between these two individuals.
Individual S919/V1883 (AHM001, plague-positive), osteologically indeterminate but genetically demonstrated to be female, was also originally included in the human aDNA analyses for an assessment of kinship, because of a contextual relationship to a non-adult individual (S919/V1884; Fig. S15). These latter remains were of foetal age, estimated 20–26 weeks *in utero*. Unfortunately, human aDNA analysis to assess familial relationship did not yield sufficient coverage in the case of individual S919/V1884. Therefore, the hypothesis of a mother-child relationship could not be confirmed.

Individual S919/V1883 (AHM001) was stratigraphically dated to the late phase of the churchyard, between 1650 and 1829. Radiocarbon analysis of the remains of this individual specifically yielded insufficient results to narrow down this estimate. Macroscopic osteological analysis estimated age-at-death between 26 and 35 years old, with a stature of approximately 160 cm tall. This puts this individual, just like individual S710/V1401 (AHM011), in line with the average female height in this population. No pathology was macroscopically apparent. Regardless, as mentioned, human aDNA analysis of skeletal materials belonging to both these adult female individuals has demonstrated that they both suffered from and likely died from the plague.

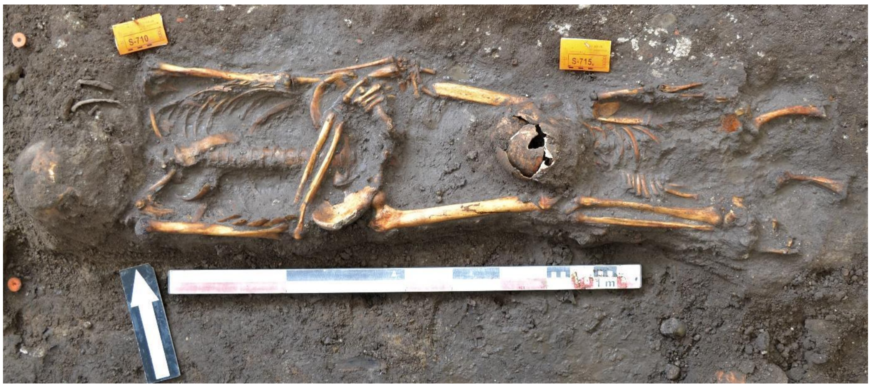

**Fig. S14: Burial positions of individuals S710/V1401 (AHM011, left) and S715/V1413 (right).**
DNA analysis did not show any familial relationships (image source: Baetsen & Scheib, 2020: 607).

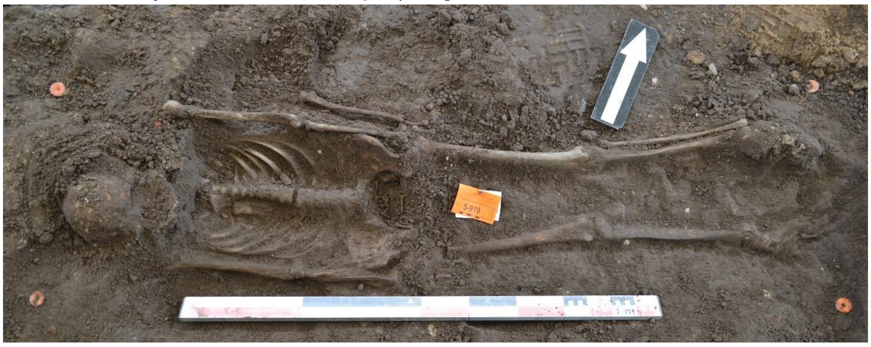

**Fig. S15: Burial position of individual S919/V1883 (AHM001).**
The foetal remains of S919/V1884 can be made out in the pelvic opening. DNA analysis could not yield sufficient coverage for confirmation of kinship. Image by RAAP.

### 3 Selection of modern *Yersinia pestis* genomes

*Marcel Keller*

In addition to *Yersinia pseudotuberculosis* IP32953, 237 published or publicly available modern genomes were considered for the reference data set (Auerbach et al. 2007; Chain et al. 2004; Chain et al. 2006; Cui et al. 2013; Deng et al. 2002; Eppinger et al. 2009; Eroshenko et al. 2017; Garcia et al. 2007; Kislichkina et al. 2015, 2017; Kislichkina, Bogun, Kadnikova, Maiskaya, Solomentsev, Sizova, et al. 2018; Kislichkina, Bogun, Kadnikova, Maiskaya, Solomentsev, Dentovskaya, et al. 2018; Kutyrev et al. 2018; Morelli et al. 2010; Parkhill et al. 2001; Song et al. 2004; Zhgenti et al. 2015). We produced a SNP alignment containing 7150 positions using MultiVCFAnalyzer (v0.85.2 with outgroup mode for *Y. pseudotuberculosis* IP32953, minimum genotyping quality of 30, minimal coverage of 3-fold and a minimal allele frequency of 0.9 for homozygous calls) and used IQTree v2.1.2 to build a phylogenetic tree using bootstrapping (1000 replications) and the GTR+F+ASC+R2 model (Fig. S16).

The inclusion of *Y. pestis* assemblies from recent publications resulted in phylogenies with extremely long tips observed for a number of said assemblies which do not seem to occur with previously published genomes with raw sequencing files available, even for strains in the same clade. This raised the suspicion that the responsible SNPs are either an artifact of data generation, contamination, or laboratory propagation with elevated mutagenesis. Artificially long tips interfere with phylogenetic analyses because of long-branch attraction, and molecular dating approaches due to the introduction of extreme rate heterogeneity. Since original sequencing raw data is not available and the cause for this phenomenon could not be investigated, we searched for a metric to flag problematic genomes for exclusion. Problematic genomes identified upon visual inspection of the phylogenetic tree coincided well with the number of homoplastic sites, which are not to be expected in higher numbers in a clonal organism such as *Y. pestis*. In an analysis including only modern genomes, we identified in total 633 homoplastic sites with up to 95 per genome. *Y. pseudotuberculosis* IP32953, used as an outgroup, showed 65 homoplastic sites. The highest number of homoplastic sites for a genome called from raw sequencing data was 24 (average 17.05). For further analyses, all genomes with equal or more than 44 homoplastic sites except *Y. pseudotuberculosis* IP32953 (25 of 238 genomes) were excluded.

In addition, IQTree identified 0.ANT5_A-1691 (Eroshenko et al. 2017) to be identical with 0.ANT5_5M (Kutyrev et al. 2018) and 0.PE5_I-2238 (Kislichkina, Bogun, Kadnikova, Maiskaya, Solomentsev, Sizova, et al. 2018) to be identical with 0.PE5_I-2231 (Kislichkina, Bogun, Kadnikova, Maiskaya, Solomentsev, Sizova, et al. 2018). Therefore, the genomes 0.ANT5_5M, 0.PE5_I-2238 were excluded from further analyses. Conversely, genomes with the ID I-2422 (assigned to 0.PE5) were published three times by (Kislichkina et al. 2015), (Kislichkina, Bogun, Kadnikova, Maiskaya, Solomentsev, Sizova, et al. 2018) and (Kutyrev et al. 2018) reveal a different issue. Whereas the strains published in 2018 were identical and contained the pMT1 plasmid, the strain published in 2015, lacking pMT1 was not identified as identical by IQTree. They were therefore labelled 0.PE5_I-2422a (Kislichkina et al. 2015) and 0.PE5_I-2422b (Kislichkina, Bogun, Kadnikova, Maiskaya, Solomentsev, Sizova, et al. 2018) and both included in the phylogenetic analyses.

To identify problematic genomes, we also calculated the number of multiallelic sites with MultiVCFAnalyzer (v0.85.2, 0.1 and 0.9 as thresholds for multiallelic sites). Of all genomes called from raw sequencing data, 0.PE7a_CMCC05009 was identified as outlier with 441 multiallelic sites (second highest 173, median 41.5).

After the exclusion of 25 genomes with the highest number of homoplatic sites, 0.PE7a_CMCC05009 due to excess of multilallelic sites as well as the duplicates 0.PE5_I-2238 and 0.ANT5_5M, 3676 sites of 209 sequences remained (Table S2). In this reduced dataset, in total 185 homoplastic sites were identified. The maximum number of homoplastic sites was reduced to 46 (*Y. pseudotuberculosis* IP32953) with 9.57 on average.

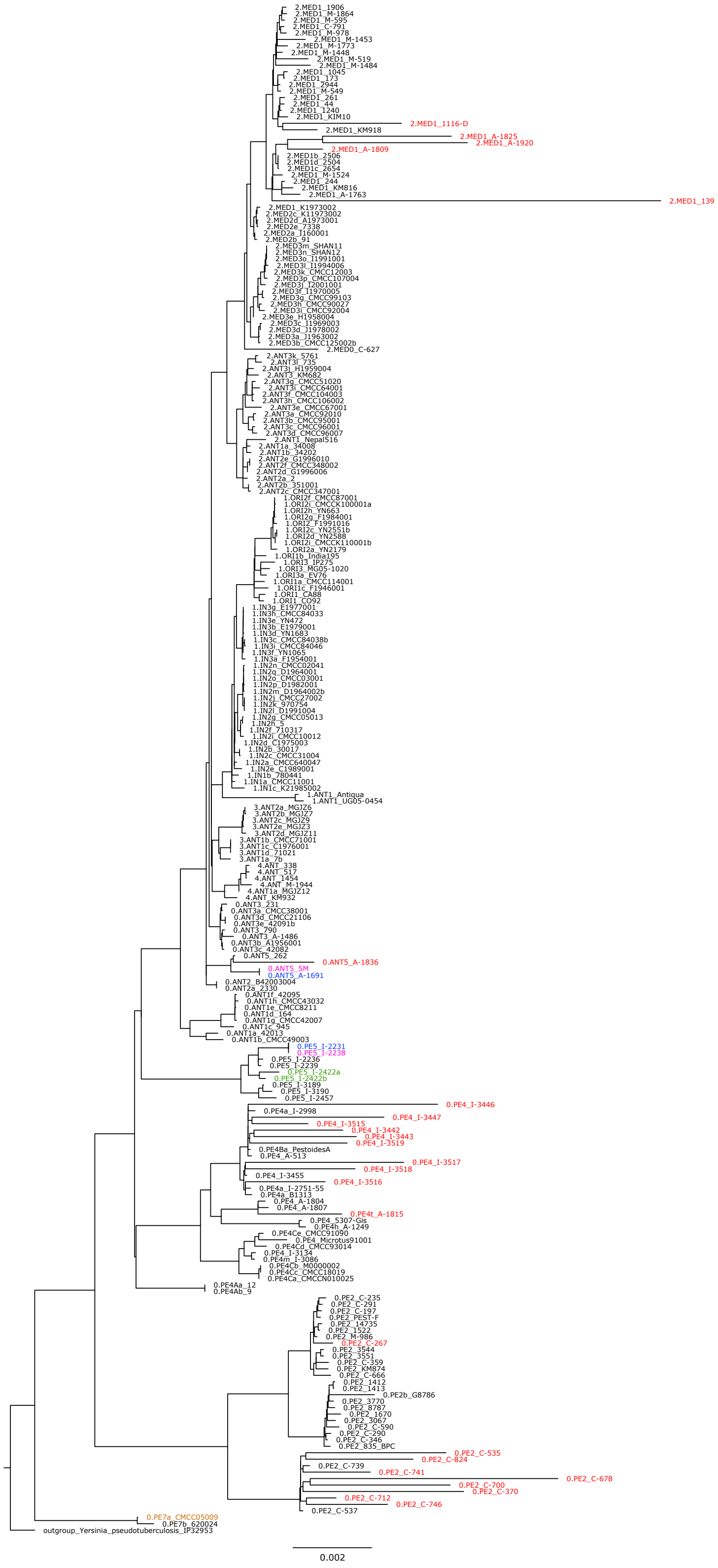

**Fig. S16: Phylogenetic tree of all considered modern Y. pestis genomes.**
Maximum likelihood tree generated with IQTree with 1000 bootstraps based on a full SNP alignment (7150 positions) of 236 modern *Y. pestis* genomes and *Y. pseudotuberculosis* IP32953 as outgroup. Genomes highlighted in red were excluded due to high numbers of homoplastic sites, the genome highlighted in brown was excluded due to an excessive number of multiallelic sites. Genomes highlighted in purple were excluded since they appeared identical with the genomes highlighted in blue. The genomes highlighted in green have the same ID in the original publication but were both kept in (with the suffixes a and b) since they occupy different phylogenetic positions.

### 4 Comparison of different in-silico treatments of non-UDG data

*Marcel Keller*

DNA damage in form of deamination of Cytosine to Uracil, read by polymerases as Thymine, can infer with SNP calling. Therefore, UDG treatment is often used in ancient pathogen genomics. Nevertheless, a significant part of published ancient genomes derive from non-UDG libraries, and also in the present study we present newly sequenced genomes reconstructed from non-UDG libraries. In the past, several strategies were used to accommodate for DNA damage in ancient *Y. pestis* genomes: bwa-aln with -n 0.01, -l 16 is a commonly used strategy allowing for higher edit distance (Andrades Valtueña et al. 2017; Spyrou et al. 2019), while others used -n 0.1 with seed disabled (-l 1000) to remove reads with excessive damage (Guellil et al. 2020; Namouchi et al. 2018). Another strategy is rescaling of base quality scores of potential deaminated sites with mapDamage2 (Rascovan et al. 2019). Other options are the trimming of fastq-files prior to mapping or soft-clipping of bam-files after mapping, as both implemented as possibilities in nf-core/eager (Fellows Yates et al. 2021). For this study, we compared seven different in-silico treatments of non-UDG libraries on two newly sequenced libraries (MAL003, MAL004) and twelve previously published genomes from different studies which we integrated in our phylogenetic analyses with coverages ranging from around 4 to almost 200-fold.

The tested treatments are:

- [0.01] mapping with bwa-aln -n 0.01, -l 16 (common for non-UDG libraries).
- [0.04] mapping with bwa-aln -n 0.04, -l 1000 (default settings, seeding disabled).
- [0.1] mapping with bwa-aln -n 0.1, -l 32 (common fpr UDG libraries).
- [clip fastq] trimming of 3 bases on both ends on fastq files followed by mapping with mapping with bwa-aln -n 0.1, -l 32.
- [clip bam] mapping with bwa-aln -n 0.01, -l 16 followed by soft-clipping of 3 bases on both ends on bam-files.
- [Rs] mapping with bwa-aln -n 0.01, -l 16, rescaling with mapDamage2.
- [RsRm] mapping with bwa-aln -n 0.01, -l 16, rescaling with mapDamage2, remapping with bwa-aln -n 0.1, -l 32.
-

To compare the different treatments, we generated a SNP table with MultiVCFAnalyzer (v0.85.2; Bos et al. 2014) both on the full dataset including modern *Y. pestis* genomes and on Second Pandemic genomes alone, and generated a phylogenetic tree including modern genomes on the full SNP alignment. Based on the Second Pandemic SNP alignment, we identified private SNPs forming terminal branches for each library-treatment combination and 132 phylogenetically informative positions that form internal branches (positions called at least in 14 out of 56 Second Pandemic genomes, representing 25% of Second Pandemic genomes or 5% of 265 genomes including modern genomes), as shown in Fig. S17. For private SNPs, we differentiated between SNPs potentially deriving from deamination (C>T, G>A; red) and other SNPs (grey), for phylogenetically informative positions between SNPs potentially misidentified as damage (C>T, G>A; green), other SNPs/reference calls (grey), and Ns (coverage <3X or multiallelic site). In addition, we also generated the plots on other ancient genomes included in our phylogenetic analyses derived from UDG libraries, partial

UDG-treated libraries (G488/G701) and heterogeneous, pre-treated data (BSK001.A) for comparison.

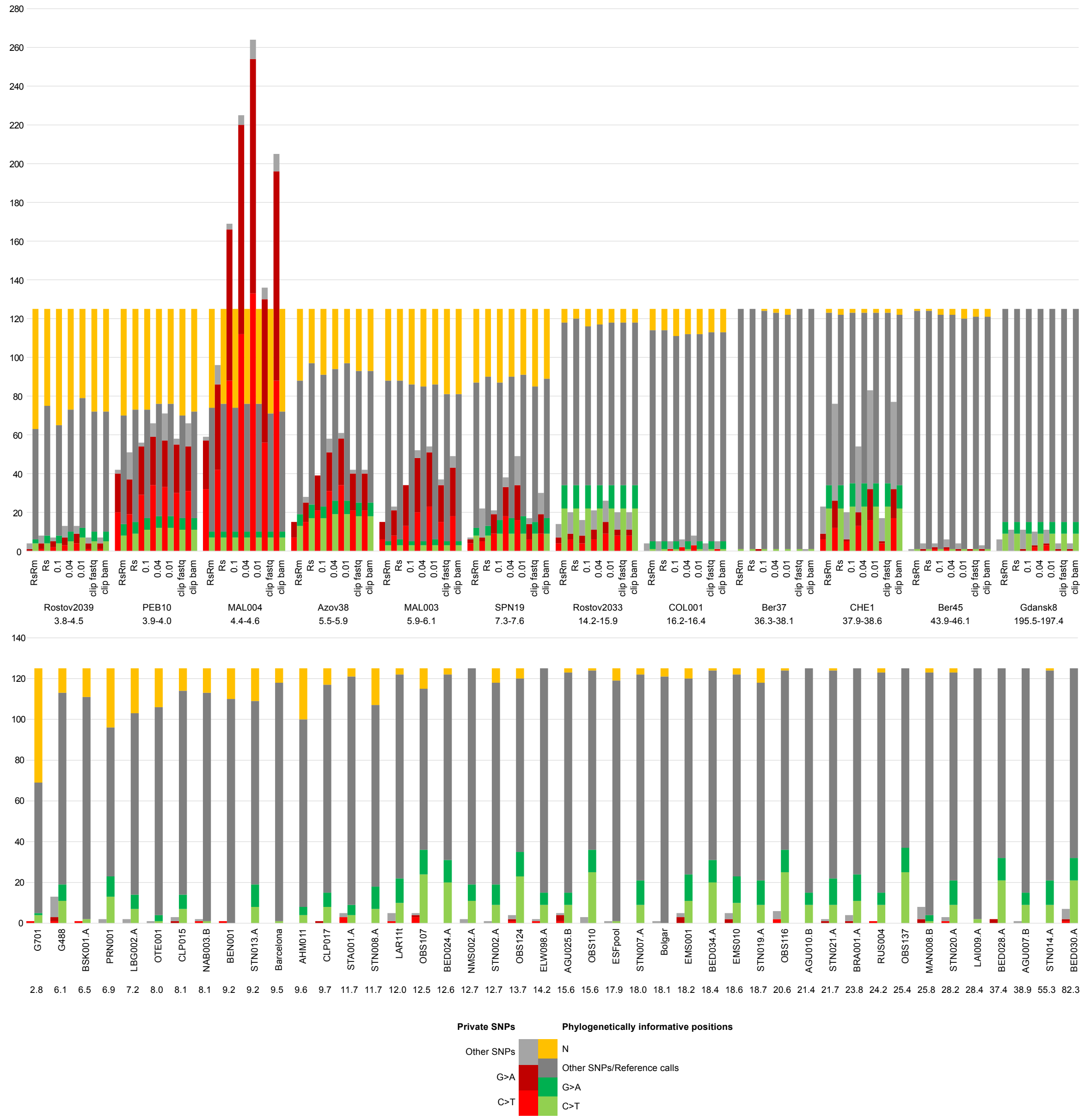

**Fig. S17: Comparison of private SNPs and phylogenetically informative positions with different non-UDG treatments.**

Upper panel: effects of different non-UDG data in-silico treatments on the absolute numbers of private SNPs and phylogenetically informative positions. Lower panel: comparison for genomes derived from full-UDG, partial-UDG or heterogeneous pre-treated data. The share of private SNPs that might be caused by DNA damage is shown in red; the share phylogenetically informative positions that could be misidentified as DNA damage is shown in green; the share of phylogenetically informative positions that could not be called due to low coverage or ambiguous base calls are shown in yellow.

In particular, non-UDG libraries with lower than 4–5-fold coverage show excessive numbers of private SNPs. The fact that the vast majority of them are either C>T or G>A substitutions suggests that most of these SNPs derive from deamination. In general, the number of private SNPs decreases with more stringent mapping (0.01 to 0.1). Trimming of fastq-files prior to mapping performs generally better in reducing SNPs compared to clipping of bam-files. The lowest number of private SNPs is generally achieved by rescaling and remapping, followed by rescaling only. In both cases, the reduction compared to 0.1 is primarily driven by the reduction of potentially deaminated sites. Reduction through remapping, however, is also characterized by loss of positions that cannot be derived from damage and are likely caused by contamination with highly similar reads of other bacteria (cf. RsRm and 0.1). Considering phylogenetically informative positions, the number of called positions generally increases with higher coverage. However, when comparing different in-silico treatments, no clear pattern is visible. Considering C>T and G>A substitutions, potentially misinterpreted by mapDamage2 as deamination and therefore downscaled, some samples show a slight decrease in Rs and RsRm proportional to other called positions. Therefore, we see no clear indication for rescaling and remapping to introduce a reference bias.

We generated a phylogenetic tree comparing all treatments together (Fig. S18, full SNP alignment) and all treatments separately (Fig. S19–25, SNP alignment with 95% partial deletion) to investigate how the different treatments affect the tree topology and branch lengths. In Fig. S18 in particular, it is apparent that “Rs” and “RsRm” can significantly reduce terminal branch lengths, whereas there is only little effect on the topological position of the respective genomes in the tree. To avoid artifacts of long branch attraction, we also present individual phylogenetic trees for each treatment (Fig. S19–25). Also here, the gross topologies are identical, although partial deletion will also affect SNPs of other ancient and modern genomes differently in the individual datasets. Affected genomes are MAL004, branching off either from the polytomy giving rise to Branches 1A1 and 1A2 (RsRm, 0.04. 0.01), or from the Gdansk8 cluster (Rs, 0.1, clip fastq, clip bam); and CHE1 and Rostov2033, which share a short branch (Rs, 0.04, 0.01, clip bam), or emerge from a polytomy (RsRm, 0.1, clip fastq).

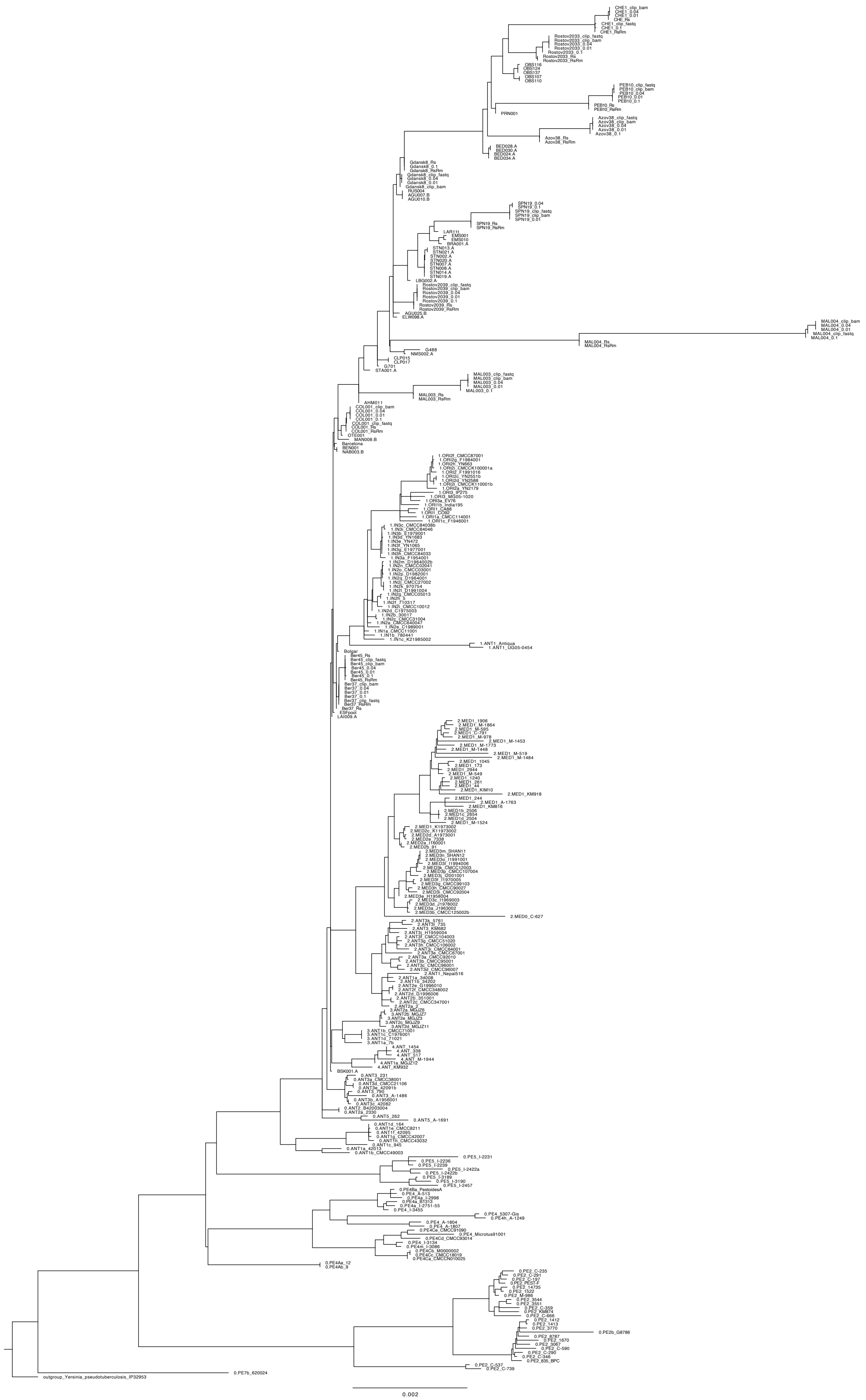

**Fig. S18: Phylogenetic tree comparing all tested non-UDG treatments.**
Maximum likelihood tree generated with IQTree with 1000 bootstraps based on a full SNP alignment (7150 positions) of 128 ancient *Y. pestis* genomes (including 12 genomes reconstructed from non-UDG * 7 different in-silico non-UDG data treatments = 84 genomes), 208 modern *Y. pestis* genomes and *Y. pseudotuberculosis* IP32953 as outgroup.

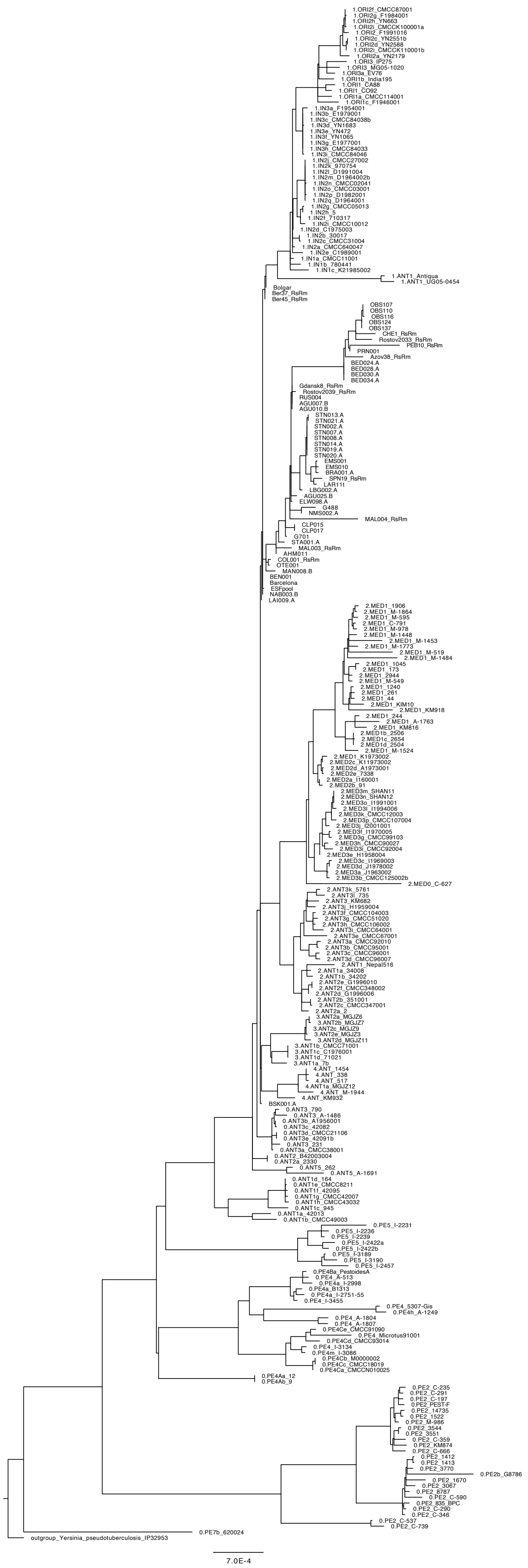

**Fig. S19: Phylogenetic tree with "RsRm" treatment.**
Maximum likelihood tree generated with IQTree with 1000 bootstraps based on a 95% partial deletion SNP alignment (3524 positions) of 56 ancient *Y. pestis* genomes, including 12 genomes reconstructed from non-UDG data and with treatment "RsRm", 208 modern *Y. pestis* genomes and *Y. pseudotuberculosis* IP32953 as outgroup.

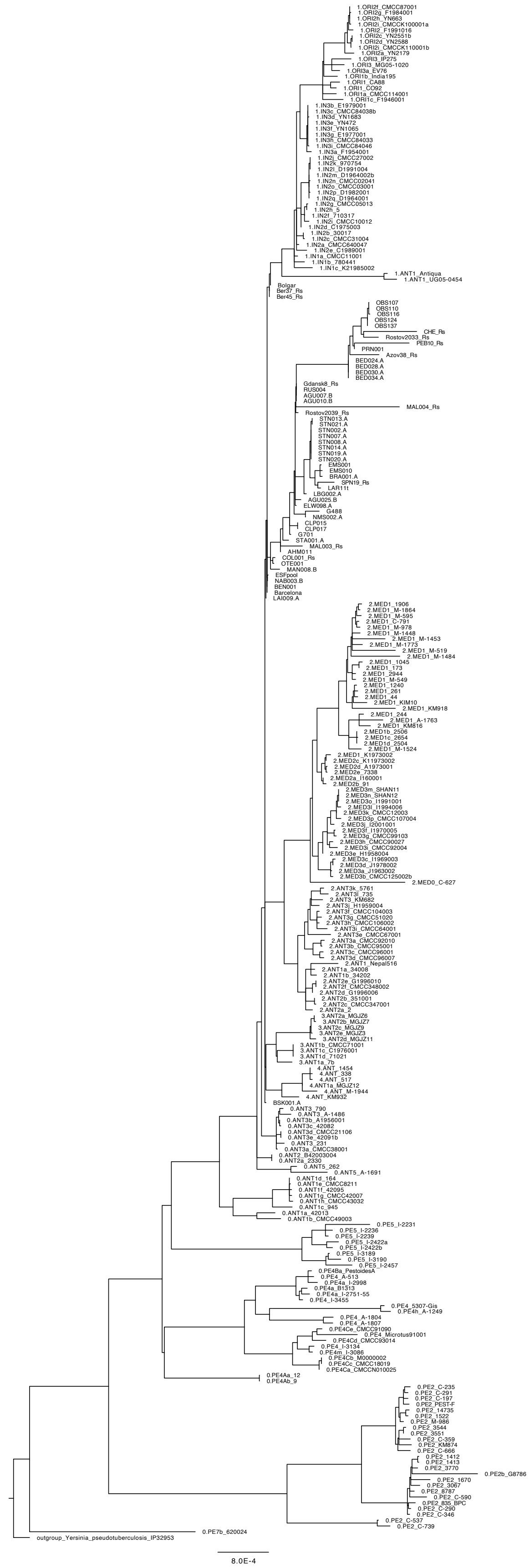

**Fig. S20: Phylogenetic tree with “Rs” treatment.**
Maximum likelihood tree generated with IQTree with 1000 bootstraps based on a 95% partial deletion SNP alignment (3625 positions) of 56 ancient *Y. pestis* genomes, including 12 genomes reconstructed from non-UDG data and with treatment “Rs”, 208 modern *Y. pestis* genomes and *Y. pseudotuberculosis* IP32953 as outgroup.

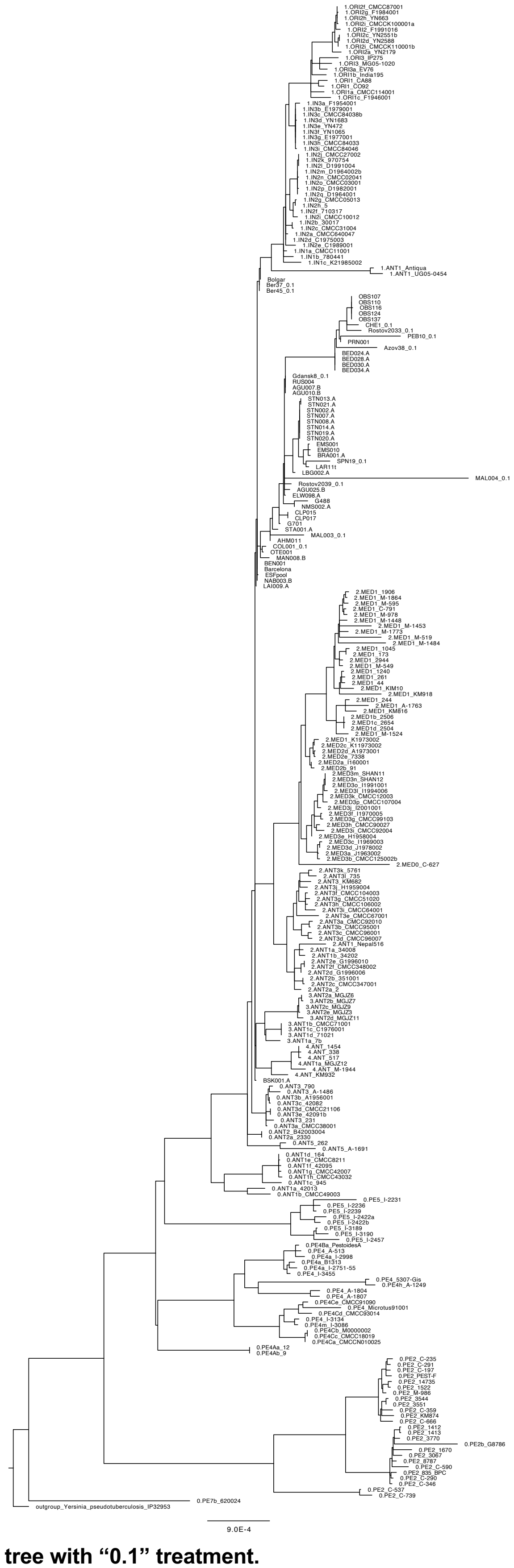

**Fig. S21: Phylogenetic tree with “0.1” treatment.**

Maximum likelihood tree generated with IQTree with 1000 bootstraps based on a 95% partial deletion SNP alignment (3802 positions) of 56 ancient *Y. pestis* genomes, including 12 genomes reconstructed from non-UDG data and with treatment “0.1”, 208 modern *Y. pestis* genomes and *Y. pseudotuberculosis* IP32953 as outgroup.

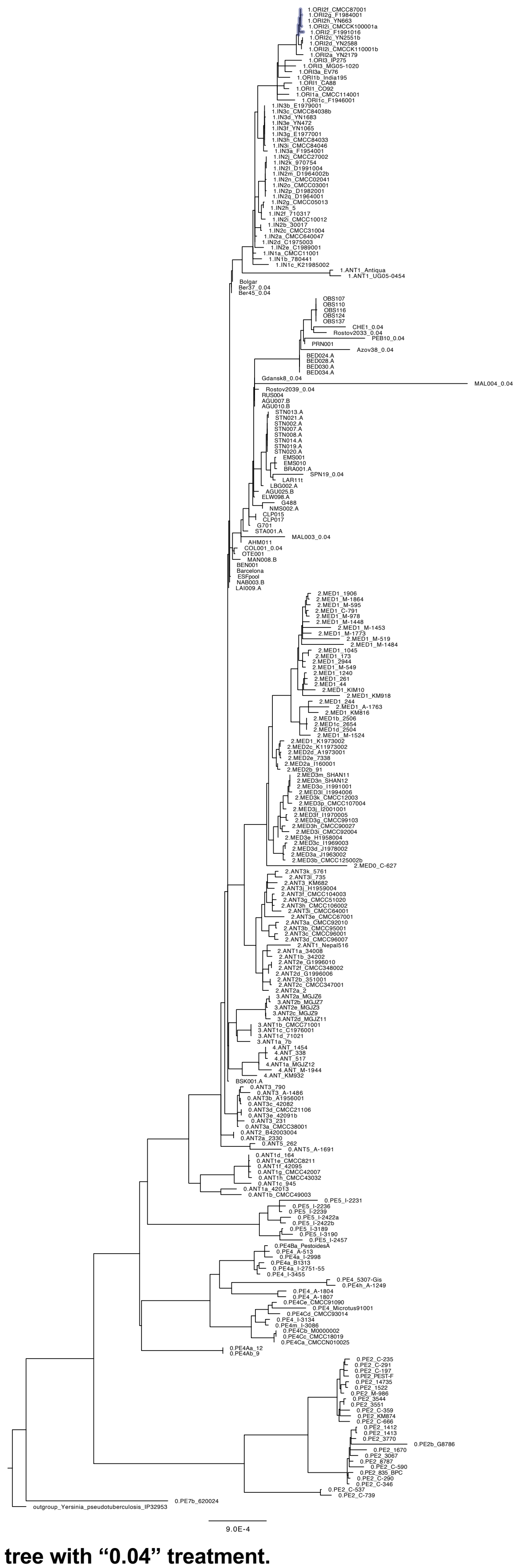

**Fig. S22: Phylogenetic tree with “0.04” treatment.**
Maximum likelihood tree generated with IQTree with 1000 bootstraps based on a 95% partial deletion SNP alignment (3936 positions) of 56 ancient *Y. pestis* genomes, including 12 genomes reconstructed from non-UDG data and with treatment “0.04”, 208 modern *Y. pestis* genomes and *Y. pseudotuberculosis* IP32953 as outgroup.

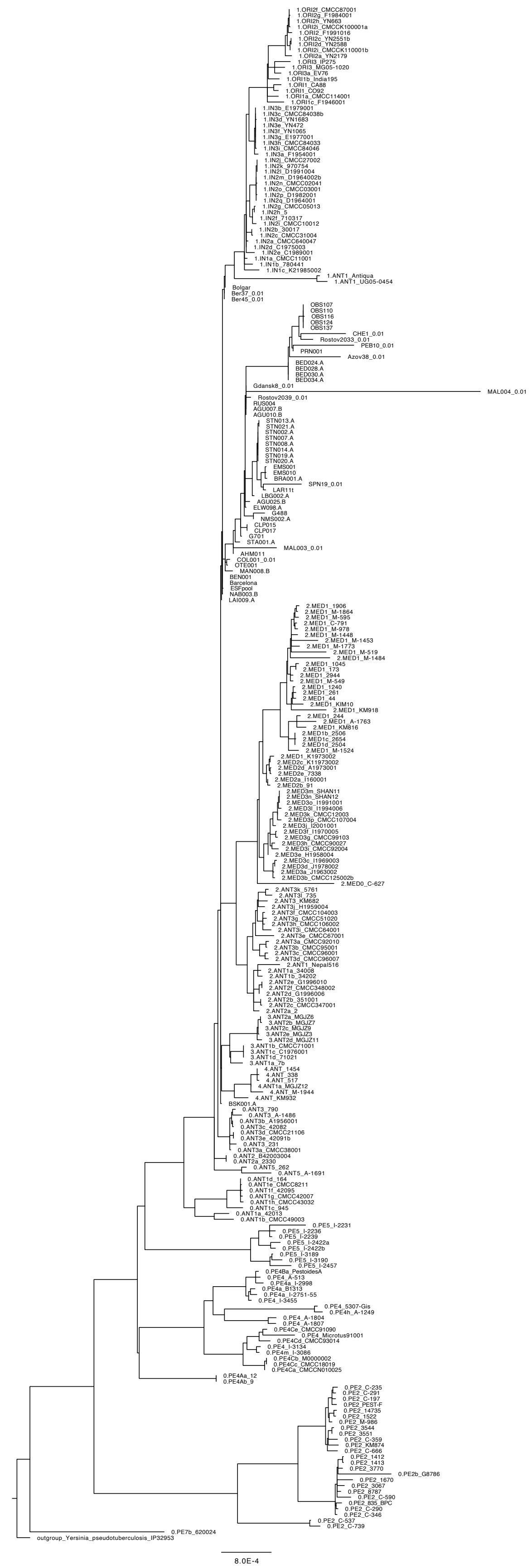

**Fig. S23: Phylogenetic tree with "0.01" treatment.**
Maximum likelihood tree generated with IQTree with 1000 bootstraps based on a 95% partial deletion SNP alignment (4033 positions) of 56 ancient *Y. pestis* genomes, including 12 genomes reconstructed from non-UDG data and with treatment "0.01", 208 modern *Y. pestis* genomes and *Y. pseudotuberculosis* IP32953 as outgroup.

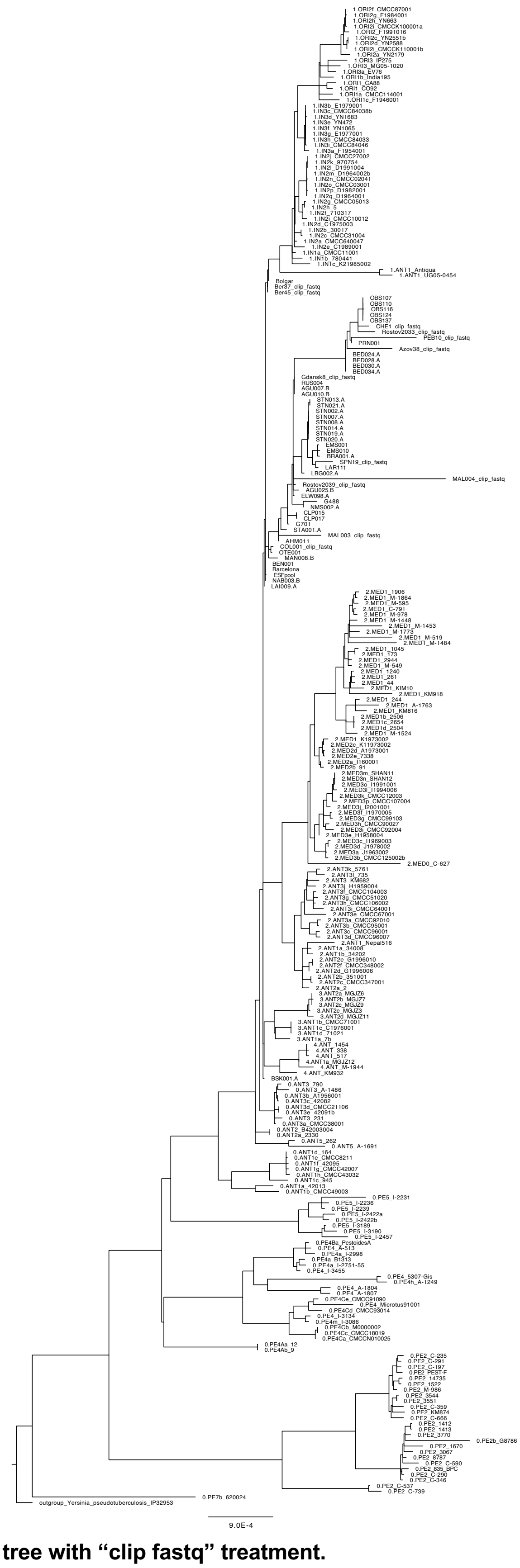

**Fig. S24: Phylogenetic tree with "clip fastq" treatment.**

Maximum likelihood tree generated with IQTree with 1000 bootstraps based on a 95% partial deletion SNP alignment (3736 positions) of 56 ancient *Y. pestis* genomes, including 12 genomes reconstructed from non-UDG data and with treatment "clip fastq", 208 modern *Y. pestis* genomes and *Y. pseudotuberculosis* IP32953 as outgroup.

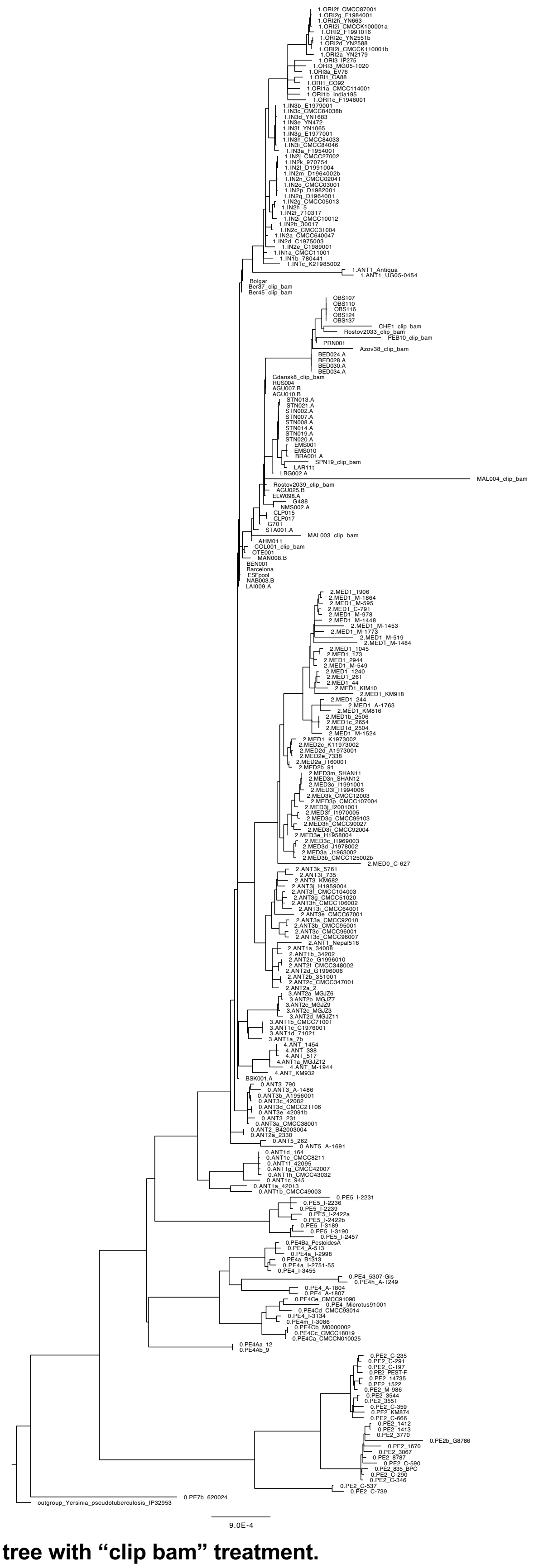

**Fig. S25: Phylogenetic tree with "clip bam" treatment.**

Maximum likelihood tree generated with IQTree with 1000 bootstraps based on a 95% partial deletion SNP alignment (3908 positions) of 56 ancient *Y. pestis* genomes, including 12 genomes reconstructed from non-UDG data and with treatment "clip bam", 208 modern *Y. pestis* genomes and *Y. pseudotuberculosis* IP32953 as outgroup.

### 5 Schematic tree

*Marcel Keller*

Since the maximum likelihood tree (Fig. 2B) was missing resolution on individual nodes (e.g. regarding the short, shared branch of MAL004 with NMS002.A and G488), we created a schematic tree after manually checking relevant phylogenetically informative positions with IGV (v2.8.13), where the original SNP table used for the phylogenetic analyses contained an ‘N’ (Fig. S27–30). The schematic tree as shown in Fig. 2A and Fig. S26 is based on shared SNPs when classified as phylogenetically informative and private SNPs, after filtering with SNPEvaluation (see main text, Material and Methods). For genomes reconstructed from non-UDG data, we determined the terminal branch lengths by comparing the initial bam file (bwa-aln -n 0.01, -l 16) with the remapped (bwa-aln -n 0.1, -l 32) bam files in SNPEvaluation, but flagged sites as potential damage if they are C>T or G>A substitution (in these cases shown as dashed lines in Fig. 2A and S26; see also Tables S5 and S6). In addition to the schematic tree, we established a new ad-hoc node numbering system for this study to allow facilitate the description of shared SNPs. Importantly, this numbering system is different from the system established by Cui et al. 2013 – the node termed N07 by Cui et al. 2013 corresponds to N01 in this study.

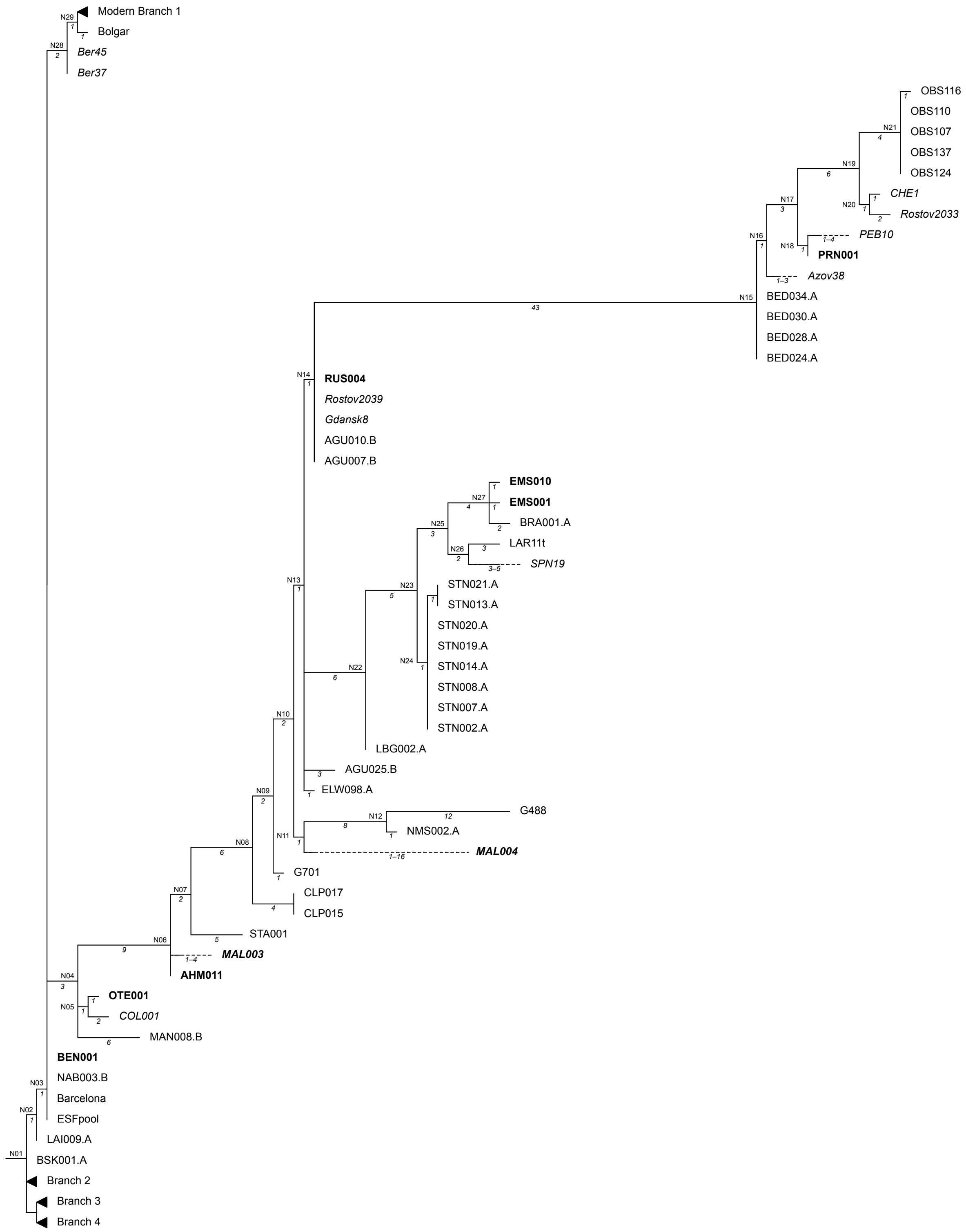

**Fig. S26: Schematic tree of the Second Pandemic.**
Manually drawn tree based on Table S6. Branch lengths correspond to numbers of SNPs. Dashed lines indicate SNPs on terminal branches that might be derived from deamination in case of non-UDG-derived genomes.

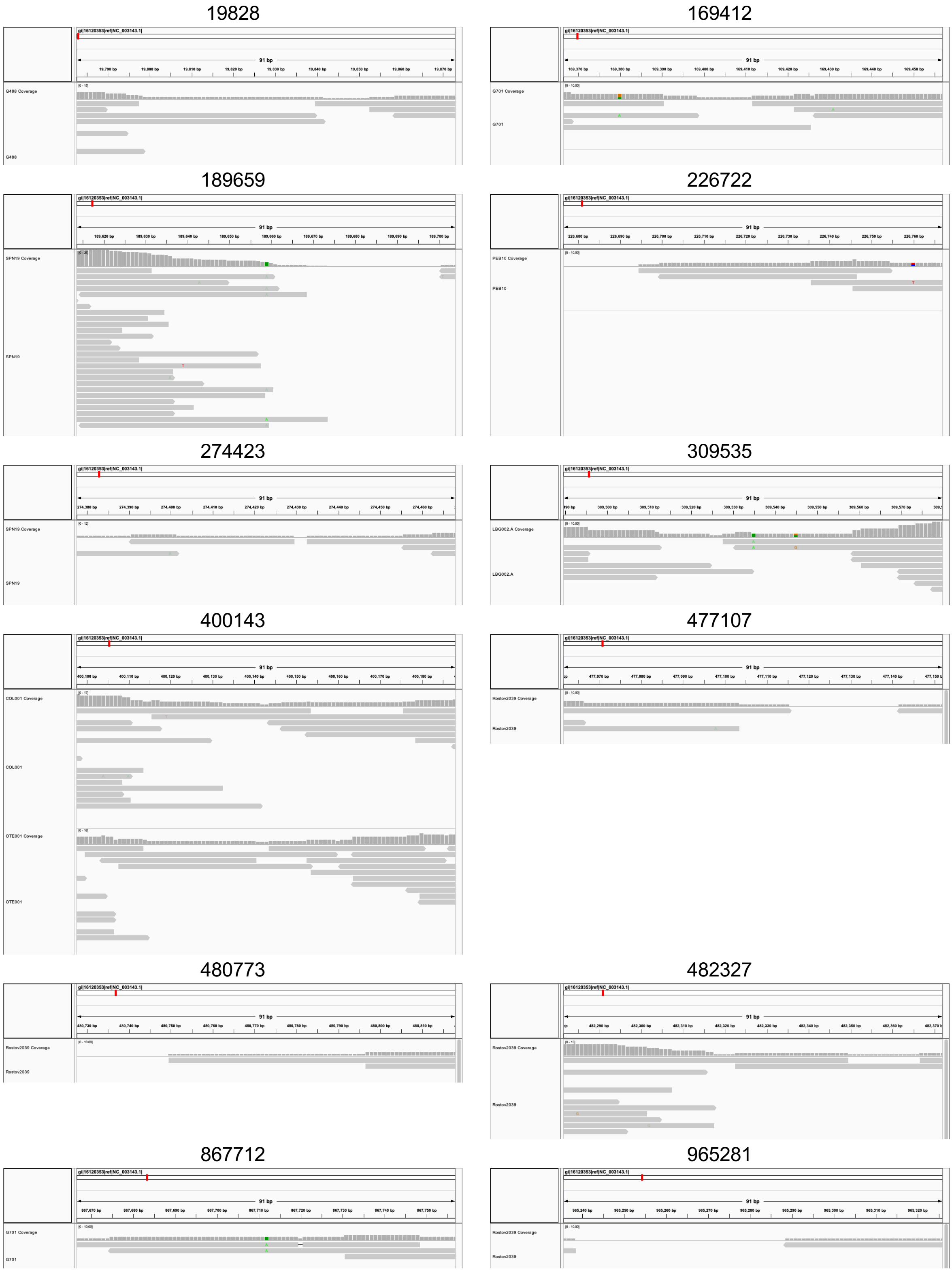

**Fig. S27: IGV Screenshots of 'N' positions manually checked for the schematic tree Figs. 2A and S11 (1).**

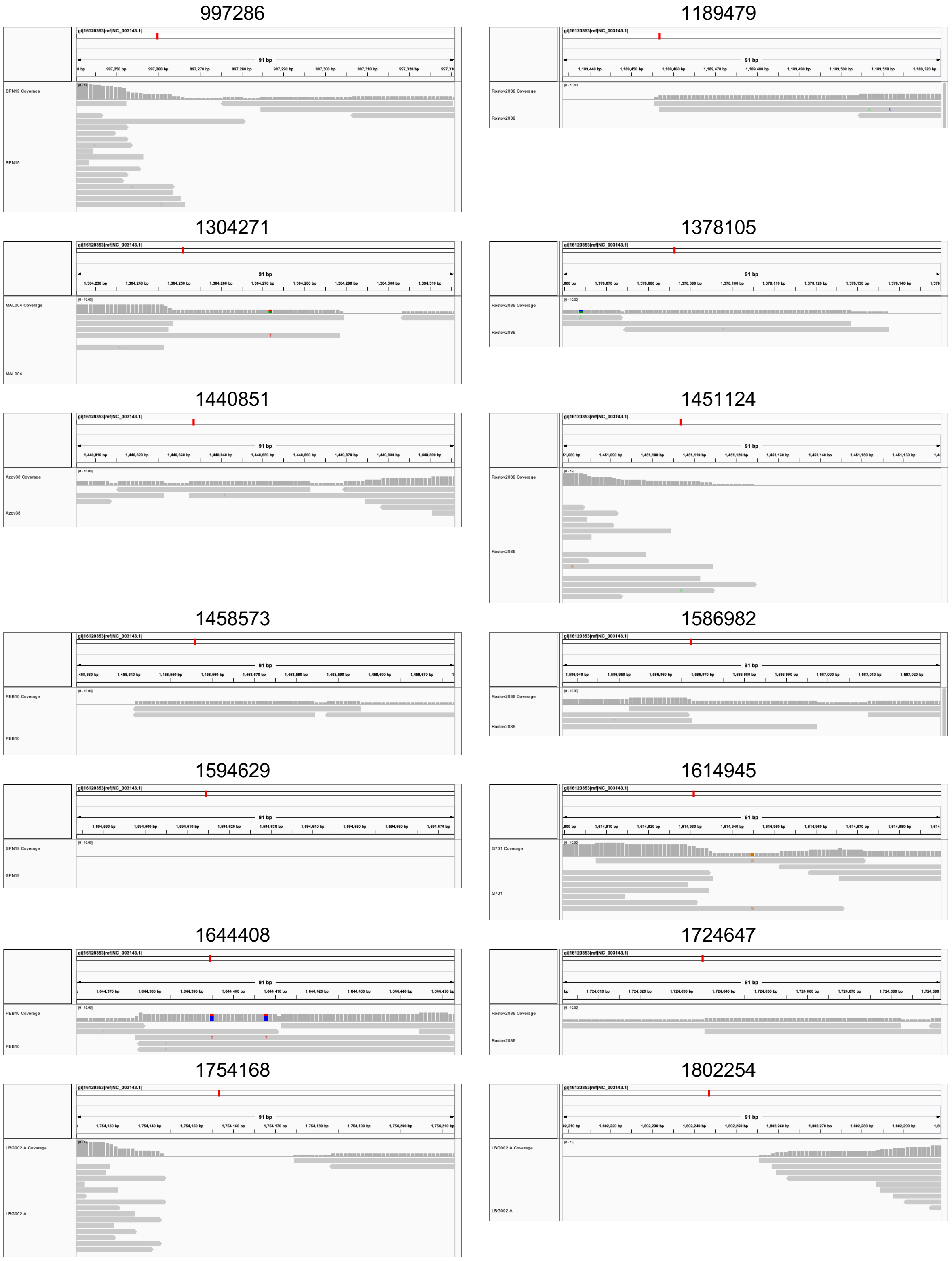

**Fig. S28: IGV Screenshots of 'N' positions manually checked for the schematic tree Figs. 2A and S11 (2).**

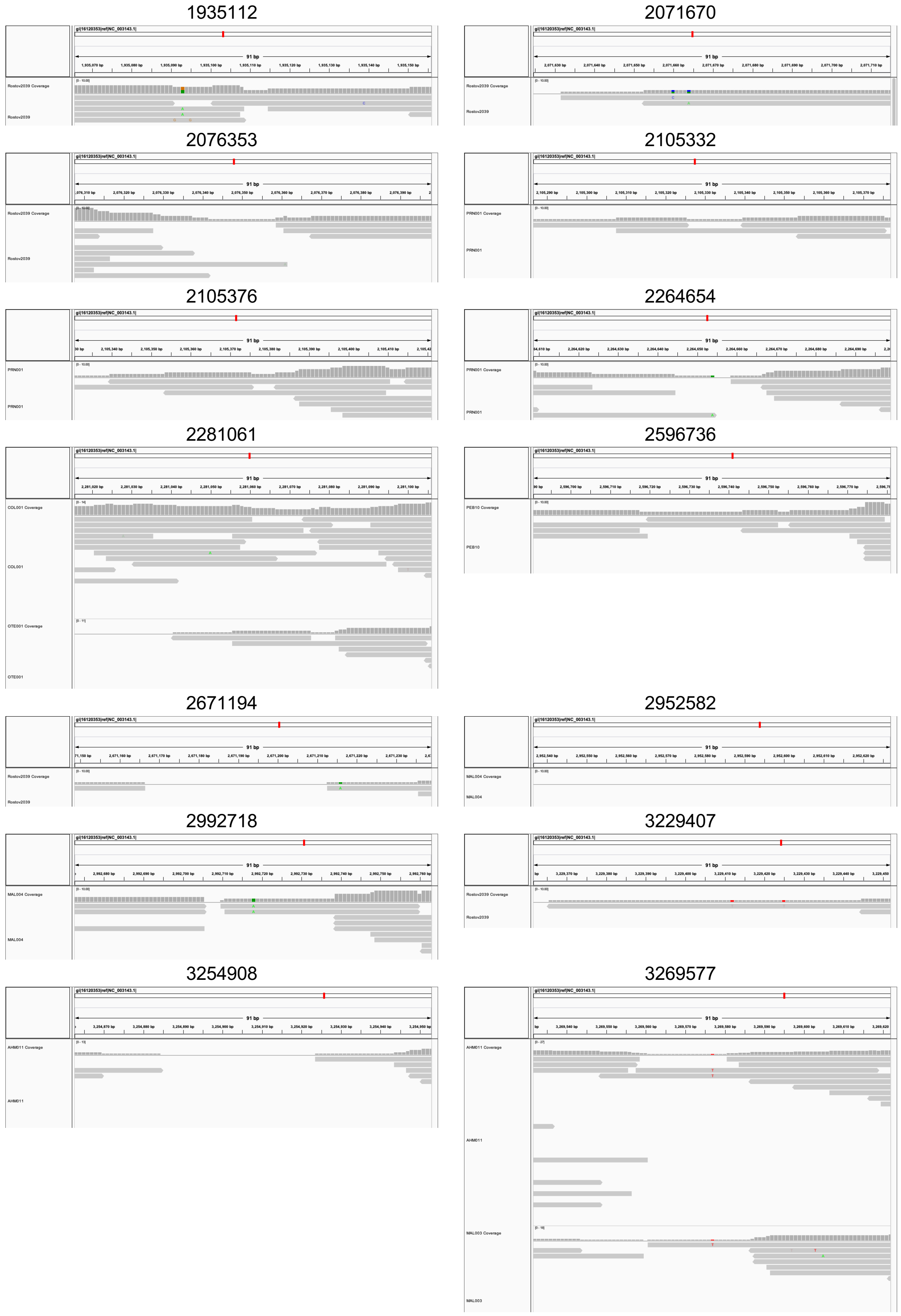

**Fig. S29: IGV Screenshots of ‘N’ positions manually checked for the schematic tree Figs. 2A and S11 (3).**

3610371

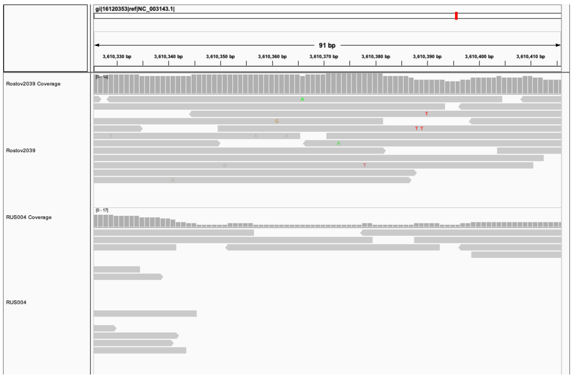

3872698

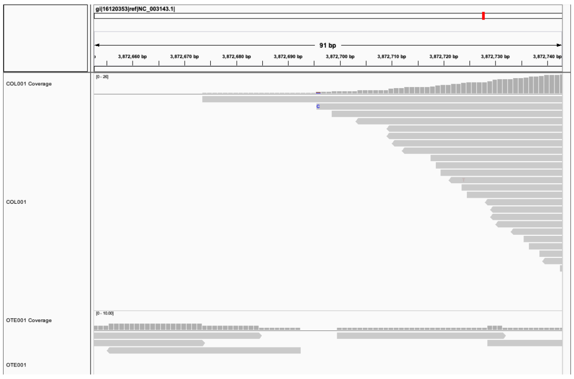

3944305

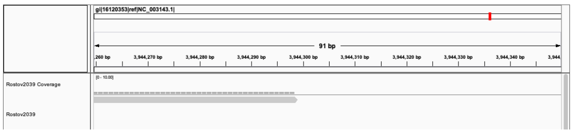

4134121

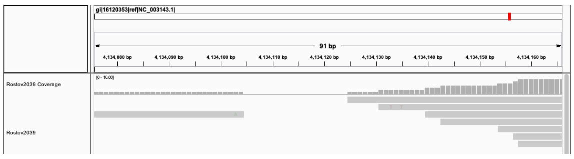

4190286

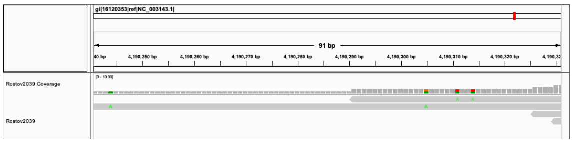

4208536_Rostov

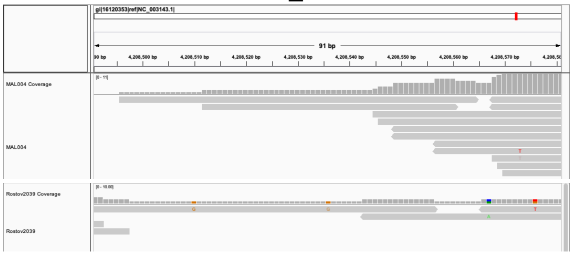

**Fig. S30: IGV Screenshots of ‘N’ positions manually checked for the schematic tree Figs. 2A and S11 (4).**

### 6 Heterozygosity plots

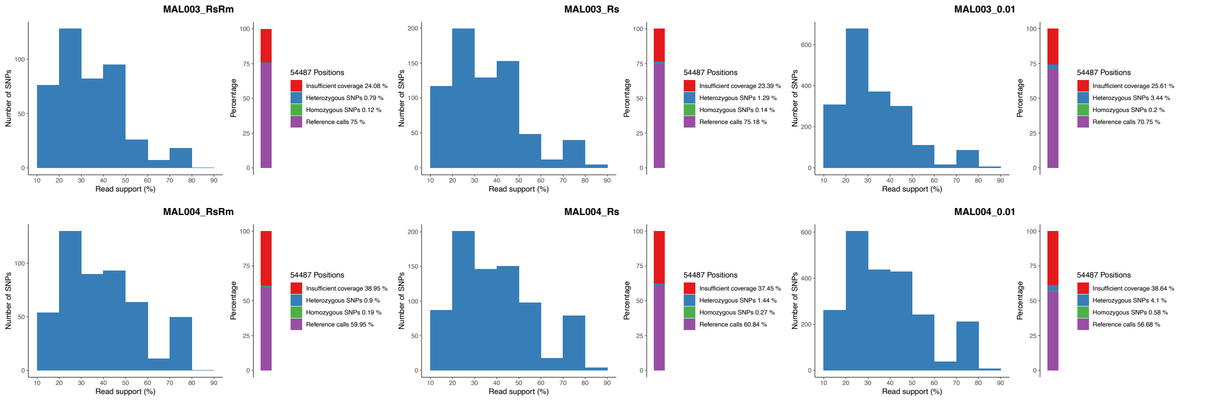

**Fig. S31: Heterozygosity plots for MAL003 and MAL004.**
Histograms of heterozygous positions (left within each panel) and percentages of reference calls, homozygous SNPs, heterozygous SNPs, and sites with insufficient coverage (right within each panel) for the newly sequenced nonUDG libraries MAL003 and MAL004 with “RsRm” treatment (left panels), “Rs” treatment (middle panels), and “0.01” treatment (right panels).

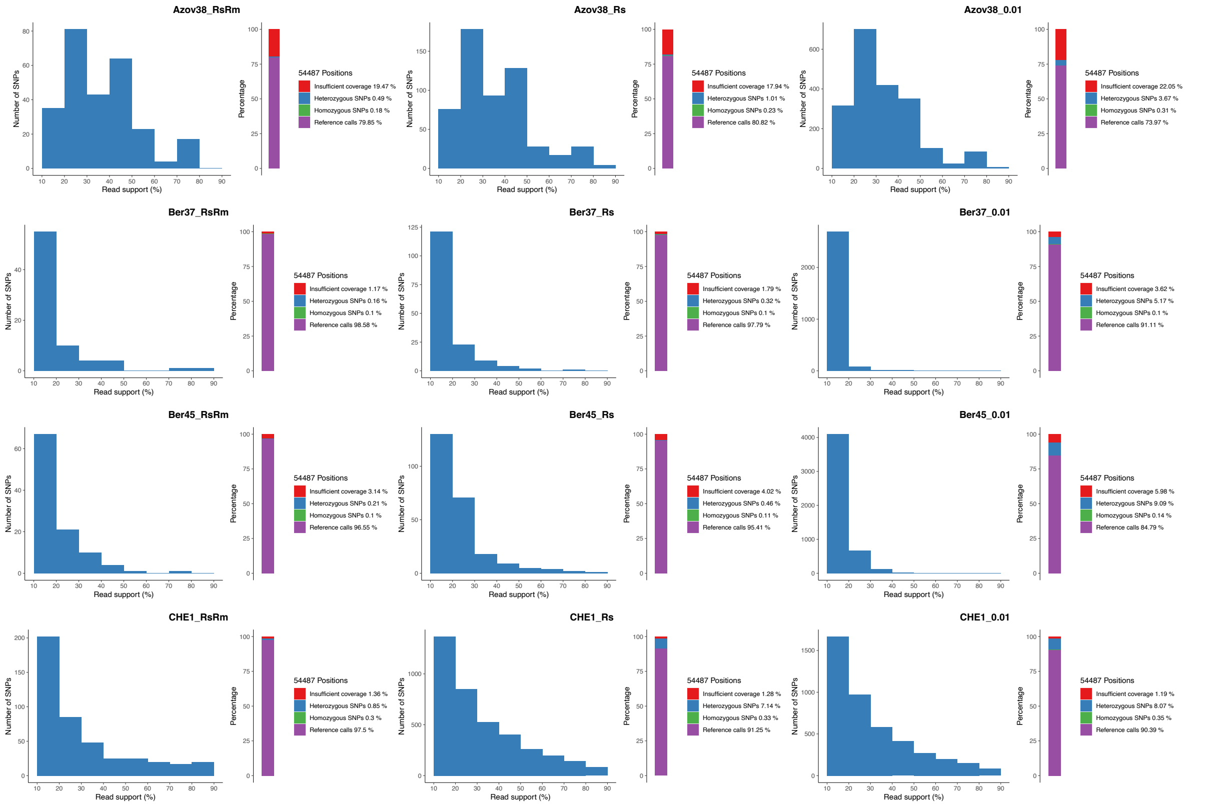

**Fig. S32: Heterozygosity plots for Azov38, Ber37, Ber45, and CHE1.**
Histograms of heterozygous positions (left within each panel) and percentages of reference calls, homozygous SNPs, heterozygous SNPs, and sites with insufficient coverage (right within each panel) for the previously published nonUDG libraries Azov38, Ber37, Ber45, and CHE1 with “RsRm” treatment (left panels), “Rs” treatment (middle panels), and “0.01” treatment (right panels).

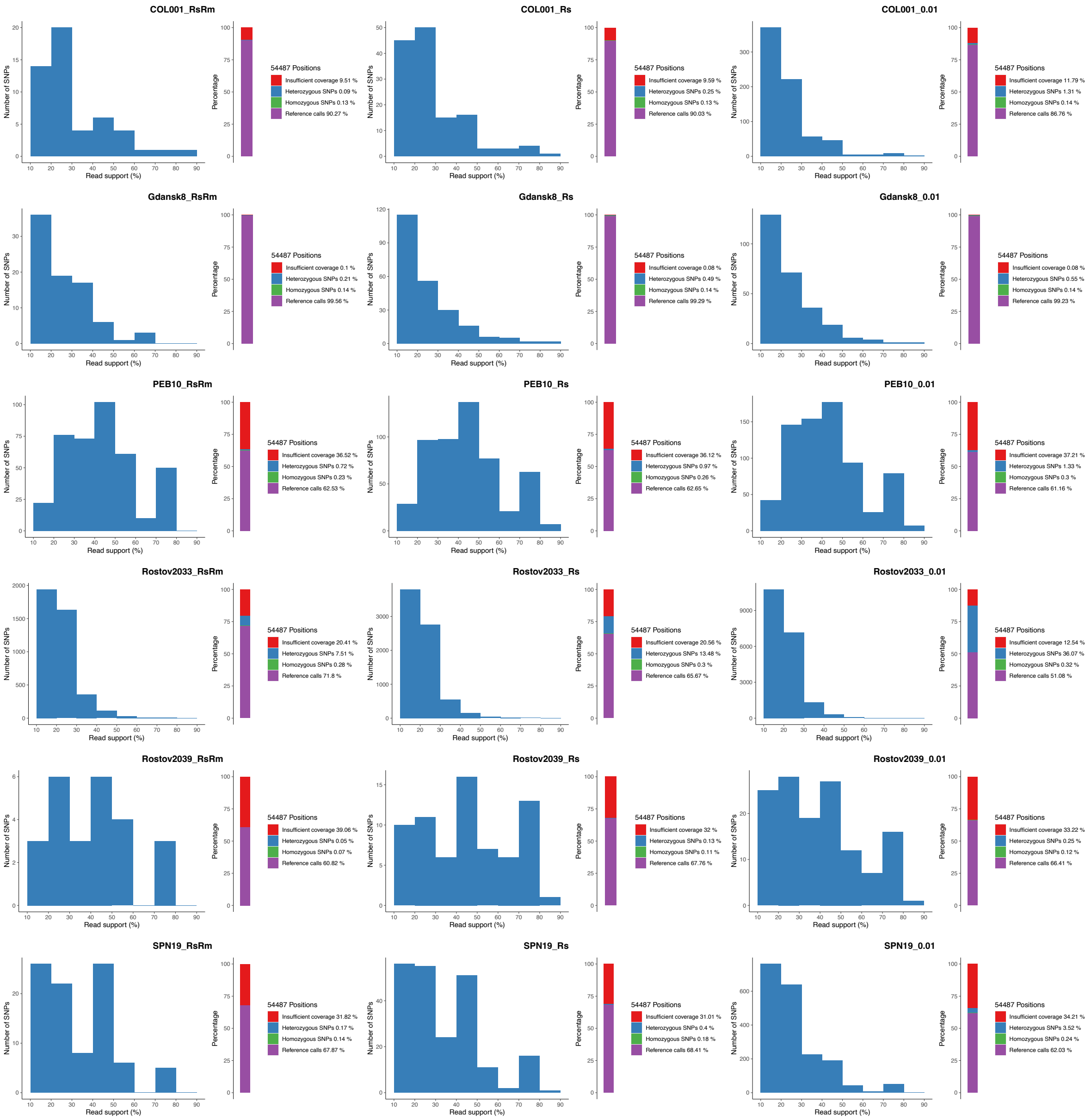

**Fig. S33: Heterozygosity plots for COL001, Gdansk8, PEB10, Rostov2033, Rostov2039, and SPN19.** Histograms of heterozygous positions (left within each panel) and percentages of reference calls, homozygous SNPs, heterozygous SNPs, and sites with insufficient coverage (right within each panel) for the previously published nonUDG libraries COL001, Gdansk8, PEB10, Rostov2033, Rostov2039, and SPN19 with “RsRm” treatment (left panels), “Rs” treatment (middle panels), and “0.01” treatment (right panels).

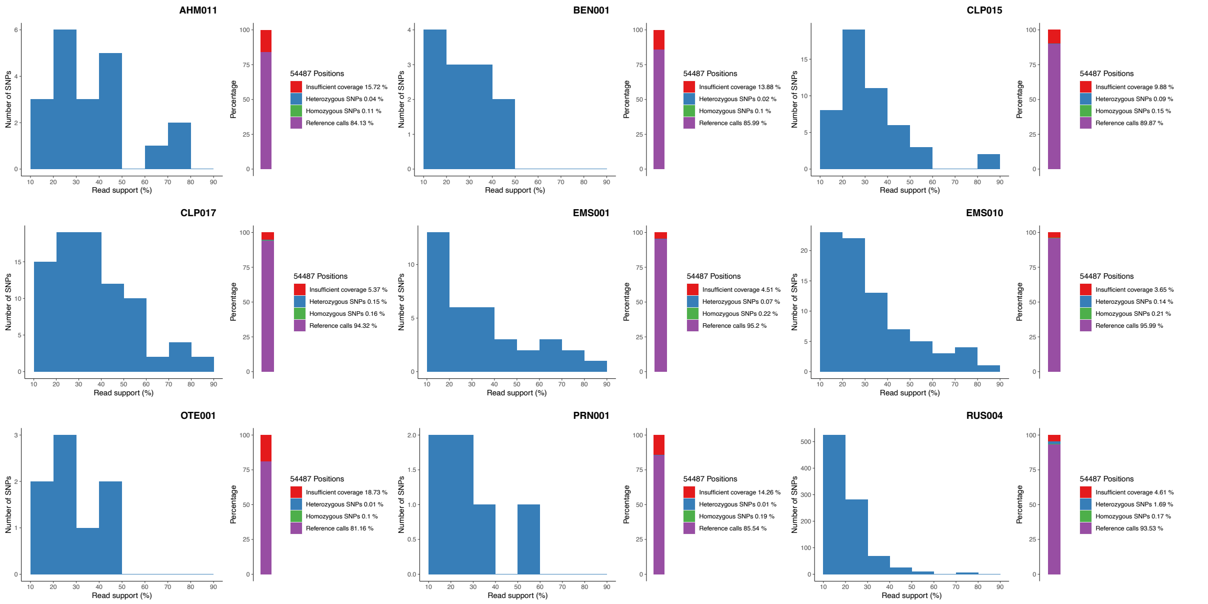

**Fig. S34: Heterozygosity plots for AHM011, BEN001, CLP015, CLP017, EMS001, EMS010, OTE001, PRN001, and RUS004.**

Histograms of heterozygous positions (left within each panel) and percentages of reference calls, homozygous SNPs, heterozygous SNPs, and sites with insufficient coverage (right within each panel) for the newly sequenced UDG libraries AHM011, BEN001, CLP015, CLP017, EMS001, EMS010, OTE001, PRN001, and RUS004.

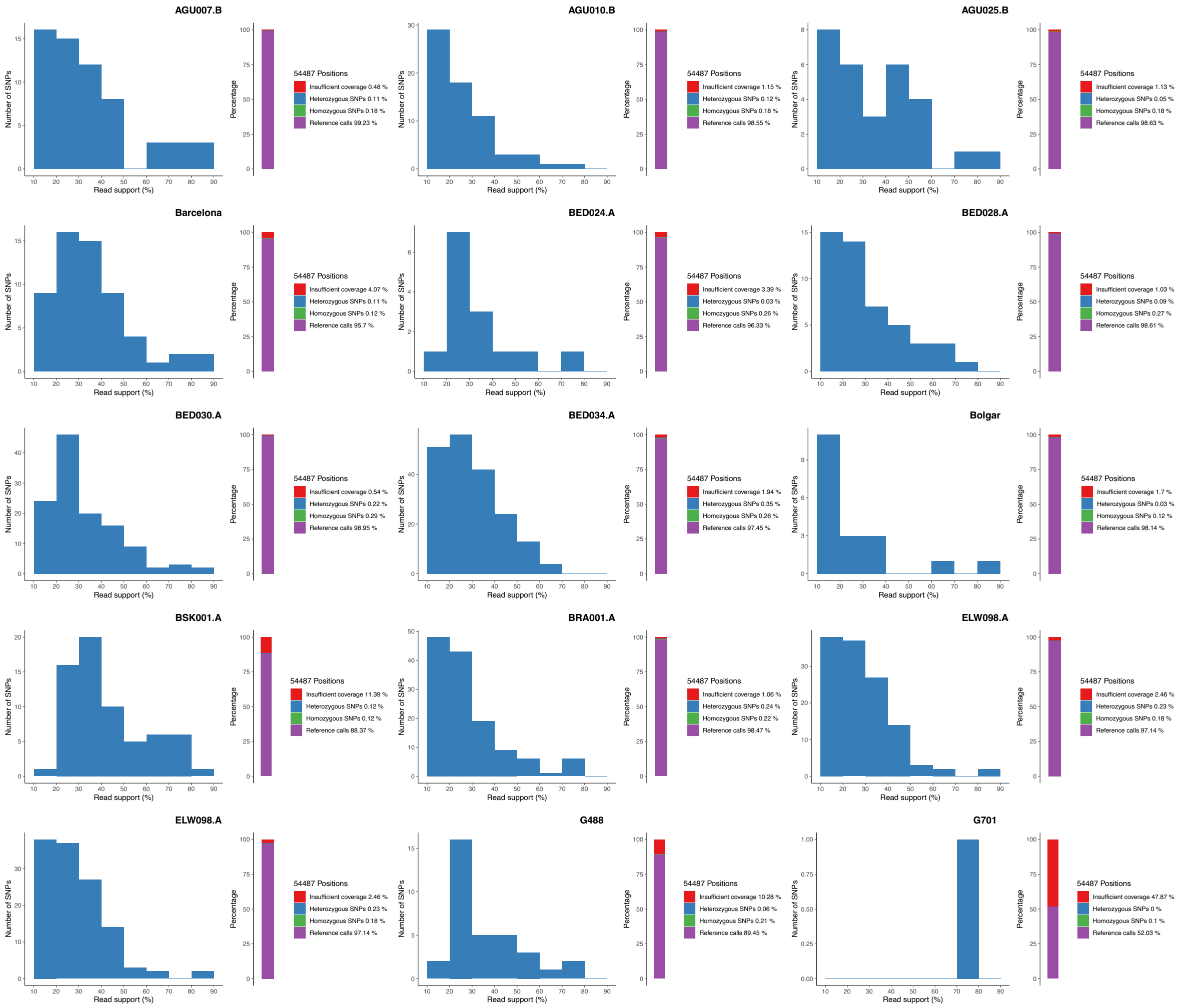

**Fig. S35: Heterozygosity plots for AGU007.B, AGU010.B, AGU025.B, Barcelona, BED024.A, BED028.A, BED030.A, BED034.A, Bolgar, BRA001.A, BSK001.A, ELW098.A, and ESFpool, G488, and G70.** Histograms of heterozygous positions (left within each panel) and percentages of reference calls, homozygous SNPs, heterozygous SNPs, and sites with insufficient coverage (right within each panel) for the previously published UDG libraries AGU007.B, AGU010.B, AGU025.B, Barcelona, BED024.A, BED028.A, BED030.A, BED034.A, Bolgar, BRA001.A, ELW098.A, and ESFpool; the halfUDG libraries G488, and G70; and the mixed dataset of BSK001.A.

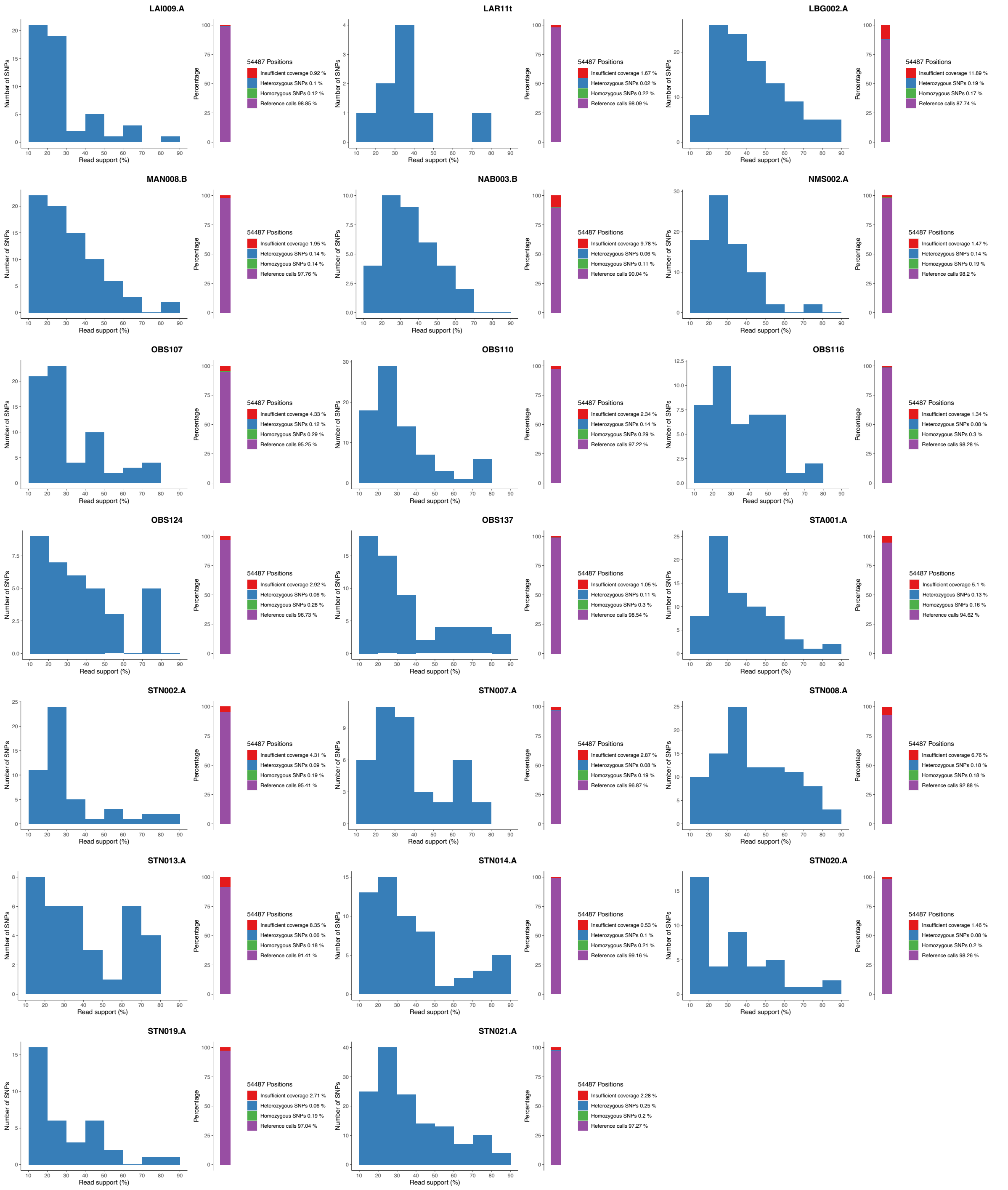

**Fig. S36: Heterozygosity plots for LAI009.A, LAR11t, LBG002.A, MAN008.B, NAB003.B, NMS002.A, OBS017, OBS110, OBS116, OBS124, OBS137, STA001.A, STN002.A, STN007.A, STN008.A, STN013.A, STN014.A, STN019.A, STN020.A, and STN021.A.**

Histograms of heterozygous positions (left within each panel) and percentages of reference calls, homozygous SNPs, heterozygous SNPs, and sites with insufficient coverage (right within each panel) for the previously published UDG libraries LAI009.A, LAR11t, LBG002.A, MAN008.B, NAB003.B, NMS002.A, OBS017, OBS110, OBS116, OBS124, OBS137, STA001.A, STN002.A, STN007.A, STN008.A, STN013.A, STN014.A, STN019.A, STN020.A, and STN021.A.

### 7 Newly sequenced low coverage genomes

*Marcel Keller*

To assess the phylogenetic positions of newly sequenced low coverage genomes, we generated a phylogenetic tree with newly sequenced genomes of a mean coverage higher than 1-fold. Fig. S37 shows the maximum likelihood tree generated with IQTree with 1000 bootstraps based on a full SNP alignment (4247 positions), including the genomes of BEN002, COM042, AHM001, MAL007, EMS003, PRN004, PRN005, PRN008, PRN009 and LHP001. The respective SNP table is shown in Table S16. A phylogenetic tree based on the full SNP alignment (without partial deletion) allows for a better resolution of the topology but is also susceptible to wrong placement of low coverage genomes due to long branch attraction/homoplastic sites. In addition, low coverage genomes are particularly susceptible to contamination by environmental bacteria with high sequence similarity since positions classified as ‘heterozygous’ (N) in higher coverage genomes can appear ‘homozygous’ because of missing coverage of the position with *Y. pestis* reads. Therefore, terminal branches and the responsible private SNPs were not considered.

The genomes of BEN002 and COM042 fall on very basal positions on Branch 1 in the phylogenetic tree (Fig. S37) and do not share any of the phylogenetically informative post-Black Death SNPs, implying that they are most likely identical with BEN001 and other European Black Death genomes.

AHM001 is most likely identical with AHM011, since it shares at least some of the N04–N06 defining SNPs (Table S16). The shared branch with MAL003 (Fig. S37) is an artifact and not based on exclusively shared SNPs between these two genomes as expected for an internal branch for a clonal species.

Since the site of Mäletjärve yielded two distinct genomes which most likely date to different periods, we investigated the position of MAL007 with more scrutiny. Figs. S38–39 show the positions leading from N06 (the branching node of MAL003) to N11 (the branching node of MAL004). All positions covered show the derived SNP ofMAL004, implying that MAL004 and MAL007 appear to be identical. The shared branch of MAL007 with NMS002.A and G488 is an artifact and not based on exclusively shared SNPs.

In the phylogenetic tree, RUS003 shared a short branch with ELW098.A, although not supported by shared SNPs in the SNP table (Table S16). Therefore, we investigated whether it is identical with RUS004 and the other basal Branch 1A1 genomes. Fig. S40 shows that the respective SNP is present in RUS003 as well, implying that RUS003 and RUS004 are most likely identical.

For the genomes of Domat/Ems, only EMS003 had a sufficient coverage for integration in the phylogenetic tree, where it falls close to EMS010. However, when investigating the private SNPs of EMS001 and EMS010, none of them appeared in either EMS003 or EMS002 (Fig. S41). Therefore, it implies that EMS003 and EMS002 are most likely basal to EMS001 and EMS10, falling on node N27. Alternatively, considering the microdiversity present at this site, few private SNPs forming terminal branches cannot be excluded.

For the site of Tallinn and the close-by site of Lehmja, the genomes PRN004, PRN005, PRN008, PRN009 (Tallinn) and LHM001 (Lehmja) had enough coverage to be integrated into the phylogenetic tree (Fig. S37), where they cluster together with PRN001. Two positions are exclusively shared among low coverage genomes of Tallinn, but both appear in local peaks of coverage, showing conflicting reads without the SNP (1737499) or 'heterozygous' positions nearby (4198695), and are therefore dismissed. Again, it implies that all genomes of Tallinn and the genome LHM001 of Lehmja are most likely identical.

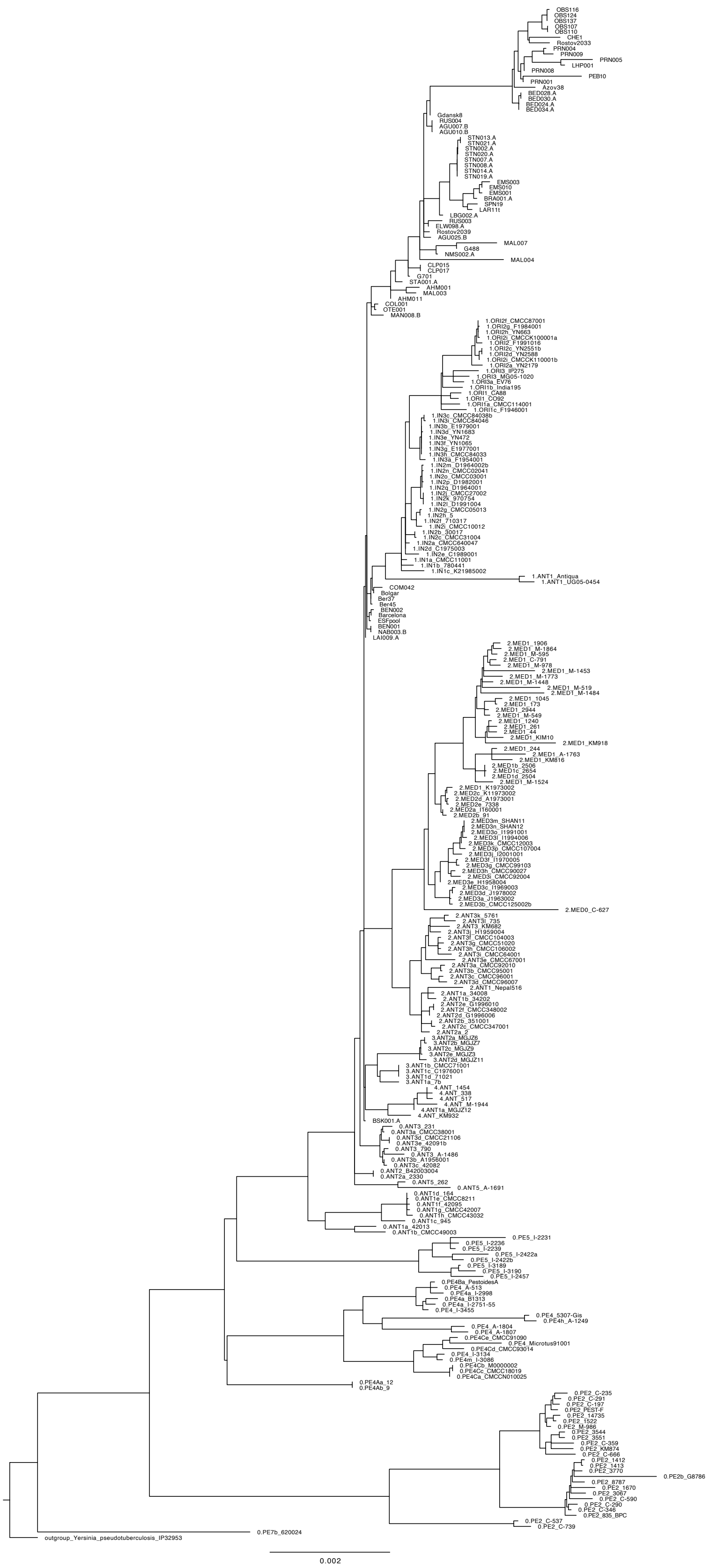

**Fig. S37: Phylogenetic tree including newly sequenced low coverage genomes (>1x)**
Maximum likelihood tree generated with IQTree with 1000 bootstraps based on a full SNP alignment (4247 positions) of 67 ancient *Y. pestis* genomes (including 11 low coverage genomes), 208 modern *Y. pestis* genomes and *Y. pseudotuberculosis* IP32953 as outgroup.

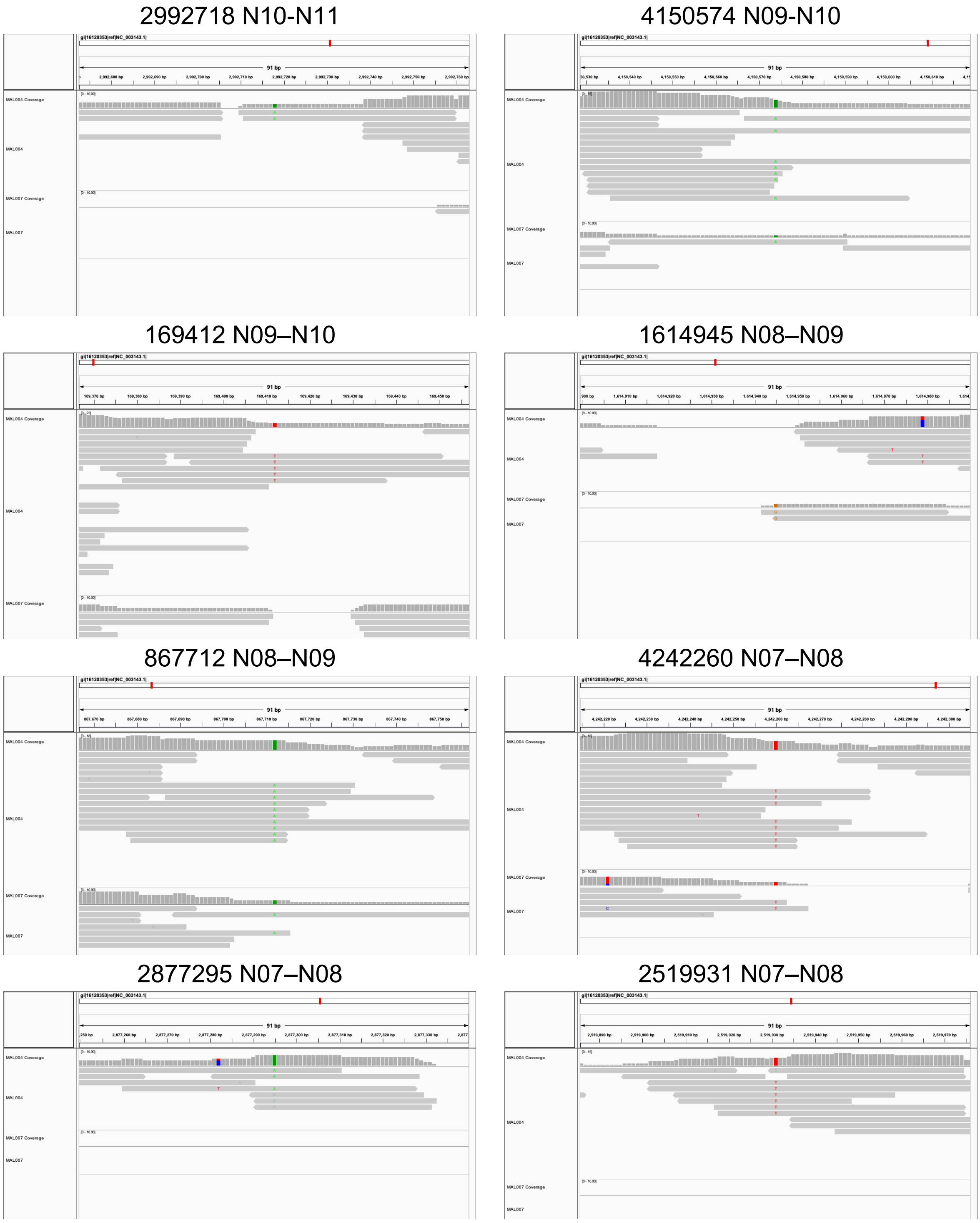

**Fig. S38: IGV screenshots of MAL004 and MAL007 for positions leading from N06–N11 (1)**
Shown are the respective bam-files of the mappings with ‘RsRm’ treatment, as used for the phylogenetic analyses.

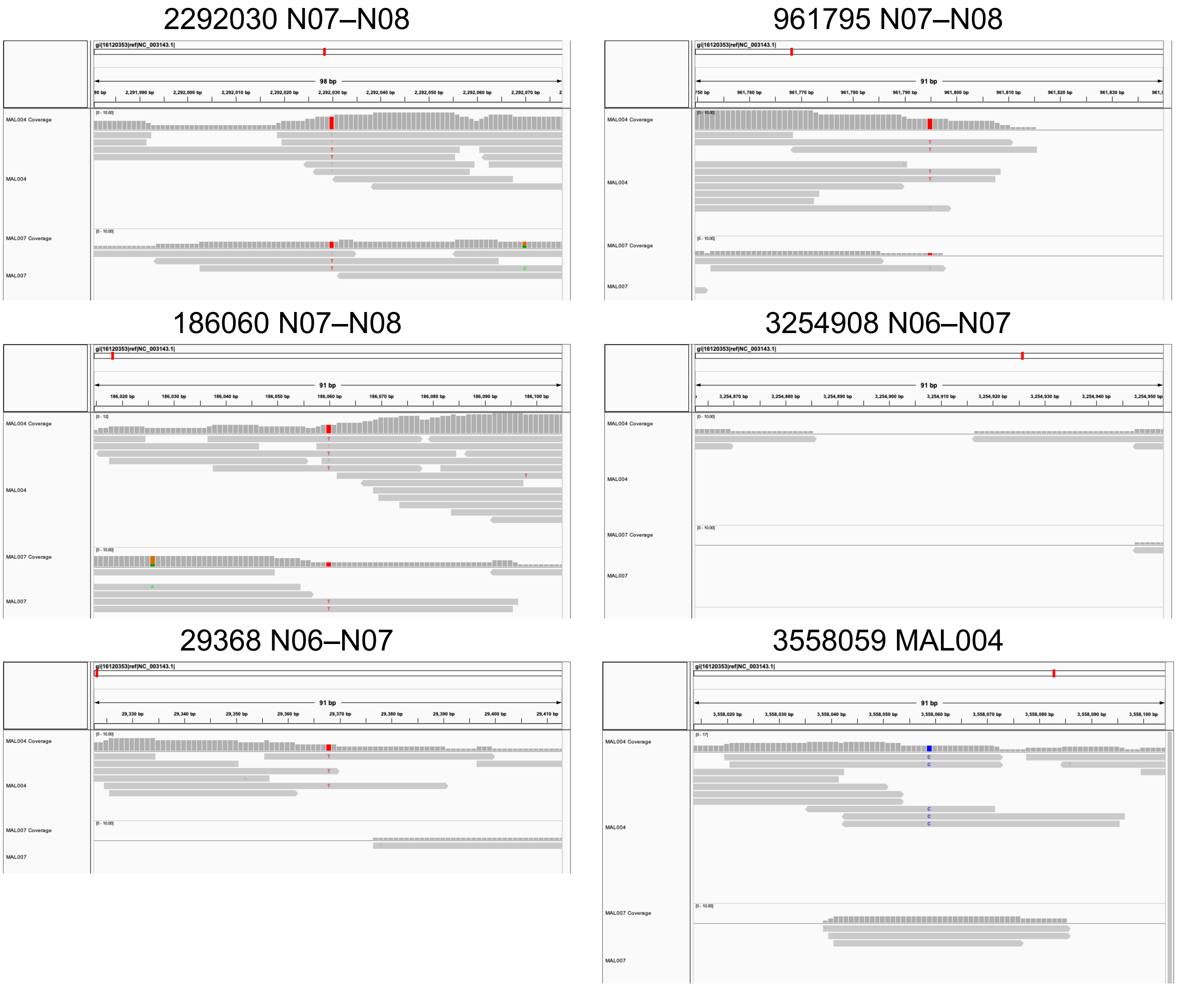

**Fig. S39: IGV screenshots of MAL004 and MAL007 for positions leading from N06–N11 (2)**
Shown are the respective bam-files of the mappings with ‘RsRm’ treatment, as used for the phylogenetic analyses.

4208536 N13–N14

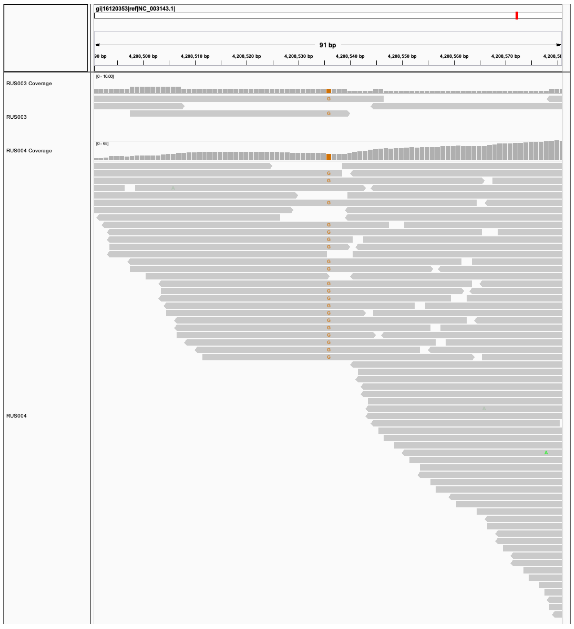

**Fig. S40: IGV screenshot of RUS003 and RUS004 for the SNP defining the basal Branch 1A1 cluster**

1526655 N27–EMS001

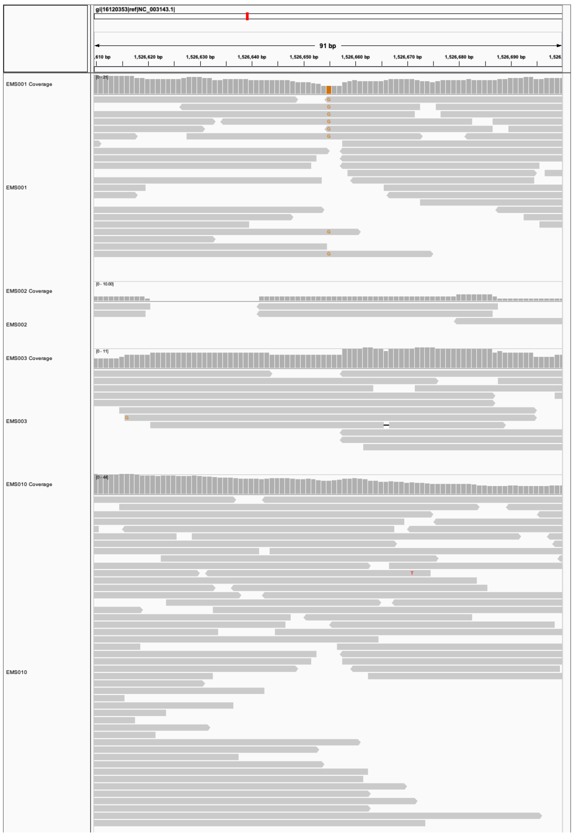

2839674 N27–EMS010

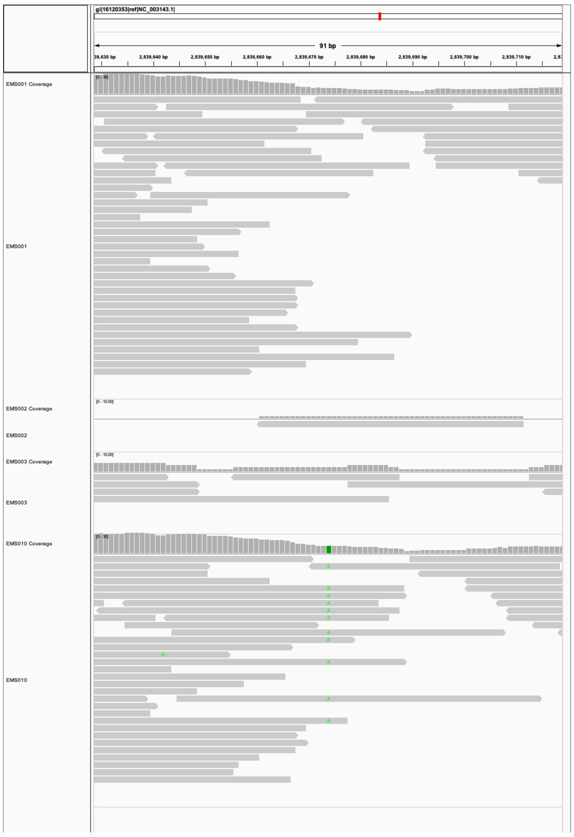

**Fig. S41: IGV screenshot of EMS001, EMS002, EMS003 and EMS010 for the private SNPs of EMS001 and EMS010.**

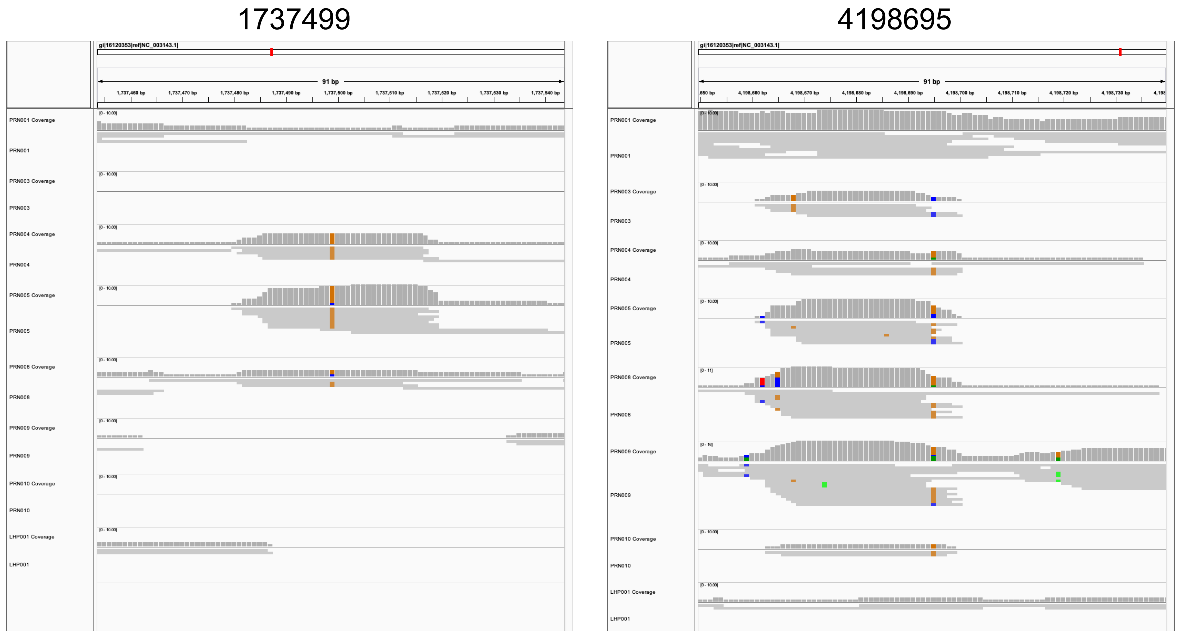

**Fig. S42: IGV screenshot of PRN001, PRN003, PRN004, PRN005, PRN008, PRN009 and PRN010 for two positions exclusively shared among low coverage PRN genomes.**

### 8 Deletion analysis and functional analysis of newly identified SNPs

*Marcel Keller*

To check for previously observed or novel deletions in ancient genomes of the Second Pandemic, we generated mappings without filtering for mapping quality in combination with bedtools and respective annotation files within the nf-core/eager (v2.3.0) pipeline. The list of virulence-related genes was adapted from Andrades Valtueña et al. 2022. The pseudogenes YPO0389/YPO0392a and YPO0394a were excluded since they did not appear in the used annotation file. The (pseudo)genes YPO1986-YPO1987, YPO2096-YPO2135, YPO2487-YPO2489, YPO3046-YPO3047 and yadA were evaluated individually or as individual copies. The results are shown in Table S13 and visualized as heatmap in Fig. S43. The previously described (Spyrou et al. 2019) large deletion, including the genes *mgtB* and *mgtC* in younger genomes of the Branch 1A1, is also observed in the genome PRN001, falling into the same clade.

The deletion of *inv*, previously described for NMS002.A (Spyrou et al. 2019), appears also in G488 and the newly reconstructed genome of MAL004, further confirming its phylogenetic position sharing a short branch with the two aforementioned genomes (see Fig. 2).
We also performed a functional analysis on all SNPs of the Second Pandemic genomes with SNPEff (v3.1), the results are shown in Table S14.

Eight private SNPs of MAL004 were classified as potential false-positive SNPs due to damage and fall into coding regions causing nonsynonymous mutations (ftsW, YPO1362, lepA, YPO2756, YPO3146, ibeB, mtta2, glpB). ftsW is a peptidoglycan glycosyltransferase and as part of the divisome important for cell division; YPO1362 is a putative ATP-dependent endonuclease of the OLD family; lepA is a ribosomal elongation factor and part of the T6SS locus (Andersson et al. 2017); YPO2756 is a tRNA 5-methylaminomethyl-2-thiouridine biosynthesis bifunctional protein; YPO3146 is a probable AsnC-family regulatory protein; ibeB is a probable outer membrane efflux lipoprotein; mtta2 is a Sec-independent protein translocase protein TatB; glpB is a putative anaerobic glycerol-3-phosphate dehydrogenase subunit B. The genomes of CLP015 and CLP017 show a nonsynonymous mutation in YPO1033, a hypothetical pyrimidine/purine-5'-nucleotide nucleosidase. For the genome of OTE001, a nonsynonymous mutation in the gene rrmA was observed, a 23S rRNA (guanine745-N1)-methyltransferase.

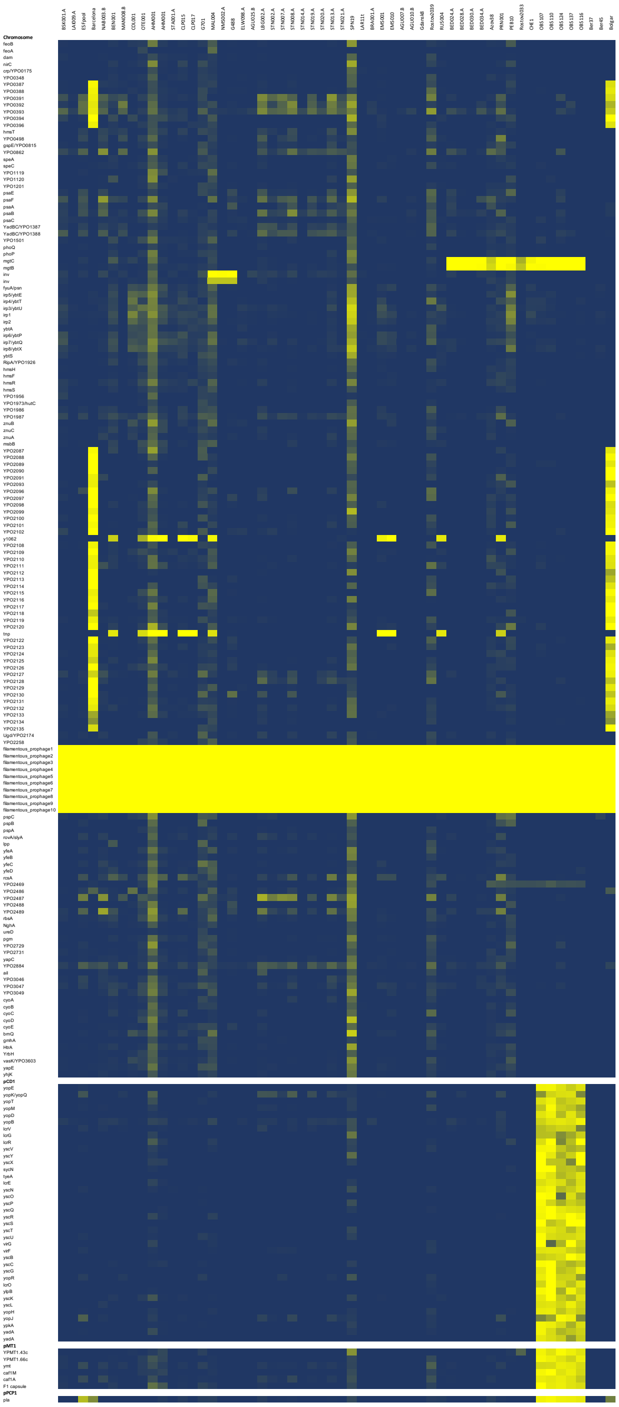

**Fig. S43: Heatmap showing breadth of coverage over virulence-associated genes on the chromosome and the plasmids pCD1, pMT1 and pPCP1 (yellow: no coverage; dark blue: gene fully covered).**

### *9* Coinfection with *Yersinia pestis* and *Treponema pallidum*

*Meriam Guellil, Marcel Keller*

The metagenomic screening with Kraken-Uniq/Kraken2 (see main text, Material and Methods) revealed the sample PRN008 (associated with Great Northern War plague burial and dated to 1710; see Supplementary Section 11.13) not only to be positive for *Y. pestis* but also *Treponema pallidum* (Table S1). To verify the screening results, we performed individual and competitive mappings to the reference sequences *Treponema pallidum ssp. pallidum* Nichols (NC_021490.2, causative agent of syphilis), *Treponema pallidum ssp. pertenue* CDC2 (NC_016848.1, causative agent of yaws), and *Treponema denticola* ATCC 35405 (AE017226.1, a closely related oral pathogen). Mappings were performed on the non-UDG shotgun data with bwa-aln (v0.7.17, -n 0.01, -l 16) with deduplication (DeDup v0.12.7), filtering for mapping quality 30 and damage calculation with mapDamage2 (Jónsson et al. 2013).

The competitive mappings of *T. denticola* vs. *T. pallidum pallidum* (75 vs. 1173 uniquely mapped reads), *T. denticola* vs. *T. pallidum pertenue* (75 vs. 1182 uniquely mapped reads) as well as the individual mapping data and the edit distance support the identification of a *T. pallidum ssp.* strain instead of the oral pathogen *T. denticola* (Fig. S44). Due to the low coverage and the high similarity of the *T. pallidum pallidum* and *T. pallidum pertenue*, we did not attempt a further characterization.

With this case, we present the second coinfection of plague and treponematosis identified through ancient DNA. The first case of a coinfection with *Treponema pallidum pertenue* was reported in Giffin et al. 2020 reported a in an individual from Vilnius, Lithuania, dating to the second half of the 15th century.

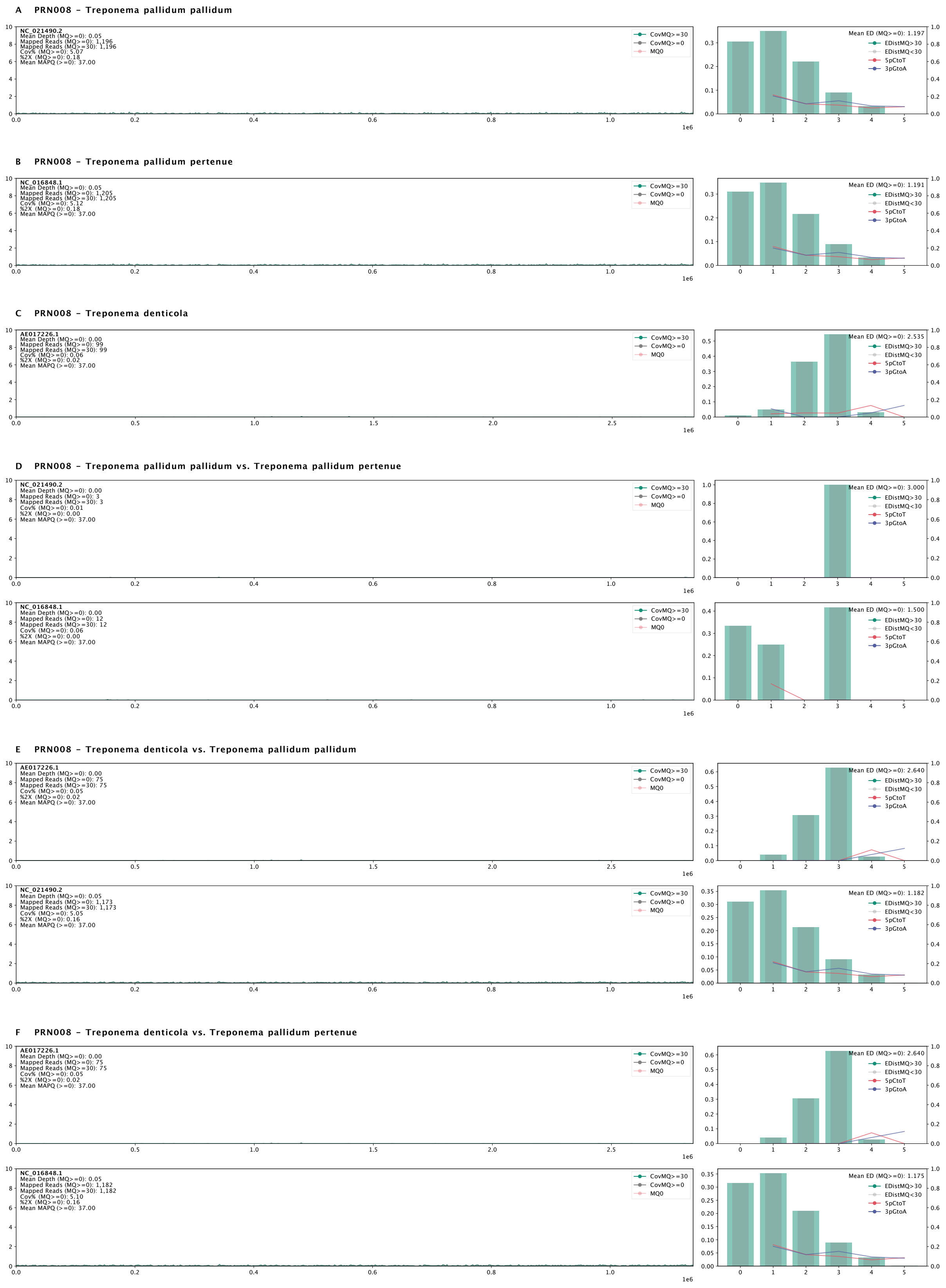

**Fig. S44: Mapping plots of PRN008 for *T. pallidum pallidum, T. pallidum pertenue* and *T. denticola.***
Mappings against the reference sequences of *T. pallidum pallidum* (A); *T. pallidum pertenue* (B); *T. denticola* (C); and for competitive mappings of *T. pallidum pallidum* vs. *T. pallidum pertenue* (D); *T. denticola* vs. *T. pallidum pallidum* (E); and *T. denticola* vs. *T. pallidum pertenue* (F). The panels on the left show the coverage distributed over the reference sequence, the right panels the edit distance (bar plots) and deaminiation damage (line plots).

### 10 Radiocarbon Modelling

**Code S1: Oxcal code for individually calibrated dates**

```
Plot()
{
 R_Date("MAN008", 653, 22);
 R_Date("OTE001", 709, 24);
 R_Date("AHM001", 579, 25);
 R_Date("AHM011", 616, 23);
 R_Date("MAL003", 445, 24);
 R_Date("STA001", 417, 42);
 R_Date("CLP015", 505, 22);
 R_Date("CLP017", 563, 25);
 R_Date("MAL004", 363, 25);
 R_Date("MAL007", 351, 24);
 R_Date("NMS002", 404, 26);
 R_Date("G488", 353, 27);
 R_Date("ELW098", 350, 22);
 R_Date("AGU025", 405, 18);
 R_Date("LBG005", 367, 22);
 R_Date("LBG007", 361, 22);
 R_Date("LBG002", 358, 22);
 R_Date("EMS010", 426, 22);
 R_Date("EMS001", 411, 28);
 R_Date("EMS_17", 361, 22);
 R_Date("EMS_99A", 360, 22);
 R_Date("STN001", 349, 30);
 R_Date("STN002", 327, 30);
 R_Date("STN005", 313, 30);
 R_Date("STN006", 351, 29);
 R_Date("STN011", 326, 33);
 R_Date("STN012", 367, 30);
 R_Date("STN_I_101", 375, 37);
 R_Date("STN015", 334, 35);
 R_Date("STN017", 368, 35);
 R_Date("STN028", 326, 30);
 R_Date("STN029", 341, 32);
 R_Date("STN030", 321, 18);
 R_Date("STN_II_107", 369, 31);
 R_Date("STN031", 313, 18);
 R_Date("STN032", 343, 18);
 R_Date("Gdansk8", 443, 20);
 R_Date("AGU010", 426, 19);
 R_Date("RUS003", 381, 28);
 R_Date("RUS004", 377, 27);
 R_Date("AGU007", 353, 18);
 R_Date("BED_8147", 442, 43);
 R_Date("BED_8198", 424, 39);
 R_Date("BED_8216", 370, 33);
 R_Date("BED_8219", 355, 31);
};
```

**Code S2: Oxcal code for individually calibrated dates with ΔR (30, 20)**

```
Delta_R("MAN008_R",30,20);
R_Date("MAN008", 653, 22);
Delta_R("OTE001_R",30,20);
R_Date("OTE001", 709, 24);
Delta_R("AHM001_R",30,20);
R_Date("AHM001", 579, 25);
Delta_R("AHM011_R",30,20);
R_Date("AHM011", 616, 23);
Delta_R("MAL003_R",30,20);
R_Date("MAL003", 445, 24);
Delta_R("STA001_R",30,20);
R_Date("STA001", 417, 42);
Delta_R("CLP015_R",30,20);
R_Date("CLP015", 505, 22);
Delta_R("CLP017_R",30,20);
R_Date("CLP017", 563, 25);
Delta_R("MAL004_R",30,20);
R_Date("MAL004", 363, 25);
Delta_R("MAL007_R",30,20);
R_Date("MAL007", 351, 24);
Delta_R("NMS002_R",30,20);
R_Date("NMS002", 404, 26);
Delta_R("G488_R",30,20);
R_Date("G488", 353, 27);
Delta_R("ELW0098_R",30,20);
R_Date("ELW098", 350, 22);
Delta_R("AGU025_R",30,20);
R_Date("AGU025", 405, 18);
Delta_R("LBG005_R",30,20);
R_Date("LBG005", 367, 22);
Delta_R("LBG007_R",30,20);
R_Date("LBG007", 361, 22);
Delta_R("LBG002_R",30,20);
R_Date("LBG002", 358, 22);
Delta_R("EMS010_R",30,20);
R_Date("EMS010", 426, 22);
Delta_R("EMS001_R",30,20);
R_Date("EMS001", 411, 28);
Delta_R("EMS_17_R",30,20);
R_Date("EMS_17", 361, 22);
Delta_R("EMS_99A_R",30,20);
R_Date("EMS_99A", 360, 22);
Delta_R("STN001_R",30,20);
R_Date("STN001", 349, 30);
Delta_R("STN002_R",30,20);
R_Date("STN002", 327, 30);
Delta_R("STN_005_R",30,20);
R_Date("STN005", 313, 30);
Delta_R("STN006_R",30,20);
R_Date("STN006", 351, 29);
Delta_R("STN011_R",30,20);
R_Date("STN011", 326, 33);
Delta_R("STN012_R",30,20);
R_Date("STN012", 367, 30);
Delta_R("STN_I_101_R",30,20);
R_Date("STN_I_101", 375, 37);
```

```
Delta_R("STN015_R",30,20);
R_Date("STN015", 334, 35);
Delta_R("STN017_R",30,20);
R_Date("STN017", 368, 35);
Delta_R("STN028_R",30,20);
R_Date("STN028", 326, 30);
Delta_R("STN029_R",30,20);
R_Date("STN029", 341, 32);
Delta_R("STN030_R",30,20);
R_Date("STN030", 321, 18);
Delta_R("STN_II_107_R",30,20);
R_Date("STN_II_107", 369, 31);
Delta_R("STN031_R",30,20);
R_Date("STN031", 313, 18);
Delta_R("STN032_R",30,20);
R_Date("STN032", 343, 18);
Delta_R("Gdansk8_R",30,20);
R_Date("Gdansk8", 443, 20);
Delta_R("AGU010_R",30,20);
R_Date("AGU010", 426, 19);
Delta_R("RUS003_R",30,20);
R_Date("RUS003", 381, 28);
Delta_R("RUS004_R",30,20);
R_Date("RUS004", 377, 27);
Delta_R("AGU007_R",30,20);
R_Date("AGU007", 353, 18);
Delta_R("BED_8147_R",30,20);
R_Date("BED_8147", 442, 43);
Delta_R("BED_8198_R",30,20);
R_Date("BED_8198", 424, 39);
Delta_R("BED_8216_R",30,20);
R_Date("BED_8216", 370, 33);
Delta_R("BED_8219_R",30,20);
R_Date("BED_8219", 355, 31);
```

**Code S3: Oxcal sequence model using KDE_Plots and ΔR (30, 20)**

```
Sequence()
{
 C_Date("BD",AD(1350),2);
 Boundary("T1");
 Phase()
 {
  Delta_R("MAN008_R",30,20);
  R_Date("MAN008", 653, 22);
  Delta_R("OTE001_R",30,20);
  R_Date("OTE001", 709, 24);
  Sequence()
  {
   Boundary("T2");
   KDE_Plot("AHM")
   {
    Delta_R("AHM001_R",30,20);
    R_Date("AHM001", 579, 25);
    Delta_R("AHM011_R",30,20);
    R_Date("AHM011", 616, 23);
   };
   Phase()
```

```
{
 Delta_R("MAL003_R",30,20);
 R_Date("MAL003", 445, 24);
 Sequence()
 {
  Boundary("T3");
  Phase()
  {
   Delta_R("STA001_R",30,20);
   R_Date("STA001", 417, 42);
   Sequence()
   {
    Boundary("T4");
    Phase()
    {
     KDE_Plot("CLP")
     {
      Delta_R("CLP015_R",30,20);
      R_Date("CLP015", 505, 22);
      Delta_R("CLP017_R",30,20);
      R_Date("CLP017", 563, 25);
     };
     Sequence()
     {
      Boundary("T5");
      Phase()
      {
       KDE_Plot("MAL")
       {
        Delta_R("MAL004_R",30,20);
        R_Date("MAL004", 363, 25);
        Delta_R("MAL007_R",30,20);
        R_Date("MAL007", 351, 24);
       };
       Delta_R("NMS002_R",30,20);
       R_Date("NMS002", 404, 26);
       Delta_R("G488_R",30,20);
       R_Date("G488", 353, 27);
       Sequence()
       {
        Boundary("T6");
        Phase()
        {
        Phase()
        {
        Delta_R("ELW0098_R",30,20);
        R_Date("ELW098", 350, 22);
        Delta_R("AGU025_R",30,20);
        R_Date("AGU025", 405, 18);
        };
        Sequence()
        {
        KDE_Plot("LBG")
        {
        Delta_R("LBG005_R",30,20);
        R_Date("LBG005", 367, 22);
        Delta_R("LBG007_R",30,20);
        R_Date("LBG007", 361, 22);
```

```
Delta_R("LBG002_R",30,20);
R_Date("LBG002", 358, 22);
};
Boundary("T7");
Phase()
{
KDE_Plot("EMS")
{
Delta_R("EMS010_R",30,20);
R_Date("EMS010", 426, 22);
Delta_R("EMS001_R",30,20);
R_Date("EMS001", 411, 28);
Delta_R("EMS_17_R",30,20);
R_Date("EMS_17", 361, 22);
Delta_R("EMS_99A_R",30,20);
R_Date("EMS_99A", 360, 22);
};
KDE_Plot("STN")
{
Delta_R("STN001_R",30,20);
R_Date("STN001", 349, 30);
Delta_R("STN002_R",30,20);
R_Date("STN002", 327, 30);
Delta_R("STN_005_R",30,20);
R_Date("STN005", 313, 30);
Delta_R("STN006_R",30,20);
R_Date("STN006", 351, 29);
Delta_R("STN011_R",30,20);
R_Date("STN011", 326, 33);
Delta_R("STN012_R",30,20);
R_Date("STN012", 367, 30);
Delta_R("STN_I_101_R",30,20);
R_Date("STN_I_101", 375, 37);
Delta_R("STN015_R",30,20);
R_Date("STN015", 334, 35);
Delta_R("STN017_R",30,20);
R_Date("STN017", 368, 35);
Delta_R("STN028_R",30,20);
R_Date("STN028", 326, 30);
Delta_R("STN029_R",30,20);
R_Date("STN029", 341, 32);
Delta_R("STN030_R",30,20);
R_Date("STN030", 321, 18);
Delta_R("STN_II_107_R",30,20);
R_Date("STN_II_107", 369, 31);
Delta_R("STN031_R",30,20);
R_Date("STN031", 313, 18);
Delta_R("STN032_R",30,20);
R_Date("STN032", 343, 18);
};
};
};
Phase()
{
Sequence()
{
KDE_Plot("1A1_basal")
{
```

```
          Delta_R("Gdansk8_R",30,20);
          R_Date("Gdansk8", 443, 20);
          Delta_R("AGU010_R",30,20);
          R_Date("AGU010", 426, 19);
          Delta_R("RUS003_R",30,20);
          R_Date("RUS003", 381, 28);
          Delta_R("RUS004_R",30,20);
          R_Date("RUS004", 377, 27);
          Delta_R("AGU007_R",30,20);
          R_Date("AGU007", 353, 18);
          };
          Boundary("T8");
          KDE_Plot("BED")
          {
          Delta_R("BED_8147_R",30,20);
          R_Date("BED_8147", 442, 43);
          Delta_R("BED_8198_R",30,20);
          R_Date("BED_8198", 424, 39);
          Delta_R("BED_8216_R",30,20);
          R_Date("BED_8216", 370, 33);
          Delta_R("BED_8219_R",30,20);
          R_Date("BED_8219", 355, 31);
          };
          C_Date("GNW",AD(1710),1);
          };
          };
          };
         };
        };
       };
      };
     };
    };
   };
  };
 };
 };
 Boundary("E");
};
```

**Code S4: Oxcal sequence model using KDE_Plots and ΔR U(0, 50)**

```
 Sequence()
 {
  C_Date("BD",AD(1350),2);
  Boundary("T1");
  Phase()
  {
   Delta_R("MAN008_R",U(0,50));
   R_Date("MAN008", 653, 22);
   Delta_R("OTE001_R",U(0,50));
   R_Date("OTE001", 709, 24);
   Sequence()
   {
    Boundary("T2");
    KDE_Plot("AHM")
    {
     Delta_R("AHM001_R",U(0,50));
```

```
 R_Date("AHM001", 579, 25);
 Delta_R("AHM011_R",U(0,50));
 R_Date("AHM011", 616, 23);
};
Phase()
{
 Delta_R("MAL003_R",U(0,50));
 R_Date("MAL003", 445, 24);
 Sequence()
 {
  Boundary("T3");
  Phase()
  {
   Delta_R("STA001_R",U(0,50));
   R_Date("STA001", 417, 42);
   Sequence()
   {
    Boundary("T4");
    Phase()
    {
     KDE_Plot("CLP")
     {
      Delta_R("CLP015_R",U(0,50));
      R_Date("CLP015", 505, 22);
      Delta_R("CLP017_R",U(0,50));
      R_Date("CLP017", 563, 25);
     };
     Sequence()
     {
      Boundary("T5");
      Phase()
      {
       KDE_Plot("MAL")
       {
        Delta_R("MAL004_R",U(0,50));
        R_Date("MAL004", 363, 25);
        Delta_R("MAL007_R",U(0,50));
        R_Date("MAL007", 351, 24);
       };
       Delta_R("NMS002_R",U(0,50));
       R_Date("NMS002", 404, 26);
       Delta_R("G488_R",U(0,50));
       R_Date("G488", 353, 27);
       Sequence()
       {
        Boundary("T6");
        Phase()
        {
        Phase()
        {
        Delta_R("ELW0098_R",U(0,50));
        R_Date("ELW098", 350, 22);
        Delta_R("AGU025_R",U(0,50));
        R_Date("AGU025", 405, 18);
        };
        Sequence()
        {
        KDE_Plot("LBG")
```

```
{
Delta_R("LBG005_R",U(0,50));
R_Date("LBG005", 367, 22);
Delta_R("LBG007_R",U(0,50));
R_Date("LBG007", 361, 22);
Delta_R("LBG002_R",U(0,50));
R_Date("LBG002", 358, 22);
};
Boundary("T7");
Phase()
{
KDE_Plot("EMS")
{
Delta_R("EMS010_R",U(0,50));
R_Date("EMS010", 426, 22);
Delta_R("EMS001_R",U(0,50));
R_Date("EMS001", 411, 28);
Delta_R("EMS_17_R",U(0,50));
R_Date("EMS_17", 361, 22);
Delta_R("EMS_99A_R",U(0,50));
R_Date("EMS_99A", 360, 22);
};
KDE_Plot("STN")
{
Delta_R("STN001_R",U(0,50));
R_Date("STN001", 349, 30);
Delta_R("STN002_R",U(0,50));
R_Date("STN002", 327, 30);
Delta_R("STN_005_R",U(0,50));
R_Date("STN005", 313, 30);
Delta_R("STN006_R",U(0,50));
R_Date("STN006", 351, 29);
Delta_R("STN011_R",U(0,50));
R_Date("STN011", 326, 33);
Delta_R("STN012_R",U(0,50));
R_Date("STN012", 367, 30);
Delta_R("STN_I_101_R",U(0,50));
R_Date("STN_I_101", 375, 37);
Delta_R("STN015_R",U(0,50));
R_Date("STN015", 334, 35);
Delta_R("STN017_R",U(0,50));
R_Date("STN017", 368, 35);
Delta_R("STN028_R",U(0,50));
R_Date("STN028", 326, 30);
Delta_R("STN029_R",U(0,50));
R_Date("STN029", 341, 32);
Delta_R("STN030_R",U(0,50));
R_Date("STN030", 321, 18);
Delta_R("STN_II_107_R",U(0,50));
R_Date("STN_II_107", 369, 31);
Delta_R("STN031_R",U(0,50));
R_Date("STN031", 313, 18);
Delta_R("STN032_R",U(0,50));
R_Date("STN032", 343, 18);
};
};
};
Phase()
```

```
          {
          Sequence()
          {
          KDE_Plot("1A1_basal")
          {
          Delta_R("Gdansk8_R",U(0,50));
          R_Date("Gdansk8", 443, 20);
          Delta_R("AGU010_R",U(0,50));
          R_Date("AGU010", 426, 19);
          Delta_R("RUS003_R",U(0,50));
          R_Date("RUS003", 381, 28);
          Delta_R("RUS004_R",U(0,50));
          R_Date("RUS004", 377, 27);
          Delta_R("AGU007_R",U(0,50));
          R_Date("AGU007", 353, 18);
          };
          Boundary("T8");
          KDE_Plot("BED")
          {
          Delta_R("BED_8147_R",U(0,50));
          R_Date("BED_8147", 442, 43);
          Delta_R("BED_8198_R",U(0,50));
          R_Date("BED_8198", 424, 39);
          Delta_R("BED_8216_R",U(0,50));
          R_Date("BED_8216", 370, 33);
          Delta_R("BED_8219_R",U(0,50));
          R_Date("BED_8219", 355, 31);
          };
          C_Date("GNW",AD(1710),1);
          };
          };
          };
         };
        };
       };
      };
     };
    };
   };
  };
 };
 };
 Boundary("E");
};
```

**Code S5: Oxcal sequence model using KDE_Plots without ΔR**

```
Sequence()
{
 C_Date("BD",AD(1350),2);
 Boundary("T1");
 Phase()
 {
  R_Date("MAN008", 653, 22);
  R_Date("OTE001", 709, 24);
  Sequence()
  {
   Boundary("T2");
```

```
KDE_Plot("AHM")
{
 R_Date("AHM001", 579, 25);
 R_Date("AHM011", 616, 23);
};
Phase()
{
 R_Date("MAL003", 445, 24);
 Sequence()
 {
  Boundary("T3");
  Phase()
  {
   R_Date("STA001", 417, 42);
   Sequence()
   {
    Boundary("T4");
    Phase()
    {
     KDE_Plot("CLP")
     {
      R_Date("CLP015", 505, 22);
      R_Date("CLP017", 563, 25);
     };
     Sequence()
     {
      Boundary("T5");
      Phase()
      {
       KDE_Plot("MAL")
       {
        R_Date("MAL004", 363, 25);
        R_Date("MAL007", 351, 24);
       };
       R_Date("NMS002", 404, 26);
       R_Date("G488", 353, 27);
       Sequence()
       {
        Boundary("T6");
        Phase()
        {
        Phase()
        {
        R_Date("ELW098", 350, 22);
        R_Date("AGU025", 405, 18);
        };
        Sequence()
        {
        KDE_Plot("LBG")
        {
        R_Date("LBG005", 367, 22);
        R_Date("LBG007", 361, 22);
        R_Date("LBG002", 358, 22);
        };
        Boundary("T7");
        Phase()
        {
        KDE_Plot("EMS")
```

```
{
R_Date("EMS010", 426, 22);
R_Date("EMS001", 411, 28);
R_Date("EMS_17", 361, 22);
R_Date("EMS_99A", 360, 22);
};
KDE_Plot("STN")
{
R_Date("STN001", 349, 30);
R_Date("STN002", 327, 30);
R_Date("STN005", 313, 30);
R_Date("STN006", 351, 29);
R_Date("STN011", 326, 33);
R_Date("STN012", 367, 30);
R_Date("STN_I_101", 375, 37);
R_Date("STN015", 334, 35);
R_Date("STN017", 368, 35);
R_Date("STN028", 326, 30);
R_Date("STN029", 341, 32);
R_Date("STN030", 321, 18);
R_Date("STN_II_107", 369, 31);
R_Date("STN031", 313, 18);
R_Date("STN032", 343, 18);
};
};
};
Phase()
{
Sequence()
{
KDE_Plot("1A1_basal")
{
R_Date("Gdansk8", 443, 20);
R_Date("AGU010", 426, 19);
R_Date("RUS003", 381, 28);
R_Date("RUS004", 377, 27);
R_Date("AGU007", 353, 18);
};
Boundary("T8");
KDE_Plot("BED")
{
R_Date("BED_8147", 442, 43);
R_Date("BED_8198", 424, 39);
R_Date("BED_8216", 370, 33);
R_Date("BED_8219", 355, 31);
};
C_Date("GNW",AD(1710),1);
};
};
};
 };
 };
 };
 };
 };
 };
 };
};
```

```
  };
 };
 Boundary("E");
};
```

**Code S6: Oxcal code for KDE_Plots using ΔR (30, 20)**

```
KDE_Plot("AHM")
{
 Delta_R("AHM001_R",30,20);
 R_Date("AHM001", 579, 25);
 Delta_R("AHM011_R",30,20);
 R_Date("AHM011", 616, 23);
};
KDE_Plot("CLP")
{
 Delta_R("CLP015_R",30,20);
 R_Date("CLP015", 505, 22);
 Delta_R("CLP017_R",30,20);
 R_Date("CLP017", 563, 25);
};
KDE_Plot("MAL")
{
 Delta_R("MAL004_R",30,20);
 R_Date("MAL004", 363, 25);
 Delta_R("MAL007_R",30,20);
 R_Date("MAL007", 351, 24);
};
KDE_Plot("LBG")
{
 Delta_R("LBG005_R",30,20);
 R_Date("LBG005", 367, 22);
 Delta_R("LBG007_R",30,20);
 R_Date("LBG007", 361, 22);
 Delta_R("LBG002_R",30,20);
 R_Date("LBG002", 358, 22);
};
KDE_Plot("EMS")
{
 Delta_R("EMS010_R",30,20);
 R_Date("EMS010", 426, 22);
 Delta_R("EMS001_R",30,20);
 R_Date("EMS001", 411, 28);
 Delta_R("EMS_17_R",30,20);
 R_Date("EMS_17", 361, 22);
 Delta_R("EMS_99A_R",30,20);
 R_Date("EMS_99A", 360, 22);
};
KDE_Plot("STN")
{
 Delta_R("STN001_R",30,20);
 R_Date("STN001", 349, 30);
 Delta_R("STN002_R",30,20);
 R_Date("STN002", 327, 30);
 Delta_R("STN_005_R",30,20);
 R_Date("STN005", 313, 30);
 Delta_R("STN006_R",30,20);
 R_Date("STN006", 351, 29);
```

```
 Delta_R("STN011_R",30,20);
 R_Date("STN011", 326, 33);
 Delta_R("STN012_R",30,20);
 R_Date("STN012", 367, 30);
 Delta_R("STN_I_101_R",30,20);
 R_Date("STN_I_101", 375, 37);
 Delta_R("STN015_R",30,20);
 R_Date("STN015", 334, 35);
 Delta_R("STN017_R",30,20);
 R_Date("STN017", 368, 35);
 Delta_R("STN028_R",30,20);
 R_Date("STN028", 326, 30);
 Delta_R("STN029_R",30,20);
 R_Date("STN029", 341, 32);
 Delta_R("STN030_R",30,20);
 R_Date("STN030", 321, 18);
 Delta_R("STN_II_107_R",30,20);
 R_Date("STN_II_107", 369, 31);
 Delta_R("STN031_R",30,20);
 R_Date("STN031", 313, 18);
 Delta_R("STN032_R",30,20);
 R_Date("STN032", 343, 18);
};
KDE_Plot("1A1_basal")
{
 Delta_R("Gdansk8_R",30,20);
 R_Date("Gdansk8", 443, 20);
 Delta_R("AGU010_R",30,20);
 R_Date("AGU010", 426, 19);
 Delta_R("RUS003_R",30,20);
 R_Date("RUS003", 381, 28);
 Delta_R("RUS004_R",30,20);
 R_Date("RUS004", 377, 27);
 Delta_R("AGU007_R",30,20);
 R_Date("AGU007", 353, 18);
};
KDE_Plot("BED")
{
 Delta_R("BED_8147_R",30,20);
 R_Date("BED_8147", 442, 43);
 Delta_R("BED_8198_R",30,20);
 R_Date("BED_8198", 424, 39);
 Delta_R("BED_8216_R",30,20);
 R_Date("BED_8216", 370, 33);
 Delta_R("BED_8219_R",30,20);
 R_Date("BED_8219", 355, 31);
};
```

**Code S7: Oxcal code for KDE_Plots using ΔR U(0, 50)**

```
KDE_Plot("AHM")
 {
  Delta_R("AHM001_R",U(0,50));
  R_Date("AHM001", 579, 25);
  Delta_R("AHM011_R",U(0,50));
  R_Date("AHM011", 616, 23);
 };
KDE_Plot("CLP")
```

```
 {
  Delta_R("CLP015_R",U(0,50));
  R_Date("CLP015", 505, 22);
  Delta_R("CLP017_R",U(0,50));
  R_Date("CLP017", 563, 25);
 };
KDE_Plot("MAL")
 {
  Delta_R("MAL004_R",U(0,50));
  R_Date("MAL004", 363, 25);
  Delta_R("MAL007_R",U(0,50));
  R_Date("MAL007", 351, 24);
 };
KDE_Plot("LBG")
 {
  Delta_R("LBG005_R",U(0,50));
  R_Date("LBG005", 367, 22);
  Delta_R("LBG007_R",U(0,50));
  R_Date("LBG007", 361, 22);
  Delta_R("LBG002_R",U(0,50));
  R_Date("LBG002", 358, 22);
 };
KDE_Plot("EMS")
 {
  Delta_R("EMS010_R",U(0,50));
  R_Date("EMS010", 426, 22);
  Delta_R("EMS001_R",U(0,50));
  R_Date("EMS001", 411, 28);
  Delta_R("EMS_17_R",U(0,50));
  R_Date("EMS_17", 361, 22);
  Delta_R("EMS_99A_R",U(0,50));
  R_Date("EMS_99A", 360, 22);
 };
KDE_Plot("STN")
 {
  Delta_R("STN001_R",U(0,50));
  R_Date("STN001", 349, 30);
  Delta_R("STN002_R",U(0,50));
  R_Date("STN002", 327, 30);
  Delta_R("STN_005_R",U(0,50));
  R_Date("STN005", 313, 30);
  Delta_R("STN006_R",U(0,50));
  R_Date("STN006", 351, 29);
  Delta_R("STN011_R",U(0,50));
  R_Date("STN011", 326, 33);
  Delta_R("STN012_R",U(0,50));
  R_Date("STN012", 367, 30);
  Delta_R("STN_I_101_R",U(0,50));
  R_Date("STN_I_101", 375, 37);
  Delta_R("STN015_R",U(0,50));
  R_Date("STN015", 334, 35);
  Delta_R("STN017_R",U(0,50));
  R_Date("STN017", 368, 35);
  Delta_R("STN028_R",U(0,50));
  R_Date("STN028", 326, 30);
  Delta_R("STN029_R",U(0,50));
  R_Date("STN029", 341, 32);
  Delta_R("STN030_R",U(0,50));
```

```
 R_Date("STN030", 321, 18);
 Delta_R("STN_II_107_R",U(0,50));
 R_Date("STN_II_107", 369, 31);
 Delta_R("STN031_R",U(0,50));
 R_Date("STN031", 313, 18);
 Delta_R("STN032_R",U(0,50));
 R_Date("STN032", 343, 18);
};
KDE_Plot("1A1_basal")
{
 Delta_R("Gdansk8_R",U(0,50));
 R_Date("Gdansk8", 443, 20);
 Delta_R("AGU010_R",U(0,50));
 R_Date("AGU010", 426, 19);
 Delta_R("RUS003_R",U(0,50));
 R_Date("RUS003", 381, 28);
 Delta_R("RUS004_R",U(0,50));
 R_Date("RUS004", 377, 27);
 Delta_R("AGU007_R",U(0,50));
 R_Date("AGU007", 353, 18);
};
KDE_Plot("BED")
{
 Delta_R("BED_8147_R",U(0,50));
 R_Date("BED_8147", 442, 43);
 Delta_R("BED_8198_R",U(0,50));
 R_Date("BED_8198", 424, 39);
 Delta_R("BED_8216_R",U(0,50));
 R_Date("BED_8216", 370, 33);
 Delta_R("BED_8219_R",U(0,50));
 R_Date("BED_8219", 355, 31);
};
```

**Code S8: Oxcal code for KDE_Plots without ΔR**

```
KDE_Plot("AHM")
{
 R_Date("AHM001", 579, 25);
 R_Date("AHM011", 616, 23);
};
KDE_Plot("CLP")
{
 R_Date("CLP015", 505, 22);
 R_Date("CLP017", 563, 25);
};
KDE_Plot("MAL")
{
 R_Date("MAL004", 363, 25);
 R_Date("MAL007", 351, 24);
};
KDE_Plot("LBG")
{
 R_Date("LBG005", 367, 22);
 R_Date("LBG007", 361, 22);
 R_Date("LBG002", 358, 22);
};
KDE_Plot("EMS")
{
```

```
 R_Date("EMS010", 426, 22);
 R_Date("EMS001", 411, 28);
 R_Date("EMS_17", 361, 22);
 R_Date("EMS_99A", 360, 22);
};
KDE_Plot("STN")
{
 R_Date("STN001", 349, 30);
 R_Date("STN002", 327, 30);
 R_Date("STN005", 313, 30);
 R_Date("STN006", 351, 29);
 R_Date("STN011", 326, 33);
 R_Date("STN012", 367, 30);
 R_Date("STN_I_101", 375, 37);
 R_Date("STN015", 334, 35);
 R_Date("STN017", 368, 35);
 R_Date("STN028", 326, 30);
 R_Date("STN029", 341, 32);
 R_Date("STN030", 321, 18);
 R_Date("STN_II_107", 369, 31);
 R_Date("STN031", 313, 18);
 R_Date("STN032", 343, 18);
};
KDE_Plot("1A1_basal")
{
 R_Date("Gdansk8", 443, 20);
 R_Date("AGU010", 426, 19);
 R_Date("RUS003", 381, 28);
 R_Date("RUS004", 377, 27);
 R_Date("AGU007", 353, 18);
};
KDE_Plot("BED")
{
 R_Date("BED_8147", 442, 43);
 R_Date("BED_8198", 424, 39);
 R_Date("BED_8216", 370, 33);
 R_Date("BED_8219", 355, 31);
};
```

**Code S9: Oxcal code for KDE_Models using ΔR (30, 20)**

```
KDE_Model("AHM")
{
 Delta_R("AHM001_R",30,20);
 R_Date("AHM001", 579, 25);
 Delta_R("AHM011_R",30,20);
 R_Date("AHM011", 616, 23);
};
KDE_Model("CLP")
{
 Delta_R("CLP015_R",30,20);
 R_Date("CLP015", 505, 22);
 Delta_R("CLP017_R",30,20);
 R_Date("CLP017", 563, 25);
};
KDE_Model("MAL")
{
 Delta_R("MAL004_R",30,20);
```

```
 R_Date("MAL004", 363, 25);
 Delta_R("MAL007_R",30,20);
 R_Date("MAL007", 351, 24);
};
KDE_Model("LBG")
{
 Delta_R("LBG005_R",30,20);
 R_Date("LBG005", 367, 22);
 Delta_R("LBG007_R",30,20);
 R_Date("LBG007", 361, 22);
 Delta_R("LBG002_R",30,20);
 R_Date("LBG002", 358, 22);
};
KDE_Model("EMS")
{
 Delta_R("EMS010_R",30,20);
 R_Date("EMS010", 426, 22);
 Delta_R("EMS001_R",30,20);
 R_Date("EMS001", 411, 28);
 Delta_R("EMS_17_R",30,20);
 R_Date("EMS_17", 361, 22);
 Delta_R("EMS_99A_R",30,20);
 R_Date("EMS_99A", 360, 22);
};
KDE_Model("STN")
{
 Delta_R("STN001_R",30,20);
 R_Date("STN001", 349, 30);
 Delta_R("STN002_R",30,20);
 R_Date("STN002", 327, 30);
 Delta_R("STN_005_R",30,20);
 R_Date("STN005", 313, 30);
 Delta_R("STN006_R",30,20);
 R_Date("STN006", 351, 29);
 Delta_R("STN011_R",30,20);
 R_Date("STN011", 326, 33);
 Delta_R("STN012_R",30,20);
 R_Date("STN012", 367, 30);
 Delta_R("STN_I_101_R",30,20);
 R_Date("STN_I_101", 375, 37);
 Delta_R("STN015_R",30,20);
 R_Date("STN015", 334, 35);
 Delta_R("STN017_R",30,20);
 R_Date("STN017", 368, 35);
 Delta_R("STN028_R",30,20);
 R_Date("STN028", 326, 30);
 Delta_R("STN029_R",30,20);
 R_Date("STN029", 341, 32);
 Delta_R("STN030_R",30,20);
 R_Date("STN030", 321, 18);
 Delta_R("STN_II_107_R",30,20);
 R_Date("STN_II_107", 369, 31);
 Delta_R("STN031_R",30,20);
 R_Date("STN031", 313, 18);
 Delta_R("STN032_R",30,20);
 R_Date("STN032", 343, 18);
};
KDE_Model("1A1_basal")
```

```
{
 Delta_R("Gdansk8_R",30,20);
 R_Date("Gdansk8", 443, 20);
 Delta_R("AGU010_R",30,20);
 R_Date("AGU010", 426, 19);
 Delta_R("RUS003_R",30,20);
 R_Date("RUS003", 381, 28);
 Delta_R("RUS004_R",30,20);
 R_Date("RUS004", 377, 27);
 Delta_R("AGU007_R",30,20);
 R_Date("AGU007", 353, 18);
};
KDE_Model("BED")
{
 Delta_R("BED_8147_R",30,20);
 R_Date("BED_8147", 442, 43);
 Delta_R("BED_8198_R",30,20);
 R_Date("BED_8198", 424, 39);
 Delta_R("BED_8216_R",30,20);
 R_Date("BED_8216", 370, 33);
 Delta_R("BED_8219_R",30,20);
 R_Date("BED_8219", 355, 31);
};
```

**Code S10: Oxcal code for KDE_Models using ΔR U(0, 50)**

```
KDE_Model("AHM")
{
 Delta_R("AHM001_R",U(0,50));
 R_Date("AHM001", 579, 25);
 Delta_R("AHM011_R",U(0,50));
 R_Date("AHM011", 616, 23);
};
KDE_Model("CLP")
{
 Delta_R("CLP015_R",U(0,50));
 R_Date("CLP015", 505, 22);
 Delta_R("CLP017_R",U(0,50));
 R_Date("CLP017", 563, 25);
};
KDE_Model("MAL")
{
 Delta_R("MAL004_R",U(0,50));
 R_Date("MAL004", 363, 25);
 Delta_R("MAL007_R",U(0,50));
 R_Date("MAL007", 351, 24);
};
KDE_Model("LBG")
{
 Delta_R("LBG005_R",U(0,50));
 R_Date("LBG005", 367, 22);
 Delta_R("LBG007_R",U(0,50));
 R_Date("LBG007", 361, 22);
 Delta_R("LBG002_R",U(0,50));
 R_Date("LBG002", 358, 22);
};
KDE_Model("EMS")
{
```

```
 Delta_R("EMS010_R",U(0,50));
 R_Date("EMS010", 426, 22);
 Delta_R("EMS001_R",U(0,50));
 R_Date("EMS001", 411, 28);
 Delta_R("EMS_17_R",U(0,50));
 R_Date("EMS_17", 361, 22);
 Delta_R("EMS_99A_R",U(0,50));
 R_Date("EMS_99A", 360, 22);
};
KDE_Model("STN")
{
 Delta_R("STN001_R",U(0,50));
 R_Date("STN001", 349, 30);
 Delta_R("STN002_R",U(0,50));
 R_Date("STN002", 327, 30);
 Delta_R("STN_005_R",U(0,50));
 R_Date("STN005", 313, 30);
 Delta_R("STN006_R",U(0,50));
 R_Date("STN006", 351, 29);
 Delta_R("STN011_R",U(0,50));
 R_Date("STN011", 326, 33);
 Delta_R("STN012_R",U(0,50));
 R_Date("STN012", 367, 30);
 Delta_R("STN_I_101_R",U(0,50));
 R_Date("STN_I_101", 375, 37);
 Delta_R("STN015_R",U(0,50));
 R_Date("STN015", 334, 35);
 Delta_R("STN017_R",U(0,50));
 R_Date("STN017", 368, 35);
 Delta_R("STN028_R",U(0,50));
 R_Date("STN028", 326, 30);
 Delta_R("STN029_R",U(0,50));
 R_Date("STN029", 341, 32);
 Delta_R("STN030_R",U(0,50));
 R_Date("STN030", 321, 18);
 Delta_R("STN_II_107_R",U(0,50));
 R_Date("STN_II_107", 369, 31);
 Delta_R("STN031_R",U(0,50));
 R_Date("STN031", 313, 18);
 Delta_R("STN032_R",U(0,50));
 R_Date("STN032", 343, 18);
};
KDE_Model("1A1_basal")
{
 Delta_R("Gdansk8_R",U(0,50));
 R_Date("Gdansk8", 443, 20);
 Delta_R("AGU010_R",U(0,50));
 R_Date("AGU010", 426, 19);
 Delta_R("RUS003_R",U(0,50));
 R_Date("RUS003", 381, 28);
 Delta_R("RUS004_R",U(0,50));
 R_Date("RUS004", 377, 27);
 Delta_R("AGU007_R",U(0,50));
 R_Date("AGU007", 353, 18);
};
KDE_Model("BED")
{
 Delta_R("BED_8147_R",U(0,50));
```

```
 R_Date("BED_8147", 442, 43);
 Delta_R("BED_8198_R",U(0,50));
 R_Date("BED_8198", 424, 39);
 Delta_R("BED_8216_R",U(0,50));
 R_Date("BED_8216", 370, 33);
 Delta_R("BED_8219_R",U(0,50));
 R_Date("BED_8219", 355, 31);
};
```

**Code S11: Oxcal code for KDE_Models without ΔR**

```
KDE_Model("AHM")
{
 R_Date("AHM001", 579, 25);
 R_Date("AHM011", 616, 23);
};
KDE_Model("CLP")
{
 R_Date("CLP015", 505, 22);
 R_Date("CLP017", 563, 25);
};
KDE_Model("MAL")
{
 R_Date("MAL004", 363, 25);
 R_Date("MAL007", 351, 24);
};
KDE_Model("LBG")
{
 R_Date("LBG005", 367, 22);
 R_Date("LBG007", 361, 22);
 R_Date("LBG002", 358, 22);
};
KDE_Model("EMS")
{
 R_Date("EMS010", 426, 22);
 R_Date("EMS001", 411, 28);
 R_Date("EMS_17", 361, 22);
 R_Date("EMS_99A", 360, 22);
};
KDE_Model("STN")
{
 R_Date("STN001", 349, 30);
 R_Date("STN002", 327, 30);
 R_Date("STN005", 313, 30);
 R_Date("STN006", 351, 29);
 R_Date("STN011", 326, 33);
 R_Date("STN012", 367, 30);
 R_Date("STN_I_101", 375, 37);
 R_Date("STN015", 334, 35);
 R_Date("STN017", 368, 35);
 R_Date("STN028", 326, 30);
 R_Date("STN029", 341, 32);
 R_Date("STN030", 321, 18);
 R_Date("STN_II_107", 369, 31);
 R_Date("STN031", 313, 18);
 R_Date("STN032", 343, 18);
};
KDE_Model("1A1_basal")
```

```
{
 R_Date("Gdansk8", 443, 20);
 R_Date("AGU010", 426, 19);
 R_Date("RUS003", 381, 28);
 R_Date("RUS004", 377, 27);
 R_Date("AGU007", 353, 18);
};
KDE_Model("BED")
{
 R_Date("BED_8147", 442, 43);
 R_Date("BED_8198", 424, 39);
 R_Date("BED_8216", 370, 33);
 R_Date("BED_8219", 355, 31);
};
```

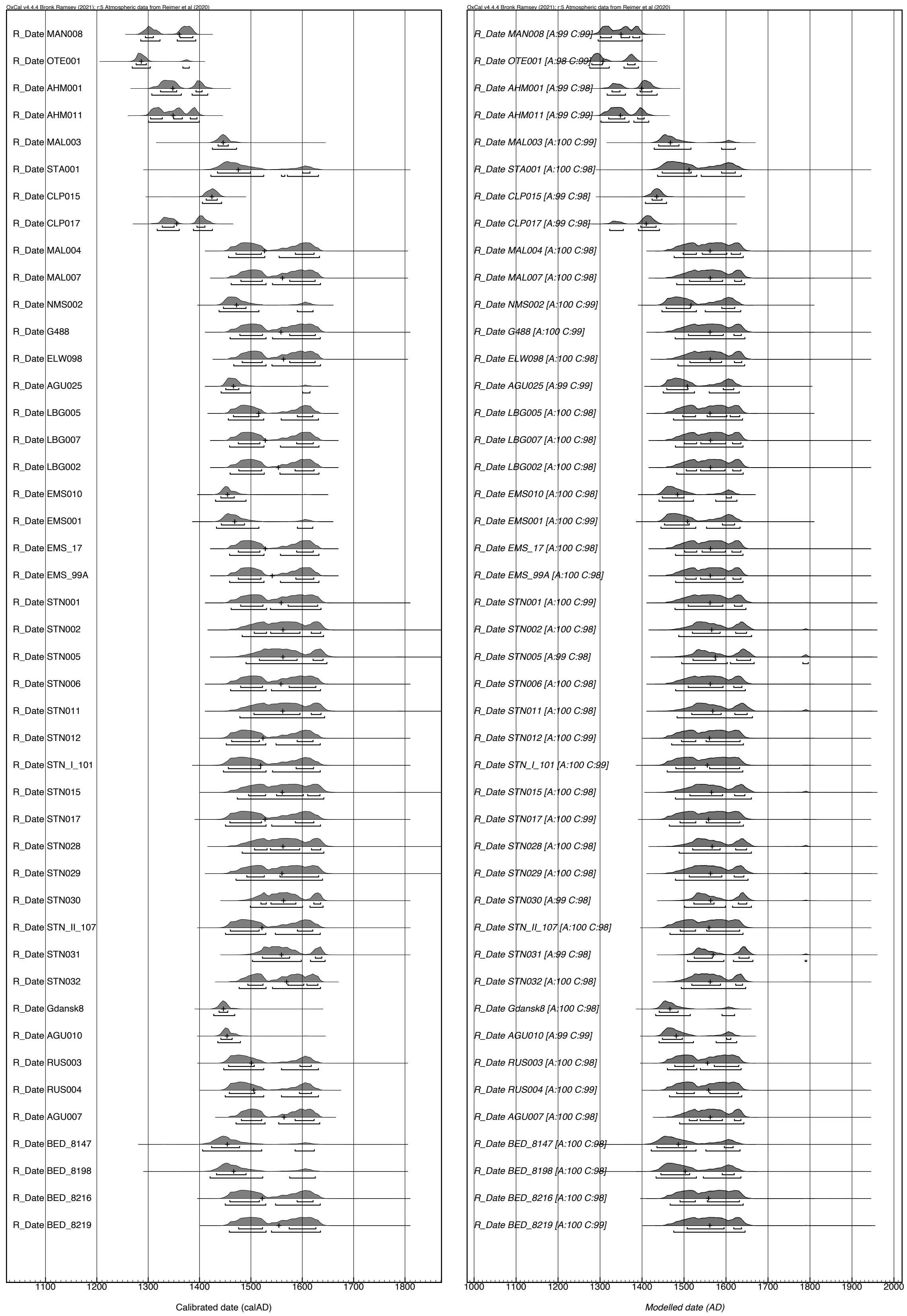

**Fig. S45: Oxcal plot of individually calibrated radiocarbon dates without ΔR [left] and with ΔR (30, 20) [right].**

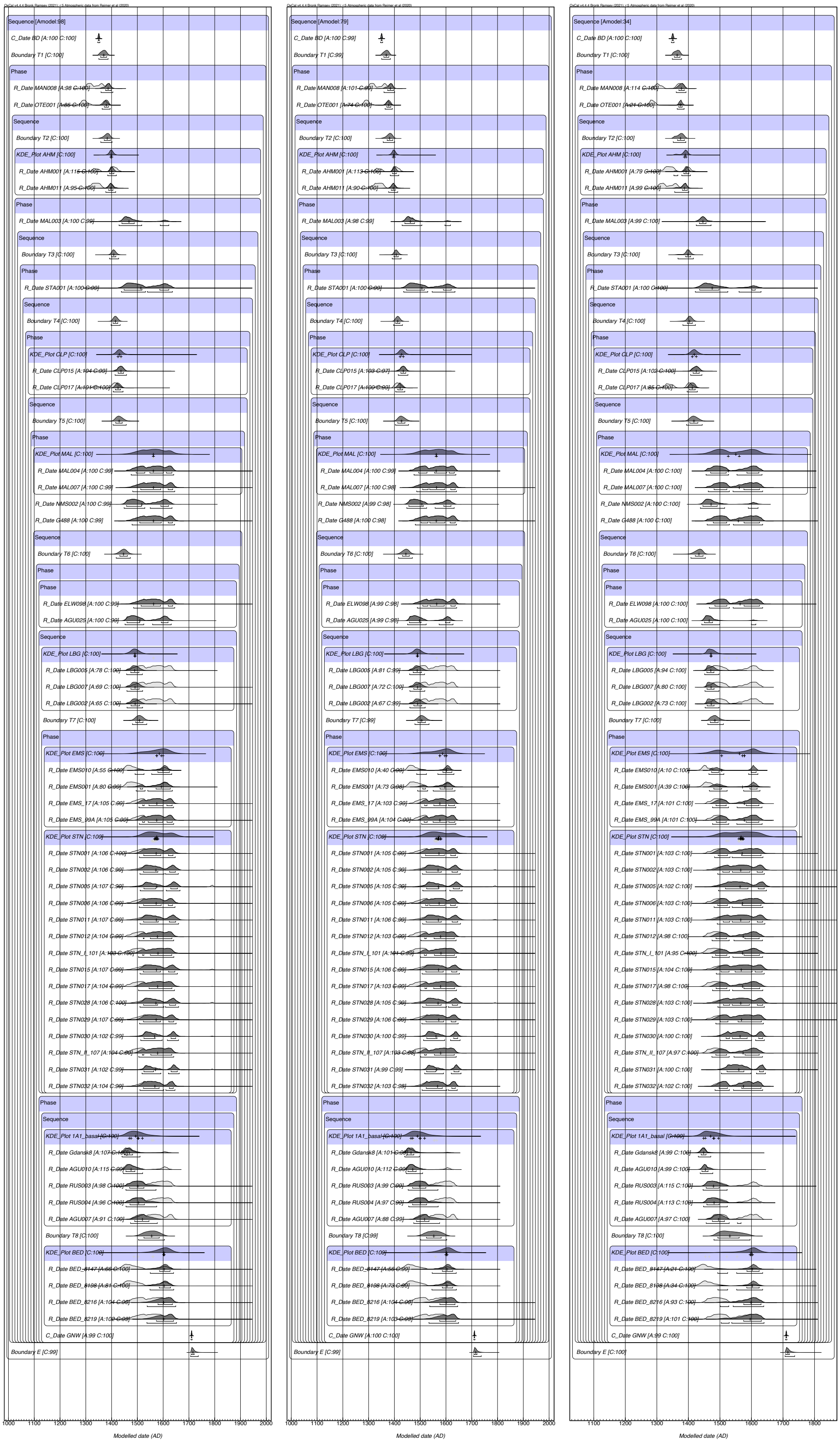

**Fig. S46: Oxcal plot of sequence models using KDE_Plots with ΔR (30, 20) [left], ΔR U(0, 250) [middle] and no ΔR [right].**

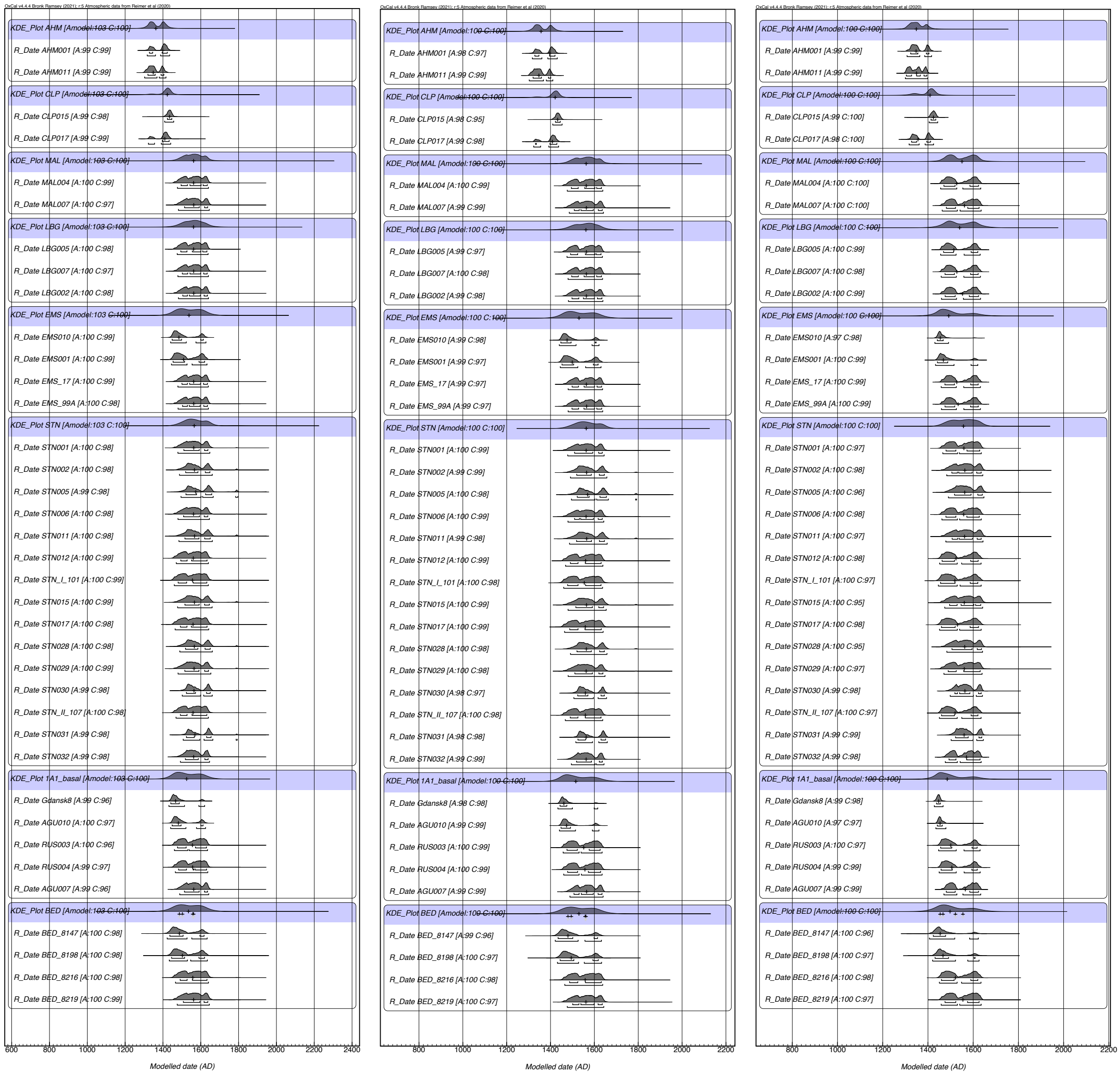

**Fig. S47: Oxcal KDE_Plots with ΔR (30, 20) [left], ΔR U(0, 50) [middle] and no ΔR [right].**

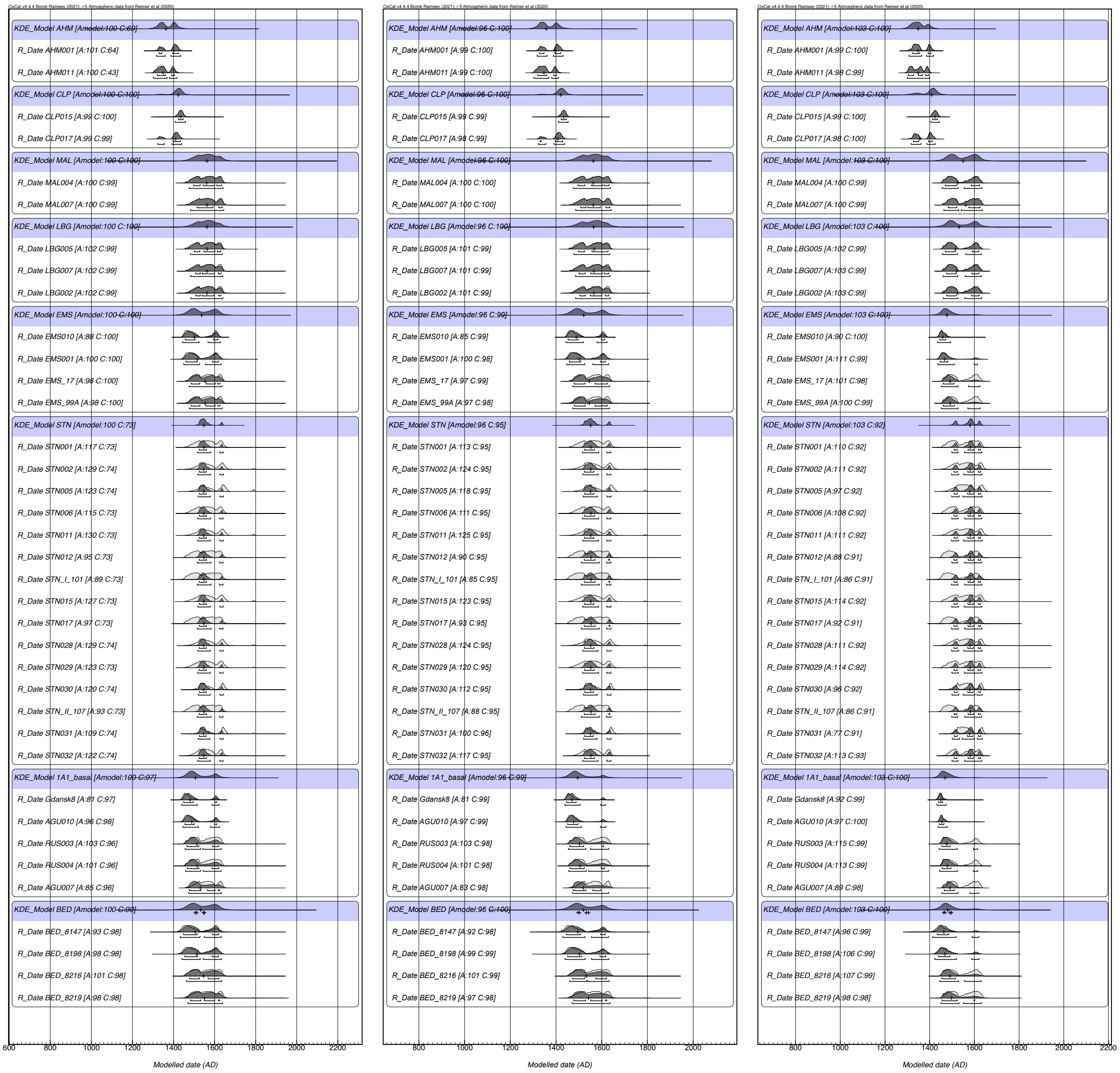

**Fig. S48: Oxcal KDE_Models with ΔR (30, 20) [left], ΔR U(0, 50) [middle] and no ΔR [right].**

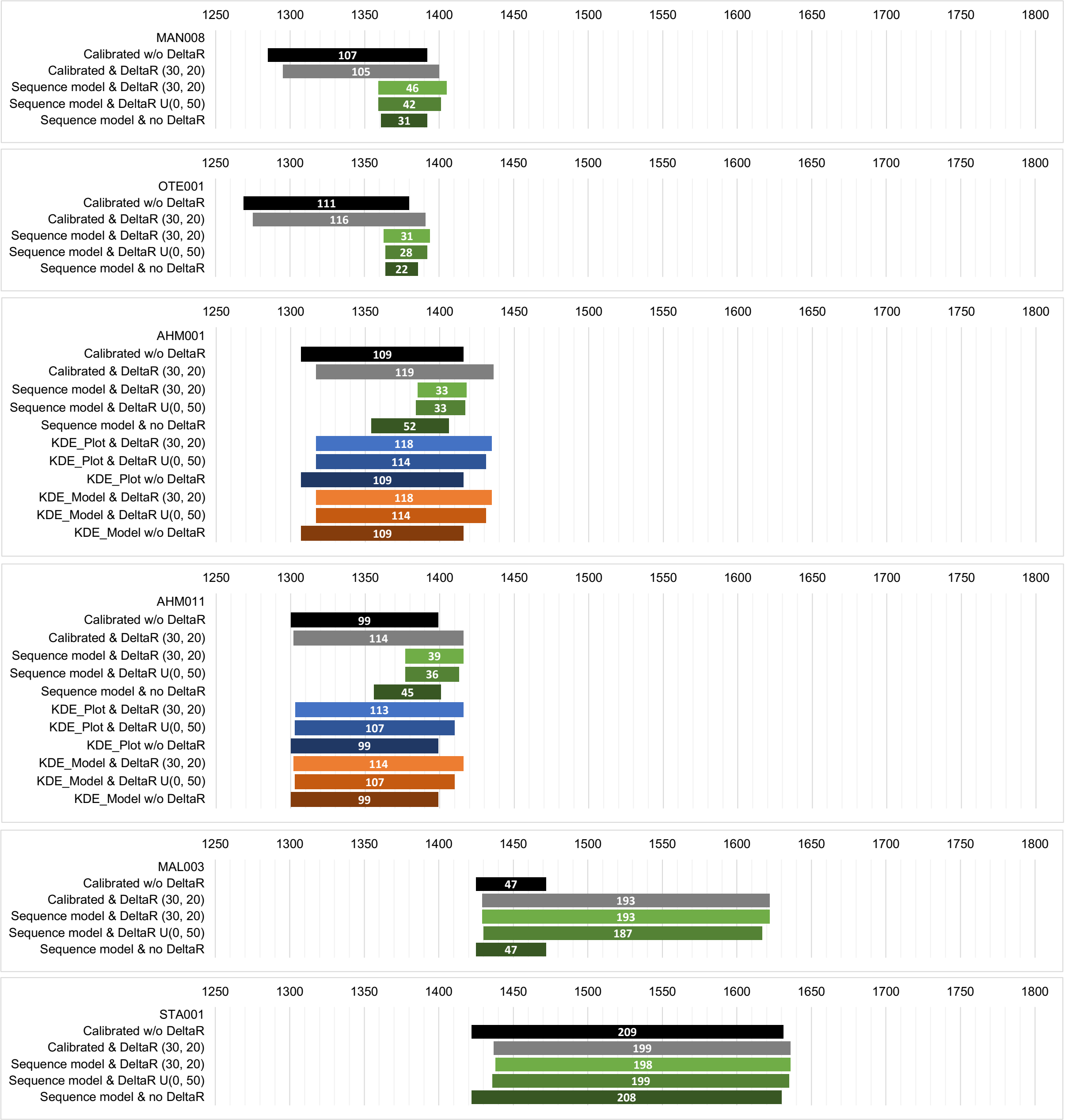

**Fig. S49: Plots showing the 2σ intervals for calibrated radiocarbon dates of the individuals MAN008, OTE001, AHM001, AHM011, MAL003, and STA001 with different modelling approaches.**

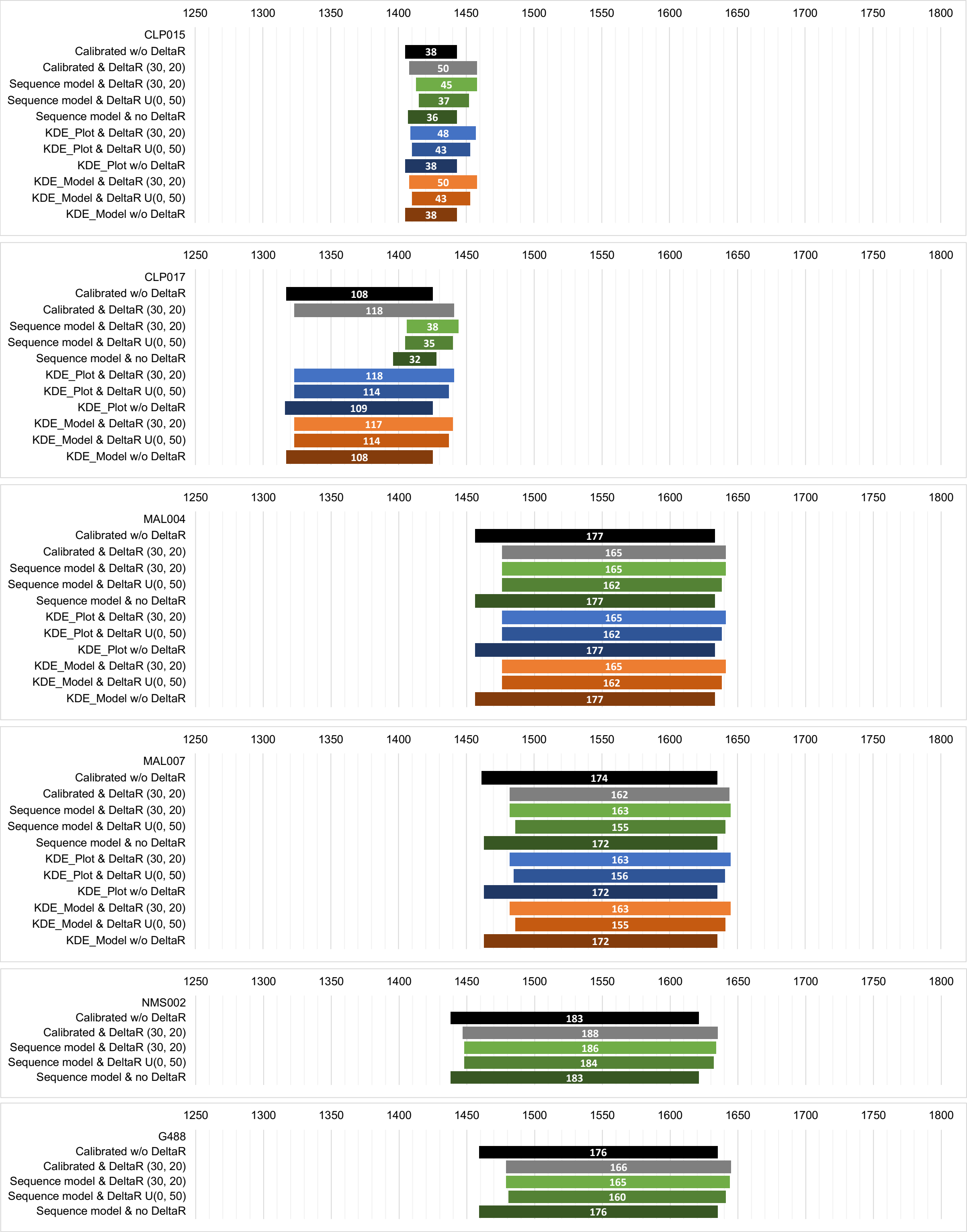

**Fig. S50: Plots showing the 2σ intervals for calibrated radiocarbon dates of the individuals CLP015, CLP017, MAL004, MAL007, NMS002, and G488 with different modelling approaches.**

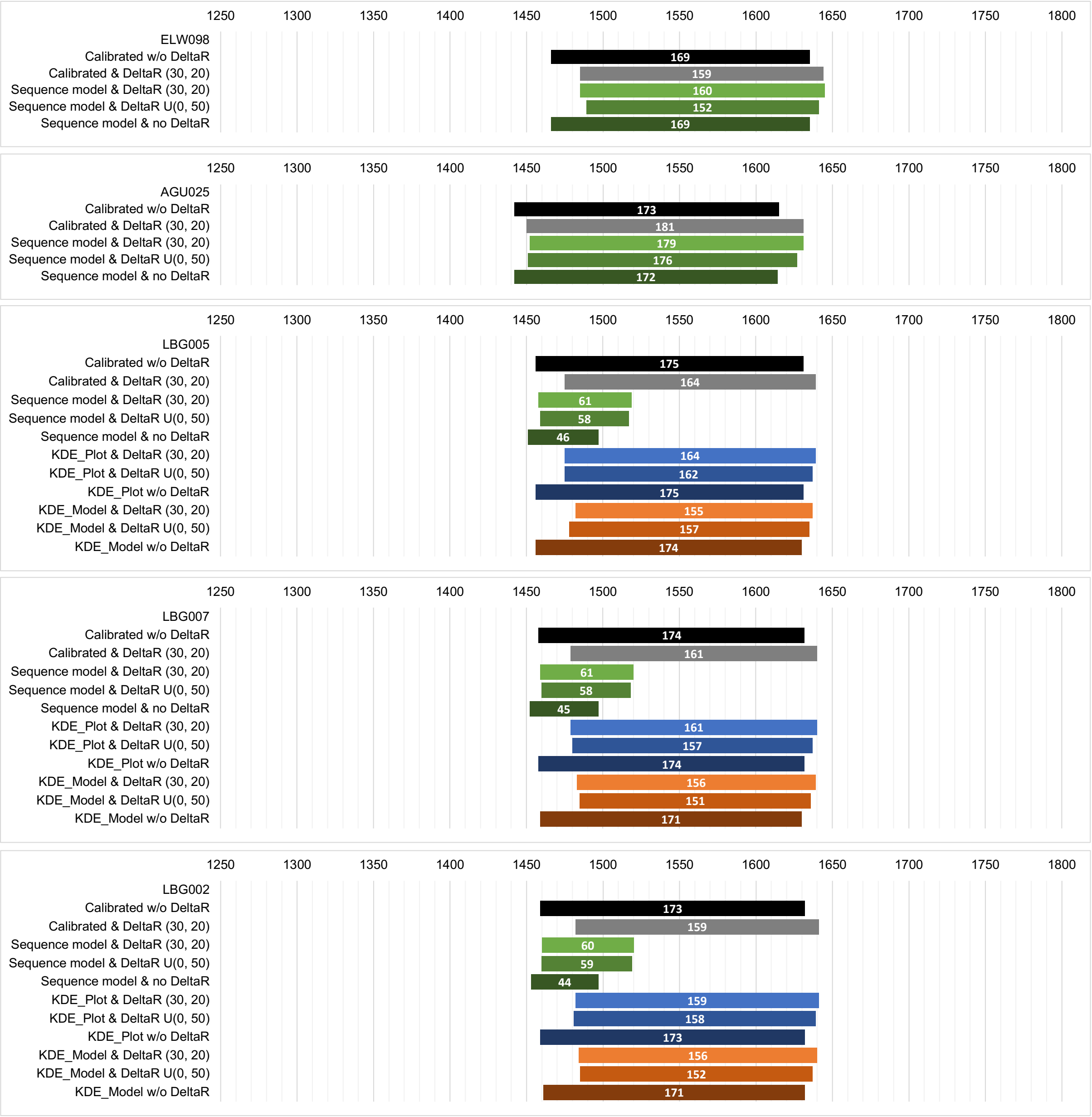

**Fig. S51: Plots showing the 2σ intervals for calibrated radiocarbon dates of the individuals ELW098, AGU025, LBG005, LBG007, and LBG002 with different modelling approaches.**

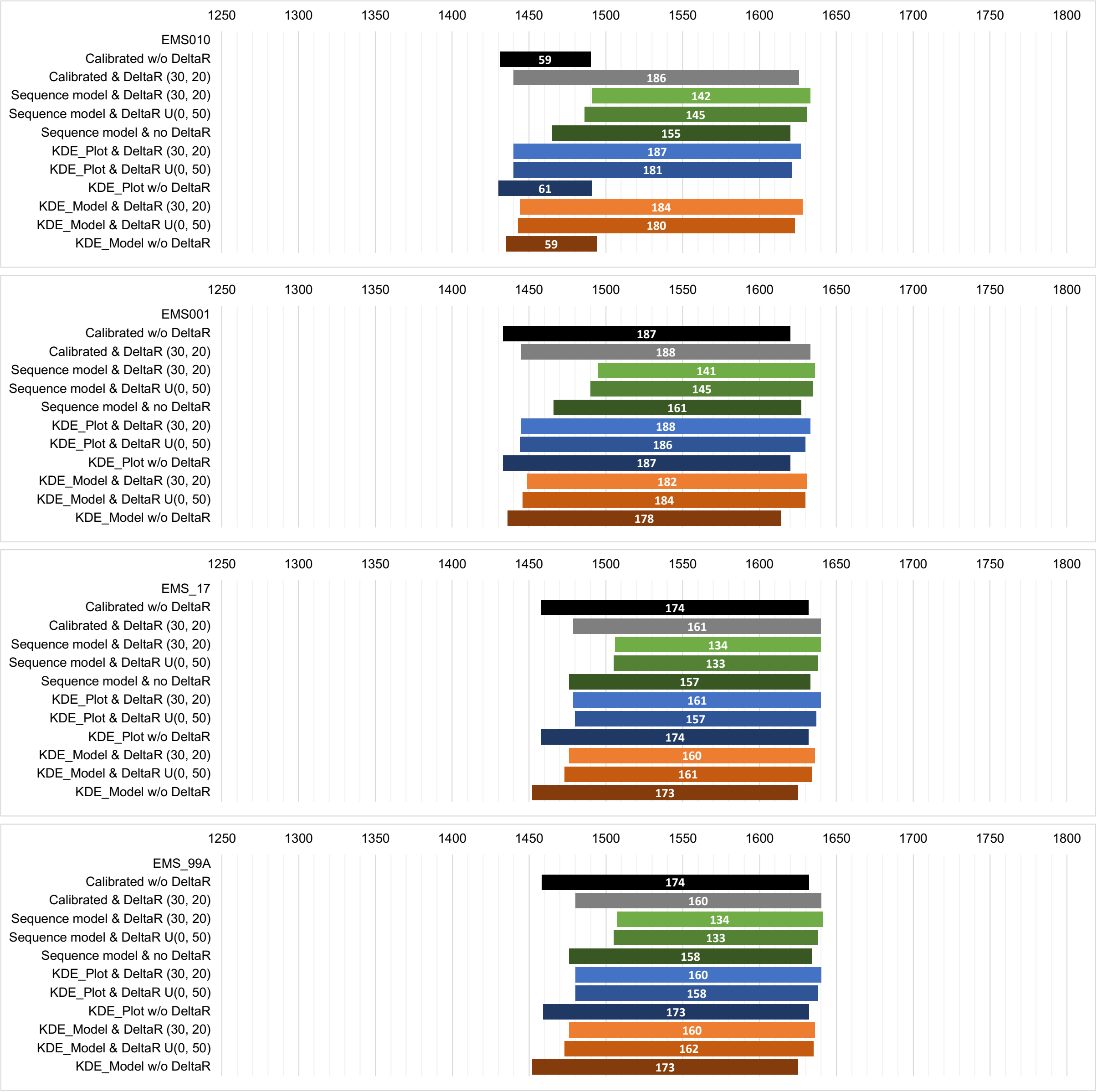

**Fig. S52: Plots showing the 2σ intervals for calibrated radiocarbon dates of the individuals EMS010, EMS001, EMS_17 and EMS_99A with different modelling approaches.**

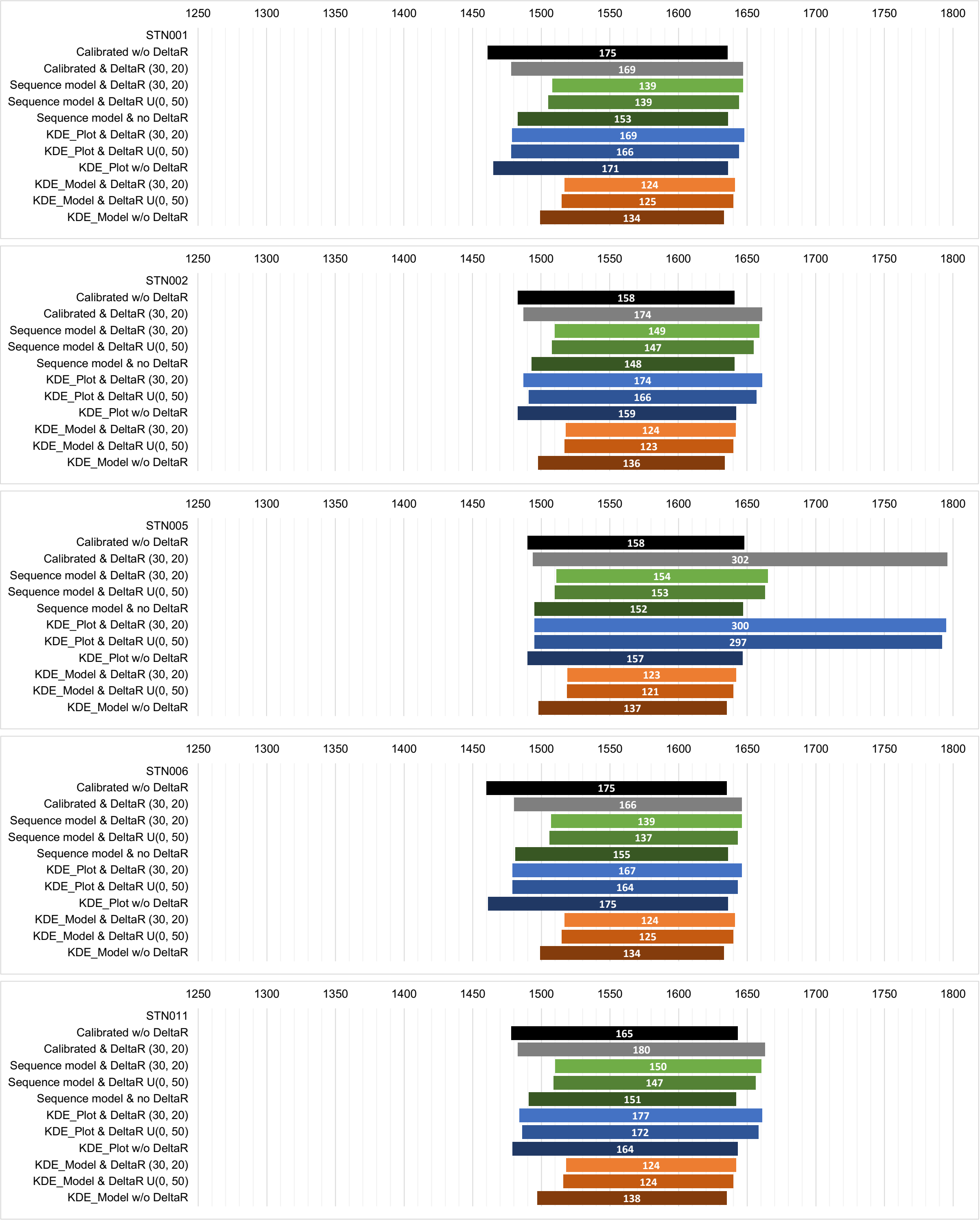

**Fig. S53: Plots showing the 2σ intervals for calibrated radiocarbon dates of the individuals STN001, STN002, STN005, STN006 and STN011 with different modelling approaches.**

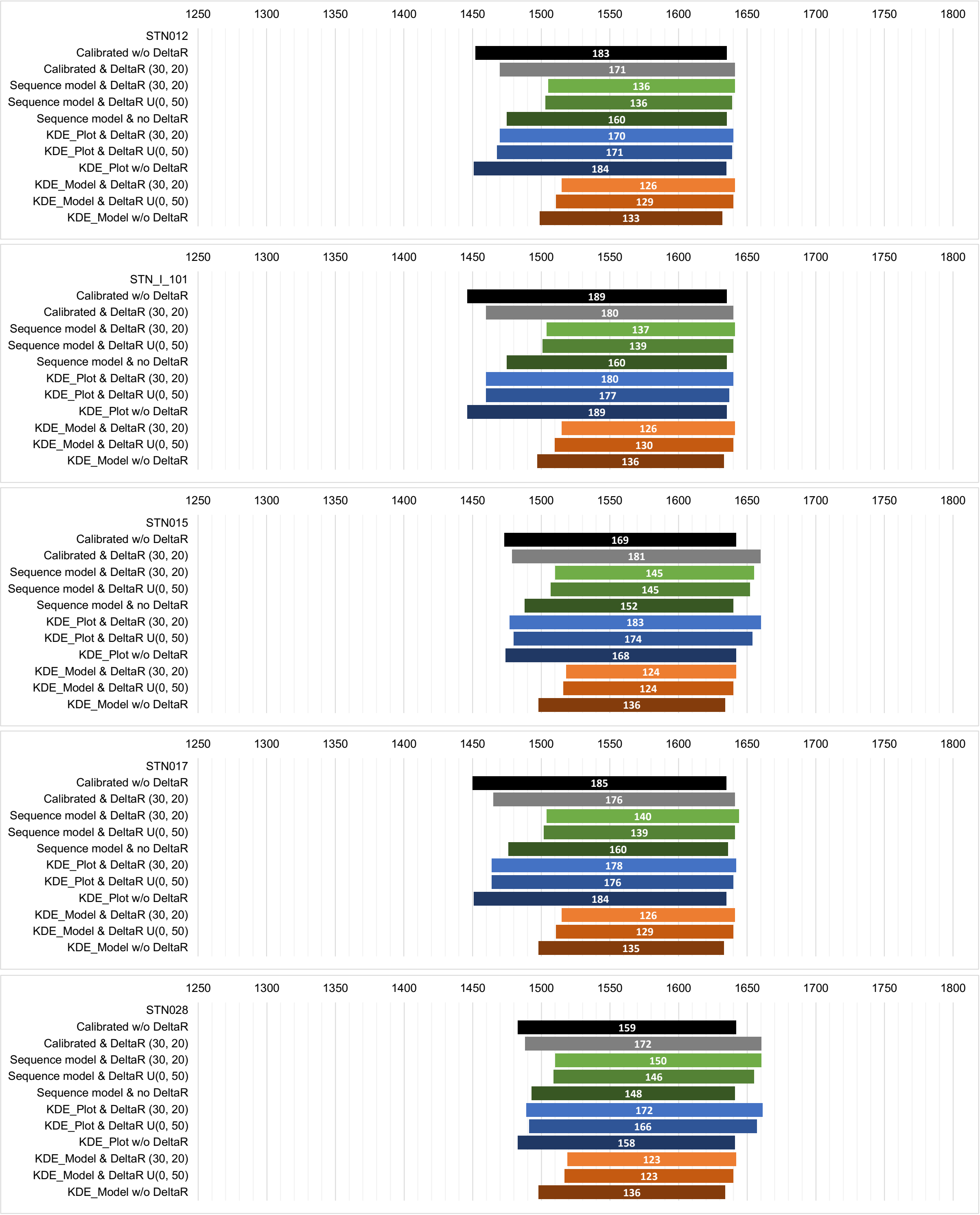

**Fig. S54: Plots showing the 2σ intervals for calibrated radiocarbon dates of the individuals STN012, STN_I_101, STN015, STN017, and STN028 with different modelling approaches.**

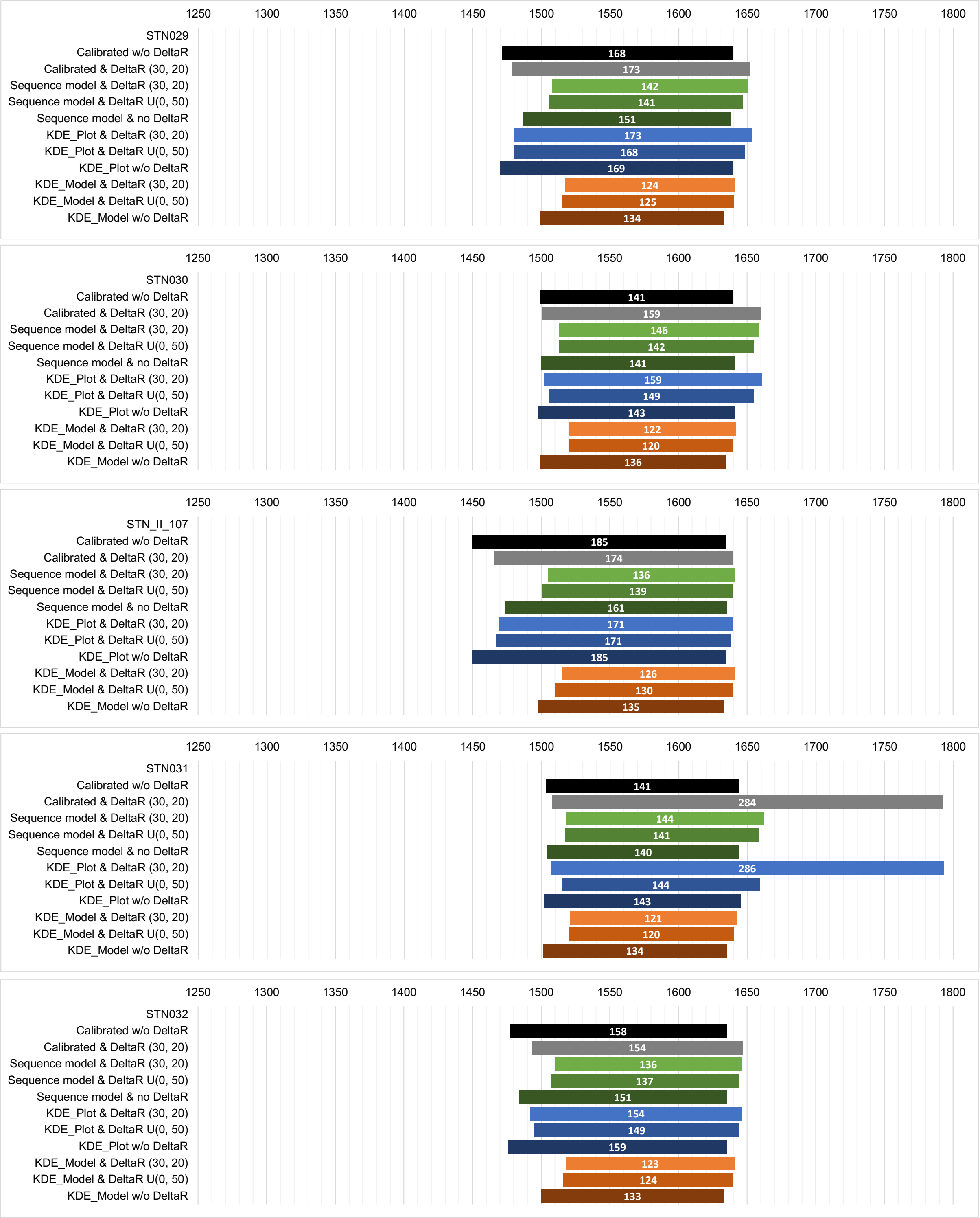

**Fig. S55: Plots showing the 2σ intervals for calibrated radiocarbon dates of the individuals STN030, STN_I_107, STN031 and STN031 with different modelling approaches.**

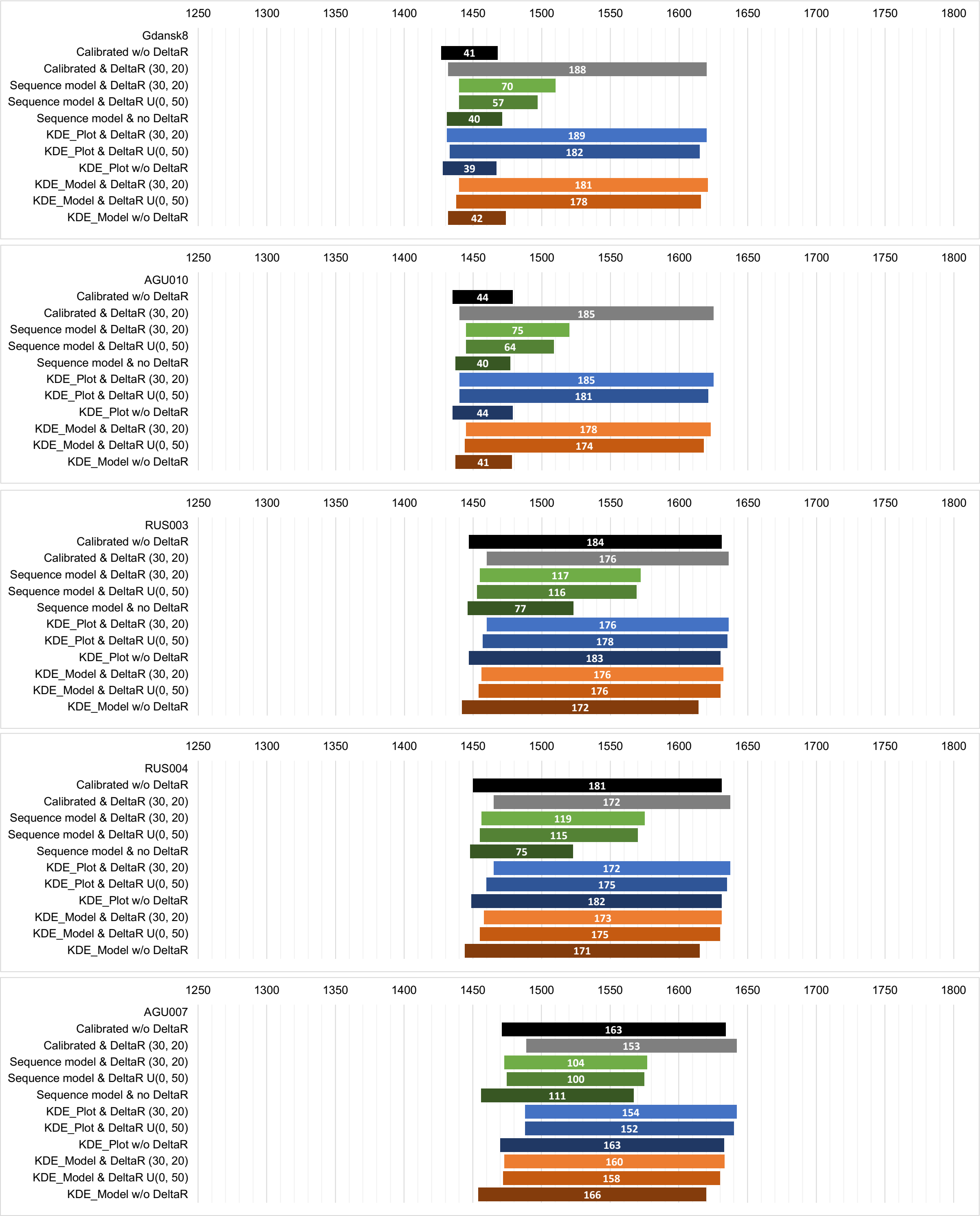

**Fig. S56: Plots showing the 2σ intervals for calibrated radiocarbon dates of the individuals Gdansk8, AGU010, RUS003, RUS004, and AGU007 with different modelling approaches.**

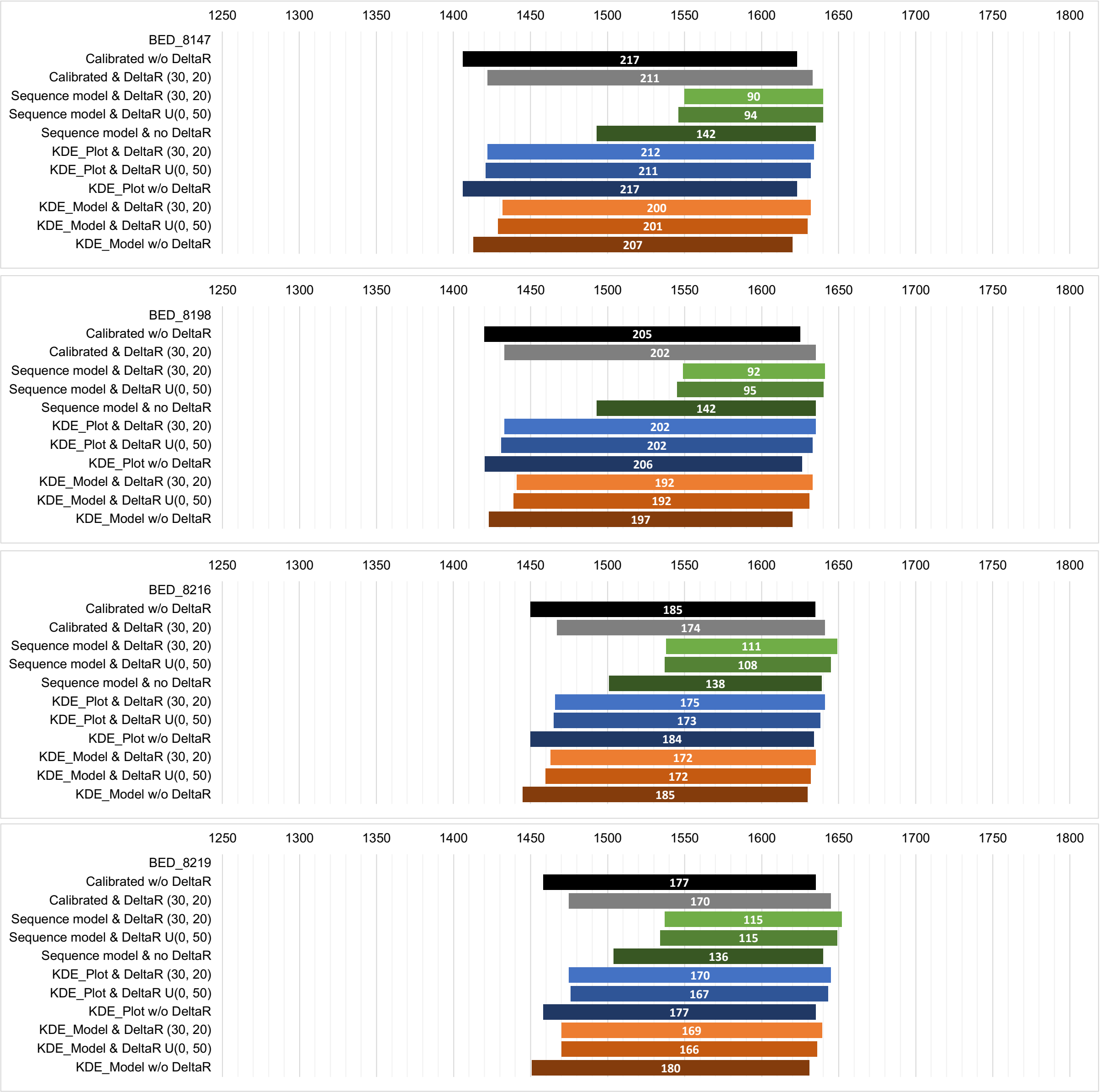

**Fig. S57: Plots showing the 2σ intervals for calibrated radiocarbon dates of the individuals BED_8147, BED_8198, BED_8216, and BED_8219 with different modelling approaches.**

### 11 Historical contextualization of the Second Pandemic plague genomes

*Marcel Keller and Philip Slavin*

#### 11.1 The genomes of the Black Death – BSK001.A, LAI009.A, ESFpool, Barcelona, NAB003.B, the BEN genomes and COM042

The genomic history of the Black Death – i.e. the massive outbreak of plague in the mid-14th century in Europe as the onset of the Second Pandemic – is well-established by now (Bos et al. 2011; Namouchi et al. 2018; Spyrou et al. 2016; Spyrou et al. 2019, 2022). With BSK001.A, a direct progenitor not only of all ancient Second Pandemic genomes and Branch 1, but also Branches 2–4, has been found in Kara-Djigach, modern-day Kyrgyzstan, dating to 1338–1339 (Spyrou et al. 2022), situated on a node giving birth to the "Great Polytomy" - designated as node N01 in this study. This confirms that its positioning on the same node in the original paper by Spyrou et al. 2022 was correct.

Another well-dated genome (pooled from the individuals 8124, 8291, 11972; here called ESFpool) was reconstructed for the site of London East-Smithfield, a cemetery founded in late 1348 or early 1349 and in use during the Black Death outbreak in the city, ending in July 1349 (Sloane 2011; Grainger et al. 2008). It appears to be the most recent common ancestor of all Branch 1 genomes, although Morozova et al. 2020 presented a phylogenetic tree with a short terminal branch for ESFpool and reported the private SNP 2356292 in the SI. It is unclear why this SNP, also identified in our analyses, was not called in other studies, but the SNP evaluation identified it as a false positive SNP due to contamination (Table S5), and therefore it is not shown in the schematic tree (Fig. 2A). Other genomes, not included in this analysis for various reasons (see Table S3), have been shown to be identical with ESFpool in previous studies, such as BSS31 (Abbadia San Salvatore, Siena, Italy; Namouchi et al. 2018), OSL1 (Oslo, Norway; Namouchi et al. 2018), SLC1006 (Saint-Laurent-de-la-Cabrerisse, France; Namouchi et al. 2018), TRP002.A (Toulouse, France; Spyrou et al. 2019). Also, the newly sequenced genomes of Bene't Street, Cambridge, United Kingdom (BEN001 and BEN002) as well as Cambridge All Saints, United Kingdom (COM042) are identical with other European Black Death genomes (see Supplementary Section 7). This is in line with a previously published low-coverage Black Death genome from Cambridge New Museums site/Augustinian Friary NMS003.A (Spyrou et al. 2019), indicating the usage of different burial sites within the city of Cambridge during the local Black Death outbreak in 1349, as discussed also in Cessford et al. 2021. The genome of Laishevo (Tatarstan), LAI009, falls in one SNP derived of the Kara-Djigach genome (BSK001.A) and one SNP ancestral to the Black Death cluster. The original publication offered only a rough dating into the 14th century (1300–1400) based on ceramic and a bronze earring pointing towards the period of the Golden Horde (Spyrou et al. 2019). However, an archaeological report of the Laishevo burials reported these to be Muslim (Rudenko 2013). It was not until the 1320s that Turkic-speaking inhabitants of the Volga-Kama region started converting to Islam, in a piecemeal fashion, following the conversion of their ruler Khan Öz Beg in 1320. Hence, this fact narrows down the dating of LAI009 to

1320–1400. However, with the precise dates of the direct progenitor (BSK001) and descendant (ESFpool), the dating can be narrowed down to around 1338–1349. As all the available textual evidence suggests, plague ravaged the Volga-Kama region in 1346 (see comments to the reviewers published along with Spyrou et al. 2022). If the Laishevo outbreak can be dated to 1346, then it would mean that the one SNPs separating BSK001.A from LAI009 would have been acquired within an interval of around 7–8 years, while the additional SNP defining the ESFpool and other Black Death genomes would have been acquired within the next 2–3 years, between 1346 and 1348–1349 (given that the Black Death came to Saint-Laurent-de-la-Cabrerisse, Toulouse, Sienna and Barcelona, as well as London, in 1348).

#### 11.2 Basal Branch 1B – Ber37, Ber45 and Bolgar

The basal Branch 1B genomes in the phylogeny presented here (Fig. 2) comprise the two genomes from Bergen op Zoom, Netherlands (Ber37 and Ber45; Namouchi et al. 2018) as well as the genome from Bolgar (Tatarstan, Russia; Spyrou et al. 2016). The previously published genome of London St. Mary Graces 6330 (Bos et al. 2011) has been shown to occupy a position on a short terminal branch directly descendant of the Bergen op Zoom genomes (Namouchi et al. 2018) but was excluded in this analysis due to low coverage. The Bergen op Zoom genomes represent the most recent common ancestor (MRCA) of all other Branch 1B genomes, two SNPs derived from the Black Death cluster (Fig. 2A), and have been associated by Namouchi et al. 2018 with the first post-Black Death wave, the *pestis secunda* in Flanders and Antwerp, dated by the authors between 1358 and 1363 (Namouchi et al. 2018). The dating of London St. Mary Graces was corrected to 1350–1400 (Bos et al. 2016) after it was erroneously attributed to the East Smithfield site and the Black Death outbreak of 1348 in the original publication (Bos et al. 2011). The genome of Bolgar was originally dated to 1362–1400 based on numismatic and archaeological evidence (Spyrou et al. 2016). Namouchi et al. 2018 associated this genome with a plague outbreak starting in Nizhnii Novgorod in 1364 and spreading through Russia until 1366. While all of these basal genomes were already associated with the *pestis secunda* dated by the original authors to 1357–1366 (Namouchi et al. 2018), an in-depth analysis of historical data by Slavin was able to establish a geo-chronological framework of the *pestis secunda* outbreaks in Europe, Middle East and North Africa, starting in or around southern Hesse, Germany, in the year 1356 (Slavin 2021). This study also established a chronology for the outbreaks corresponding to ancient *Y. pestis* genomes: Bergen op Zoom in 1359, London St. Mary Graces in 1361, and Bolgar around 1364. This chronology is also consistent with the phylogenetic tree with the ancestral genome – Bergen op Zoom – dating the oldest. As none of the previous publications presented radiocarbon dates for the respective individuals that the plague genomes are derived from, these were not used for PhIRM. However, the contextual data offers a level of precision which cannot be surpassed with radiocarbon dating.

So far, no other ancient genomes have been associated with Branch 1B despite the increasing sampling density of European sites (see Fig. 1). Therefore, it has been proposed that this lineage, emerging in Central Europe and traveling eastwards (Slavin 2021; Spyrou et al. 2016), disappeared from Europe after 1366, but established a reservoir in Asia, where it later gave rise to the Third Pandemic emerging in Yunnan province in 1772 and becoming global in 1894 (Cui et al. 2013; Slavin 2021; Benedict 1996; Xu et al. 2014). However, one sub-branch of Branch 1B, comprising 1.ANT cluster branching off basal to the cluster associated with the global dissemination of the Third Pandemic (1.ORI), has so far been found exclusively in Africa (Uganda, Kenya and Congo; Morelli et al. 2010; Cui et al. 2013; Green 2018) and most likely established a local reservoir prior to the Third Pandemic. So far there is no solid evidence of when this lineage reached Africa; for an in-depth discussion see Green 2018.

#### 11.3 OTE001, COL001 and MAN008.B

Individuals from the site of Manching-Pichl were first identified as plague victims by Garrelt and Wiechmann 2003. In a PCR-based study by Seifert et al. 2016, the causative *Y. pestis* strain was already identified as identical or derived from the Black Death on a separate branch, which could be confirmed by whole-genome reconstruction by (Spyrou et al. 2019). The reconstructed genome (MAN008), assumed to represent the same strain as the other identified plague victims of this site, forms a terminal branch that splits from Branch 1A briefly after the Black Death. Gutsmiedl-Schümann et al. 2018 offered the first radiocarbon dates, (cited prior to publication by Seifert et al. 2016 as “14th century”). However, two out of four radiocarbon date intervals fell completely before the Black Death (2-sigma intervals of 1027–1154 and 985–1026), while the other two covered the Black Death as well as the late 14th century (1287–1391 and 1298–1403), explained here by erroneous dating of skeletal element introduced into the mass grave from older burials at the site (Gutsmiedl-Schümann et al. 2018). To prevent such a mix-up, the same tooth was used in Spyrou et al. 2019 for both the DNA extraction and radiocarbon dating, giving a calibrated 2-sigma range of 1283–1390.

The genome of Colalto Sabino COL001, published by Guellil et al. 2020, forms a terminal branch emerging from the same node (N04, Fig. 2A, 3 SNPs derived from the European Black Death cluster) as MAN008, forming a trifurcation. The authors of the original paper associated COL001 with a local outbreak in 1363 as part of the *pestis secunda* (Guellil et al. 2020), therefore proposing the parallel circulation of two *Y. pestis* lineages during the *pestis secunda* (see Supplementary Section 11.2).

The site of Otepää was first studied for human DNA by Saag et al. 2019. Since none of the graves, mostly containing two to four individuals and laid out in irregular rows, cut into each other, a short usage time and a possible epidemic context of this cemetery were assumed upon excavation. A coin dating to the last third of the 14th century (minted in Tallinn/Reval between 1360 and 1390) gave already a *terminus post quem*, in line with a documented plague outbreak in 1378 in the bishopric of Tartu/Dorpat (Saag et al. 2019). The *Y. pestis* genome reconstructed from the sample OTE001 shares one SNP with COL001 and has an additional private SNP (Fig. 2A). Radiocarbon dating of OTE001 as part of this study yielded a 2-sigma interval of 1269–1380. However, the sequence model without offset (Code S5, Table S11, Fig. 46 right) resulted in a poor model agreement (Amodel: 34.0) and also a low agreement for the OTE001 radiocarbon date (A: 21.4) caused by the year 1350 (±2 years) for the Black Death as prior. The sequence model with an offset (Fig. 3, Code S3, Table S11, Fig. S46 left) performed much better (Amodel: 97.7; OTE001 A: 84.7), and an investigation of the posterior probability of the offset revealed that the median for the offset is at 43 radiocarbon years, shifted by 13 years compared to the prior distribution (30 years), see Fig. 58. This is a strong indicator that this radiocarbon sample has a significant offset, although the cause (HBCO, freshwater or marine reservoir effect) remains unclear.

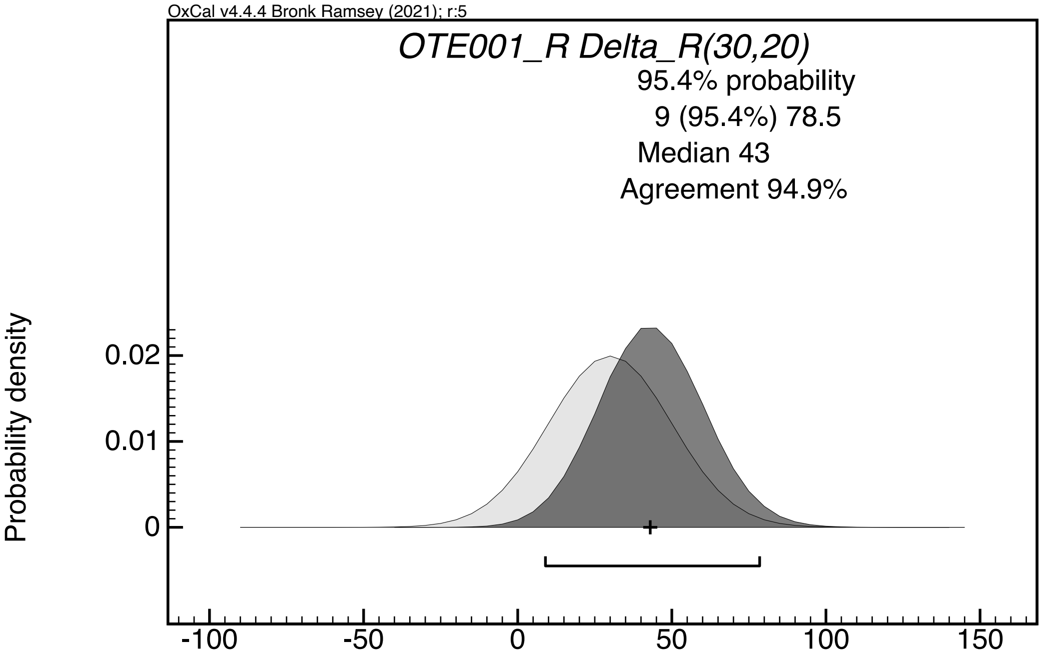

**Fig. S58: Delta_R offset of OTE001 with prior distribution (light grey) and posterior distribution (dark grey).**

For OTE001, the posterior 2-sigma distribution of this model is 1363–1394; the posterior interval of MAN008 is shifted to 1361–1392 (without offset) and 1359–1405 (with offset).

Although the original radiocarbon dates, because of their wide chronological bracketing, support an association of the genomes of Manching-Pichl, Collalto Sabino and Otepää with, potentially, any outbreak in second half of the fourteenth century, the results of the PhIRM approach rather suggest an association of these genomes with post-*pestis secunda* ones. Moreover, during the *pestis secunda* wave, plague reached Bavaria in 1357, making it one of the first German regions to be affected. If the Manching-Pichl genome were associated with the *pestis secunda*, we would expect to find it positioned basally somewhere between N03 (the 1A–1B split between the Black Death and *pestis secunda*) and N28 (the Bergen op Zoom genomes, dated, most likely, to 1359).

To estimate the most likely association of OTE001 – and other closely positioned genomes MAN008 (Manching-Pichl, in Bavaria) and COL001 (Collalto Sabino, in Lazio, Central Italy) – with particular outbreaks, it is important to consider the spatio-temporal contours of both the third (*pestis tertia*) and fourth (*pestis quarta*) waves of the Second Pandemic, and have the phylogenetic positioning of these genomes matched against them.

Contrary to the two preceding waves (the Black Death of c.1338–1353 spreading in a 'linear' manner and the *pestis secunda* of 1356–1366 spreading in a 'concentric' fashion), the *pestis tertia* appears to have been characterized by a long and repeated circulation in the German territories, attacking, in the course of its journey, some regions twice. Also, contrary to the *pestis secunda*, it appears to have commenced either in Frankonia or northern Swabia, rather than in Hesse, in 1362. In the same year, the plague was reported in Nuremberg and Augsburg (*Die Chroniken Der Fränkischen Städte. Nürnberg. Erster Band* 1862, 64.20, *Die Chroniken Der Schwäbischen Städte. Augsburg, Vierter Band* 1894, 29), spreading into the Würzburg region in 1363 (Fries 1994, 362), and then into Hessan and eastern Rhineland-Palatinate towns, including Frankfurt am Main, Fulda and Mainz (Papke 1947, 8; Zeiller 1646, 38; *Die Chroniken Der Mittelrheinischen Städte. Mainz, Zweiter Band* 1882, 167).

The spatio-temporal discrepancy between the *pestis secunda* and *pestis tertia* is of significance, and it may reflect a possibility that the reservoir may have moved south-eastwards between its initial seeding, presumably in southern Hesse, during the Black Death outbreak there in 1349 and the commencement of the *pest tertia* wave in 1362, potentially expanding its range and seeding several local micro-foci in southern Germany. There are about 225 km between Frankfurt am Main and Nuremberg and 360 km between Frankfurt am Main and Augsburg. The median distance of ~290 km and 13 years (as the absolute maximum between 1349 and 1362), hints that if there indeed was that hypothesized migration of the wild rodent reservoir from its initial home in Hesse to a new one in or around the Nuremberg-Augsburg region, then it may have moved at the speed of about ~22 km a year - certainly compatible with the annual speed of 10–120 km a year reported for prairie dog reservoirs in the western US in the later 20th century (Adjemian et al. 2007). Also, given that as of 2023, of 56 sequenced genomes from the post-*pestis secunda* period, not a single one falls on the 1B branch hints that the putative Hessan reservoir, associated with the *pestis secunda* and 1B, may have died out shortly after 1356, with 1B strains commencing their eastward migration, to re-emerge during the Third Plague Pandemic. Meanwhile, 1A strains may have continued their southbound migration into their new reservoir, seeded in or around the Nuremberg-Augsburg region at some point between 1349 and 1362. Indeed, as we shall see, the subsequent late-medieval and early-modern waves appear to have commenced in South Germany, indicating that the same putative reservoir may have persisted for several centuries. However, without securely dated aDNA material from specifically that region, this scenario remains, at this point, purely hypothetical.

Between 1364 and 1368 the plague was spreading northwards, westwards and eastwards through most of the German *Reich*; there is, however, no evidence of its concurrent southbound movement. It reached Bremen in 1366, Lübeck in 1367 and Stralsund a year later (Schwarz 1996, 104–105; Ibs 1994, 99–104). From the Baltic coast, it appears to have spread in two directions: (a) northwards into Scandinavia (Bisgaard 2009, 97; Myrdal 2003, 58–61, 127); and (b) southwards along the Oder into Brandenburg, Silesia and Bohemia (Białecki and Rymar 2005, 540–545). Traversing along the Oder route, the plague reemerged in Bohemia (Emler 1884, 540–545) and the Salzburg region in 1369, and a year later (1370) in Vienna (Fuhrmann 1738, 552), before reaching Regensburg in the following year (1371; Leidinger 1918, 174). It is unclear if the 1371 outbreak in Regensburg and its subsequent northbound spread via eastern parts of Bavaria, reflects either a commencement of a new wave, or continuation of the *pestis tertia*. Importantly, there is no evidence of a commencement of a new wave or any outbreak around the putative new reservoir in or around the Nuremberg-Augsburg region in or around 1371 (and indeed until 1378–1379, as noted below). The 1371 outbreak is the earliest recorded event after the beginning of the pestis tertia in 1362 that may potentially be associated with the MAN008

genome – although, as we shall suggest below, the said genome could have been associated with a later outbreak in the Munich region, occurring, sequentially, in 1380 and 1396 (see below, this section).
Concurrently, the *pestis tertia* wave was making its way into the West Mediterranean, reaching Italy via two routes. Firstly, it reached Lombardy and Venice, seemingly via Tyrol, in 1371. Secondly, it reached Genoa in 1372, with South-East France (Auvergne-Rhône-Alpes and Alpes-Côte d'Azur provinces), Catalonia and Corsica remaining equally possible candidates of pathogen importation. From North Italy, it penetrated into Parma, Bologna, Pisa and Lucca in 1373, before spreading all over Central and South Italy, including the Lazio region in the following year (1374; Bethmann 1866; Slavin 2022, 338–340; Cohn 2002, 164). As we shall argue below, the COL001 genome may be associated with the 1374 outbreak.

As far as an outbreak in Estonia is concerned, the textual evidence in the fourteenth century is frustratingly thin, compared to later centuries. The only documented case is the outbreak of 1378/1379 in the bishopric of Tartu/Dorpat (Napiersky 1846; von Bunge 1857, 346; Koppmann 1884, 563; Schwalm 1895, 73). It is unclear how the plague entered the Tartu/Dorpat bishopric, but three possible routes are to be considered. Firstly, it could have potentially come from the west, via the Hanseatic-Pomeranian route. In 1376, the plague is reported in Lübeck and Stralsund (Ibs 1994, 104–105). Secondly, it may have entered Estonia from Sweden, where one witnesses an increased number of annual wills and dated gravestones in 1378 (Myrdal 2003, 58–61, 127), or, alternatively, from Finnish territories – say, from Turku. However, it is not until the early 15th century that we have any textual evidence of plague there (Aalto 2017). The third possibility is that it could have arrived from the east via the Moscow-Tver'-Pskov route. Here, the plague is attested in Central Russia in Summer 1374 (*Letopisnyi Sbornik Imenuyemyi Patriarsheyu Ili Nikonovskoyu Letopis'yu* 1885, 21, *Rogozhskii Letopisets* 2000, 106), Tver' in 1375 (*Rogozhskii Letopisets* 2000, 112) and in Smolensk in Summer 1377 (Bychkov and Bestuzhev-Ryumin 1889, 103). There is no information about the situation in nearby Pskov, 140 km to the south-east of Tartu/Dorpat: unfortunately, all three versions of the Pskov Chronicle are missing a 1378–1379 entry (6887, corresponding to 1 September 1378 – 31 August 1379, according to the Byzantine-Old Russian calendar; Nasonov 1955, 29 and 106; Nasonov 1955, 106; Pogodin 1837, 35). Importantly, if the 1378/1379 outbreak was imported from Pomerania or Scandinavia, where it is attested shortly before the same outbreaks, then it is to be associated with the *pestis quarta*, rather than the *pestis tertia*, as the latter visited the region in the later 1360s, rather than the 1370s, as shown above. If we are to associate both the COL001 and OTE001 with the same wave, attacking both regions in the 1370s (in 1374 and 1378/1379, respectively), then the same wave is more likely to be associated with the *pestis tertia*, implying that the plague came to the Tartu region from the east (the Russian principalities), rather than from the west (Pomerania or Scandinavia).

Although the association of the COL001 and OTE001 genomes with the *pestis tertia*-associated outbreaks of, respectively, 1374 and 1378/1379 remains a plausible scenario, we may also consider a less likely possibility of the genomes to be linked to the subsequent wave - the *pestis quarta*. For COL001, we hear about a plague outbreak in Lazio in 1384, reported in both L'Aquila and Rome (Cassese 1941, 769; Muratori 1734), with Collalto-Sabino situated halfway between both cities, about 70 km from each. As far as OTE001, the region appears to have been affected by plague around 1389–1390, as witnessed in Pskov chronicles, describing an outbreak in the city in that year. Although there is no direct evidence from the Tartu region in general and Otepää in particular, the latter's proximity to Pskov (~140km) may indicate that the plague spread to that region, too.

The *pestis quarta* wave, associated with the 1384 outbreak in Lazio and the 1389–1390 one in Pskov (and possibly in Otepää), appears to have commenced in South-Central Germany either in late 1378 or in 1379. The wave's westbound and southbound spread was remarkably fast: having passed through Saarland and Lorraine, the plague was in Paris already by August 1379 (Delachenal 1916 Vol. 2, 362–363, 382). In the south, having crossed the Alps in 1381, it made its appearance in northern Italy in the following year (Nada Patrone and Naso 1978, 34). From there, it spread into Liguria, Emilia-Romagna and Tuscany in 1383, before arriving in Central Italy, including Lazio, in 1384.

By contrast, its northbound spread into the Baltic coast was slow, and it was not until 1387 that it appeared in Hamburg and Wismar and 1388 in Lübeck, where it allegedly killed 16,000 people (Koppmann 1899, 8, 24; Ibs 1994, 105–107). In the same year, the plague crossed to southern Denmark and southern Sweden (Bisgaard 2009, 97; Myrdal 2003, 58–61, 127). Although it is possible that the plague came to Pskov (and potentially Otepää) either from the West – from Pomerania or from Scandinavia – it may, alternatively, have spread from the South, from Ruthenian territories, via Smolensk. Several Russian chronicles mention a devastating plague in Smolensk, dated by some to 6894 (1385–1386) and by some to 6895 (1386–1387; *Simeonovskaya Letopis'* 1913, 136, *Pskovskiya I Sofiiskiya Letopisi* 1851, 5:242–243).

Given their close phylogenetic positioning, we may assume COL001 and OTE001 are associated with the same wave - either *pestis tertia* or *pestis quarta*. In the case of the former, the respective outbreaks occurred in 1374 and 1378/1379, while in that of the latter – in 1384 and 1389–1390. One indirect argument against the identification of OTE001 with the *pestis quarta* is the fact that (1) the excavated site contained a bracteate minted 1370–1375 and several others from the second or third quarter of the 14th century, and (2) the use of north German bracteates fell out of circulation in the course of the 1370s following their ban in Livonia by the Livonian Diet in 1374 (Molvygin 1969, 50–51). Although the ban was imposed in 1374, it takes, as a rule, several years for discontinued coins to fall completely out of circulation, meaning that the north German bracteates may, in fact, have continued to circulate until c.1380 – but not until c.1390. Hence, a plausible association of COL001 and OTE001 is with, respectively, the 1374 and 1379–1380 outbreaks, is more likely than with the 1384 and 1389–1390 ones. If this is correct, then N04 may represent the beginning of the *pestis tertia* wave.

The MAN008 genome may be trickier to associate. On the one hand, it could be associated with the 1371 outbreak, reported in Regensburg, the best-known geospatial proxy for MAN008. As mentioned above, it is unclear if the 1371 outbreak (and its subsequent northbound spread via eastern parts of Bavaria in the 1370s), reflects either a commencement of a new wave, or a continued (and intense) circulation of the *pestis tertia* in German territories. One argument against the beginning of the new wave is the fact that there is no historical evidence of a plague outbreak anywhere in the Nuremberg-Augsburg region, a putative home of a local reservoir, in 1371 or in the immediately preceding years. As shown above, one possibility is that the 1371 outbreak was imported into Regensburg and other nearby regions in Bavaria from Bohemia or Austria. However, there is also a possibility that the 1371 outbreak represents a later spillover event, whereby N04-associated strains migrated into the Regensburg region from their putative reservoir in/around the Nuremberg-Augsburg region. The distance between Regensburg on the one hand and Nuremberg and Augsburg on the other is, respectively, 110 km and 150 km. This may also hint that 6 individual SNPs of MAN008 were acquired while plague was circulating in rodents, before crossing to humans. If N04 is indeed associated with the beginning of the *pestis tertia* in 1362 and MAN008 is indeed associated with the 1371 outbreak, then the acquisition of 6 SNPs in the course 9 years implies a very rapid substitution rate of one SNP every 1.5 years - about thrice as faster as the ‘average’ speed of one SNP every 4.8 years along the entire length of 1A branch (between the Black Death and the ‘Little Polytomy’, or N13; see Supplementary Section 11.7). The elevated speed of substitution rate on the MAN008 terminal sub-branch has been already noted in its original publication by (Spyrou et al. 2019), and it may indeed reflect the circulation and evolution of *Yersinia pestis* strains in wild rodent reservoir, before spilling over to humans. Indeed, as some evidence from natural plague reservoirs in Qinghai Province and Madagascar show, the speed of SNP acquisition in wild rodent reservoirs appears to be faster compared to that when circulating in humans (Rahelinirina et al. 2017; Dai et al. 2018).

One other possibility is that the MAN008 genome is associated with a later outbreak. The closest documented outbreak for Manching-Pichl, in space and time, is from Munich in 1380, associated with the *pestis quarta* wave, whose beginnings are mentioned in Swabia and the Nuremberg region in 1378–1379, and which spread to Osterhofen, Augsburg, as well as the Munich and Regensburg regions in 1380 (*Die Chroniken Der Fränkischen Städte. Nürnberg. Erster Band* 1862, 354, *Die Chroniken Der Schwäbischen Städte. Augsburg. Erster Band* 1865, 66, *Die Chroniken Der Mittelrheinischen Städte. Mainz, Zweiter Band* 1882200)(Solleder 1938, 391–392; S. Krämer 1972, 243)(*Die Chroniken Der Fränkischen Städte. Nürnberg. Erster Band* 1862, 354, *Die Chroniken Der Schwäbischen Städte. Augsburg. Erster Band* 1865, 66, *Die Chroniken Der Mittelrheinischen Städte. Mainz, Zweiter Band* 1882, 200). Given a short time-lag between the beginning of the wave and its spread to the Munich region, we would expect the MAN008 genome to be positioned basally on its branch. However, as MAN008 is, at present, the only genome representing this lineage, an approximate terminal branch length of MAN008 relative to a hypothetical genome representing the initial outbreak of this wave cannot be determined. The next outbreak in Munich took place in 1396, still falling within the brackets of its 2-sigma range dating (1359–1405; Solleder 1938, 391–392; Krämer 1972, 243). It is unclear, at this point, if the 1396 outbreak is associated with the fifth wave, the *pestis quinta*, which may have commenced in 1388 in its South German reservoir, as attested in Augsburg, Nuremberg and Heilbronn sources (von Oefele 1763, 327; Mencke 1728, 1:1531, 1538; *Chroniken von Closener und Koeningshoven* 1845, 172; Steinhilber 1956, 198), or with a later wave of the 1390s, whose origins cannot, at this point be established, but which may have possibly commenced in South Germany in 1395 (*Die Chroniken Der Fränkischen Städte. Nürnberg. Erster Band* 1862, 88, 94).

#### 11.4 The AHM genomes and MAL003

The genomes of Arnhem, Netherlands (AHM001 and the identical low-coverage genome of AHM011, see Supplementary Section 7) as well as the genome of Mäletjärve, Estonia (MAL003) occupy positions on a node (N06, Fig. 2A) and a terminal branch on Branch 1A, respectively, which were previously not represented by ancient genomes. As AHM001 does not show any private mutations, it is well-suited as calibration point for the relative chronology as utilized by PhIRM. Prior to the genome reconstruction and the radiocarbon dating, the burial was dated 1650–1829, i.e., the later phase of the cemetery, based on the stratigraphy. Radiocarbon dating resulted in 2-sigma intervals of 1307–1416 (AHM001) and 1300–1399 (AHM011), clearly suggesting an older date, which is in line with its phylogenetic position, only 12 SNPs derived from the European Black Death cluster and ancestral by at least 32 SNPs to the genomes of the Thirty Years' War plague of the early 17th century (Fig. 2A). The sequence model with offset (Fig. 3, Code S3, Table S11, Fig. S46 left) narrowed down the intervals to only 33 years (1385–1418) and 39 years (1377–1416), respectively.

The genome of Mäletjärve MAL003 has accumulated at least one SNP passing filtering (11 SNPs did not pass filtering, and 3 were classified as potential damage, see Table S5). The radiocarbon interval (2-sigma: 1425–1472) situates this sample in the mid-15th century, while PhIRM with generic offset yielded a much broader interval (1429–1622). Considering a branch length of maximum 4 SNPs (Table S5), the bimodal distribution of the posterior probability with two peaks in the late 15th and early 17th century (Fig. 3), and the bracteate with MAL003 dating to 1379–1420 (see also Supplementary Section 2.4), a date within the 15th century appears more likely. Moreover, the fact that nine SNPs separate N04, leading to the outbreaks associated with the MAN008, and COL001/OTE001 (most likely, in the context of the *pestis tertia* commencing in 1363, or, less likely, in the context of the *pestis quarta* commencing c.1377–1379) on the one hand, and N06, leading to AHM001/AHM011 and MAL003, on the other, may potentially narrow down the genomes to c.1400–1420.

Given that MAL003 is directly derived from AHM001/AHM011 and has accumulated a low number of SNPs (4 of 15 private SNPs passing filtering, thereof 3 are potential false-positives due to DNA damage), it is possible that they constituted a part of the same wave. Moreover, the basal positioning of AHM001/AHM011, compared to a more derived one of MAL003, implies that the former's associated outbreak came before that of the latter, which is further corroborated by Arnhem's closer proximity to the putative South German reservoir compared to Mäletjärve. There are several possible candidates to be associated with the genomes.

One possibility is that the AHM001/AHM011 genomes are associated with a 1398 outbreak, reported in nearby Deventer (~45 km north of Arnhem; Mertens 1999), while MAL003 is to be identified with a 1404 outbreak reported in Tartu/Dorpat and Pskov (Nasonov 1941, 27; Nasonov 1955, 31, 111). Although Pskovian chronicles allege that the plague arrived in Pskov from the German settlement in Tartu/Dorpat (in summer 1404), it may reflect an anti-German prejudice of local authors: there is no evidence of plague circulation on the southern Baltic shore, along the Hanseatic trade routes, between 1396 and 1405 (Ibs 1994, 107). Conversely, plague is attested in Smolensk in Autumn 1401 (Buganov and Rybakov 2002, 168), implying possibly a southern, rather than a northern route of introduction. In any event, the 1398 outbreak in Arnhem and the 1404 one in Tartu/Dorpat both seem to have been associated with a wave, which commenced in the putative South German reservoir in 1388, as the evidence from Nuremberg, Augsburg and Heilbronn indicates (von Oefele 1763, 327; Mencke 1728, 1:1531, 1538; *Chroniken von Closener und Koeningshoven* 1845, 172). However, while the 1398 outbreak in the Netherlands would be a good fit regarding the PhIRM dating of Arnhem, an association of the MAL003 genome with the plague outbreak reported for Tartu/Dorpat in 1404 contradicts both the unmodelled radiocarbon date and the PhIRM modelling.

The second possibility is that the AHM001/AHM011 and MAL003 genomes may have been associated with a wave, which commenced in the putative South German reservoir in 1405, as the evidence from Nuremberg indicates (*Die Chroniken Der Fränkischen Städte. Nürnberg. Erster Band* 1862, 96), spreading all over Germany in the course of the next few years. If this hypothesis is correct, then the Arnhem genomes may be associated with a c.1410–1412 outbreak documented in the region. Here, the plague was reported in Wesel in 1410 (Jankrift 2020, 196–197, 384), in Breda and Leiden in 1411 (Gooskens 1987, 31; Ladan 2012, 234) and in Gorinchem in 1412 (Zuijderduijn 2013, 21; Curtis and Roosen 2017 Appendix 1, p. 19). The Mäletjärve genome may be identified with a 1420–1421 outbreak in Tartu/Dorpat. According to one Hanseatic document, the plague was present in Tartu/Dorpat, Tallinn/Reval, Riga, and surrounding regions in winter 1420–1421 (*Die Recesse Und Andere Akten Der Hansetage von 1256-1430. Band VII* 1893, 165, 171), while Pskovian chronicles alleged, again, that the plague came to Pskov in summer 1420 from the 'Germans of Tartu', and remained in the city until early January 1421 (Nasonov 1941, 34; Nasonov 1955, 37–38). One other Russian chronicle stated that the plague was present in Tartu/Dorpat, Pskov, Novgorod, Torzhok and Tver' already in August–September 1419 (*Letopisnyi Sbornik Imenuyemyi Patriarsheyu Ili Nikonovskoyu Letopis'yu* 1885, 286), although it may be a misdating, with 1420 being a correct date. Indeed, a discrepancy of a year and sometimes two years between various Russian chronicles is a commonplace. The statement that the plague came to Pskov from the 'Germans of Tartu' may, just as in the case of the 1404 outbreak, reflect anti-German prejudice, rather than reality. Assuming the westbound movement, we would expect the plague to be mentioned in Lübeck or some other Baltic city before its arrival in Tartu/Dorpat in Summer 1419 or 1420. However, it is not attested there until 1420–1421, and there is no evidence of any plague outbreak in any neighboring regions in the preceding years. Rather, the 1420–1421 outbreak in Lübeck may be associated with a different plague wave, commencing in 1418–1419 in South Germany (Abel 1852, 44; *Die Chroniken Der Schwäbischen Städte. Augsburg. Erster Band* 1865, 119; Müllner 1984, 231) and spreading remarkably fast all over Germany in 1419–1420, reaching Lübeck in either late 1420, or 1421 (Ibs 1994, 112). It is likely, therefore, that the 1419/1420 outbreak in Tartu/Dorpat came from the south, possibly via Bohemia (attested in 1414–1415; Maur 1989), Silesia (attested in 1416; Kehrberg 1724 II: 80), and Ruthenia (no data). In summer 1417, the epidemic arrived in north-western Russian cities, initiating the so-called 'St Anastasia's Pestilence' ravaging Russian lands for the next few years (*Letopisnyi Sbornik Imenuyemyi Patriarsheyu Ili Nikonovskoyu Letopis'yu* 1885, 232–233), and it was just a matter of time before it reached Tartu/Dorpat – either in summer 1419 or 1420. The outbreaks in question appear to be associated with the same wave commencing in South Germany in 1405 that was responsible for the 1410–1412 outbreak in the Netherlands, discussed above. Here, again, the PhIRM interval for AHM001/AHM011 would be compatible with an outbreak in 1410–1412, whereas the outbreak reported for 1420–1421 in Tartu/Dorpat falls slightly outside of the PhIRM and raw radiocarbon interval of MAL003.

The third possibility, albeit a less likely one (assuming the upper range of the Mäletjärve bracteate minted between 1379 and 1420), is that the genomes may have been associated with the subsequent wave, which appears to have commenced, as noted, in 1418–1419, spreading quickly all over Germany and the Low Countries in 1420–1421. In Arnhem, it is directly attested in 1421 (Van Veen 1903, 4–10; Curtis and Roosen 2017 Appendix 1, p. 21–22), while in Tartu/Dorpat it was reported, in one Hanseatic document, in October 1425 (*Die Recesse Und Andere Akten Der Hansetage von 1256-1430. Band VII* 1893, 598–599), while Pskovian chronicles mention the outbreak in their city in 6933 (1 September 1424 – 31 August 1425; Nasonov 1955, 40, 121). Given a gap of some four years between the 1420–1421 outbreak in Lübeck and the 1424–1425 outbreak in Tartu/Dorpat and Pskov, it is clear that the pathogen had not been imported directly there from the Hanseatic city via the maritime route, but (at least partially) via the inland one – by the way of Pomerania, Poland, Lithuania and Ruthenia. Indeed, in 1422 the plague is reported in Toruń (Baczkowski and Turkowska 2000, 181), an important trade centre connecting Hanseatic cities with North-western Russia. If the AHM001/AHM011 and MAL003 genomes are indeed associated with this wave, then they may be identified with the 1421 and 1424–1425 outbreaks in their respective locations. In this case, the identification of MAL004 with the 1424–1425 outbreak assumes that the bracteate (minted between 1379 and 1420) is dated towards or around its upper dating range and remained in circulation for a few extra years.

#### 11.5 STA001, CLP15, CLP017 and G701

The genomes of Starnberg, Germany (STA001), Clopton, Cambridgeshire, United Kingdom (CLP015, CLP017, both identical), and G701 from Riga, Latvia all represent offshoots from the basal Branch 1A prior to the polytomy N13 with short terminal branches.
STA001 was published in Spyrou et al. 2019 with a dating interval of 1433–1523. STA001 corresponds to individual 207 of a simultaneous triple burial, containing also the individuals 208 and 214. The dating interval is based on the radiocarbon date of individual 214 and on a rosary (*Paternoster*) found with it. According to the archaeological publication (Later 2010), this type of rosary first appears in the 15th century, in line with the earliest 1-sigma radiocarbon interval of 1433–1494 (with 59.6% probability, and 8.7% probability for 1601–1615). Regardless of that, in Spyrou et al. 2019, the upper boundary of the earliest of the 2-sigma intervals was used (1420–1523 with 74.4%, 1557–1564 with 1.1%, 1569–1630 with 19.8%).
The recalibrated 2-sigma radiocarbon interval of this individual is 1422–1631, PhIRM shifted the interval to 1438–1636 due to the generic offset. However, the posterior probability distribution is bimodal with 54.7 % likelihood between 1437–1531 (see Fig. S59), which is in line with the dating of the rosary.
Phylogenetically, STA001 is 7 SNPs derived from N06 corresponding with AHM001/AHM011: two SNPs are shared with other younger Branch 1A genomes (except MAL003), and 5 SNPs define the terminal branch of STA001.
The fact that N06 and N07 are separated by only 2 SNPs implies that they may represent the emergence of two waves chronologically close to each other. If this is the case, then STA001 may be associated with a wave following that of AHM001/AHM011 and MAL003. However, given the radiocarbon dating of STA001 above and that of the Clopton genomes (CLP015 and CLP017) below, it is unlikely that the genome is any earlier than c.1420 and

later than c.1440. While there are no written testimonies of plague specifically from Starnberg, we can use evidence from nearby Munich (about 26 km northeast of Starnberg), assuming that the same outbreaks reported in Munich also occurred in Starnberg. Here, we hear about plague visiting the city in 1420, 1430 and 1439 (Solleder 1938, 391–392; S. Krämer 1972, 243).

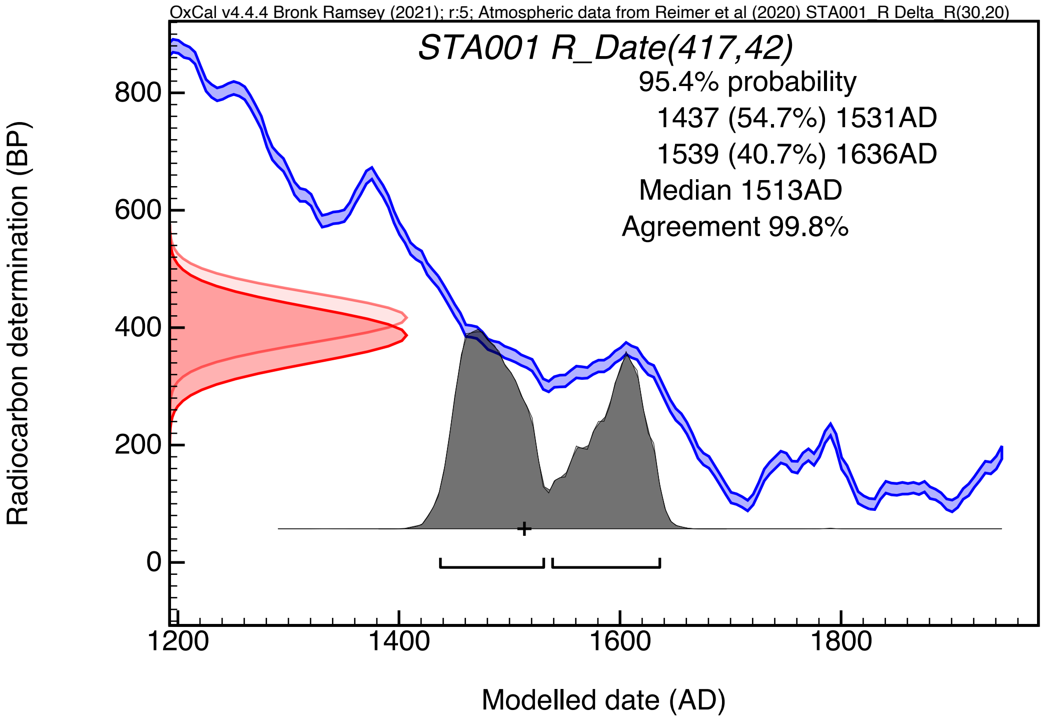

**Fig. S59: probability distributions (brackets indicating 2-sigma intervals) of STA001 within the sequence model applying a generic offset (Fig. 3, Code S3, Table S11, Fig. S31 left).**

Since the two genomes from Clopton CLP015 and CLP017 are identical, we can assume that they belong to the same local plague outbreak. The archaeological dating ofthe Clopton plague victims is relatively broad, with a *terminus ante quem* of 1561 for the usage time of the cemetery (Cessford et al. 2021). The radiocarbon dates of the Clopton plague victims prior to modelling hint towards the first half of the 15th century (CLP015: 1405–1443, CLP017: 1317–1425, both 2-sigma, Fig 2A, Table S11). Applying PhIRM on those dates results in slightly younger intervals (1413–1458 and 1406–1444, respectively). However, given that the STA001 genome does not come, in all likelihood, from a pre-1420 context (but rather, as suggested above, is dated c.1420 – c.1440), and that the 'Little Polytomy' (N13), discussed in Supplementary Section 11.7, emerged likely before 1473, we may perhaps date the Clopton genomes to c.1438 – c.1458 – that is, some twenty years after the Starnberg genome. At first sight, such estimated chronological distance between the two genomes may appear too narrow, given that there are 6 SNPs separating N07 (leading to the Starnberg genome) from N08 (leading to the Clopton ones), implying a substitution rate of one SNP every 3.3 or so years – lower than the average 4.8 years for Branch 1A, taking it as a whole from N3 (the Black Death genomes) and N13 (the 'Little Polytomy', emerging, at some point, between  c.1450 and c.1500; see Supplementary Section 11.7). However, as Spyrou et al. 2019's analysis has established, there appears to be an acceleration in SNP substitution rates between N07 and N10, which is consistent with a proposed chronological distance between N07 and N08.

Unfortunately, our information on plague outbreaks in Cambridge before the early 16th century (with the exception of the Black Death) is extremely patchy, owing to poor documentation. Nevertheless, some hints can be gleaned from a few local sources, as well as from other East Anglian cities, first and foremost Ely and Norwich. A close analysis of all the available information yields several possible scenarios. One possible scenario is a nationwide (and indeed pan-West Eurasian) outbreak in 1438–1439 in England. Next, we know that in 1441 King Henry VI sent the Marquis of Suffolk to Cambridge to lay the foundation stone of King's College instead of coming himself, fearing the pestilential air that has long been reigning in the university (Cooper 1842 Vol.1, 198–199). This is corroborated by a peak in the annual number of wills probated at the Bishop of Norwich's court in 1442–1443 (Farrow 1943-1945). Then, we identify another spike in the number of annual wills in Norwich in 1456–1458, in Bury St Edmunds in 1458, and we hear about plague in Ely and elsewhere in East Anglia in 1458–1459 (Redstone 1907; Farrow 1943-1945; Atkinson 1933 Appendix X; *Abbreviata Cronica ab Anno 1377 usque ad Annum 1469* 1840: 7). Finally, plague returned to Norfolk in 1465, as reflected in another spike in the number of wills (Farrow 1943–1945). In addition, there is a local story of an old man walking along Cambridge colleges in 1462 and foretelling a great plague and famine to occur within the next two years (Gardiner 1880, 163). Indeed, the 1463–1465 outbreak, reflected in the Norfolk records and the Cambridge story, appears to have been on a national scale. However, the 1463–1465 outbreak may be too late to be associated with the Clopton genomes.

The plague genome of G701 from Riga was published by Susat et al. 2020, alongside another Riga genome (G488). Although both Riga genomes come from the same site (Riga St Gertrude Church), they derive from two different burial contexts: while G488 comes from 'mass grave 1', G701 was unearthed from a 'burial pit'. While the chronology of G488 (c.1550–1650) has been determined by both archaeological evidence (based on artifacts) and radiocarbon dating, regarding the G701 genome, the authors state that the 'chronology of the small mass burial pit with 15 buried individuals is not entirely clear'. Nevertheless, they date G701, too, to c.1550–1650 (Susat et al. 2020 Fig. 2), without justification.

In the phylogenetic tree G701 is framed by the branch leading to CLP015/CLP017 and the MAL003-MAL007-NMS002-G488 clade, but shows the shortest distance from the MRCA represented by N08 ($d=3$ compared to $d=4$ for CLP015/CLP017, $d\geq6$ for MAL004). Considering the dating interval of CLP015/CLP017 in the mid-15th century and the Boundary T5 interval of 1406–1457, representing the lower boundary of the Branch 1A3, a date within the interval of 1550–1650 for G701 seems highly unlikely when assuming a roughly clock-like accumulation of substitutions. In conclusion, we assume that the mass grave 1 and the burial pit of St Gertrude Church are not contemporary with each other, but represent two funerary structures established during two separate plague outbreaks.

Given that (1) G701 phylogenetically derives from N09; (2) G701, positioned on the terminal branch resulting from N09, has only one unique SNP; and (3) N09 is situated only two SNPs derived from N07, we may hypothesize that G701 is to be associated with an event shortly following the event giving rise to the Clopton branch, and that the associated wave reached Riga shortly after its commencement in its reservoir. Assuming the approximate timeframe of c.1445–1465, we may suggest several potential outbreaks in Riga reported in Hanseatic and

Polotsk records. The outbreaks in question occurred in 1453–1454, 1458, and 1464–1465 (Schwartz 1905 nos. 321, 325, 747; Von der Ropp 1888, 269, 435; Khoroshkevich et al. 2015 Vol. 1, nos. 167–168).

The first scenario (that of 1453–1454) is a possible one. The outbreak in question seems to have been associated with a wave commencing in South Germany in 1449 (Weiland 1877, 379; Müllner 1984, 409). In the course of 1449–1450, it spread all over Germany, reaching the Baltic coast in 1450 and ravaging coastal cities, including Lübeck in 1450–1451 (Ibs 1994, 113–116). From there, it spread northwards into Denmark (1450–1452; Bisgaard 2009, 97, 103), and subsequently Sweden (1451; Myrdal 2003, 68, 79–82, 85, 110). Concurrently, the pathogen spread eastwards, reaching Gdańsk/Danzig in 1450, before continuing southwards into mainland Poland, reaching territories around Warsaw in 1451 (Możejko 2012, 43; Walawander 1932, 164). From there, plague reached Lithuania in 1452, ravaging the local population in 1452–1453 (*Kronika Polska, Litewska, Zmodzka I Wszystkiej Rusi Macieja Stryjkowskiego* 1846 Vol. 2, p. 234), and moving northwards into Latvia in 1453 (Schwartz 1905 no. 321), Estonia and then the Pskov region in 1454 (Schwartz 1905 no. 325). Given the spatio-temporal contours of the wave, it is likely that the plague reached Riga via inland routes from Lithuania, rather than by sea from one of Hanseatic ports or Sweden. The time-lag of four years between the inception of the wave and its arrival in Riga seems to be a good fit for G701 to have acquired only one unique SNP in relation to N09.

The possibility of the 1458 outbreak is less likely. The same outbreak appears to have been exported from the Balkans, in the context of the ongoing Ottoman-Hungarian war, centred around the Siege of Belgrade, where plague had broken out in late July 1456 in the Serbo-Hungarian camp (Wattenbach 1851, 519). It is unclear, at this stage, if the same outbreak was associated with the same wave commencing in 1449, or if it was a different wave originating in a different reservoir that had in a meantime been seeded somewhere outside of Central Europe, possibly in the Balkans or further south-east in the expanding Ottoman Empire. If the former scenario is correct, and if G701 was indeed associated with the 1458 outbreak, then it would have been on a more derived phylogenetic position than what it is, with additional unique SNPs. Therefore, it is unlikely that G701 is associated with the 1458 outbreak.

The 1464–1465 outbreak appears to be associated with a wave commencing in South Germany in later 1462 (Mencke 1728, 1:1648, 1652). In the course of 1463, it spread remarkably fast all over Germany, reaching Lübeck in May of the same year, before crossing into different Baltic regions (Ibs 1994, 117–120). In the north, it is attested in Denmark in 1464 (Bisgaard 2009, 97, 104) and Sweden in 1464–1465 (Myrdal 2003, 68–69, 81–82, 85, 110). In the east, it arrived in Gdańsk/Danzig at some point between April and August 1464 (Możejko 2012, 53–55), spreading all over Poland in 1464–1465 (Walawander 1932, 167–170), and then into Lithuania in 1465 (*Kronika Polska, Litewska, Zmodzka I Wszystkiej Rusi Macieja Stryjkowskiego* 1846 Vol. 2, p.267). There is a reference to plague in Livonia in Summer 1464, without a specific location (Von der Ropp 1888, 269), although its presence in Tallinn/Reval is clearly indicated in the Autumn of the same year (Tallinna linnaarhiiv, F190, N1, S60, fol. 28v; Tallinna linnaarhiiv, F191, N2, S1, fols. 31–31a; (Hildebrand 1889 no. 922). As far as Riga is concerned, it is mentioned in several documents in early 1465, but it appears to have reached the city by Autumn 1464, given the references to plague in Livonia and Tallinn/Reval in the second half of 1464 (Khoroshkevich et al. 2015 Vol. 1, nos.

167–168; Von der Ropp 1888, 435). In 1465, the plague was also mentioned in the Diocese of Turku and in the Pskov region (Risberg 2008 no. 116; Nasonov 1955, 161–162). It is unclear if the plague arrived in Riga directly from one of Hanseatic ports on the Baltic littoral, or from Sweden - directly, or possibly via Estonia. The time-lag of two years between the inception of the wave and its arrival in Riga seems to be a good fit for G701 to have acquired only one unique SNP in relation to N09.

#### 11.6 Branch 1A3 – MAL004, MAL007, NMS002.A, G488

The clade consisting of MAL004 (Mäletjärve), NMS002 (Augustinian Friary, Cambridge) and G488 (Riga), henceforth called Branch 1A3, so far represents the only known offshoot of Branch A1 prior to the polytomy N13 which survived for a longer period of time and is associated with multiple, geographically distant plague outbreaks. Given that N10 is only 2 SNPs derived from N09, and only one SNP ancestral to N13 (the 'Little Polytomy'), emerging c.1450–c.1500, we may tentatively date N10 to a rather narrow chronological bracketing on c.1440–c.1460.

The most basally branching genome of MAL004 comes from a child (Grave 39) buried next to an adult female, MAL003 (Grave 38). Due to the lack of disturbances and the fact that both individuals were buried next to each other in the same orientation on the same horizon, it was interpreted as a simultaneous double burial. However, the respective plague genomes occupy different positions in the phylogenetic tree (Fig. 2) with different genetic distances from their MRCA (N06). Since both genomes are derived from non-UDG data, the terminal branch lengths cannot be determined with certainty (Table S5). While for MAL003 4 out of 15 private SNPs passed the filtering (with 3 of 4 classified as potential damage), for MAL004 16 out of 59 private SNPs passed the filtering (with 15 of 16 classified as potential damage). The genetic distances from their MRCA (N06) therefore are 1–4 SNPs for MAL003 and 13–28 SNPs for MAL004.

This is further corroborated by the radiocarbon dates: MAL003 (2-sigma interval 1425–1472, bracteate dating to the late 14th or early 15th century, see Supplementary Sections 2.4 and 11.4) dates significantly older than MAL004 (1456–1633), while those of MAL004 and MAL007 (1461–1635) are more similar. MAL007 (Grave 28), however, belongs to a different period of the cemetery than MAL003, as indicated by a different orientation of the graves. In addition, two coins were found in the grave of MAL007, minted in 1537 and 1561–1568. We further tested the contemporaneity of MAL004 with MAL003 or MAL007 with the function "Combine" in Oxcal while applying generic offsets to none, both, or only one of the radiocarbon dates. As shown in Table S17 and Fig. 60, none of the combinations of MAL003 and MAL004 reached an agreement value over 60, while all combinations of MAL004 and MAL007 passed this threshold. The analysis of the low-coverage plague genome of MAL007 (see Supplementary Section 7) confirmed that it shares several derived positions with MAL004 and, thus, might be identical. However, the jewelry from MAL004 is not characteristic for the 16th century, but is of earlier origin than MAL007, coin-dated later than 1561.

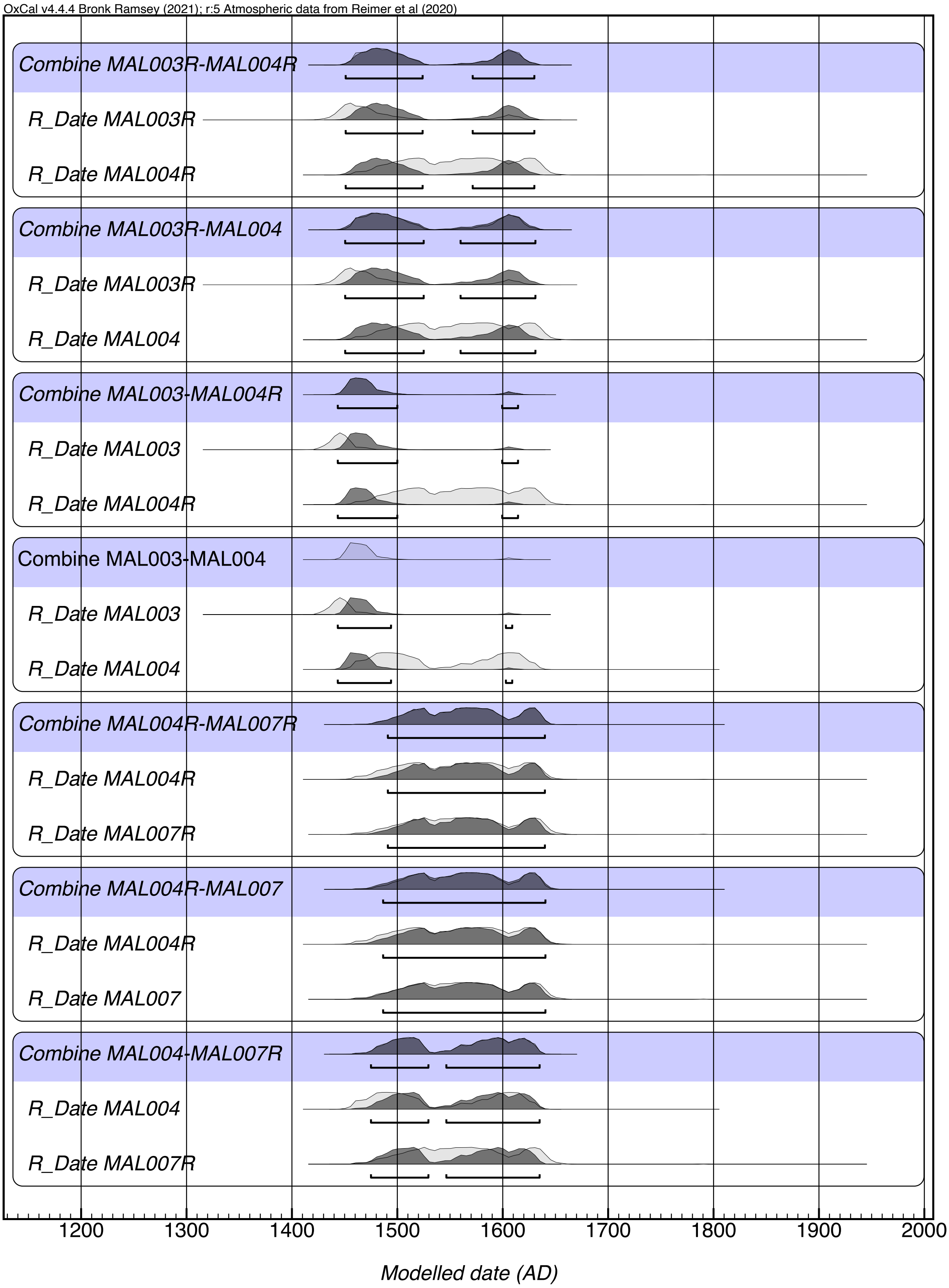

**Fig. S60: Radiocarbon intervals for the different combinations of MAL004 with MAL003 or MAL007, corresponding to Table S17.**

The identification of two vastly different *Y. pestis* genomes in Burial 38 (MAL003) and burial 39 (MAL004) led to the suspicion that the skeleton of Burial 39 might have been mixed up with another subadult skeleton of the cemetery or that a mix-up happened after sampling during laboratory work or during data processing. However, by comparison of the skeletal remains with original photographs of the excavation in 1984 we were able to exclude a possible mix-up of the skeletal remains, as the missing teeth in the jaws matched the documentation prior to sampling. Also, we were able to exclude a mix-up in the laboratory or during data processing by resampling a second tooth root and comparing mitochondrial haplogroups.
After validating that the *Y. pestis* genome of MAL003 belongs to Burial 38 and MAL004 to burial 39, two possible scenarios remain to explain the discrepancy: (1) 38 and 39 were not a simultaneous double burial but 39 was added later to the burial, although no disturbances were observed and the individuals were found on the same horizon right next to each other in the same orientation; or (2), the plague genomes stem from a simultaneous double burial with two highly divergent strains, and one of the radiocarbon date is a statistical outlier or the MAL003 appears older due to a massive offset (stronger than modeled here). Assuming the contemporaneity of 38 and 39, both strains would have had to circulate in Mäletjärve at the same time despite their disparate genetic distance.

For PhIRM, as applied for Fig. 3 and the following section, we considered the genomes of MAL004 and MAL007 as identical, which is supported by the similar radiocarbon ranges and at least 3 shared SNPs (see Supplementary Section 7), and ignored the archaeological context, suggesting that MAL004 was buried simultaneously with MAL003. Applying PhIRM on the radiocarbon dates of MAL004 and MAL007 shifts their intervals even younger to 1476–1641 and 1482–1645, respectively. The broad intervals can be explained by a wiggle in the calibration curve and the lack of a *terminus ante quem* prior. Assuming the two genomes to be identical, we may apply 1561 as a *terminus post quem* on the respective plague outbreak based on the coin found with MAL007, which corresponds well with the median of 1562 of the KDE_Plot combining the two dates (Fig. 61).

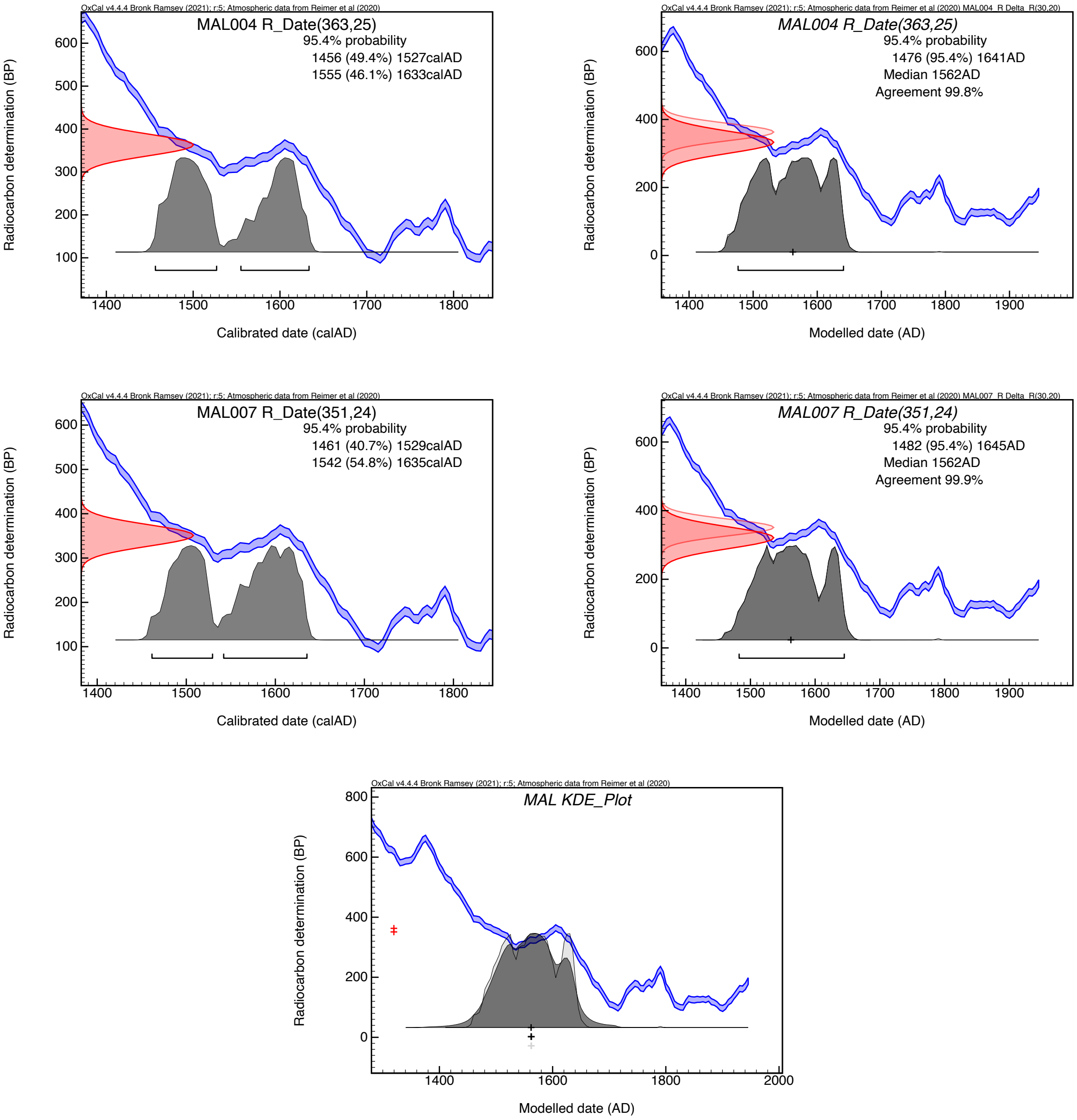

**Fig. S61: probability distributions (brackets indicating 2-sigma intervals) of MAL004 and MAL007 without offset (left) and after integrating into the sequence model with a generic offset of 30±20 RC years (right) and the respective KDE_Plot (row 3).**

Although – technically – Boundaries within PhIRM can indicate only an upper boundary for the oldest genome of a clade (and not the MRCA of a clade), the dense sampling in this part of the phylogenetic tree allows to consider the Boundary T5 as an approximation of the emergence of the Branch 1A3, represented here by node N10. Boundary T5 is constrained through the radiocarbon dates of AHM001 and AHM011 as direct ancestors on the lower end (distance 12 SNPs), through Boundary T6 and the radiocarbon dates of Branch 1A3 on the upper end (shortest distance of a derived genome – the Basal Branch 1A1 cluster – 2 SNPs). The 2-sigma interval for Boundary T5 in our sequence model is 1406–1457 with a median date of 1429 (see Fig. 62).

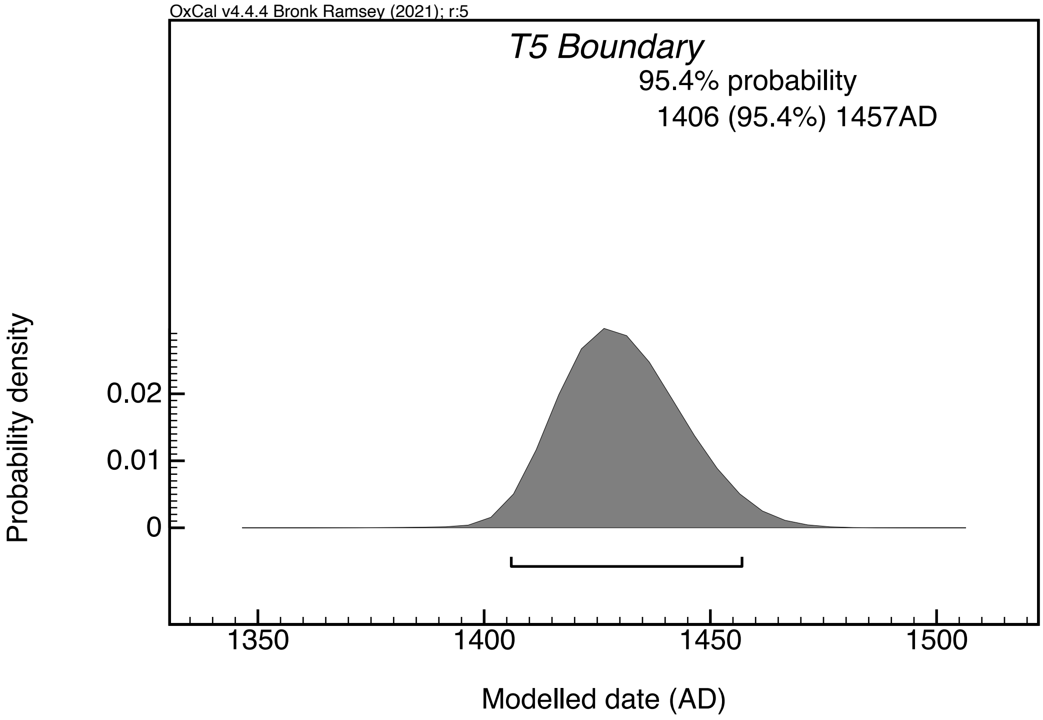

**Fig. S62: probability distribution (brackets indicating 2-sigma intervals) of Boundary T5 as part of the sequence model with a generic offset of 30 ± 20 RC years.**

MAL004 and MAL007 are positioned on a terminal branch deriving from N11. Given that N11 is only 1 SNP derived of N10 (the 1A3 branch emergence, possibly dating c.1440–c.1460), it can possibly be dated to the later 15th century, perhaps c.1460–1480. However, the fact that MAL004 has potentially as many as 16 private SNPs indicates a chronological distance of MAL004/007 from N11. According to the proposed model above, the range for MAL007 is 1482–1645, with 1562 as the median date for the KDE plot for both MAL004 and MAL007. Given the presence of two coins next to MAL007 - the one from 1537 and the other one from 1561–1568 - this wide range can be narrowed down considerably to c.1561–1575 (to allow a few extra years for a possible circulation of the 1561–1568 coin). For that period, no direct references to plague outbreaks in the Tartu/Dorpat region could be found. Hence, we may infer about their chronology from reference to plague outbreaks in other regions in Estonia and in the Pskov region. Thus, we hear about a plague outbreak in Tallinn in 1566 and 1570–1571 (Russow 158460r, 75r, 77v), in Pärnu/Pernau in 1566 (Stefan Hartmann 2008 no. 3460 (p. 78), no. 3463 (p. 79–80)), and in Pskov in 1566–1567 (Nasonov 1955, 249; Goryushkina 2021, 48–51).

The origins of the 1566–1567 and 1570–1571 outbreaks are difficult to establish. As far as the 1566–1567 outbreak is concerned, it may be associated with the wave that may have commenced in or around the Balkans in 1559 (Varlık 2015, 181), spreading into Hungary in 1560–1561, then Bratislava and Vienna (1561–1562; Magyary-Kossa 1931, 200–204; Schmölzer 2015, 81, 191). In 1562–1563, the plague was ravaging southern Germany (*Der Feind in Der Stadt. Vom Umgang Mit Seuchen in Augsburg, München Und Nürnberg. Eine Ausstellung Der Bayerischen Archivschule Der Generaldirektion Der Staatlichen Archive Bayerns* 2016, 41; Erdel, Specker, and Winckelmann 1986; Sturm 2014, 37; Eckert 1996, 81–83), before spreading all over Germany in 1564–1565 (Eckert 1996, 81–85). It was attested on the south-eastern Baltic Coast (including Gdańsk/Danzig, Elbląg/Elbing, Braniewo/Braunsberg and Königsberg) in the summer of 1564 (Stefan Hartmann 1991, 248–249; Sahm 1905, 16), but it was not until the next summer that it commenced in Riga (Stefan Hartmann 2008 no. 3389/2-3390 (p. 40)), before spreading to Tallinn/Reval and Pärnu/Pernau, where it is attested in, respectively, Spring and Summer 1566 (Russow 158460r; Stefan Hartmann 2008 no. 3463 (p. 79–80)). In Pskov, the plague arrived in Autumn 1566 (Nasonov 1955, 269). It is unclear, at this point, if the 1562 outbreak in South Germany was a continuation of the 'Ottoman' wave, or the beginning of a new wave radiating out of the putative South German reservoir in the same year. Again, any such identification is hindered further by our lack of knowledge about the geographic whereabouts of the reservoir associated with the 1A3 branch, as well as additional genomes associated with the branch in question.

The origins of the 1570–1571 outbreak in Tallinn/Reval seem to be more straightforward to track. Contrary to the 1566 outbreak, there is no evidence of plague activity in any western or southern regions in the preceding years (1568–1570). As the chronicler Balthasar von Rüssow (c.1536–1600), himself a native of Tallinn/Reval, stated, the outbreak first commenced in early November 1570, and after abating in Spring 1571, broke out again in late July of the same year. According to von Rüssow, the locals called the disease the 'Russian or Muscovite Plague' ('Rüssche edder Muscoswitissche Plage'; Russow 1584, 75r and 77v). Indeed, the plague appears to have been imported by Ivan IV the Terrible's armies during the Siege of Tallinn/Reval (21 August 1570 – 16 March 1571). According to contemporary eye-witnessing sources, plague in and around Moscow began at some point in 1568, intensifying and then abating in 1569, and breaking out again in mid-July 1570 lasting until early December of the same year (Zimin 1950, 21; Sincera 1810, 332; Penskoi et al. 2021).

The 1568–1570 outbreak appears to have been associated with the same wave that spread into Estonia in 1566 (see the previous paragraph). The same wave appears to have spread into Russian territories from Lithuania, where it is attested in 1564–1565 (*Kronika Polska, Litewska, Zmodzka I Wszystkiej Rusi Macieja Stryjkowskiego* 1846 Vol. 2, p. 414; Stefan Hartmann 2008 Vol. 7, nos. 3400 (p. 46), 3396 (p. 45)), via Jezyaryshcha/Ozerishcha and Polotsk, where it ravaged in late 1565 – early 1566 (Nasonov 1955, 248). In Summer 1566, it arrived in Velikie Luki, Toropets and Smolensk; in the latter, it was present until March 1567, allegedly killing more than half of its inhabitants ("Dopolneniya K Nikonovskoi Letopisi" 1904-1906, 404, *Ustyuzhskiye I Vologodskiye Letopisi XVI-XVIII vv* 1982, 37:173). By Autumn 1566, the plague reached Pskov, Staraya Russa and Novgorod, remaining in the latter until early May 1567 (Nasonov 1955, 249; Koretskii 1980, 236). Concurrently, it was present in Mozhaisk ("Dopolneniya K Nikonovskoi Letopisi" 1904-1906, 404), and it was only a matter of time before it finally reached Moscow and its region at some point in 1568.

If the 1566 and 1570–1571 outbreaks are indeed associated with the same wave, then they represent an instance when the same wave can be introduced to a single location on more than one occasion. Indeed, Tallinn/Reval is not the only example of a possible reintroduction during the same wave. In late 1571, the wave returned to Novgorod, after its initial visitation in 1566–1567. Importantly, from Tallinn/Reval, the plague carried on both southwards and northwards. In the course of 1571, it spread all over Livonian territories (Russow 158477v) and then into Lithuania in 1572–1573 (*Kronika Polska, Litewska, Zmodzka I Wszystkiej Rusi*

*Macieja Stryjkowskiego* 1846 Vol. 2, p. 420). These regions had been previously attacked by what appears to be the same wave in 1564–1565. In the north, it was attested in southern Finland in 1571 (after its previous visit in 1566) (Aalto 2017) and eastern Sweden in 1572 (after ravaging that region in 1565; Ilmoni 1846-9 Vol. 2 p. 108, 144; Schröder 18549).

The genome of NMS002 (Augustinian Friary, Cambridge, United Kingdom) was published by Spyrou et al. 2019 with a dating interval of 1475–1536. The reported radiocarbon interval in the SI is 1437–1619, while the authors explain the shorter interval in the main text by considering marine reservoir effect, HBCO, and the *terminus ante quem* of 1538, when the chapter house, where the individual was buried, went out of use. Recalibration of the radiocarbon date as part of this study gave a 2-sigma interval of 1438–1621, after applying the sequence model with offset, the interval was shifted to 1448–1634. The probability distribution is bimodal (Fig. 63), dividing the 2-sigma range into two: 1448–1529 (53.6%) and 1551–1634 (41.8%). Accepting the *terminus ante quem* of 1538, only the first interval 1448–1529 is applicable.

A more precise dating of NMS002 is much more challenging. The main difficulty arises from the fact that NMS002 is positioned on the 1A3 branch, represented, for now, by only three genomes, all without precise dating. Here, we accept the interval of 1448–1529 as a putative date for the outbreak associated with NMS002. As the University of Cambridge records indicate, there were frequent plague outbreaks at local colleges in the period between c.1490 and 1538 (for the outbreaks predating 1465 in Cambridge and elsewhere in East Anglia, please refer to the previous section discussing the genomes of Clopton). For the post-1465 period, we hear of outbreaks in East Anglia (inferred from spikes in annual numbers of probated wills in Norwich and Bury St Edmunds, both in a vicinity of Cambridge) in 1471 (Norwich), 1473 (Bury St Edmunds), 1479 (both cities), 1492–1493 (Norwich), 1494 (Bury St Edmunds), 1500 (Norwich), 1503–1504 (Bury St Edmunds), 1504–1505 (Norwich) (Redstone 1907; Farrow 1943-1945). In addition, in 1492–1493 and 1495–1496 there were spikes in the annual number of last testaments probated at the Consistory Court of Bishop of Ely (Thurley et al. 1994–1996). In Cambridge proper, annual account books from King's Hall record plague outbreaks in 1498/1499, December 1500 – July 1501, 1502/1503, 1505/1506, 1508/1509, Summer – Autumn 1514, 1517/1518, 1520/1521, Spring 1525 or 1526, 1528–1529, 1532, and 1537–1538 (University of Cambridge, Trinity College Archives O.13, Vol. XIX, fols. 143v, 177r, 208r; Vol. XX, fols. 59v, 149r, 212v; Vol. XXI, fol. 68v; Vol. XXII, fols. 37v, 133v; Vol. XXIII, fol. 168v) (Cooper 1842; Cobban 1969, 220–222; Shrewsbury 1971, 163, 167; Leader 1988, 211–215). Given the multitude of references to plague, identifying each other's origins and possible association with NMS002 may not be feasible within the remit of the present project. It may suffice to conclude that it can be associated with one of the early sixteenth-century outbreaks. Also, given that it has only one unique SNP, the Cambridge outbreak appears to have commenced a few years after the beginning of its associated plague wave.

The genome of G488 (Riga, Latvia) was published by Susat et al. 2020, giving a date range of 1550–1650 in Susat et al. 2020 Fig. 2. While this study does not present a radiocarbon date, it can be found in the cited paper of Petersone-Gordina et al. 2020. The 2-sigma ranges given in the supplementary material are 1450–1530 (43.7%) and 1540–1640 (51.7%). The restriction to the latter interval is not justified in either publication, but probably relates to the assumption that the individuals of "Mass Grave 1" died during a famine in

1601–1602. Moreover, Susat et al. 2020 suggest that in addition to a concurrent plague outbreak in 1601–1602, the genome could also be associated with the outbreaks in 1621–1623 and 1657 in Riga (Susat et al. 2020). Unfortunately, G488 was not considered for the dietary isotope measurements presented in Petersone-Gordina et al. 2020, therefore a possible marine reservoir effect cannot be calculated.

Integrating the date of G488 into the sequence model shifted the 2-sigma interval from 1459–1635 (recalibration without offset) to 1479–1644 (PhIRM), compatible with the 1601–1602 (or, to be more accurate, the 1602) and 1621–1623 (or, to be more accurate, the August to December 1623) plague outbreaks in Riga (Fig. 63), with which the mass graves have been repeatedly associated (Susat et al. 2020; Petersone-Gordina et al. 2020; Petersone-Gordina et al. 2018). Nevertheless, in the present discussion, we have also considered earlier outbreaks from 1550 onwards, considering the terminal branch length of G488.
Between c.1550 and c.1650, we hear of several plague outbreaks in Riga, occurring in 1549/50–1552, 1565–1566, 1578–1579, 1602 and 1623. Because of the existence of at least two separate plague reservoirs (the one in the Ottoman Empire responsible for the 1A1 branch and the other one most likely in Central Europe causing 1A2 branch-associated waves – in addition to a possible reservoir responsible for 1A3 strains), the frequency and geographic pervasiveness of plague outbreaks in the 16th and early 17th centuries was much more intense compared to those in the 14th and 15th centuries, with possibly two or more strains radiating of different reservoirs, circulating concurrently or nearly concurrently across the same regions. This issue, compounded by the lack of precisely dated genomes on the 1A3 branch makes the reconstruction of each outbreak's origins hardly feasible at the moment. However, a tentative exercise may be attempted.

The 1549/50–1552 outbreak seems to have originated in South Germany in late 1546 or early 1547 (Erdel, Specker, and Winckelmann 1986; Sturm 2014, 37; Horanin 2019, 166–169), moving fast towards the Baltic Coast, arriving in Hamburg in the same year, then in Lübeck in either late 1547 or 1548 (Ibs 1994, 128–129), and finally in Gdańsk/Danzig, where it is attested in 1548–1549 (Kizik 2012, 70). It is most likely from Gdańsk/Danzig that the plague was imported into Riga in either 1549 or 1550, ravaging the city until Autumn 1552 (Russow 158427v), before moving southwards into Lithuania (*Kronika Polska, Litewska, Zmodzka I Wszystkiej Rusi Macieja Stryjkowskiego* 1846 Vol. 2, p. 754) and eastwards into Pskov and thereafter into Novgorod in the same year (Tikhomirov 1965, 180; Stefan Hartmann 2005 no. 159). We do not know, at this stage, if the South German reservoir was responsible only for the 1A2 branch, or if it is possible that the 1A3 branch, emerging before the "Little Polytomy" (N13, discussed in Supplementary Section 11.7), was being radiated out of the same reservoir, or if it had been seeded elsewhere.

By contrast, the spatio-temporal origins of the 1565–1566 outbreak are more difficult to establish, as they may potentially be associated either with a wave commencing in or around the Balkans in 1559 or with that starting in South Germany in 1562. Alternatively, the 1562 outbreak in South Germany could potentially be a continuation of the 'Ottoman' wave, rather than the beginning of a new wave radiating out of the putative South German reservoir. The details of that/those wave/s are discussed in Supplementary Section 11.6. Again, any such identification is hindered further by our lack of knowledge about the geographic whereabouts of the reservoir associated with the 1A3 branch.

The spatio-temporal origins of the 1578–1579 outbreak may be associated with two possible scenarios. It may have been associated with a wave commencing in South Germany in 1570–1571, spreading all over Germany in 1572–1577 (Eckert 1996, 87–93), crossing fast into Denmark in 1575 (Christensen 2003, 418) and then Sweden either in the same or following year (Ilmoni 1846-9 Vol. 2, p. 98–99). Concurrently, it was spreading eastwards along the southern Baltic coast, reaching Szczecin/Stettin in 1577 and Tapiau (today's Gvardeysk) in 1578, but it was not until 1580 that it reached neighboring Gdańsk/Danzig and Königsberg (Kizik 2012, 71; Sahm 1905, 16–17). It was, most likely, from one of these Baltic ports that the plague reached Riga in Autumn 1578, remaining in the city until after summer 1579 (Russow 1584, 117r). The second scenario is that the 1578–1579 outbreak may be associated with a different albeit concurrent plague wave circulating in Southern and Central Europe in the 1570s. Originating in the Ottoman territories – possibly in or around the Aegean region in 1570 – the wave spread northwards all over the Balkans and Wallachia in 1571–1574 (Varlık 2015, 189–196; Binder 1983, 100–102), reaching Hungary in 1576 (Magyary-Kossa 1931, 230). To reach the Baltic coast in the following year, the same wave would need to have traveled via Carpathians into Poland. There is, however, no evidence of plague outbreaks in Southern and Central Poland in 1576–1577 (Walawander 1932, 285).

Establishing the origins of the 1602 outbreak is much more straightforward. It may be associated with a wave beginning in South Germany in 1592, and spreading all over the country in 1593–1598 (Eckert 1996, 100–112). Although reaching Lübeck in 1598, its maritime spread in the North and Baltic seas was slow, owing to quarantine measures implemented by local authorities. It was not until Autumn 1601 that it arrived in southern Denmark, Gdańsk/Danzig and Königsberg (Christensen 2003, 418; Kizik 2012, 72; Sahm 1905, 20–21), while in southern Sweden the plague was not attested until the following year (Ilmoni 1846-9 Vol. 2, p. 159). The plague may have been imported into Riga in 1602 from either the south-eastern Baltic or from Sweden (Gerhards 2011, 47–48, 54), and its spread may have been facilitated by Polish-Lithuanian troops marauding in the city, after the Swedish retreat from there in August-September 1601.

By contrast, establishing the spatio-temporal contours of the 1623 outbreak, ravaging the city between August and December (Gerhards 2011, 48), is much more difficult. It may have arrived in the city from Estonia, where it is attested in late 1622 – June 1623 (Tallinna linnaarhiiv, *Begravna i St.-Olai kyrkan i Reval, 1603–1821*: 41–42), possibly imported from Sweden, where it had been attested earlier that year. But it could have also been imported from the south – from eastern Poland and Lithuania, where it had been reported in 1622–1623 (Karpiński 2000, 314). The origins of the 1623 outbreak are even less clear. If it had been imported from Sweden, by the way of Estonia, then it was clearly associated with one of the waves of the Thirty Years' War plague. As the phylogenetic evidence indicates, four genomes from later Thirty Years' War plague outbreaks (c.1628–1632) were caused by strains of Branch 1A2, rather than 1A3. This does not preclude the concurrent circulation of strains from both branches. If, however, the 1623 outbreak was imported from the south, then the origins of its associated wave appear to have commenced in the Ottoman territories. It appears to have been imported into South Poland from Hungary, where it was attested in 1621–1622 (Magyary-Kossa 1931, 331–332), where, in turn, it seems to have been imported from the Balkans, ravaged by the plague in the preceding years (Bazala 1962, 73; Nikolov and Michev 1978; Hrabak 1989).

The South German origin of at least three out of five outbreaks suggests that if the identification of NMS002 with one of the proposed outbreaks is correct, then it is possible that the reservoir associated with 1A3 branch may have emerged in South-Central Germany, in or around the region, in which Branch 1 had previously seeded a reservoir between c.1349 and 1356, out of which both the *pestis secunda* (commencing in 1356) and (after a putative southwestern migration) the subsequent waves caused by Branch 1A (commencing with the *pestis tertia* in 1363) radiated.

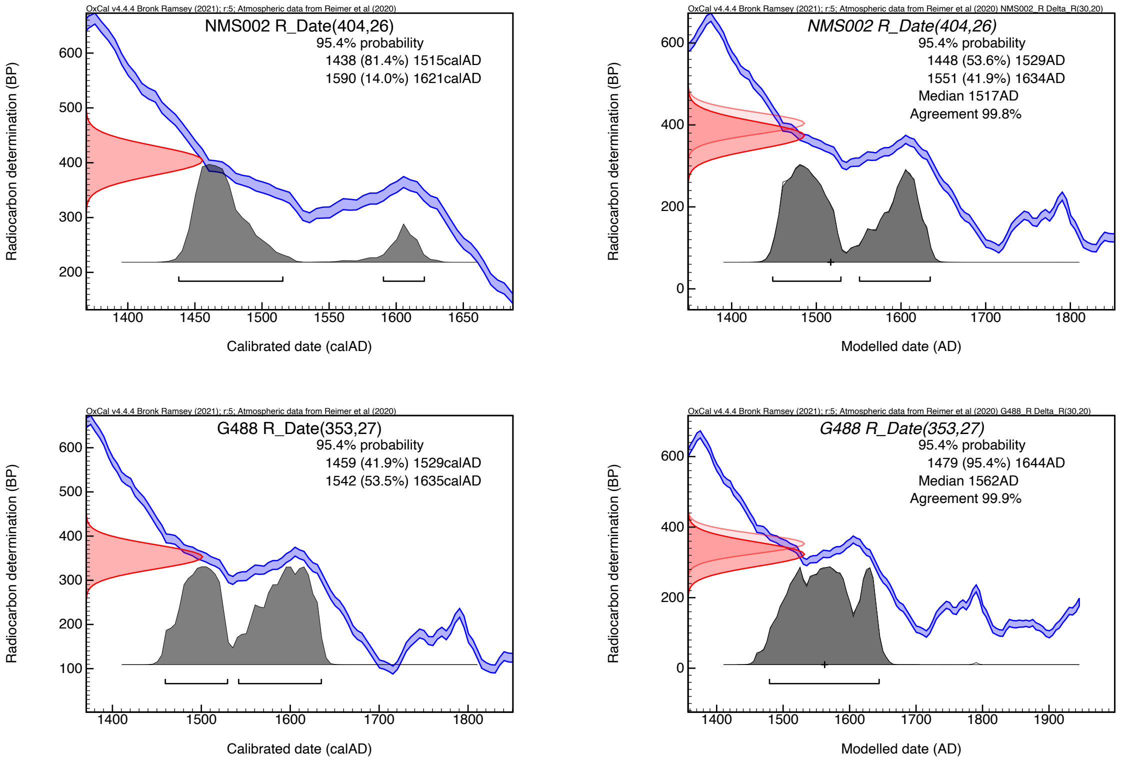

**Fig. S63: probability distributions (brackets indicating 2-sigma intervals) of NMS002 and G488 without offset (left) and after integrating into the sequence model with a generic offset of 30 ± 20 RC years (right).**

#### 11.7 The “Little Polytomy” (N13), ELW098.A, AGU025.B

The polytomy N13 gave rise to two major sub-branches of Branch 1A causing major plague outbreaks in Europe at least until the 1630s (Branch 1A2) and the 18th century (Branch 1A1), but also two lineages so far only represented by single genomes on short terminal branches (ELW098.A, Ellwangen, Germany; AGU025.B, Vilnius, Lithuania). This is a remarkable diversification event, especially when also considering Branch 1A3, emerging just one SNP prior to the polytomy.

Unfortunately, no genome falling on node N13 has been sequenced so far, implying that we have to approach the dating of this diversification event indirectly. PhIRM offers the possibility to use radiocarbon dates of all genomes post-dating the polytomy to determine an upper boundary. Since both the genome of ELW098.A and the basal 1A1 cluster are only one SNP derived from the polytomy, this upper boundary is expected to come close to the date of the polytomy itself. The boundary was implemented in the sequence model as Boundary T6 with a 2-sigma range of 1418–1473 (median at 1446, Fig. 64), thus placing the polytomy roughly in that period.

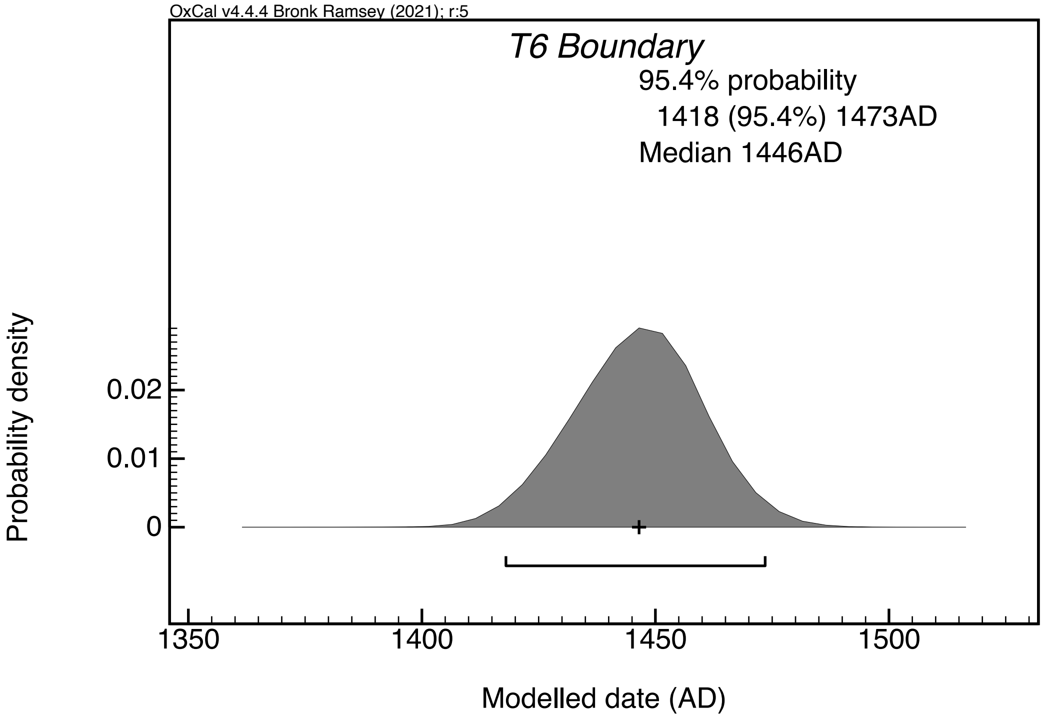

**Fig. S64: probability distribution (brackets indicating 2-sigma intervals) of Boundary T6 as part of the sequence model with a generic offset of 30 ± 20 RC years.**

The sequence model does not affect the two dates of the genomes on terminal branches emerging from node N13: ELW098 has calibrated 2-sigma ranges of 1466–1635 without offset and 1485–1644 with a generic offset of 30 ± 20 RC years, and they remain basically unchanged within the model (1466–1635 and 1485–1645, respectively). The same is true for AGU025 with 2-sigma ranges outside of the model of 1442–1615 (without offset) and 1450–1631 (with a generic offset of 30 ± 20 RC years), within the sequence model of 1442–1614 (without offset) and 1452–1631 (with a generic offset of 30 ± 20 RC years). Considering the length of the terminal branches of 1-3 SNPs and the bimodal distributions of the radiocarbon ranges of both ELW098 (without offset) and AGU025 (both with and without offset), a date in the late 15th or the early 16th century seems more likely - that is up to half a century later than the suggested dating of the Little Polytomy (N13), in line with their phylogenetic distance.

Accepting the modelled intervals of c.1377–1418 for AHM001/AHM011/N06, 1406–1457 for Boundary T5 as an approximation for N10, 1418–1473 for Boundary T6 as an approximation for N13, and 1440–1577 for the basal Branch 1A1 cluster/N14, we can tentatively narrow down the respective intervals when considering their phylogenetic distance. The nodes are not equidistantly distributed: N05–N10 are 12 SNPs apart, while N10–N13 and N13–N14 are only 1 SNP apart, respectively. Assuming a relatively constant substitution rate along this internal branch, we would expect much shorter intervals between N10, N13, and N14 than N05 and N10, which would push them towards the mid/end of the 15th century.

One potential difficulty with this dating is the fact that one of the post-Little Polytomy genomes, AGU007 (Vilnius, Aguonu St. graveyard), positioned on N14, one 1 SNP after N13, shows the presence of *Treponema pallidum* (Giffin et al. 2020). Despite some recent advances in aDNA analysis of *Treponema pallidum*, the bacterium's presence in Europe before Columbus' voyages has not yet been decisively established, although it cannot be precluded either (Majander et al. 2020). Also with the new data and PhIRM, this question cannot be answered: integrating all dates of the basal Branch 1A1 genomes into a KDE_Plot within the sequence model, all individual dates cover the year 1498 (Gdansk8 1440–1510; AGU010 1445–1520; RUS003 1455–1672; RUS004 1456–1575, and AGU007 1473–1577). The first textual reference to an outbreak of syphilis in Vilnius comes from 1498 (*Kronika Polska, Litewska, Zmodzka I Wszystkiej Rusi Macieja Stryjkowskiego* 1846 Vol. 2, p. 304). Therefore, neither a pre- nor post-Columbian appearance of treponematosis in Europe can be excluded.

Taking all these facts and uncertainties, it is safest to widen the potential chronological bracketing of the Little Polytomy to c.1450 – c.1500. At present, without securely dated genomes falling either on the Little Polytomy itself or nearby nodes, or positioned on nearby branches, no narrower range can be suggested with any certainty.

The Little Polytomy (N13; to be distinguished from the "Great Polytomy" emerging in the early 14th century, most likely c.1340, and giving birth to four long-lived internal branches) was a major evolutionary event, whereby at least four branches emerged concurrently. Two of these branches are currently only represented by single genomes with short genetic distance to N13 (ELW098 with 1 SNP; AGU025 with 3 SNPs), while the other two gave rise to bigger clades: Branch 1A1 currently represented by 19 genomes from 11 different contexts and Branch 1A2 currently represented by 14 genomes from 6 different contexts). Polytomies are still poorly understood phenomena, and it is unclear if these diversification events are coincidental, or if they are triggered by exogenous (natural) factors, biotic and abiotic. It has been suggested that the exposure of *Yersinia pestis* to ecological stress may result in its evolutionary changes - a process known as stress-induced mutagenesis (SIM; Suntsov 2019, 2021).

The proposed dating of the Little Polytomy between c.1450 and c.1500 coincides with one of the most pronounced climatic anomalies in Central European history of the last 2,500 years - the so-called 'Great Renaissance Drought'. This event of almost megadrought proportions consisted of 17 straight years of excessively dry springs and summers between 1457–1473, preceded by the drought spells of 1437–1448 and 1450–1453 and followed by those of 1476–1479, 1483, 1491–1494, 1498 and 1502–1504 (Cook et al. 2015; Büntgen et al. 2021). To make things even worse, the period between 1451 and 1470 stands out as one of the coldest ones in Central Europe between 500 BCE and 2000 CE, with annual summer temperatures dropping -1.5°C below their average over the entire period (Büntgen et al. 2011). Conversely, the period of 1471–1474 saw four back-to-back excessively hot summers, with some far-reaching impact on the ecosystem (Camenisch et al. 2020). A wide array of sources from all over Europe, from Italy to Poland, narrate about extensive wildfires (Camenisch 2015, 295; Camenisch et al. 2020; Kiss 2017; Malewicz 1980, 137–138), with one Nuremberg source noting the mortality of deer and other sylvatic animals (Glaser 2001, 70). One Polish chronicler reported mass mortality of herbivores ingesting gravel, because of grass scarcity, in the same year (Malewicz 1980, 137). Some German and Swiss sources revealed that very little hay could be made because of the drought (Dierauer 1900, 257). A Göttingen chronicle stated that the fruit harvest of 1471 was destroyed by worms (Glaser 2001, 70), and there are several sources indicating the death of trees in the same year. In addition, all of Central and Eastern Europe was devastated by locust invasion, coming from south-eastern Europe, between 1473 and 1480 (Brázdil et al. 2013). The extreme drought is noted by several chronicles from the Nuremberg-Augsburg region (*Die Chroniken Der Fränkischen Städte. Nürnberg. Vierter Band* 1872, 336; Müllner 2003, 19–20; *Die Chroniken Der Schwäbischen Städte. Augsburg. Dritter Band* 1892, 236; Glaser 2001, 70), presumably in the vicinity of the putative South German reservoir.

The adverse combination of excessive drought with either excessive cold or heat can severely reduce the available biomass, by depressing grass growth, both in meadows and woodland, and tree mast, by destroying woodland vegetation through either fire or drought. This would lead to mass die-offs of sylvatic animals or/and their migration to arable fields and hence, closer to human habitat. It could be in this context that the Little Polytomy may have occurred, with *Yersinia pestis* experiencing stress-induced mutagenesis (SIM) and spilling over from infected sylvatic rodents – possibly common voles (*Microtus arvalis*) – to humans, leading to an outright outbreak in the latter, which unfolded into a new plague wave, reflected in the ELW098 and AGU025 genomes.

Given that ELW098 is only one SNP away from the Little Polytomy (N13), we may assume that (1) the Little Polytomy might have emerged not far from Ellwangen, and potentially in the same South German reservoir radiating previous 1A branch-associated outbreaks; (2) it could be a wave commencing in South Germany in 1462, 1472, 1482, or 1492, given the suggested dating of the Little Polytomy to c.1450 – c.1500; (3) the Ellwangen outbreak itself occurred shortly after the Little Polytomy. While there are no direct references to plague outbreaks in the town, we have indirect evidence from nearby places. Thus, in 1462 there was a plague in Schorndorf (~65 km south-west of Ellwangen; *Beschreibung Des Oberamts Schorndorf* 1851, 108), while in the following year was reported in Nördlingen (~35 km south-east of Ellwangen) (Sturm 2014, 37) and Heilbronn (100 km north-west of Ellwangen; Steinhilber 1956, 198). For the next wave, we are told of a 1472–1473 outbreak in Rothenburg ob der Tauber (~60 km north of Ellwangen; Schnurrer 1987), Esslingen (~100 km south-west of Ellwangen) and a 1473–1474 outbreak in Nördlingen (Sturm 2014, 37). For the 1482 wave, we know about a 1482 outbreak in Heilbronn and Esslingen, and a 1483 outbreak in Nördlingen, Schwäbisch Hall and Rothenburg ob der Tauber (Steinhilber 1956, 198; Sturm 2014, 37; Schnurrer 1987). Finally, for the 1492 wave, we should consider a 1492–1493 wave in Heilbronn (von Rauch 1913, 266; Steinhilber 1956, 198), and the 1493 one in Stuttgart (~95 km south-west of Ellwangen; Pfaff 1845, 240) and Esslingen and the 1494 one in Nördlingen, Schwäbisch Hall and Rothenburg ob der Tauber (Sturm 2014, 37; Schnurrer 1987).

The phylogenetic position of AGU025 on a terminal branch, born out of the Little Polytomy, implies that its associated outbreak occurred shortly after the Little Polytomy, just as the Ellwangen one. However, the fact that AGU025 is 3 SNPs derived from N13 – in contrast with ELW098, one SNP derived – may indicate that chronologically, AGU025 came later than ELW098. Given that the AGU025 and ELW098 are situated on different terminal branches, it is unclear if the two genomes are associated with the same outbreak or with two different ones. Also, given that AGU025 is positioned on a different branch from other genomes from the same graveyard (AGU007 and AGU010, both positioned basally on N14; see Supplementary Section 11.12), is unclear if AGU025 is to be associated with the same outbreak as AGU007/010 (Giffin et al. 2020). Importantly, AGU025 comes from a different grave (Grave 8) than AGU007 (Grave 104) and AGU010 (Grave 94; Giffin et al. 2020 Supplementary Table 1), indicating that both scenarios are possible. As in Supplementary Section 11.12, the AGU007/010 genomes are dated, together with other genomes positioned together basally on N14, to the last third of the 15th century – the first decade of the 16th century. In particular, it is suggested that AGU007/010 could be associated with the 1474, 1495, and 1505–1506 outbreaks reported in Vilnius and elsewhere in Lithuania, in addition to a possible outbreak of c.1482, whose textual evidence could not, at this point, be found, but which is reported in both Riga and Poland. If AGU025 and AGU007/010 come from the same outbreak, then AGU025 could be associated with any of the proposed outbreaks. The origins of the 1474, 1495, and 1505–1506 outbreaks are discussed in Supplementary Section 11.12; the 1465 outbreak is associated with the same wave (commencing in South Germany in later 1462) as the 1464–1465 outbreak in Riga, discussed in Supplementary Section 11.5.

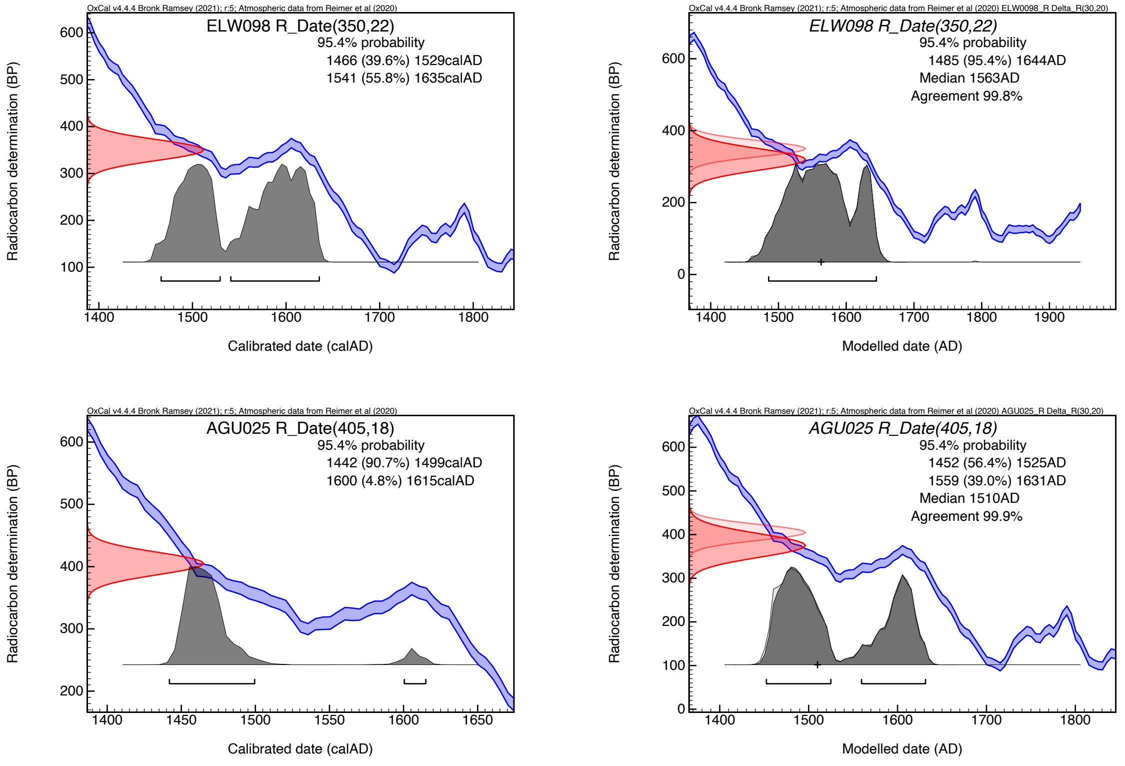

**Fig. S65: probability distributions (brackets indicating 2-sigma intervals) of ELW098 and AGU025 without offset (left) and after integrating into the sequence model with a generic offset of 30 ± 20 RC years (right).**

#### 11.8 Branch 1A2, LBG002.A and the STN genomes

The Branch 1A2, one of the major lineages born out of the polytomy N13 includes the genome of LBG002.A (Landsberg am Lech, Germany), which represents the direct ancestor of all other Branch 1A2 genomes of Stans, Switzerland and the plague of the Thirty Years' War. Therefore, the analysis of LBG002.A profits the most from the PhIRM approach. In the original publication (Spyrou et al. 2019), a date range of 1455–1632 is given in the main text, based on the 2-simga radiocarbon interval. However, the supplementary text states that the churchyard of St. John in Landsberg am Lech, where the plague victim was recovered from, had been in use from 1507–1806. The recalibrated 2-sigma range of LBG002 is 1456–1631; in addition, we have radiocarbon dates for two more individuals that tested positive for plague (Spyrou et al. 2019) and are probably identical: while LBG002 (arch. ID 460) was buried in a mass burial containing 11 individuals, LBG005 (arch. ID 572) coming from a single burial covered with quicklime was dated to 1456–1631, and LBG007 (arch. ID 598) deriving from a quintuple burial covered with quicklime was dated to 1458–1632 (2-sigma ranges). These broad intervals of more than 173–175 years get drastically reduced by applying PhIRM to 60–61 years: 1458–1519 (LBG005), 1459–1520 (LBG007), and 1460–1520 (LBG002).
In the original publication, local outbreaks of plague in 1586, 1592, 1627–1629, and 1634 were mentioned, none of which fall into the range of the refined dating. Accepting the *terminus post quem* of 1507 for the foundation of this cemetery, the interval could be narrowed down even further to 1507–1520, corresponding well with the fact that the multiple burials represent the lowest of eight burial levels (Spyrou et al. 2019).

Although direct references to plague outbreaks in Landsberg am Lech for that period could not be found for the suggested narrow range of 1507–1520, they survive for nearby Swabian and Upper Bavarian cities and towns. Thus, we hear about plague outbreaks in 1509 (Memmingen), 1511–1512 (Augsburg and Memmingen), 1515–1517 (Munich) and 1520–1521 (Augsburg, Kempten, Kaufbeuren, Memmingen, Munich and Ulm) (Müller 2006; Berning 1987-9; Erdel, Specker, and Winckelmann 1986; Horanin 2019, 125–151; Stahleder 2020, 213). The basal positioning of LBG002 on N22 hints that its associated wave may have commenced in a Central European reservoir – perhaps in South Germany – persevering in that region after the Little Polytomy and being responsible for LBG002 and subsequent 1A2 branch-associated outbreaks. However, if the Little Polytomy (N13) emerged later, c.1485–1495 (if we were to assume the presence of *Treponema pallidum* in AGU007 to be a result of Columbian Exchange), then the acquisition rates on the N13–N22 segment of 1A2 Branch were considerably faster than if N13 emerged closer to the mid-15th century – about one SNP every 4.2 or so years. This would be consistent with Spyrou et al.'s analysis indicating that there was an acceleration in speed of substitution rate between the two nodes (Spyrou et al. 2019).

The Stans cluster is only 1 SNP derived from node N23. For the dating of N23, we can use the Boundary T7 as an approximation. Boundary T7 is constrained on the lower boundary by the radiocarbon dates of Landsberg am Lech and on the upper boundary by the Stans cluster and the Thirty Years' War cluster. The respective 2-sigma interval is 1481–1536 (Fig. S66). However, accepting 1507 as *terminus post quem* for Landsberg am Lech, this interval can be constrained to 1507–1536. However, it needs to be noted that the suspiciously old radiocarbon intervals of Domat/Ems (discussed in Supplementary Section 11.9) might push this boundary towards an older date and that N23 is 5 SNPs derived from LBG002/N22. Therefore, we can tentatively date this node into the second quarter of the 16th century.

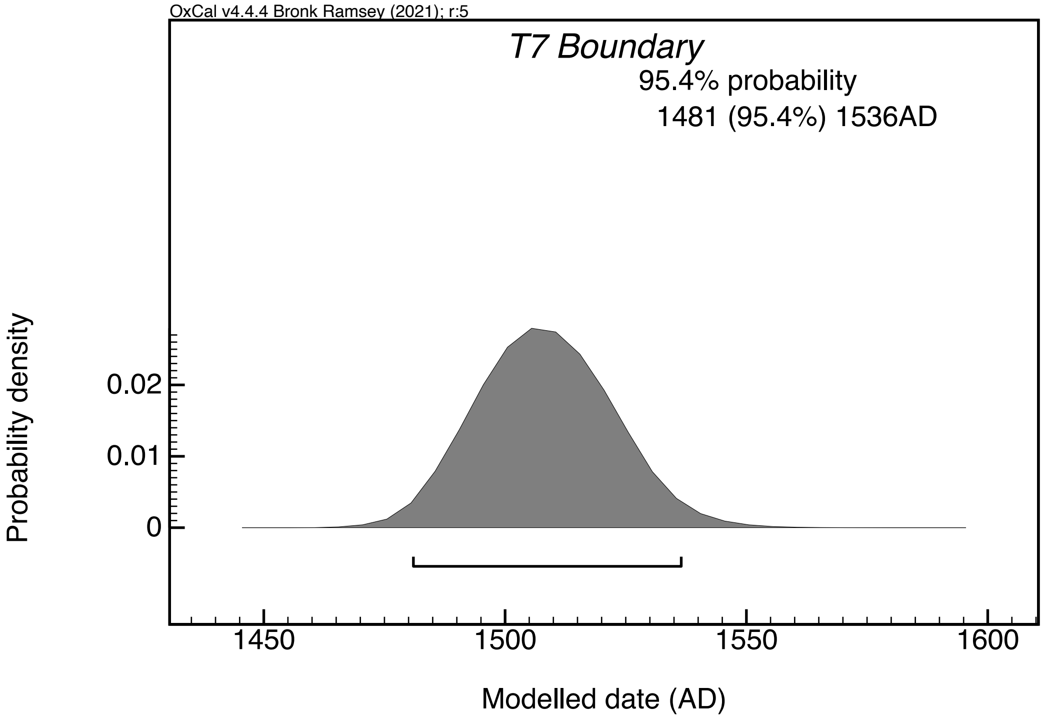

**Fig. S66: probability distribution (brackets indicating 2-sigma intervals) of Boundary T7 as part of the sequence model with a generic offset of 30 ± 20 RC years.**

The genomes of Stans (STN) belong to three different mass graves, based on the low diversity (maximum 1 additional SNP) we can assume that they are all contemporaneous. In the original publication (Spyrou et al. 2019) a date range of 1485–1635, based on a combination (R_Combine in Oxcal) of available radiocarbon dates, has been given. The respective raw and calibrated dates can be found in Tables S10 and S11, the 2-sigma ranges reach from 1480 to 1638 (Table S11). Through PhIRM, the 2-sigma ranges get slightly shortened and shifted towards younger dates, with an absolute lower boundary of 1504 and an absolute upper boundary of 1665. A comparison of the KDE_Models and KDE_Plots of all radiocarbon dates of Stans reveals that the probability distribution shows itself between two and three peaks (Fig. S67). For the plots/models including a generic offset, the first, broader peak falls into the mid-16th century and the second, narrower peak into the 1610s–1630s. Taking the genetic distance of the Stans genomes to the MRCA (N23) with the Thirty Years' War plague cluster into account (1–2 SNPs vs. min. 8 SNPs), the Stans outbreak seems more likely to date to the mid-16th century.

Surviving records from both Stans and neighboring towns record several plague outbreaks in that period. In 1548 there was an outbreak in Engelberg (20 km south of Stans); in 1564–1565 there was a major outbreak all over Switzerland, reported in Stans and nearby Lucerne (14 km north of Stans); finally, in 1573–1574 there was an outbreak in Engelberg, spreading in 1575 to Lucerne (Krämer 2014; Schnyder 1932, 107–108). The 1548, 1564–1565 and 1575 outbreaks appear to be associated with the waves commencing in the South German reservoir in, respectively, 1546–1547, 1563, 1572, discussed in the previous section dealing with the sixteenth-century outbreaks in Riga.

Six out of eight Stans genomes (STN002, STN007, STN008, STN014, STN019 and STN020) appear to be identical; two (STN013 and STN021) have acquired an additional SNP. This implies that the additional SNP was acquired either via local evolution during the outbreak in the town, or via a (re)introduction in a more derived form from outside. There are some instances, when *Yersinia pestis* genomes from a single spatio-temporal context show a genetic variation of one SNP between them - as was indeed the case of the Great Marseille Plague (1720–1722), with one genome (out of the total five) possessing an additional SNP in relation to its four counterparts.

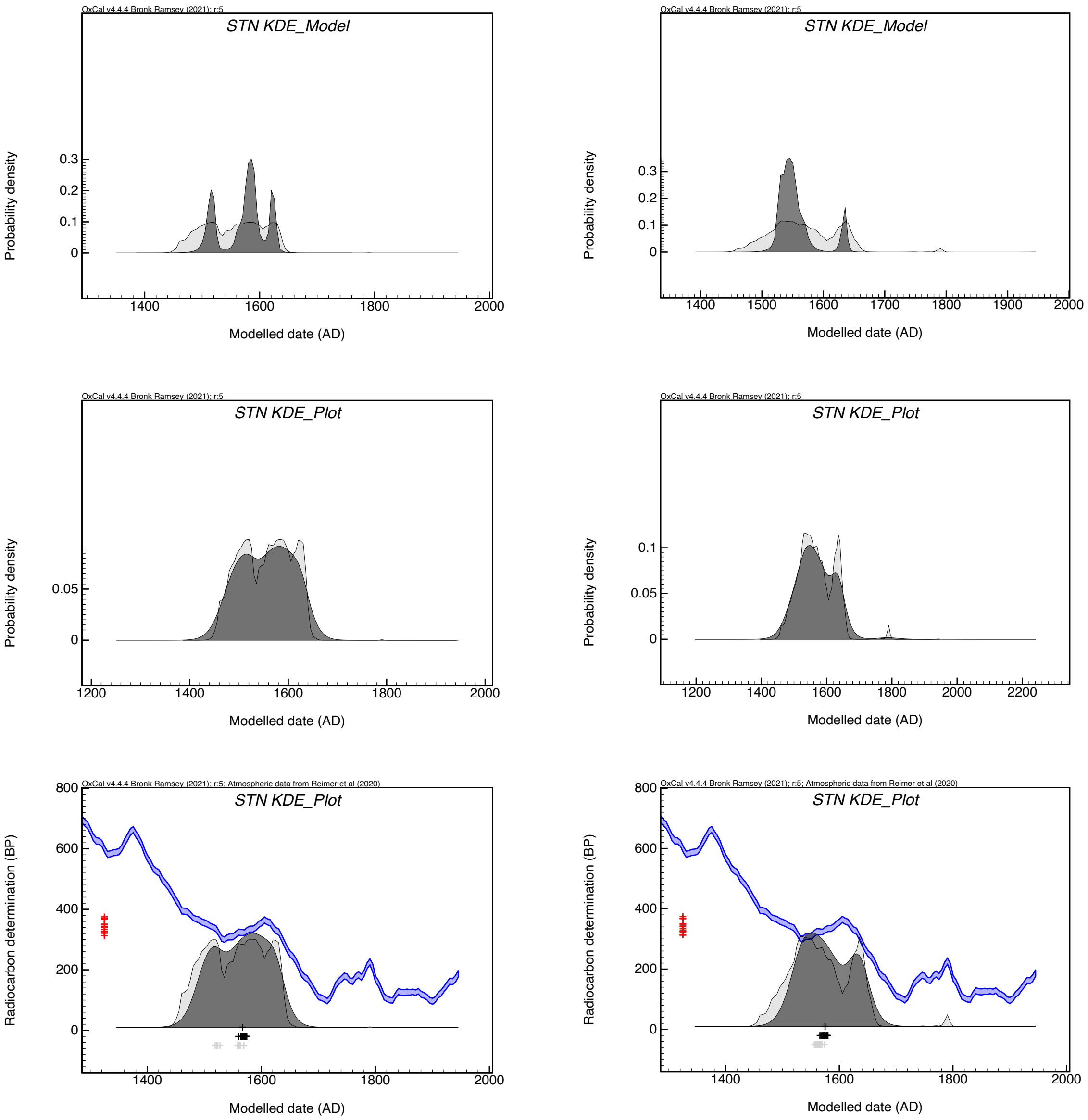

**Fig. S67: Probability distributions for KDE_Models (top row), separate KDE_Plots (middle row) and KDE_Plots within the sequence model (bottom row) without (left) and with (right) a generic offset of 30 ± 20 RC years.**

#### 11.9 Genomes of the Plague of the Thirty Years' War – SPN19, LAR11t, BRA001.A and the EMS genomes

The first published genome of this clade was BRA001.A (Brandenburg an der Havel). There is no radiocarbon date available for the burial this genome is derived from, but the contextual information provides a relatively precise date. According to a first PCR-based study (Seifert et al. 2016), the triple burial dates to the Thirty Years' War (1618–1648) based on a publication presenting the archaeological and anthropological findings (Dalitz, Grupe and Jungklaus 2012). The burial was found in the backyard of a bourgeois house and contained the remains of three young men with signs of healed traumatic injuries. Furthermore, a clay pipe with a stamp providing a *terminus post quem* of 1630–1640 was found with one of the

skeletons, and strontium and oxygen isotope analyses confirmed a non-local origin of the individuals. This led to the identification of the individuals as soldiers of the Swedish troops that were garrisoned in Brandenburg an der Havel in 1626–1631 during the Thirty Years' War. While the publication of the fully reconstructed genomes gives the latter time span in the main text (Spyrou et al. 2019), the supplementary text mentions local plague outbreaks in 1625–1627 and 1631. Taken together with the dating of the clay pipe, the burial is most likely associated with the 1631 outbreak.
A closely related strain from the site of San Procolo a Naturno (Naturns) in Northern Italy was published by Guellil et al. 2020, dated by the authors to ca. 1636 in their Fig. 3 (Guellil et al. 2020) following numismatic evidence, as described in the supplementary text. Another closely related genome was found at the site of Lariey, Puy-St-Pierre, Hautes-Alpes, France, around 390 km southwest of Naturno/Naturns (Seguin-Orlando et al. 2021). According to the supplementary text (Seguin-Orlando et al. 2021), radiocarbon dating, "rare ceramic artefacts" and archival information allows to associate the respective burials unambiguously to the years 1629–1630 of the local plague epidemic of 1628–1632.

The site of Domat/Ems, presented in this study, represents a small cemetery at the Church of Sogn Pieder/St. Peter, used for a short period of time, as inferred from the fact that none of the single and multiple burials intersect or superpose. The high number of young adult male individuals, the presence of different tools for weapons in the burials and pilgrimage coins giving a *terminus post quem* of 1610 suggest that the burials date to the "Bündner Wirren" (1618–1639), a local conflict in the Canton of Graubünden, forming a part of the Thirty Years' War (see Supplementary Section 2.10). Since soldiers were garrisoned in Domat/Ems in 1629–1631 and the only plague outbreak in the Graubünden region around that time occurred in 1628–1632, we may argue that two distinct plague genomes recovered from this site, EMS001 and EMS010, most likely date to the years 1628–1632 (or, to be more precise, to 1629–1630, as indicated below). In total, four individuals of this cemetery were radiocarbon dated: the skeletons of burial 17 and 99A were dated at the radiocarbon laboratory of the ETH Zurich, while the skeletons of burial 24/EMS001 and 98B/EMS010 were dated at the radiocarbon laboratory in Belfast (see Table S10). Whereas the 2-sigma intervals of 17 and 99A overlap with the expected date range (both 1458–1632 after calibration), the interval of 24/EMS001 does so slightly (1433–1620), and the interval of 98B/EMS010 is significantly younger (1431–1490). The latter dates even older than the Landsberg am Lech plague genome (LBG002, see Supplementary Section 11.8), although the genome EMS010 is 13 SNPs derived from LBG002.A (Fig. 2A). This discrepancy within the site and with the phylogenetic information could be caused by several effects or a combination thereof: an offset due to freshwater or marine reservoir effect, a human bone collagen offset, a statistical outlier or a perhaps measurement error.

Since we can assume a relatively short usage period for the cemetery, we integrated all four radiocarbon dates into our sequence model within a KDE_Plot. Interestingly, even the sequence model without generic offset shifted the 2-sigma intervals younger, giving low agreement values for the two Belfast dates (24/EMS001: 1466–1627, A=38.7; 98B/EMS010: 1465–1620, A=10.0, see Code S5, Table S11, Fig. 46 right). Adding a generic offset of 30 ± 20 RC years, all the intervals are overlapping with the expected range of 1628–1632 (Fig. 3, Code S3, Table S11, Fig. S46 left), but the date of 98B/EMS010 still receives a low agreement value (A=55.1) and a significant shift towards a higher offset (see Fig. 68). Although the archaeological and historical information available for this site was not used as prior in this model, all four radiocarbon dates were modelled to conform to the presumed dating around 1631, while flagging one of the dates as problematic. This, again, highlights the methodological advantage of PhIRM.

As far as the phylogenetic positions of these genomes are concerned, assuming all dating to a few years between 1629 and 1636, and the geographic proximity between Naturno/Naturns, Lariey and Domat/Ems, the diversity is remarkable. This is especially striking for the site of Domat/Ems where two distinct genomes were found, both deriving from a common ancestor with BRA001, with a terminal branch of one private SNP, respectively (Fig. 2A). None of the low coverage genome EMS002 and EMS003 reconstructed from this site seem to share either of these private SNPs, meaning that this direct ancestor, representing the node N27, might have been present at the site as well. Surprisingly, the genetically closest genome was found in Brandenburg an der Havel, 810 km north of Domat/Ems, while the sites of Naturno/Naturns (180 km) and Lariey (430 km) are much closer to Domat/Ems. We may therefore conclude that the plague epidemics ravaging through Central Europe in the 1620s–1630s show a more complex phylogeography and chronology than previously assumed.

In order to understand this phylogenetic complexity, it is important to historically contextualize each of these genomes, and the routes by which they arrived at sites associated with their specimens' burials. Although both the BRA/EMS and the LAR/SPN clusters are associated with the same source (N25), perhaps initiating the same wave, their spatiotemporal trajectories were different. One distinctive feature of the Thirty Years' War plague is that outbreaks were both more geographically widespread and pervasive, and longer than those during preceding waves (Eckert 1996, 132–154). Some cities and regions were ravaged by plague for several years in a row, and it is unclear if such lengthy outbreaks were caused by repetitive reintroductions by warring armies, or by the inability of local authorities to cope with the crisis caused by poor hygiene of locally stationed garrisons. The constant movement of different armies spreading plague bacteria into the places they were invading or stationed at makes it very difficult, at this stage, to establish a clear association between particular outbreaks and their wider waves, and it is unclear how many waves radiated out of their putative South German reservoir in the 1620s and 1630s. Hence, determining whether the two clusters are associated with the same wave or with two different ones, as well as establishing the chronological dates of the associated wave(s), cannot be undertaken in the present study.

Conversely, the transmission routes of the strains associated with the two clusters may be tentatively reconstructed. The BRA001 genome has been associated by the original authors with a 1631 burial of a Swedish soldier stationed at Brandenburg an der Havel (Spyrou et al. 2019). The EMS samples come, as indicated in Supplementary Section 2.10, from soldier burials at Sogn Pieder /St. Peter's Church, putatively associated with the presence of Imperial Austrian troops (the Alwig Count von Sulz Regiment) stationed at Ems, between about June 1629 and April 1631 (although we do not know the exact arrival date of the troops in the town, we may still assume it happened shortly after the occupation of Chur on 28 May 1629). It is important to note that the plague may have already been present in Domat/Ems before the arrival of the Austrian troops and hence, its initial introduction to the region should not be strictly associated with the Austrian invasion. Although, as noted above, records contemporaneous with the burials have been destroyed by a town fire in 1776, the chronology of plague spread in the region immediately around Domat/Ems can easily be reconstructed from records of neighbouring places. Thus, we know that plague began in Chur (only 7.2 km north-east of Ems) in November 1628, continuing into 1629, in the course of which it lost about 1,300 citizens. In Summer 1629, the plague spread into other settlements in the region, including Domleschg (13 km south of Domat/Ems), Thusis (20 km south of Domat/Ems), Ronggellen (23 km south of Domat/Ems) and Safiental (20 km west of Domat/Ems) (Lorenz 1868–1869, 36–37; Eckert 1996, 67–68). In Thusis, the first outbreak had been reported on 17 August 1629 (Lorenz 1868–1869, 36). Given that Domat/Ems lies about halfway between Chur and Thusis, along a trade route connecting the Graubünden in the north and the Como Lake region in the south, we may assume that the plague entered the town at some point between November 1628 and August 1629 (but we cannot establish, if it did before the arrival of the Austrian troops, presumably in early June 1629, or thereafter). We also know that the plague abated in the region in the course of 1630, although, as noted in Section 2.10, a reconstructed parish register mentions that the priest of Domat/Ems died of plague in 1631. This potentially allows to narrow down the date of the EMS samples to the period of May 1629–1631.

The pandemic arrived in Domat/Ems from the North, most likely from South Bavaria via the Lake Constance and the Vorarlberg regions. In all three regions, the plague was spreading rife in 1628 (Eckert 1996, 137, 141–143; Baltzarek and Pradel 1973, 81, 95, 116, 129) and it was only a matter of time that it reached Chur in November of the same year, and Ems thereafter. As we have seen, by the time the Austrian soldiers invaded the region (late May-early June 1629), the plague had already been present all over the Graubünden region. But it is also possible that the invading soldiers aggravated the situation by reintroducing an additional plague strain, distinct from the one that had already been circulating in the town during the 1629-1631 outbreak. It is known that the plague was present in the Vorarlberg region from 1628 until 1630, and by the time the Austrian troops were marching through the region on the way to invade Graubünden, in May 1629, the plague was peaking in local towns and villages (Pieth 1945, 216; Baltzarek and Pradel 1973, 81, 95, 116, 129). Indeed, according to local records, the ‘patient zero’ at Thusis, was alleged to have been a soldier’s wife, dying on 17 August 1629 (Lorenz 1868-9, 36–67). Hence, we may assume that around the time of death of the individuals associated with EMS001 and EMS010 (as well as their comrades-in-arms, buried next to them at Sogn Pieder/St. Peter’s Church), there may have been two different (but closely related) strains, imported from South Germany, circulating in Ems and the wider Graubünden region – the one arriving from Chur in late 1628/early 1629 before the Austrian invasion, and the other one being imported by the Austrian troops in late May/early June 1629. This, in turn, may explain why EMS001 and EMS010 are associated with different, but closely related, plague strains.

The LAR/SPN cluster has a different story. Contrary to the BRA/EMS cluster, associated with the Austrian ‘proximate’ origin, the LAR/SPN one can be associated with the French ‘proximate’ origin, in conjunction with the French theatre of the Thirty Years’ War. In the course of the 1620s and 1630s, plague was ravaging France incessantly, on, intermittently, local and national scale, with 1628–1632 and 1636–1637, standing out as the years of outbreaks on a national scale. Just as with the Graubünden outbreak of 1628–1630, the 1628–1632 outbreak in France appears to have arrived from Germany. In that year, plague was reported all over East France, from Alsace in the North, through Burgundy, Auvergne-Rhône-Alpes to Provence in the south (Eckert 1996, 133, 137; Biraben 1975, 386–387). By

1629 it established itself in the French Alps, including Lariey, where local evidence points to an outbreak in 1629–1630, with which the LAR11t genome is associated.

The SPN19 genome is associated with a 1636 outbreak in Naturno/Naturns. More specifically, it is known that the plague arrived there either in later July or in August 1636 (Nothdurfter, Kersting, and Gebauer 2019, 41). There are two possible scenarios for the plague's arrival there and in other regions of South Tyrol. The first possibility is that it may have arrived from North Tyrol, where it is attested at various towns between 1634 and 1636, including Hall in Tirol (1634), Hötting (1634–1635), Imst (1635–1636), Innsbruck (1634–1635), Kitzbühel (1634–1635), Landeck (1634–1636), Telfs (1634–1635), and Vils (1635) (Hye 1980, 36, 58, 84, 137, 167, 233; Gapp 1986). Importantly, the sanitary situation at Naturno/Naturns was, as other Tyrol towns and villages, supervised and regulated by the Health Authority (*Provisores Sanitatis*) based at Innsbruck (Nothdurfter, Kersting, and Gebauer 2019, 41). The 1634–1636 outbreak in Tyrol appears to have arrived from Bavaria, where it was attested in 1634, possibly initiating a new wave out of its putative South Germany reservoir (Eckert 1996, 149). To associate the SPN19 genome with that wave is somewhat problematic, given its phylogenetic position in the same cluster with the LAR11t genome, shown to have been associated with the previous plague wave.

The second possibility is that the pandemic came to Naturno/Naturns and other South Tyrol regions from the West, in conjunction with the French invasion of the Valtellina region (April 1635 – March 1637), led by the Huguenot leader Henri, Duke of Rohan. Rohan's armies, mustered in late March 1635 in Alsace, marched via Muhlhausen, Basle, Bern, Zurich, St. Gallen, Mellingen, reaching Coire on 12 April (Clarke 1966, 198–199). Around that time, plague was present at some locations in the Alsace region (Eckert 1996, 149), as well as in the Basel-Jura, including Dornach, Welschenrohr and Oberdorf, although larger cities, such as Basel and Bern were spared (Eckert 1978, 71). The French advances into Switzerland and the occupation of Valtellina led to an inevitable conflict with the Austrian army, which, having mustered its forces in Tyrol, clashed with the French at the Val Federia (27 June 1635) and the Val Fraela (31 October 1635), resulting, in both instances, in a disastrous defeat of the Austrians. After the defeat at Val Fraela, the latter retreated back to Tyrol (Clarke 1966, 199–202). In his letters, Duke of Rohan, noted the challenges his army was facing between June and December 1636, because of provision shortage and plague outbreaks in Valtellina and the neighbouring regions of Tyrol and Engadin (*Mémoires et Lettres Sur La Guerre de La Valteline de Henri Duc de Rohan* 1758 Vol. 1, pp. 282–283, Vol. 2 pp. 218–220, 316–318, 339). Given the presence of plague in the French army and in occupied Valtellina region, it is possible that the infection was passed on to the Austrian troops in the course of one of the two engagements in the region, and it was then spread by them on their way back to Tyrol in November 1635, reaching Naturno/Naturns in the later summer of 1636. This scenario, assuming the French 'proximate' origin of the outbreak, is supported by the phylogenetic positioning of the SPN19 genome, forming one cluster with another genome (LAR11t) of the French 'proximate' origin, with both genomes being possibly associated with the same wave, ravaging France in the late 1620s and early 1630s. Importantly, LAR11t has three individual SNPs, while SPN19 has either three or potentially five SNPs (with two SNPs classified as potential damage). If the SPN19 sub-branch is indeed longer than the LAR11t, then it could be explained by the fact that the former is associated with a later outbreak (1636), rather than the previous one (1629–1630).

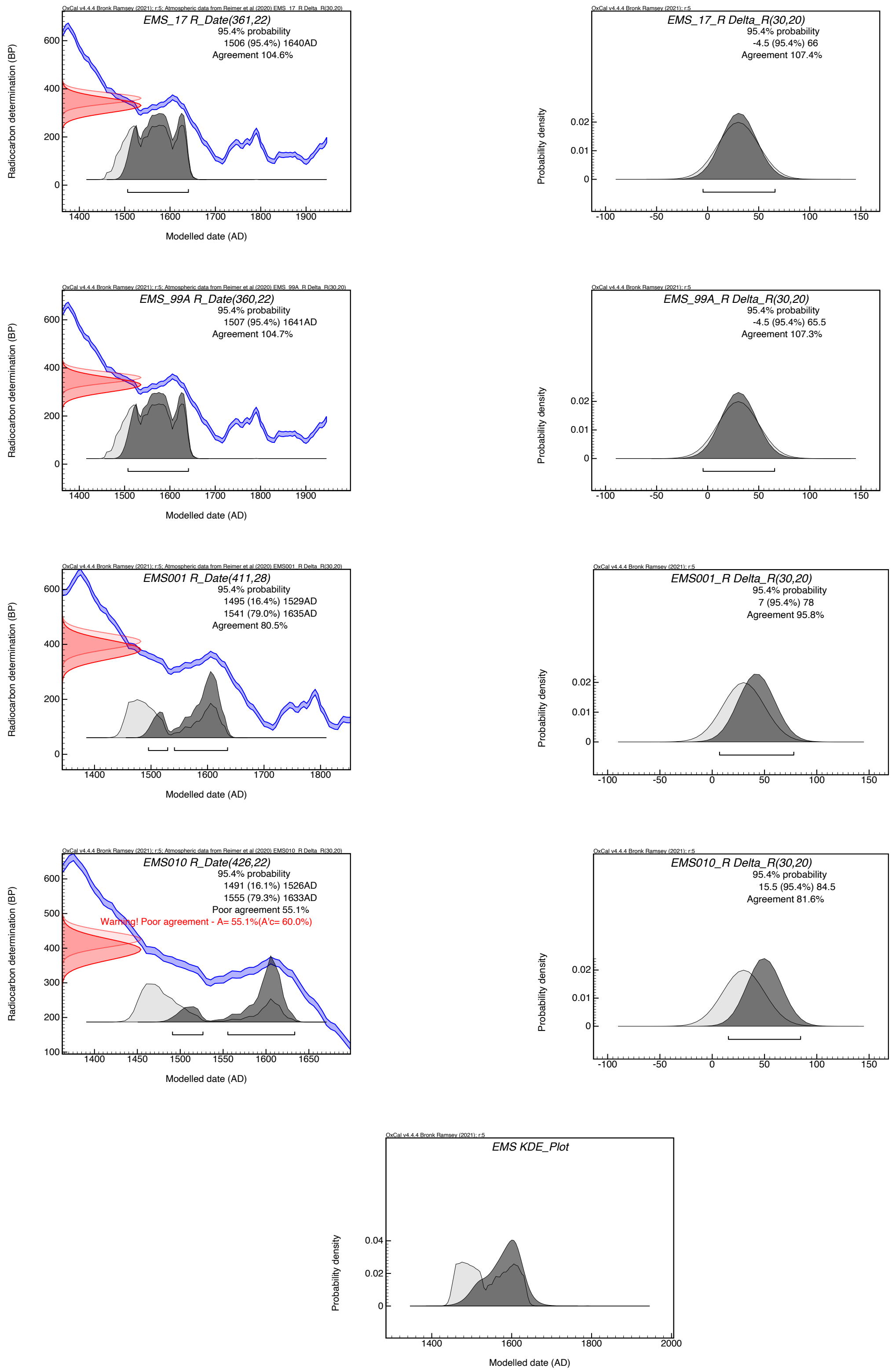

**Fig. S68: Probability distributions of radiocarbon dates (row 1-4, left), the respective offsets (row 1-4, right) as well as the KDE_Plot (row 5) for Domat/Ems within the sequence model with generic offset of 30 ± 20 RC years.**

#### 11.10 Basal Branch 1A1 – AGU007.B, AGU010.B, Gdansk8, the RUS genomes – and the provenance of Rostov2039

At the very basis of Branch 1A1, only one SNP derived from the polytomy N13, sits a cluster of five identical genomes (see Supplementary Section 5 for the respective analysis), stemming from presumably four different sites (regarding the provenance of Rostov2039 see below). Since radiocarbon dates are available for three of the sites, and the genomes act both as *terminus ante quem* for N13 and *terminus post quem* for all other Branch 1A1 genomes, they are particularly valuable for PhIRM.

The first published genomes of this cluster were the genomes AGU007.B and AGU010.B, both from Vilnius, Lithuania (Giffin et al. 2020). The date of AGU007.B has also relevance beyond the dating of the respective plague outbreak, as the individual associated with this genome was coinfected with *Treponema pallidum pertenue*, the causative agent of yaws, and its date, therefore, holds clues about a possible New World origin of European treponematoses. By combining the two dates of AGU007 and AGU010 (R_Combine function), the authors of the original study got a 2-sigma range of 1447–1616, with 83% of the probability density within the interval 1448–1498, which remains ultimately inconclusive regarding an association of the introduction of treponematosis via the Columbian exchange. While the range rather supports a pre-Columbian date for the cluster, taking any kind of offset into account will shift the range towards younger dates: adding a generic offset of 30 ± 20 RC years causes a shift of the 2-sigma range of AGU007 from 1471–1634 to 1489–1642 (see Table S11, Fig. 69).

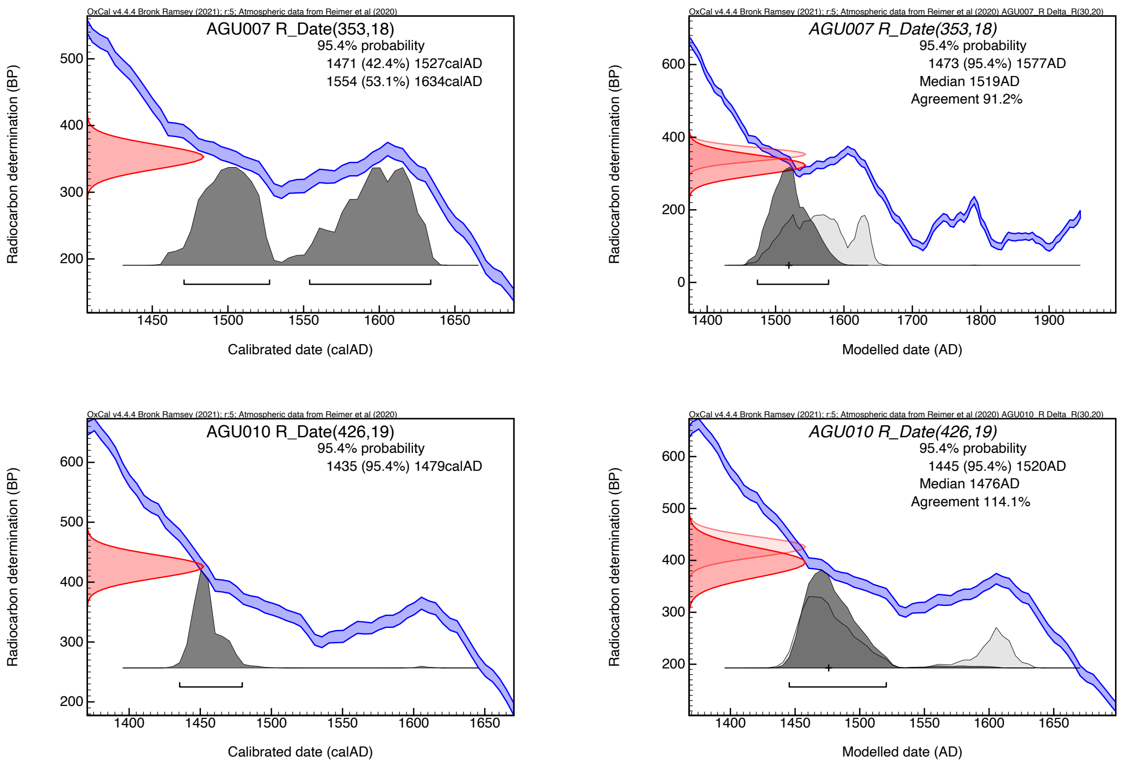

**Fig. S69: probability distributions (brackets indicating 2-sigma intervals) of AGU007 and AGU010 without offset (left) and after integrating into the sequence model with a generic offset of 30 ± 20 RC years (right).**

The genome of Gdańsk/Danzig, Poland (Gdansk8) was published shortly after the publication of the Vilnius genomes (Morozova et al. 2020) and therefore could not include the latter. The original paper gives a radiocarbon range of 1425–1469 (almost identical to the recalibration in this study to 1427–1468). The ossuaries containing respective plague victims are dated by the authors to the 15th–18th centuries in the supplementary texts, with no further chronological information. Conversely, but the main text of the publication suggests that the genome corresponds to the 1464 plague outbreak, regarded by the authors as the most severe out of six or seven plague outbreaks in the 15th century. In the same paper, the authors also published, along with Gdansk8, the genome Rostov2039. In the original paper (Morozova et al. 2020 Fig. 2), it forms a clade at the base of Branch 1A1 together with Gdansk8 and Ellwangen (ELW098). This, along with other inconsistencies with previous publications (see also the phylogenetic positions of, e.g., LAI009 and OSL1) can be traced back to a different treatment of the underlying data, and most prominently the fact that non-core regions were not excluded. Unfortunately, no radiocarbon date was provided for the sample Rostov2039, and hence, it could not be considered for PhIRM. For further discussion concerning the provenance of the Rostov2039 genome, see below.

In this study, we reconstructed the two genomes RUS003 and RUS004 from the site of Stankeyevo (Pskov Oblast, Russia), which are identical to the two Vilnius genomes, as well as the Gdańsk/Danzig genome. The Stankeyevo genomes are radiocarbon dated to 1447–1631 and 1450–1631, respectively (individual calibration without model or offset). Relying on radiocarbon date of Gdansk8 and its hypothesized association with the outbreak of 1464 would set also the Vilnius genomes well into the mid-15th century, before the Columbian exchange. However, this ignores a potential (marine reservoir) offset of the Gdansk8 radiocarbon date, all the more given the location of Gdańsk/Danzig on the Baltic coast.

Partially overlapping with wiggles in the calibration curve, the individually calibrated dates of this cluster are rather disparate with 2-sigma ranges from 1427 (absolute lower boundary) to 1634 (absolute upper boundary). Integrating them into a sequence model helped to restrict the intervals significantly: without any offset, the 2-sigma intervals range from 1431 to 1567, with a generic offset of 30 ± 20 RC years, it restricts them from 1440 to 1577. While the ranges are still relatively broad, the probability densities of the KDE_Plots are all left-skewed (see Fig. S70): the medians are at 1470 (no offset) and 1493 (generic offset of 30 ± 20 RC years). Therefore, a broad dating to the last third of the 15th century or the first decade of the 16th century is most likely, which is also in line with the distance of only one SNP from the polytomy N13, broadly dated to c.1450 – c.1500 (Fig. 2A).

Interestingly, running KDE_Plot outside of the sequence models yield younger dates still (median 1485 without offset, median 1525 with generic offset of 30 ± 20 RC years, Fig. S70). KDE_Models, which cannot be integrated into sequence models due to confounding effects, perform slightly better (in terms of providing shorter intervals) when compared to the sequence model (median 1469 without offset, 1506 with generic offset of 30 ± 20 RC years, Fig. S70). This shows that the generic offset of 30 ± 20 RC years is not strong enough to overrule the “soft” priors of the sequence model (i.e., broad radiocarbon intervals as opposed to “hard” calendar dates), also evident in the disappearance of a second peak in the probability distributions. Furthermore, the integration into the sequence model also offers an estimate of the offset of the individual radiocarbon dates – as expected, strongest for the Gdansk8 date and weaker for the AGU007 date, which is also the youngest one (Fig. S71).

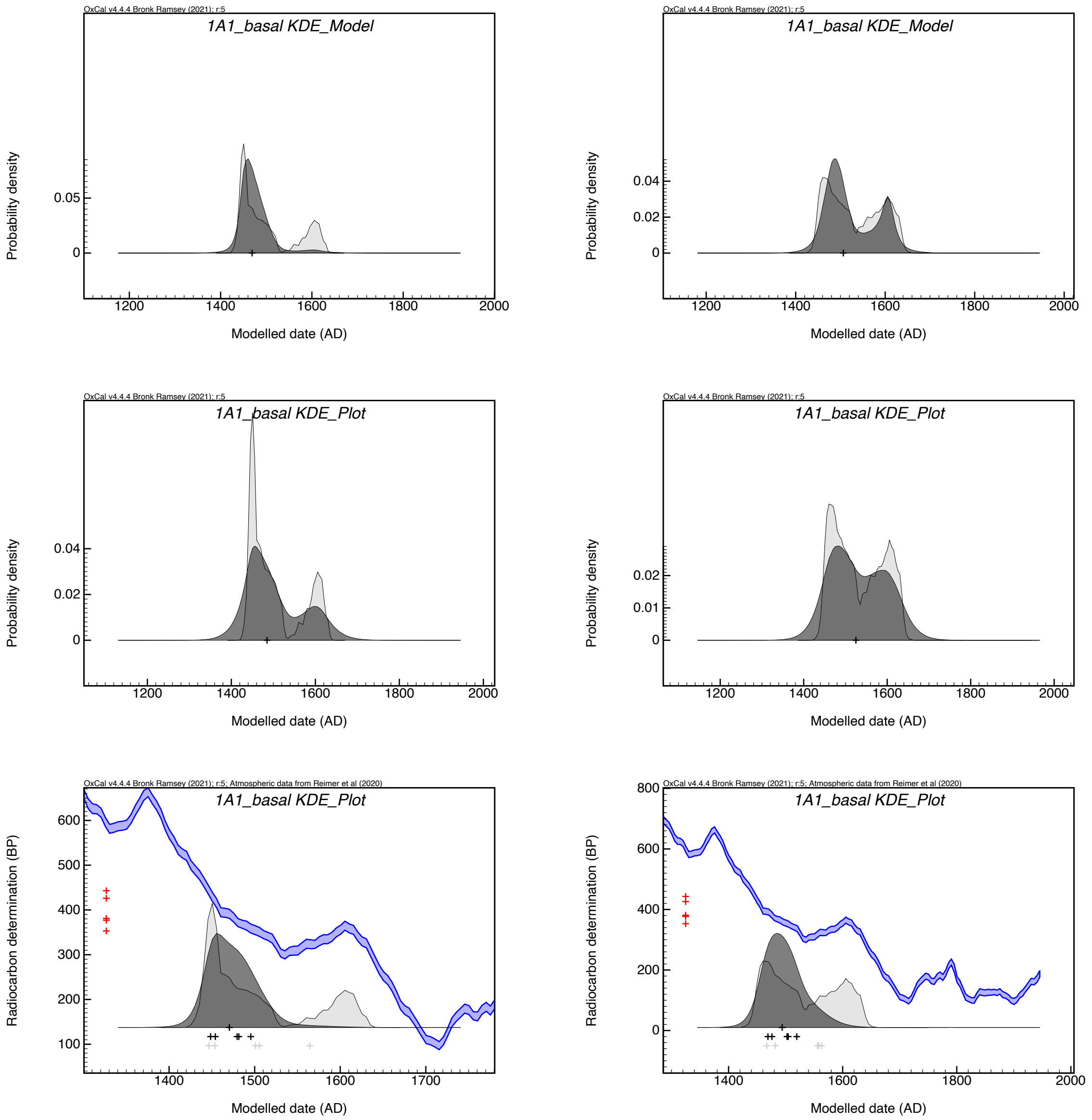

**Fig. S70: Probability distributions for KDE_Models (top row), separate KDE_Plots (middle row) and KDE_Plots within the sequence model (bottom row) without (left) and with (right) a generic offset of 30 ± 20 RC years.**

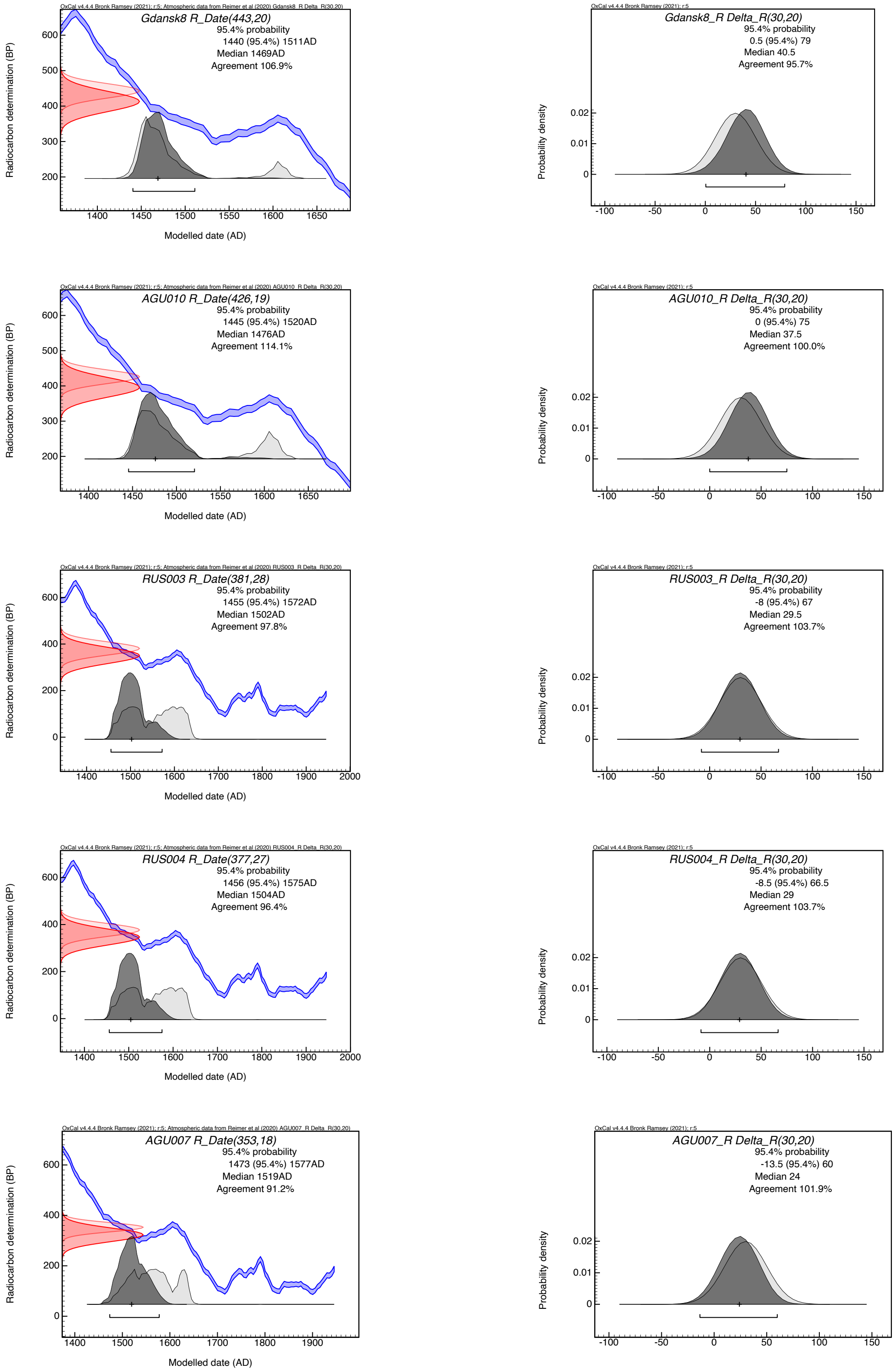

**Fig. S71: Probability distribution for the calibrated radiocarbon dates (left) and the offsets (right) within the sequence model and a generic offset of 30 ± 20 RC years.**

The genome Rostov2039 was published along with the genomes of Gdansk8, Rostov2033 and Azov38 in Morozova et al. 2020. According to the original paper, the sample comes from the cemetery of the St Dmitrii of Rostov Fortress, Rostov on Don, Russia, specifically from a ‘section of the burials’ ‘dated more precisely to 1762–1773 AD’. According to the supplementary text, “[d]ocumentary sources provided a means to connect at least a few of the burials with plague epidemics in Rostov-on-Don in the winter of 1771 (Elena Batieva, personal communication)". In the Table 1 (Morozova et al. 2020), the authors date both genomes to 1762–1773, despite the fact that they occupy vastly different positions on the phylogenetic tree. The authors acknowledged this discrepancy in their paper and ascribed it to the fact that Rostov on Don is situated “on the crossroads of multiple water and land routes” (Morozova et al. 2020), supposedly explaining the observed diversity. As established through a reanalysis of the data, none of the private SNPs of Rostov2039 passed the filtering as true-positive (Table S5), and therefore the genome must be considered identical with Gdansk8, AGU007, AGU010, RUS004 and potentially RUS003. As shown above, none of the respective radiocarbon dates for these individuals reach into the 18th century, and combining them within the sequence model with a generic offset of 30 ± 20 RC years gives an upper boundary of 1519 for the 2-sigma range of the youngest sample AGU007 (see Table S11 and above).

Therefore, the dating interval of 1762–1773 for Rostov2039 can be refuted. This discrepancy could be explained by different scenarios: 1) the site was in use already in the late 15th/early 16th century and the individual was wrongly attributed to the burials of the plague outbreaks 1762–1771; 2) the human remains, its sample, extract or library associated with Rostov2039 were misassigned and the genome stems from a different site. Two lines of evidence hint towards the second scenario: 1) the Rostov-on-Don fortress was founded only in 1761 (and completed in 1763) around the so-called Temernik Customs House (*Temernitskaya Tamozhnya*), which, in turn, had been founded in 1749. Both the Custom House and the St Dmitrii of Rostov Fortress came to replace the older St Anne’s Fortress, built in 1730–1731 (Stefanov 2011). The Custom House and St Dmitrii of Rostov Fortress (coordinates: 47.225, 39.73) were founded ~40 km to the west of St Anne’s Fortress (coordinates: 47.253333, 40.089167) on previously uninhabited sites, by the decree of Empress Elizabeth of Russia. Therefore, it seems unlikely that there were earlier burials at the site that could have been misassigned to the 1762–1771 burials (Slavin 2022). In Supplementary Figure 6 of the original paper (Morozova et al. 2020), the genome is wrongly labeled as Azov2039, whereas Azov38 is mislabeled as Rostov38 and Rostov2033 as Azov2033, which might be an indication that the genomes were assigned different IDs during data processing and the final labels were only added for the figures. Since an identical genome from Gdańsk/Danzig was published along with Rostov2039, one possibility would be that the respective sample originally comes from the same archaeological context as Gdansk8 and was mixed up in the laboratory.

Given that all five genomes (RUS004, Rostov2039, Gdansk8, AGU010.B and AGU007.B) are all situated on N14, and given that N14 is only one SNP away from the Little Polytomy (N13), we may assume that (1) the genomes come from the same wave; (2) the same wave appears to be the subsequent one after that associated with the Little Polytomy; (3) the local outbreaks associated with the genomes occurred close to the origin of the same wave; (4) the same outbreaks were, more or less, concurrent. As we have suggested, the Little Polytomy (N13) appears to have emerged c.1450 – c.1500, implying that N14 and its associated basal genomes may be dated to the last third of the 15th century or the first decade of the 16th century. To putatively associate these genomes with possible outbreaks, it is essential to identify years, in which plague outbreaks are attested simultaneously or nearly simultaneously in or around all three places - Gdańsk/Danzig, Vilnius and Stankeyevo. As the available textual evidence indicates, there is only one instance between the 1470s and the 1500s, when plague outbreaks are recorded in or around all three locations around the same time, as the following discussion, considering each wave, shows.

During the wave commencing in the South German reservoir in 1472 (see Supplementary Section 11.7), plague was reported in Gdańsk in 1473–1474 (Możejko 2012, 56–57) and Kaunas in 1474 (Lewicki 1894 no. 188 (p. 212)), but not in Pskov or anywhere near. These outbreaks appear to be associated with a wave commencing in its putative South German reservoir in 1472 (*Die Chroniken Der Fränkischen Städte. Nürnberg. Vierter Band* 1872, 330; Leidinger 1915, 621) and spreading swiftly all over Germany and Central Europe in 1473–1474. Although covering most of Germany, the Baltic ports, the same wave appears to have spared Hamburg and Lübeck this time. This implies that the epidemic was imported into Gdańsk/Danzig and Lithuania via a different route. It is known that plague was present in Gdańsk/Danzig, where it is reported in 1473–1474 (Możejko 2012, 56–57), having arrived there most likely from either south Poland or from Brandenburg. At some point in 1474, it also reached Königsberg, from where it was carried to Kaunas in the autumn of the same year (Lewicki 1894 no. 188 (p. 212)). In Riga, the plague is attested in two Hanseatic documents in early February 1475, but it may have reached the city in later 1474 (Mahling, Neitmann, and Thumse 2018 no. 276 (p. 275), 304 (p. 299)). Although there is no direct reference to a plague outbreak in Vilnius in 1474, given its documented presence in Kaunas, a relatively short distance between the two towns (~100 km), and political and commercial ties between the two, it is likely that the outbreak ravaged both Kaunas and Vilnius.

During the next wave, plague was present in Gdańsk/Danzig from March until Autumn 1484 (Możejko 2012, 57–58). The origins of this outbreak remain, at this stage, unclear. On the one hand, it is possible that it may have been associated with a wave commencing in South Germany in 1482 (*Die Chroniken Der Fränkischen Städte. Nürnberg. Vierter Band* 1872, 369; Müllner 2003, 64; Sturm 2014, 37), spreading all over the country in 1482–1483 and reaching Lübeck in early 1483 (Ibs 1994, 122–124). On the other hand, the 1484 outbreak in Gdańsk/Danzig may have been associated with a different wave. By the time the putative South German reservoir was radiating its wave all over Germany in 1482–1483, southern and central Poland appears to have been ravaged by a different (concurrent) plague wave (Walawander 1932, 179–180). As several Polish and Russian sources indicate, the plague came to Poland from Hungary (*Kronika Polska, Litewska, Zmodzka I Wszystkiej Rusi Macieja Stryjkowskiego* 1846 Vol. 2, p. 286, *Ipat'yevskaya Letopis'* 1843, 2:359). A close analysis of the available source material reveals that the plague ravaged Hungary in 1479–1480 (Magyary-Kossa 1931, 101–102), having arrived there from Wallachia, where it had circulated in 1476–1479 (Iorga 1970, 80; Andreescu 1977, 267–268; Vătămanu 1972, 1972) – possibly introduced there by refugees from Caffa fleeing to Moldovan and Wallachian territories on the north-western Black Sea littoral, following the city's fall to the Ottomans in 1475. From Hungary, the wave carried on northwards, via Slovakia (then part of the Kingdom of Hungary; Majorossy 2012, 105) into southern Poland, where it is attested in the Carpathian region in 1480–1481, before spreading all over southern and central parts of the country in 1482 (Walawander 1932, 179–181). In other words, it is possible that around

1484, two concurrent waves were circulating around Gdańsk/Danzig (and other regions in Baltic Poland), spreading two strains originating in two different reservoirs.

Although there is no data about the spread of the same wave into Vilnius or elsewhere in Lithuania in 1482, plague is attested in both Riga (Gahlbeck et al. 2020 no. 528 (p. 504)) and Tallinn/Reval (Tallinna linnaarhiiv, F191, N2, S1, fol. 42; Johansen and von zur Mühlen 1973, 275) in the same year. To reach Latvia and Estonia from Poland, it was bound to travel via Lithuanian territories, implying that the presence of plague in Vilnius in 1482 is a possibility. Conversely, it is not until 1487 that plague is attested in Pskov (Nasonov 1941, 80; Nasonov 1955, 223).

During the next wave commencing in South Germany in 1492 (see Supplementary Section 11.7), plague was reported in 1494–1495 Gdańsk/Danzig (Możejko 2012, 58–59), and in 1495 in Lithuania (Ulashchik 1980, 35:123), but nowhere near Pskov or elsewhere in West Russia. During the subsequent wave, however, plague is known to have ravaged in all three cities around the same time: in 1505 in Gdańsk/Danzig (Możejko 2012, 59–60), in 1505–1506 in Vilnius (and elsewhere in Lithuania) (*Kronika Polska, Litewska, Zmodzka I Wszystkiej Rusi Macieja Stryjkowskiego* 1846 Vol. 2, p. 328, 345), and in 1506–1507 in Pskov (Pogodin 1837, 173; Nasonov 1941, 91; Nasonov 1955, 225).

Obviously, the lack of surviving references to plague outbreaks does not mean that these outbreaks did not happen. We may be, more or less, certain that the lack of any mention of plague in western Russian towns, except Novgorod, between 1467 and 1487 in over 50 Russian chronicles and other documents may indicate that *major* western Russian towns were spared during these years. However, the same chronicles would record events in more minor settlements very seldom, mostly in conjunction with military conflicts. The Stankeyevo context is to be associated with the town of Krasnogorsk, founded by Ivan Alexandrovich Zvenigorodskii, Prince of Pskov, in 1464 as a minor fort with an attached settlement (Kal'yu 2004, 15–17; Derkach and Shun'gina 2018). Although population figures are lacking, it is unlikely that the young town had a population of more than a few hundreds by the time of the plague outbreak. As such, its early history, including plague outbreaks, remained unrecorded. However, if there was an undocumented outbreak in Krasnogorsk around 1474–1475 or 1495–1496, then it did not spread from there to major West Russian cities and trade hubs, such as Pskov, Velikie Luki, and Staraya Russa. There was an outbreak in Novgorod in 1477–1478 (Pogodin 1837, 173: 151). However, it was most likely imported there from northern Estonia, via Narva, given that it was not reported in Pskov. In other words, while it is possible that there was an outbreak in Krasnogorsk around 1474–1475 or/and 1495–1496 (arriving there from, say, Latvian or Belorussian territories), there is no textual witness to that, because the same wave reached neither Pskov, nor other cities in the region, if we are to accept the silence of local chronicles as an indication of that.

The same can be said about Vilnius regarding the lack of reference to an outbreak around 1482–1484, reported in both Riga (1482) and Polish territories (1484). Although there is no evidence that there was such an outbreak, it is important to bear in mind that our information about pre-sixteenth-century outbreaks in Lithuania is very patchy, deriving either from sixteenth-century Polish chronicles or from occasional references in state papers of the Duchy of Lithuania. Hence, it is possible that it may have ravaged Vilnius and other

Lithuanian territories, around the same time it had been present in neighboring Latvian and Polish territories, but the same outbreak remains undocumented.

Regardless of the precise identification of the outbreak related to the basal Branch 1A1 cluster, all of the waves appear to have originated in South Germany, as the discussion above indicates. We assume that the Ottoman Reservoir was established thereafter and the observed cluster is an offshoot of the Branch 1A1 before or while moving towards the Ottoman territory. However, the currently available genomic and historical data do not allow to test this against alternative hypotheses.

#### 11.11 The Ottoman Reservoir of Branch 1A1

Regarding the phylogenetic structure of the Branch 1A, the internal branch between the basal Branch 1A1 cluster (N14) and the BED cluster (N15) is remarkable for several reasons. In the present analysis, a branch length of 43 SNPs was determined, by far the longest internal branch within this clade. As shown in Spyrou et al. 2019, this branch is characterized by an elevated substitution rate, as reflected in the different genetic distance of the likely contemporary BED cluster and the Thirty Years' War plague cluster from their MRCA (N13) – 44 SNPs versus 19–20 SNPs (see Fig. 2A). Since the BED cluster is directly derived from the basal Branch 1A1 cluster, we can, in this instance, use the boundary function in the sequence model to get a rough estimate of the timespan between the two events, with the Boundary T8 is estimated to 1504–1605 (2-sigma interval, see Fig. S72). In addition, a large deletion in the chromosome of all genomes of this clade, which – by law of parsimony – must have happened between nodes N14 and N15. Whether there is a causal relationship between these two phenomena or whether they merely coincidentally remain an open question, but a hypothesis may be offered.

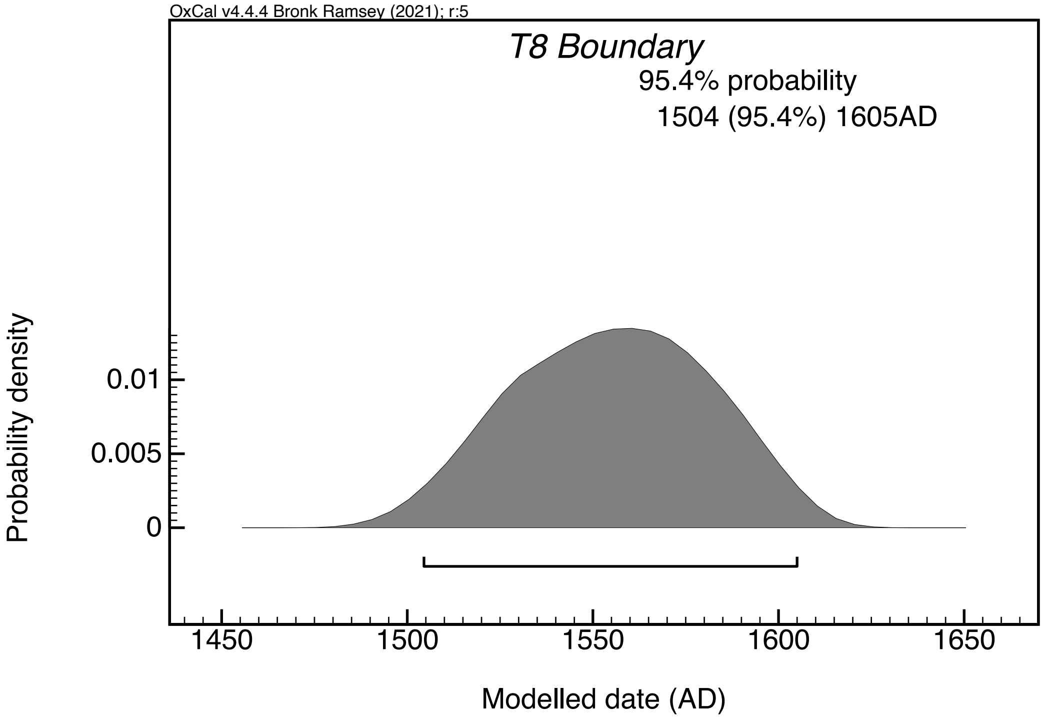

**Fig. S72: Probability distribution for Boundary T8 as part of the sequence model with a generic offset of 30 ± 20 RC years.**

One possible causal relation is that both phenomena are independently or dependently linked to the migration of 1A1 strains to a new reservoir, situated in a new natural environment, characterized by a different ecological and climatic niche compared to the original reservoir in South Germany. As the reconstruction of the spatio-temporal origins and spread of plague waves associated with 1A1 genomes below indicates, they all point into the Ottoman territories as their point of origin. This is particularly apparent in relation to those genomes that are securely or nearly securely dated – the Great Northern War plague genomes (PRN001 from Tallinn/Reval and PEB10 from Pestbacken in Sweden, both dated to 1710), Rostov2033 from Rostov-on-Don (most likely dated to 1771–1772) and OBS genomes associated with the Great Plague of Marseille (1720–1722) (Slavin 2022). The analysis of undated genomes (the BED genomes from London, Azov38 (from Azov, South Russia) and CHE1 (from the Chechen-Georgian border) reveals their Ottoman provenance too (see below, Sections 10.12 and 10.14).

The existence of a putative Ottoman reservoir has been already suggested by historians, with Moldova-Wallachia, South-West Balkans, the Istanbul hinterland, East Anatolia, Kurdistan, the ‘Asir Region (the Saudi-Yemeni border), and Egypt all highlighted as possible candidates (Panzac 1985, 81–96; Varlık 2022). Although the exact whereabouts of the putative Ottoman reservoir(s) cannot be established at this point, it is certain that wherever they were located, they had different eco-climatic systems of biotic and abiotic components from those of the temperate forest environment of the South German reservoir. This implies not only different climate, vegetation, soils, but also different mammalian hosts. The common vole has been hypothesized as the main host in the South German reservoir (Slavin 2021), while for the Anatolian one, the jerboa and the Persian jird were suggested, among others (Varlık 2022, 168–172). Possibly, the exposure to and acclimatization into a new reservoir environment was a major factor triggering the large deletion on the chromosome and elevation in substitution rates, discussed above.

The fact that no 1A1 branch-associated European genomes from the 16th century have been uncovered on the long segment between N14 and N15 (acquiring 43 SNPs in the course of about a century) may be explained by the migration of the nascent 1A1 into a new reservoir in the Ottoman territories, at some point soon after the outbreak associated with the N14 genomes cluster (RUS004, Rostov2039, Gdansk8, AGU010.B and AGU007.B). It is also important to consider the international trade situation as a factor holding back the spread and circulation of 1A1 strains in Europe. During most of the sixteenth century, the commercial contacts between European and Ottoman merchants was dominated primarily by the Venetians and, to a lesser extent, French (Masson 1896; Munro 2007). All in all, the volume of European-Ottoman transactions and exchange was comparatively low. The situation changed rapidly from the late 16th century on, with the formation of the English Levant Company in 1592 and the Dutch-Ottoman treaty of 1612, catapulting, respectively, English and Dutch merchants to the forefront of international trade with the Ottoman Empire, at the expense of the declining power of the Venetians (Epstein 1908; De Groot 1978). With regular presence of English and Dutch merchants in key Ottoman ports (especially, Smyrna/Izmir, Aleppo, Alexandretta/Iskenderun, Constantinople/Istanbul, Algiers and

Cypriote cities) and annual shipments to and from these ports, the volume and frequency of European-Ottoman trade grew up considerably in the course of the 17th century. As noted in the next sections, the fact that textiles and textile-based goods (that could, in theory, harbour ectoparasites infested with *Y. pestis*) constituted the major products exported by the English and Dutch merchants, undoubtedly facilitated the recurrent reimportation of 1A1 strains from their putative Ottoman reservoir(s) in the 17th and 18th centuries – in contrast with the 16th century, characterized by limited commercial contacts between the Ottoman Empire and West/Central Europe.

#### 11.12 The London New Churchyard genomes (BED) and Azov38

The genomes of London New Churchyard (also known as London Bedlam) BED024, BED028, BED030 and BED034 were published by Spyrou et al. 2019. The respective site, a municipal burial ground, was opened in 1569 and closed in 1739. The plague genomes derive from human remains found in a mass burial containing at least 42 individuals, whose orientation is consistent with the early phase of the burial ground (1569–1670). Pottery found in the fill was dated to the period 1550–1610 and coffin type identified in the pit was "appeared in the last quarter of the 16th century and was ubiquitous from 1650 onwards" (Spyrou et al. 2019). The original assumption that the burials represent victims of the Great Plague of London in 1665 was however refuted by radiocarbon dates. The dating interval in Table 1 (Spyrou et al. 2019) of 1560–1635 originates in an unpublished report by Derek Hamilton (SUERC) and Peter Marshall (Historic England) providing the raw radiocarbon dates as well as δ13C and δ15N values and the results of a modelling. A first modelling attempt used the years 1569 and 1739 as priors for the lower and upper boundary and combined the dates of five individuals (8147, 8193, 8219, 8198/BED034, 8216/BED039); unfortunately, the original code is not provided in the original report of Hamilton and Marshall. This first modelling attempt was statistically not consistent, and an outlier analysis identified the date of 8193 as a misfit. Excluding 8193 resulted in a model with good agreement and resulted in a 2-sigma interval of 1590–1620. In a second model, the authors took a marine reservoir offset into account which they calculated to 31 ± 56 RC years, yielding a 2-sigma interval of 1560–1635. Therefore, in the supplementary text (Spyrou et al. 2019) plague outbreaks in 1603, 1625 and 1636 were listed as potential dates for the respective genomes. One may add the earlier outbreaks of 1574–1582 and 1592–1593 (Porter 2005), also fitting into this dating range.

In the dating approach of this study applying PhIRM, we used solely the phylogenetic information, ignoring other contextual information and also applying a generic offset of 30 ± 20 RC years to all samples except the excluded outlier (8193). The relevant priors for the BED cluster were the radiocarbon dates of the basal Branch 1A1 genomes as *terminus post quem*, as they represent the direct ancestor and the date 1710 for the genomes representing the plague of the Great Northern War (see Supplementary Section 11.13) as direct descendants. This caused a restriction of the 2-sigma ranges from 1406–1635 to 1537–1652 (lowest and upmost boundaries of all four samples, respectively, see Table S11 and Fig. S73). The median of the KDE_Plot is at 1603, corresponding perfectly with an outbreak in London reported in that year. Nevertheless, also the outbreaks in 1625 and – although less likely – in 1636, should be considered. Interestingly, the sequence model without generic offset gave similar 2-sigma ranges from 1534 to 1649 (lowest and upmost boundaries of all four samples, respectively, see Table S11) and a similar KDE_Plot median (1599), but gave low agreement values for two of the four samples. Therefore, the assumption of a marine reservoir effect by Hamilton and Mashall could be confirmed indirectly.

In contrast to the sequence models, KDE_Plots and KDE_Models give significantly older median dates (ranging from 1481 for KDE_Model without offset to 1535 for KDE_Plot with a generic offset of 30 ± 20 RC years) due to pronounced bimodal distributions (Fig. S74).

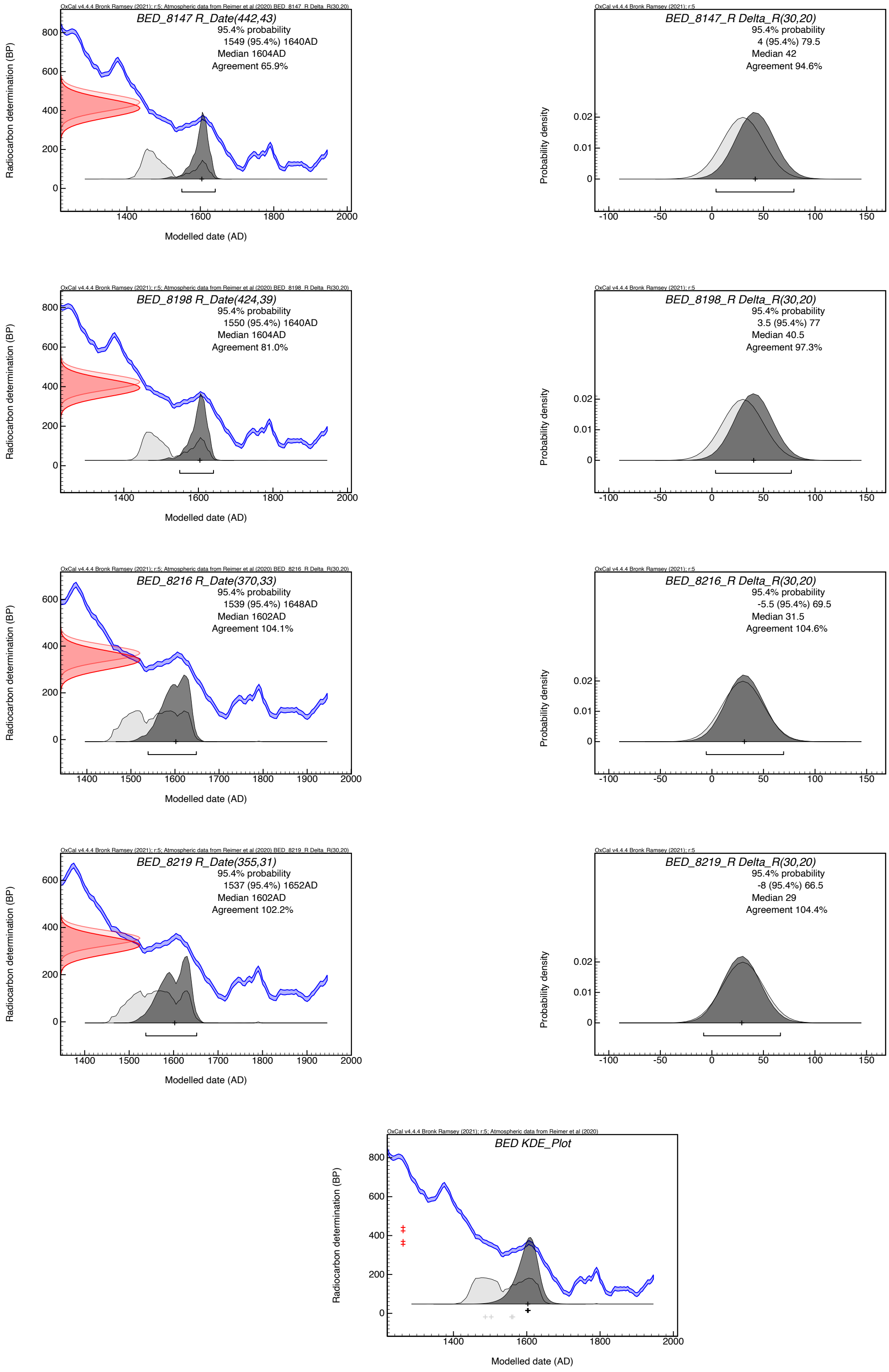

**Fig. S73: Probability distribution for the calibrated radiocarbon dates (row 1-4, left) and the respective offsets (right) as well as the KDE_Plot (row 5 left) within the sequence model and a generic offset of 30 ± 20 RC years.**

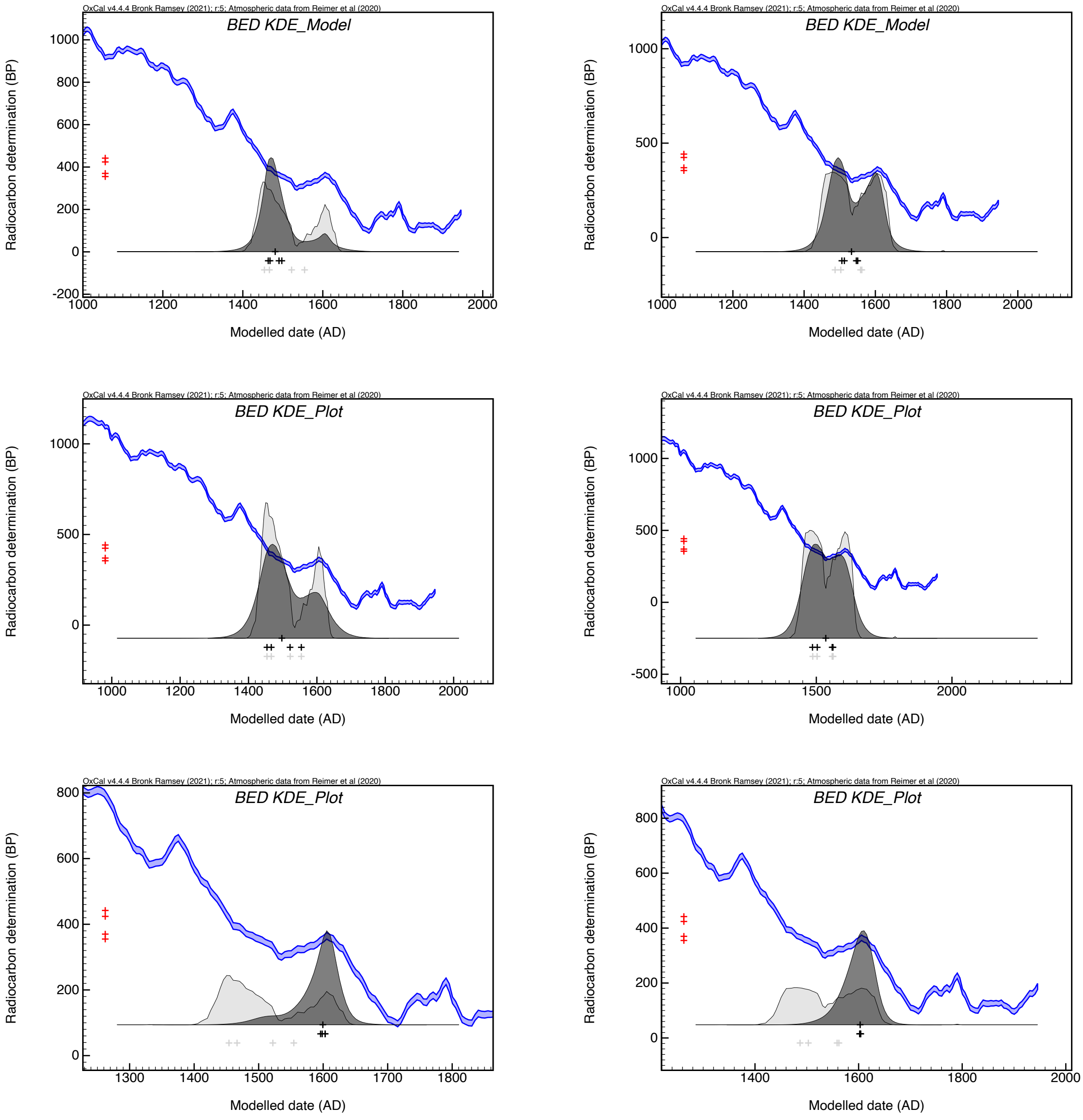

**Fig. S74: Probability distributions for KDE_Models (top row), separate KDE_Plots (middle row) and KDE_Plots within the sequence model (bottom row) without (left) and with (right) a generic offset of 30 ± 20 RC years.**

The genome of Azov38 was published by Morozova et al. 2020 and dated to the 15th–17th centuries in the main paper. The supplementary text mentions that burial artifacts and stratigraphy date the remains to the 15th–18th centuries. Phylogenetically, the genome of Azov38 is a direct descendant of the BED cluster and shares one SNP with all the other younger Branch 1A1 genomes. From this node (N16), it branches off with a terminal branch – 3 private SNPs passed the SNP filtering with two being classified as potential damage. Because of its derived position, it must date younger than the BED cluster. Considering the PhIRM results and contextual data of the London New Churchyard burials, the *terminus post quem* of the New Churchyard cemetery opening in 1569 can also be applied to Azov38. In addition, the radiocarbon date modelling – without considering the cemetery opening date as a prior – also hints, independently, towards the first decade of the 17th century for the BED cluster. Therefore, the date interval for Azov38 can be restricted to a broad period of the last third of the 16th century and the 17th century.

Although based on the radiocarbon dating of 1560–1635, combined with the opening date of the graveyard (1569) the BED cluster can be theoretically associated with any of the five outbreaks in London listed above (1574–1582, 1592–1593, 1603, 1625 and 1636), only one of these appear to have been firmly linked to the Ottoman territories as their origins, as far as the available historical evidence can tell.

The period of 1574–1582 saw a continuous circulation of plague all over the British Isles, with epidemics reported in London every year, albeit in a much lighter form compared to other 16th and 17th century outbreaks. During that wave, the annual mortality toll would fluctuate between about 2,000 and 4,000 Londoners, in comparison with about 21,000 in 1563 and 1592–1593, 30,000 in 1603, 46,000 in 1625, 17,500 in 1636 and 70,000 (perhaps 75,000) in 1665 (Cummins, Kelly, and Ó Gráda 2016; Porter 2005, 249). The 1574–1582 wave appears to have commenced with the introduction of plague into the port of Edinburgh or Leith in October 1574 (Burton 1878, 45, 49). Around the time of the 1574 outbreak, the vast majority of trade goods imported to Scotland came from either Hanseatic ports of the Baltic (primarily Gdańsk/Danzig and Stralsund), or from coastal Netherlands (primarily Veere, Ostend/Oostende, Middelburg, and Rotterdam; Lythe 1955; Ditchburn 1990; Rorke 2001). Indeed, late 16th- and early 17th-century Scottish state papers have enough references to the importation of plague-infected cargos of grain and flax, as well as sick passengers from those regions (Oram 2006, 27). Of all the aforesaid trade hubs, only coastal Low Countries ports were infected by plague in late 1574 (Noordegraaf and Valk 1996, 226). As surviving port books from several Scottish ports indicate, in the early 1570s the majority of Scottish commercial shipping was bound to and from coastal Netherlands, followed by Gdańsk (where plague is attested in 1564 and 1580, but not in 1574) (The National Records of Scotland, E71/1/6-10; E71/2/1; E71/5/1; E71/7/1-2, 4; E71/17/1-2; E71/21/2; E71/23/1; E71/25/1; Kizik 2012, 71). Hence, the 1574 outbreak was very likely to have been imported from one of the ports of coastal Low Countries. In this case, the same outbreak may have been associated with a wave that had commenced in South Germany in 1570–1571, before spreading all over Germany and the Low Countries in the next few years (see Supplementary Section 11.6). In the Netherlands, plague was attested all over the place between 1572 and 1575 (Noordegraaf and Valk 1996, 226). If this interpretation is correct, then the strains circulating in London and elsewhere in the British Isles during the same wave may have been caused by Branch 1A2 associated with the German reservoir, rather than by 1A1, on which the BED genomes are positioned.

The exact route of the 1592–1593 outbreak to London is, at this point, unclear. It may have spread into the city from one of Devon ports, perhaps Plymouth, where it had been attested in 1590–1591. The arrival of the pathogen in Devon ports appears to be a part of a long plague wave in the British Isles commencing in (most likely June or July) 1584 in Wester Wemyss (Fife, Scotland) (Masson 1880, 679–680), spreading into North-eastern Scotland in the same year and into southern Scotland (by land and sea) and some east English ports (by sea) in the following year, before carrying on to most of English regions (by both land and sea), in the course of the later 1580s and early 1590s (Shrewsbury 1971, 222–263). Around the time of the 1584 outbreak, the structure of Scottish maritime trade was the same as around the time of the 1574 importation, with the vast majority of trade goods being imported from either the Baltic ports of Gdańsk/Danzig and Stralsund, or from coastal Netherlands (Veere, Ostend/Oostende, Middelburg, and Rotterdam; Lythe 1955.; Ditchburn

1990; Rorke 2001). Of all the aforesaid trade hubs, only coastal Low Countries ports were infected by plague in 1584 (*Calendar of State Papers, Foreign: Elizabeth. Volume 19, August 1584-August 1585* 1916, 23). As surviving port books from several Scottish ports indicate, in the early 1580s the majority of Scottish commercial shipping was bound to and from coastal Netherlands, followed by Gdańsk/Danzig (where plague is attested in 1580 and 1588, but not 1584) (The National Records of Scotland, E71/1/6-10; E71/2/1; E71/5/1; E71/7/1-2, 4; E71/17/1-2; E71/21/2; E71/23/1; E71/25/1; Kizik 2012, 71). Hence, the 1584 outbreak was very likely to have been imported from one of the ports of coastal Low Countries. In this case, the same outbreak may have been associated with a wave that had been circulating all over Germany in the early 1580s. It is unclear if it was a continuation of a wave from the 1570s, or if it may have commenced in the putative South-German reservoir in 1582, before spreading fast in all directions (Eckert 1996, 94–99). In either scenario, if this interpretation is correct, then the strains circulating in London in 1592–1593 may have been caused by Branch 1A2 associated with the German reservoir, rather than by 1A1, on which the BED genomes are positioned.

The 1603 outbreak could be imported either from Amsterdam, via Great Yarmouth, where it had been imported in September, or directly from Gdańsk/Danzig in October (Noordegraaf and Valk 1996, 227; Kizik 2012, 72). In both instances, local authorities reported the arrival of infected goods and people as the cause of plague outbreak (Porter 2005, 81–82). However, it appears as if the outbreak in Gdańsk/Danzig was abating in the course of 1603, contrary to the situation in Amsterdam. Hence, it is possible that Amsterdam may have been a more likely candidate. The 1602–1603 outbreaks in Amsterdam and Gdańsk/Danzig appear to have been associated with the same wave that had begun in South Germany in 1594, and spread all over the country in 1595–1598 (discussed in Supplementary Section 11.6). If this interpretation is correct, then the strains circulating in London in 1603 may have been caused by Branch 1A2 associated with the German reservoir, rather than by 1A1, on which the BED genomes are positioned.

It is, however, the 1625 outbreak that appears to be the best candidate to be associated with the BED genomes, given its origins. Between 1619 and 1626, plague was ravaging in Egypt (Raymond 1972, 204–205) and between 1620 and 1624 in the Maghreb (Boubaker 1995, 317). In 1623–1625 it was present in Constantinople/Istanbul (*The Negotiations of Sir Thomas Roe, in His Embassy to the Ottoman Porte, from the Year 1621 to 1628 Inclusive* 1740, 459–560) and in 1624–1625 it was reported in Iskenderun (Ayalon 2015, 82–83) and Cyprus (Jennings 1993, 187). Contrary to the 1584 importation into Scotland and the 1603 and 1636 ones into England, the 1625 outbreak may have been directly imported into London in an infected cargo ship belonging to the Levant Company. Importantly, between 1621 and 1647, Dutch trade in the Levant was paralyzed because of the Spanish maritime blockade of the Mediterranean (Israel 1989, 121–196). As a result, long-distance trade was at the hands of the Venetians and the English Levant Company established in 1592 and active in various Levantine hubs, but especially in Smyrna/Izmir, Aleppo, Alexandretta/Iskenderun, Constantinople/Istanbul, Algiers and Cypriote cities (Epstein 1908). Given the chronology of plague spread in the Middle East and Maghreb, Constantinople/Istanbul, Iskenderun and Cyprus appear to be the most likely candidates to be an ‘export port’. However, the plague was also present in the Low Countries (and Amsterdam in particular) in 1623–1626, as well as in Hamburg and its surrounding region in 1625, implying that the plague could still potentially be imported from one of North Sea ports

(Noordegraaf and Valk 1996, 227). In this instance, however, the 1625 outbreak would have more likely been associated with a 1A2 rather than a 1A1 strain, given the commercial disconnect between the Dutch Republic and the Ottoman Empire in that year. The fact that the BED genomes are associated with the 1A1 branch contradicts this scenario.

The 1636 outbreak is unlikely to be associated with the BED genomes, for the following reasons. The outbreak in question had arrived in London from the Netherlands (ravaging their regions in 1635–1636; Slack 1985, 323; Noordegraaf and Valk 1996, 227), when Dutch trade and shipping in the Middle East and North Africa was still on hold, because of the Spanish maritime blockade continuing until 1647. The 1635–1636 outbreak in the Netherlands and the 1636 one in London is to be associated with one of Thirty Years' War plague waves, circulating all over Central Europe, northern Italy, the Netherlands and England, having originated in their putative South German reservoir. This is indeed corroborated by the fact that all the so-far sequenced Thirty Years' War plague genomes from four sites (see Supplementary Section 11.9) are associated with the 1A2 branch, suggesting their Central European, rather than Ottoman origins.

If the reconstruction of the spatio-temporal origins of the four London outbreaks is correct, then it appears that the 1625 one is the most likely candidate to be associated with the BED genomes, given their Ottoman origins.

The BED genomes are directly situated on the node N15 without any private SNPs, meaning that the phylogenetically younger Branch 1A1 strains are direct descendants of the BED strain. There are two possible scenarios: a) the lineage traveled to London via trade with the Ottoman Empire and bounced back there some time after the outbreak, potentially after establishing a short-lived local reservoir in or around London, and b) the lineage had established a long-term reservoir in the Ottoman Empire and the BED cluster represents an offshoot that was not able to acquire private SNPs on its maritime itinerary from an Ottoman port to London (similar to the Black Death, with identical genomes found in the Mediterranean and London East Smithfield). A third possibility – the introduction of later Branch 1A1 outbreaks such as the Great Northern War plague and the Plague of Marseille – from a long-term reservoir in England contradicts the well-established chronologies of those outbreaks and can therefore be excluded. We favour the second scenario, which is corroborated by historical contextualisation as well as spatio-temporal origins of the later Branch 1A1-associated genomes and their short genetic distance to the BED cluster, discussed in a detail in the next sections.

As noted above, the original publication dated Azov38 to the 15th–17th centuries in the main manuscript and to the 15th–18th centuries in the SI, based on burial artifacts and stratigraphy (Morozova et al. 2020). The burial associated with the Azov38 genome was unearthed during the 2012 excavations in Azov, and its precise location is Kalnin St. 38/Ulitsa Kalinina 38 (coordinates: 47.1145, 39.4136). The site in question has been mentioned in one subsequent archaeological publication, which dated the burial to the 17th century (Goncharova et al. 2014). With no finer-tuned dating, based on numismatic, ceramic or any other material, we can tentatively date the Azov38 genome to a rather wide interval 1565–1700.

For this period, there were plague outbreaks in the Azov region and other nearby territories in the Crimean Peninsula, South Ukraine, North Caucasus and the Pontic-Caspian Steppe. Thus, there was an outbreak in 1571 in Crimea (Varlık 2015, 191–192), 1579 specifically in Azov (Varlık 2015, 198), in 1607 in South Ukraine (Sultan 2004, 46; Kompaniets 2022, 30), in 1628–1629 (Prokhorov 2016, 321) and 1636 in Crimea (Vasil'yev and Segal 1960, 52), in 1640 specifically in Azov (Sultan 2004, 62), in 1648 in Crimea (Abduzhemilev 2016, 346), in 1668–1669 in Crimea and the Azov region, including the city itself (Borisenkov and Pasetskii 1983, 217; Alexander 1980, 19); in 1673, 1677 (Alexander 1980, 19), 1680–1681 (Prokhorov 2016, 325–326; Alexander 1980, 19), 1689 (*Debar Śepatayim: An Ottoman Hebrew Chronicle from the Crimea (1683-1730). Written by Krymchak Rabbi David Lekhno* 2021, 46, 81) and 1697–1698 in Crimea (*Seyid-Muhammed Riza. Sem' Planet v Izvestiyakh O Tsaryakh Tatarskikh* 2019, 258). To reach these territories from the putative Ottoman reservoir, the bacteria would have to travel either by sea from Istanbul or any other port on the south littoral of the Black Sea, or by land either via eastern Balkans, Moldova and South-West Ukraine, or via Caucasus. In case the association of the BED genomes with the 1625 outbreak in London is correct, then the 1636 and 1640 outbreaks could be potentially the best fits to be identified with Azov38, given its relative phylogenetic positioning. However, without a finer archaeological contextualisation of the Azov38-associated burial, this argument remains a pure hypothesis.

#### 11.13 Genomes of the Plague of the Great Northern War – PEB10, the PRN genomes, LHM001 and potentially MOI001

The genomes PEB10 (Pestbacken, Sweden) and PRN001 (Tallinn, Estonia) can be precisely associated with the plague of the Great Northern War. PEB10, published by Guellil et al. 2020, derives from the cemetery dated to the 18th century. In total, 15 coins minted between 1667 and 1710 were found in the graves; together with the fact that the graves were not part of a regular churchyard and were arranged in proper, undisturbed rows, it further supports the idea that the burials are associated with an outbreak of plague in southern Sweden in 1710–1711.

Similarly, the site of Pärnu mnt (Pärnu St.) 59B in Tallinn can be securely dated to 1710, to a plague outbreak during the siege of Tallinn/Reval (see Supplementary Section 2.1). This is most likely also the context of the genome recovered from Lehmja, LHM001, recovered from an exceptional burial (see Supplementary Section 2.2) of three young men who were probably part of a Russian dragoon regiment.

The plague-positive individuals of Tallinn and Lehmja were also radiocarbon dated to confirm the archaeological-historical dating. However, due to the wiggles in the calibration curve between c.1650 and 1950, the precision of calibrated radiocarbon dates is poor and also impairs modelling approaches. The radiocarbon dates of the PRN and LHM samples with and without generic offset as well as KDE_Plots and KDE_Models for PRN are presented in Table S18.

In the phylogenetic tree, PRN001 and PEB10 share a short branch of one SNP, derived from a node (N17) between the node giving rise to the Azov38 terminal branch and the node giving rise to the CHE1-Rostov2033 clade. PEB10, directly derived from PRN001 in our phylogenetic analysis (Fig. 2A), accumulated between 1 and 4 SNPs that passed SNP filtering (3 classified as potential damage, see Table S5). This would be compatible with the

timeline of the plague of the Great Northern War, with the disease reaching Tallinn/Reval in August 1710, and Stockholm shortly after – probably in September of the same year, on a ship from either Pärnu/Pernau or Helsingør/Elsinore (Jordan 1880; Oja 1996; Zapnik 2007, 46–57; Frandsen 2010, 60–64). Other low-coverage genomes from the same site in Tallinn (PRN004, PRN005, PRN008, PRN009) and Lehmja (LHM001) are most likely identical (see Supplementary Section 7).
Unfortunately, neither the contextual data of the site of Mõisaküla nor the genetic data (MOI001, mean coverage of 0.2-fold) allow us to attribute the respective burial to a certain phase of the Second Pandemic. However, the radiocarbon date suggests a date younger than 1640 (2-sigma, without offset, Fig. S75), therefore this individual might have also been a victim of the plague of the Great Northern War.

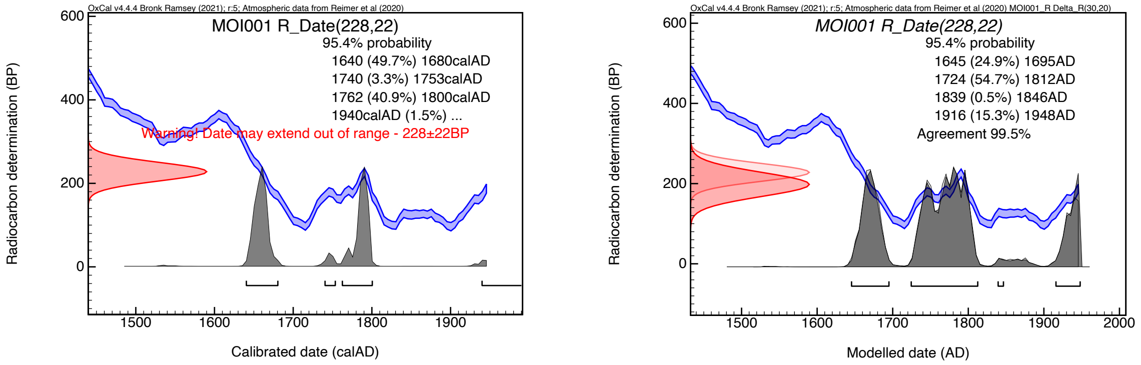

**Fig. S75: probability distributions (brackets indicating 2-sigma intervals) of MOI001 without offset (left) and with a generic offset of 30 ± 20 RC years (right).**

Although the precise geographic trajectory of the immediate origins and transmission roots of the Great Northern War plague are yet to be studied in meticulous detail, two possible scenarios can be discerned. One is that the Great Northern War plague was associated with a wave that commenced somewhere in the Ottoman Empire at the very end of the 17th century. Plague was attested all over Anatolia in 1697–1699 (Kimya 2021, 2391–2393), different parts of Greece in 1697–1702 (Kanold 1721, 63; Mertzios 1969-1970, 415–416; Kolia 1990, 225–226), Smyrna/Izmir in 1697–1698 (*The Levant Voyage of the Blackham Galley (1696–1698): The Sea Journal of John Looker, Ship's Surgeon* 2022, 32–33, 91, 98°99, 117, 164; Heywood 2007), and Constantinople/Istanbul in 1698–1701 (Kanold 1721, 63; Süreyya 1890, Vol. 1, 146). By Autumn 1700, the plague was present in Adrianople/Edirne, western Bulgaria, as well as eastern and central Serbia (Manolova-Nikolova 2004, 63), before spreading northwards into eastern Hungary (Magyary-Kossa 1940, 4–6) and Carpathian Ruthenia (Mytsyuk 1938, 24–25) in 1700–1701 and emerging some two years later in south-eastern Poland (the Subcarpathia and Lublin regions).
Another scenario is that the wave commenced in Romanian territories. It is indeed reported in Transylvania in 1698 (Petresco 1933, 8–9), before spreading into eastern Hungary and further north into Carpathian Ruthenia in 1700–1701 (Magyary-Kossa 1940, 4–6; Mytsyuk 1938, 24–25). In either scenario, it is most likely that the plague wave reached south-eastern Poland in 1703 from Carpathian Ruthenia. Between 1703 and 1706, plague is attested all over south-eastern regions (voivodeships) of Subcarpathia (Podkarpackie) and Lublin (Lubelskie) (Motylewicz 1993, 16).

The post-1702 spread of the plague in eastern-central and northern Europe wave went hand-in-hand with military campaigns of the Great Northern War (1700–1721) between Sweden led by Charles XII and Russia led by Peter the Great, and their respective allies. The perpetuated story of an alleged plague outbreak in a Swedish lazaret at Pińczów (south-eastern Poland), in the aftermath of the Battle of Kliszów (19 July 1702), as the beginning of the Great Northern Wares plague may be dismissed now, as it was not until 1706–1707 that the plague reached the Pińczów region and the 1702 outbreak at the local lazaret may have been caused by a different disease (Slavin 2022, 358). Rather, it was from the Subcarpathia and Lublin regions that the plague spread into the west Ukrainian regions of Volhynia, Galicia and Podolia in 1704 (Pękacka-Falkowska 2019, 29–30), reaching Lviv/Lwow in 1704, where about 10,000 were reported to have died in the course of 1704–1705 (Charewiczowa 1930, 67). In 1706–1707 the disease was ravaging southern Poland, arriving in Krakow in late 1706/early 1707 (claiming the lives of about 20,000 residents between 1707 and 1709), and then spreading into central Poland, attacking major cities, such as Warsaw and Poznań (killing, respectively, 30,000 and 9,000 citizens) between 1707 and 1710 (Giedroyć 1899, 58–61; Karpiński 2000, 316; Burchardt, Meissner, and Burchardt 2009; Karpacz 2012; Pękacka-Falkowska 2019).

In 1708–1709, the plague spread northwards to the Baltic coast – Pomerania in the west and Lithuania in the east. In late Autumn 1708, the plague arrived in Gdańsk/Danzig, but it was not until one year later that it appeared in the nearby Elbląg/Elbing, as well as in Szczecin/Stettin, another major Baltic port (Frandsen 2010, 19–31; Zapnik 2007, 35–41). Further west, the authorities of Stralsund managed to keep the plague away until later summer 1710 (Frandsen 2010, 59–166; Zapnik 2007, 35–41), while Hamburg and Bremen were spared until 1712; remarkably, Lübeck was spared altogether, thanks to its quarantine measures (Boyens 2004; Frandsen 2010, 475–488).

In the east, the plague arrived in summer 1709 in Vilnius/Wilno and Königsberg. In the former, 30,000 Christians and 3,700 Jews were buried between July 1709 and summer 1710, while in the latter the disease was present until early 1710 (Zagorskii 1897, 130–133; Frandsen 2010, 20, 33–38). Concurrently, the epidemic spread northwards in the course of 1710, arriving in Riga in May, in Narva and Pärnu/Pernau in June and in Tallinn/Reval in August (Frandsen 2010, 41–64; Zapnik 2007, 46–57). In all instances, the disease remained for a few months, receding by the end of the year. Also, at some point in summer 1710, the plague spread into Pskov and Novgorod regions, but not carrying on further east into central Russia. Conversely, it did continue spreading further west and north, first into Stockholm (from Pärnu/Pernau in late June 1710) and then into Helsinki/Helsingfors (from Tallinn/Reval in September 1710; Frandsen 2010, 65–69). In both Finland and Sweden, plague remained confined to the southern regions. In Denmark, the plague arrived in Helsingør/Elsinore in late 1710, before arriving in other southern parts of the country (Zealand and Hovedstaden) in early 1711. Because of efficient governmental measures, which included quarantine and tight entry control, the epidemic disappeared by mid-Autumn 1711 (Frandsen 2010, 131–472).

Important for our study is the context of the Tallinn/Reval outbreak. On the eve of the plague's arrival (early August 1710), the city, with 20,000 residents and a Swedish garrison, was besieged by a force of 5,000 Russian soldiers. The former were reportedly utilizing antisanitary practices, such as polluting local landscapes and water sources with plague corpses. By September, the situation was alarming, with 50–60 Swedish soldiers dying each day, prompting Swedish commanders to surrender the city to the Russians and leave it by the end of the month. The outbreak was both intense and short: most of the deaths occurred in September and October, after which the disease drastically receded, disappearing altogether in early December. Over the period of just four months, Tallinn lost about 5,700 people, amounting to just under 30 per cent of the total population on the eve of the outbreak (Frandsen 2010, 60–64; Zapnik 2007, 46–57).

#### 11.14 The genomes of the Great Plague of Marseille (OBS); CHE1 and Rostov2033

The so far youngest clade of Second Pandemic genomes, emerging from node N19, comprises of genomes of the Great Plague of Marseille 1720–1722 (OBS cluster) as well as the genomes of Maist (CHE1), on the Chechen-Georgian border, and Rostov-on-Don (Rostov2033) in South Russia, which share a short branch of one SNP (see Fig. 2A). Both CHE1 and Rostov2033 are derived from non-UDG libraries, but due to comparatively high coverages, their terminal branch lengths can be determined confidently. CHE1, the terminal branch length is 1 SNP after filtering; for Rostov2033, 2 private SNPs passed the filtering (in both cases no SNPs passed the filtering and were classified as potential damage).
With a genetic distance of 83–84 SNPs from the European Black Death cluster, the OBS genomes are so far the most derived ancient genomes of the Second Pandemic published so far – although they might not be the youngest (see below). As described in the original publication (Bos et al. 2016), the genomes derive from mass graves associated with the Great Plague of Marseille 1720–1722, meaning they are securely dated to those years. However, the genomes were not fully embedded into their historical context: while the authors interpreted the phylogenetic position of the OBS cluster – at the time of publication the only representatives of the Branch 1A – as evidence for long-term persistence of plague in a reservoir in Europe or western Asia, the origins of the Great Plague of Marseille can be reconstructed relatively precisely.

Leaving Marseille in July 1719, Levant-bound *Grand Saint-Antoine* ship arrived in Smyrna/Izmir in late August 1719. From Smyrna/Izmir, the ship resumed its voyage in October 1719, calling at Muskonisia/Mosconissy (port in the Gulf of Edremit), Cyprus and arriving in Sidon in December 1719. The ship left Sidon on 31 January 1720, first sailing southwards to Tyre, where it picked up passengers and merchandise (including cotton bales) on 5 February, and then returning northwards to Tripoli (Lebanon). After a two-month stopover, *Grand Saint-Antoine* left Tripoli on 3 April with 15 new passengers, of whom one was infected with plague. The passenger in question died two days later, and his body was cast to the sea, in order to avoid the spread of the contagion. Hereafter, the ship arrived in Cyprus and, having been laden with more merchandise and passengers, left back for Marseille on 18 April, with a health certificate. Few days later, seven sailors and the ship's surgeon succumbed to the disease and died. Meanwhile, the rumors of a plague-infected ship were being spread across Mediterranean ports: thus, on 4 May, Maltese Hospitallers anchoring in Alicante informed locals about *Grand Saint-Antoine*. When the ship was approaching the port of Livorno on 17 May, local authorities did not allow it to enter and anchor there, on account of the onboard epidemic. The infected ship carried on westwards, stopping at Toulon (20 May), Le Bruce (21 May), and arriving at its final destination on 25 May (Restifo 2005, 13; Devaux 2013; Signoli and Tzortzis 2018; Signoli 2022). The ensuing epidemic in Marseille, its hinterland and the wider region of Provence in 1720–1722 could theoretically have commenced with a direct contact of Marseille port workers with both infected passengers and crew members, and with infected cotton bales laden in Tyre on 5 February (Slavin 2022, 354–355). Importantly, although plague was not present in Smyrna/Izmir in 1719–1720, it was attested in both Cyprus and Lebanese ports in those years (Panzac 1973; Panzac 1985, 31, 606).

Of importance is the fact that there are 10 SNPs between N21, on which most of the OBS genomes are positioned, and N17, representing the MCRA of both the Marseille and the Great Northern Wars plague genomes (with N18, on which the PRN001 genome is situated, is only one SNP away from N17). Given only 10 years separating between the two outbreaks – implying that N17 may have emerged between c.1700 and 1709 – we may infer a very fast substitution rate between N17 and N21: around one SNP every 1–2 years.

Regarding the phylogenetic position of the other two genomes, only one private SNP of CHE1 and two private SNPs of Rostov2033 passed the SNP filtering, suggesting rather short terminal branches and a shorter genetic distance from the MRCA (N19) with the OBS cluster (CHE1: d=2; Rostov2033: d=3; OBS: d≤4). In the original paper, CHE1 is given a date of 1720+ (Guellil et al. 2020 Fig. 3) since CHE1 is directly derived from the OBS cluster (1720–1722) in their phylogenetic tree. In the supplementary text of Guellil et al.'s study, the site is only roughly dated to the 16th–18th centuries. As described above, this phylogenetic position could not be confirmed in our reanalysis, questioning also the given date of 1720 or younger. Nevertheless, the analysis of the other genomes of this clade and the closely related genomes of the Great Northern War plague (PEB10 and PRN001), yields a 18th-century dating as a plausible one.

For the genome of Rostov2033, the authors of the original study (Morozova et al. 2020) stated that its associated specimen came from St Dmitrii of Rostov Fortress cemetery, specifically from a "section of the burials" that "was dated more precisely to 1762–1773 AD", where "according to historical documents, the victims of plague ca 1762–1773 were buried". Also, they stated that "[d]ocumentary sources provided a means to connect at least a few of the burials with plague epidemics in Rostov-on-Don in the winter of 1771 (Elena Batieva, personal communication)". However, no conclusive data (such as numismatic or textual evidence) was given in the supplementary text to substantiate this argument. Furthermore, the authors did not specify the exact location of the burials within the St Dmitrii of Rostov Fortress complex.

Although no comprehensive archaeological report of the excavations of the Rostov-on-Don cemetery, conducted between 1998 and 2002, has been published, these have been mentioned in several ensuing publications. According to these, the cemetery was situated actually outside of the fortress itself - about 1 km south-west of its north-western tower, on the current intersection of Gazetny Lane and M. Gorky St. (coordiates: 47.2266, 39.7136. Also, these publications indicate that some of these burials in the excavated sections contained coins, the youngest of which date to the year 1771 (Rogudeyev 2007, 73–77;

Dedyul'kin 2010, 52; Boiko and Dedyul'kin 2013, 34). In light of this, we may indeed associate the Rostov2033 genome with the 1771–1772 outbreak in Rostov-on-Don.

The 1771–1772 outbreak in Rostov-on-Don and its region appears to have been associated with a wave that had been ravaging Anatolia and the Balkans continuously in the 1760s. As the available textual evidence reveals, it may have entered South Russian and South Ukrainian territories via Wallachia and Moldova in the west and Crimea in the east, arriving on the  Black Sea and Azov Sea coasts by autumn 1771 (Slavin 2022, 359).

Given the association between Rostov2033 and the 1771–1772 outbreak in Rostov-on-Don, and the fact that the approximate length of the terminal branches of both Rostov2033 and CHE1 is similar (two and one private SNP passing the SNP filtering, respectively), we may postulate that CHE1 is chronologically not far off from Rostov2033, but most likely somewhat earlier given its terminal branch is one SNP shorter than the Rostov2033 one. Despite the most likely association of Rostov2033 with the 1771–1772 outbreak, there is potential difficulty arising from the fact that there is only one SNP separating between N20 (leading to CHE1 and Rostov2033), and N19 (representing the MRCA of the CHE1/Rostov2033 genomes on the one hand, and the Marseille ones on the other). This does not square up with the highly elevated substitution rates of one SNP every 1–2 years as estimated above for the N17–N21 segment on the 1A1 branch – implying that N19 may have emerged at some point in the 1710s – perhaps in the second half of that decade. If Rostov2033 is indeed associated with the 1771–1772 outbreak, then contrary to N19–N21 segment characterized by a very fast substitution rate, the N19–N20 segment and the subsequent terminal branch of Rostov2033 (and potentially that of CHE1) may have been marked by very slow substitute rates. Three SNP separating N19 (emerging around 1715) and the Rostov genome (putatively associated with the 1771–1772 outbreak) implies the approximate substitution rate of one SNP every 19 or so years. Alternatively, it is possible that this discrepancy derives from the fact that the real numbers of private SNPs on the respective terminal branches of both CHE1 and Rostov2033 were actually higher and the applied SNP filtering is too stringent (for non-UDG data in general or the respective datasets specifically, since site-specific environmental background, sampling method and target-enrichment probes and protocols would affect the data).

Given all these uncertainties, it would be safe to assign CHE1 a rather wide chronological bracket of c.1720 – c.1770. During the forced deportation of the Chechens and Ingush in February 1944, local chronicles – both 'national' and 'family' ones – were destroyed by Soviet authorities (Gapurov and Umkhayev 2018, 10–11). As a result, there is no direct information about plague outbreaks in Chechnya in the 18th century, but these may be indirectly gleaned from the neighboring territories of Dagestan and Georgia, as well as Russian-controlled territories of North Caucasus. Taken together, we hear about outbreaks in 1717–1718, 1721–1722, 1728–1732 and 1734–1735 (all in Dagestan, with the two latter ones perhaps being a long circulating outbreak of the same wave; Lavrov 1984, 68; Manyshev 2015, 95). The 1734–1735 outbreak  appears to have spread north-west into Kabarda (1736–1737; Naloyeva 2015, 100, 118–119) and Georgia (1737–1738; Rayfield 2012, 220). In 1749–1750, plague was reported in the North Caucasus and Dagestan (Lavrov 1984, 68–69; Manyshev 2015, 97). In 1762, it ravaged the South Caucasus, spreading into Dagestan in the following year (1763; Manyshev 2015, 97). In 1770, it was attested in Georgia, spreading into Dagestan, where it was ravaging in 1771–1772, before carrying on into the North Caucasus in 1772–1773 (Lavrov 1984, 69; Rayfield 2012, 239; *Materialy Po Istorii Russko-Gruzinskikh Otnoshenii Vtoroi Poloviny XVIII Veka. Chast' III* 1988 Vol. 2, no. 359; Kotenev et al. 2016, 613).

In theory, any of these outbreaks could be associated with the CHE1 genome. At the same time, the 1771–1772 outbreak appears to be associated with the same wave responsible for the Rostov-on-Don outbreak and, thus, given that CHE1 and Rostov2033 are positioned on two separate terminal branches, rather than together on the same terminal branch, it appears to be the least likely of the suggested candidates to be associated with CHE1. Also, given the putative association of Rostov2033 with the 1771–1772 outbreak and the uncertainty of the real numbers of private SNPs on both Rostov2033 and CHE1 terminal branches, the 1717–1718 and 1721–1722 outbreaks may appear too early, given their chronological proximity to N19 (the MRCA of both Rostov2033/CHE1 and the Marseille genomes, emerging around 1715). Hence, the 1728–1732, 1734–1735, 1749–1750 and 1763 outbreaks appear to be the best candidates to be associated with the CHE1 genome.

Because of the paucity of the available textual sources, the exact routes of each of these outbreaks spreading across the Caucasus cannot, at this point, be reconstructed. At the same time, it appears that each of these have originated in the Ottoman territories, getting imported either by land via either Anatolia or Iran, or via the Black Sea and then via southbound inland routes. For instance, the 1717–1718, 1721–1722, 1749–1750 and 1762 outbreaks were preceded by those in West Anatolia and around Constantinople (Panzac 1985, 605–607); the 1728–1732 wave was brought from North-West Iran (Manyshev 2015, 95), while the 1771–1772 one appears to be associated with the same wave that was ravaging Anatolia and the Balkans continuously in the 1760s, and that eventually spread to Rostov-on-Don in Autumn 1771.

### 12 Historical plague reservoir theories

*Philip Slavin*

The geographic origins of historical plague outbreaks and, by extension, the whereabouts of their reservoirs have been a topic of long-lasting historical and scientific controversy. For the most part, the question focused specifically on the origins of the Black Death – the first wave of the Second Plague Pandemic. Without exception, historians and scientists alike placed these in different regions of Asia, specifically within today's borders of China (including the Qinghai-Tibet plateau and the Yunnan/Burma borderland), Central Asia (including, more specifically, the Tian Shan region), North Iraq/East Anatolia, the Pontic-Caspian region, the Volga, the Caucasus, the West Urals, Western Siberia, the Mongolian-Manchurian steppe/Gobi Desert, and India (Spyrou et al. 2022 SI 1). More recently, an aDNA analysis of three specimens from a Christian cemetery in North Kyrgyzstan, all precisely dated to 1338–1339, confirmed the Tian Shan region, with its highland marmot reservoirs, as the most likely original home of the Black Death (Spyrou et al. 2022 SI 1).

Conversely, the origins of other historical plague outbreaks, associated with both the Second Pandemic and other pandemics, have been studied much less. Here, one can, in a rough manner, distinguish between two main groups of researchers. One group rejects the possibility of the presence of a lasting European reservoir and maintains, on the basis of epidemiological modelling, that recurrent plague waves originated in their Asian reservoir(s) and would periodically get reintroduced into Europe, the Middle East, and North Africa (Bramanti et al. 2021), with one study attempting to correlate post-Black Death reintroductions with climate conditions in Central Asia (Schmid et al. 2015). Some of these studies suggested different locations for reservoirs outside of Europe, including Central Asia (Schmid et al. 2015), regions to the North-West of the Caspian Sea (Namouchi et al. 2018), Western Asia, the Black Sea region and Caucasus (Guellil et al. 2020). One recent study, while not denying the possibility of the existence of temporary (short- or medium-term) European reservoirs, argued that local soil and climatic conditions would not encourage the formation and maintenance of such a long-term reservoir – with the possible exception of West Mediterranean and Aegean parts of Europe (Stenseth et al. 2022). While some historians, too, reject the presence of European reservoir(s) and argue for the continuous reintroductions of plague from the East, there are some thinking to the contrary. Thus, several Soviet-era plague biologists hypothesised the existence of native reservoirs in southern Russia, roughly around the Black Sea region (Mironov 1958; Fyodorov 1960; Kalabukhov 1961), and this hypothesis was accepted by John Alexander, a historian of plague in early-modern Russia (Alexander 1980). In a similar vein, M. B. Suponitskii conjectured about multiple rodent reservoirs within what he referred to as the 'Great Eurasian plague fracture' and suggested that these native rodent reservoirs migrated in the aftermath of the Black Death, to form several new foci in southern and eastern Europe (Suponitskii 2004, 2005; Suponitskii and Suponitskaya 2006). Suponitskii's hypothesis was adopted by Timur Khaydarov, a historian of plague in the Golden Horde and late-medieval Russian principalities (Khaydarov 2018, 74, 82–85, 87, 162, 175, 214, 233). In his study of plague in eighteenth-century Ottoman Empire, Daniel Panzac proposed that the disease was radiating from multiple foci, situated in European, Asian and North African territories of the empire. As far as European territories are concerned, he suggested Moldova-Wallachia and South-West Balkans as the most likely candidates (Panzac 1985, 109–116). In several papers, Monica Green hypothesized that Hülegü Qan's campaigns in the 1250s led to the seeding of a reservoir somewhere in the Caucasus-Volga region of the Golden Horde khanate, and it was the same reservoir that initiated both the Black Death and the *pestis secunda* waves (Green 2020; Fancy and Green 2021).

Similarly, some researchers attempted to locate historical plague reservoirs in the Mediterranean regions of Europe. Kyle Harper proposed that some early-medieval waves may have been radiated out of a reservoir somewhere in Iberia (Harper 2017). Most recently, the same author suggested that during both the First and Second Plague pandemics, plague seeded local reservoirs "somewhere in the circum-Mediterranean" (Harper 2023). As we have seen, Stenseth et al. argued that of all European regions, the Mediterranean and Aegean zones were the most likely to have had plague reservoirs, because of their appropriate soil structure and climatic conditions (Stenseth et al. 2022). In the same vein, Peter Topping suggested that since the arrival of the Black Death in the Peloponnese, plague remained endemic there until the 18th century (Topping 1977). Also, as we have seen, in the original publication of the Marseille genomes (OBS), Bos et al. interpreted the phylogenetic position of their cluster – at that time the only representatives of the Branch 1A – as evidence for long-term persistence of plague in a reservoir in Europe or western Asia (Bos et al. 2016). This suggestion has been adopted by Nükhet Varlık, who stated that the 1720-1722 outbreak in Marseille originated "in or around Marseille itself, rather than from the eastern Mediterranean" (Varlık 2020). This interpretation is refuted by the reconstruction of the spatio-temporal origins of the Great Plague of Marseille in Section 11.14 above (see also Slavin 2022).

The possibility of a native European reservoir existing further North has, too, been advanced by several historians. In 1976, Russell advanced a hypothesis that during the First Plague Pandemic, there were plague reservoirs in the British Isles (Russell 1976). In his study of plague outbreaks in fifteenth-century England, Robert Gottfried argued for the existence of a plague reservoir in East Anglia (Gottfried 1978). This East Anglian reservoir hypothesis has been adopted and expanded, respectively, upon by Bolton 2013 and Pribyl 2017. Similarly, Paul Slack suggested that plague may have been endemic in major English towns of the 16th and 17th centuries, but its perseverance depended on constant importation of more virulent strains from outside - more specifically, from eastern Mediterranean (Slack 1985). Although for a much later period, Bramanti et al. hypothesized that the same region hosted a short-term reservoir during the Third Pandemic outbreaks there (1906–1918; Bramanti et al. 2019). In her paper on Swiss outbreaks in the 1560s, Ann Carmichael suggested that a native European reservoir may have once existed in marmot colonies of Swiss Alps (Carmichael 2014). In his study of the origins and spread of the *pestis secunda*, Philip Slavin has shown empirically that the same wave appears to have commenced in a South-Central German reservoir, possibly in South Hesse region, and hypothesized that the same reservoir may have been responsible for at least some recurrent late-medieval and early modern plague waves (Slavin 2021, 2022). Slavin's hypothesis regarding the South-Central German origins of the *pestis secunda* has been confirmed by Parker et al.'s aDNA analysis of five specimens from Krakauer Berg (near Halle), positioning three out of five specimens basal on the *pestis secunda* branch (Branch 1B; Parker et al. 2023). The present paper takes up and expands upon the South German hypothesis and through a detailed spatio-temporal reconstruction of each wave, associated with each of the newly sequenced genome, lends support to the idea of potentially a persistent long-term plague reservoir in that region.

A close analysis of the spatio-temporal context of post-Little Polytomy genomes led Slavin to hypothesize that while 1A2 genomes may have originated in a Central European reservoir (potentially, in the same or nearby reservoir in South Germany responsible for recurrent waves beginning with the *pestis secunda*), the 1A1 ones appear to be imported into Europe from Ottoman sources, seemingly outside of Europe (Slavin 2022). This hypothesis has been reiterated in the present paper. Importantly, the existence of putative Ottoman reservoirs in Asia and North Africa has already been suggested by several historians. Thus, Daniel Panzac, in addition to hypothesizing about South-East European reservoirs, mentioned above, postulated the existence about plague foci in the Istanbul hinterland, East Anatolia, Kurdistan, the ‘Asir Region (the Saudi-Yemeni border), and Egypt (Panzac 1985, 105–133). In the same vein, Paul Slack suggested that in the 16th and 17th century, plague was being repeatedly reintroduced into Europe from its original home in the eastern Mediterranean (Slack 1985). Similarly, Nükhet Varlık suggested the existence of an East Anatolian reservoir (Varlık 2022), while Timur Khaydarov hypothesized that the same reservoir may have been situated, more specifically, in the Armenian Highlands (around Van Lake) (Khaydarov 2021). For the First Plague Pandemic, Tsiamis hypothesized about a putative reservoir in Byzantine and early Islamic Middle East (Tsiamis 2010).

### 13 Archival sources

The National Records of Scotland, Edinburgh: E71/1/6-10; E71/2/1; E71/5/1; E71/7/1-2, 4; E71/17/1-2; E71/21/2; E71/23/1; E71/25/1

Tallinna linnaarhiiv (Tallinn City Archives): F190, N1, S60, fol. 28v; F191, N2, S1, fol. 42; F191, N2, S1, fols. 31–31a; Begravna i St.-Olai kyrkan i Reval, 1603–1821: 41–42

University of Cambridge, Trinity College Archives: O.13, Vol. XIX, fols. 143v, 177r, 208r; Vol. XX, fols. 59v, 149r, 212v; Vol. XXI, fol. 68v; Vol. XXII, fols. 37v, 133v; Vol. XXIII, fol. 168v

### 14 References

Aalto, Ilari. 2017. "Kulkutaudit Keskiajan Suomessa." *Arkeologia Nyt!*, 13–17.

Abduzhemilev, Refat. 2016. *Khronika Mekhmeda Denai Kak Pomyatnik Krymskotatarskoi Khudozhestvennoi Literatury XIII v*. Kazan: Institut Istorii im. Sh. Mardzhani Akademii Nauk Respubliki Tatarstan.

Abel, Otto, ed. 1852. "Chronicon Elwacense." In *Monumenta Germaniae Historica, Scriptores 10*, 34–51. Hannover: Impensis Bibliopolii Aulici Hahniani.

Achtman, Mark, Giovanna Morelli, Peixuan Zhu, Thierry Wirth, Ines Diehl, Barica Kusecek, Amy J. Vogler, et al. 2004. "Microevolution and History of the Plague Bacillus, Yersinia Pestis." *Proceedings of the National Academy of Sciences of the United States of America* 101 (51): 17837–42.

Adjemian, Jennifer Zipser, Patrick Foley, Kenneth L. Gage, and Janet E. Foley. 2007. "Initiation and Spread of Traveling Waves of Plague, Yersinia Pestis, in the Western United States." *The American Journal of Tropical Medicine and Hygiene* 76 (2): 365–75.

Alexander, John T. 1980. *Bubonic Plague in Early Modern Russia: Public Health and Urban Disaster*. Baltimore: Johns Hopkins University Press.

Andersson, Jourdan A., Jian Sha, Tatiana E. Erova, Eric C. Fitts, Duraisamy Ponnusamy, Elena V. Kozlova, Michelle L. Kirtley, and Ashok K. Chopra. 2017. "Identification of New Virulence Factors and Vaccine Candidates for Yersinia Pestis." *Frontiers in Cellular and Infection Microbiology* 7 (October): 448.

Andrades Valtueña, Aida, Alissa Mittnik, Felix M. Key, Wolfgang Haak, Raili Allmäe, Andrej Belinskij, Mantas Daubaras, et al. 2017. "The Stone Age Plague and Its Persistence in Eurasia." *Current Biology: CB* 27 (23): 3683–91.e8.

Andrades Valtueña, Aida, Gunnar U. Neumann, Maria A. Spyrou, Lyazzat Musralina, Franziska Aron, Arman Beisenov, Andrey B. Belinskiy, et al. 2022. "Stone Age Yersinia Pestis Genomes Shed Light on the Early Evolution, Diversity, and Ecology of Plague." *Proceedings of the National Academy of Sciences of the United States of America* 119 (17): e2116722119.

Andreescu, Ştefan. 1977. "L'Action de Vlad Ţepeș Dans Le Sud-Est de l'Europe En 1476." *Revue Des Etudes Sud-Est Europeennes* 15 (2): 259–72.

Archäologischer Dienst Graubünden, Lorena Burkhardt (Ed.). 2020. *Domat/Ems, Sogn Pieder: vom frühmittelalterlichen Herrenhof zum neuzeitlichen Pestfriedhof*. Chur: Somedia-Buchverlag.

Atkinson, Thomas Dinham. 1933. *Architectural History of the Benedictine Monastery of St Etheldreda at Ely*. Cambridge: Cambridge University Press.

Auerbach, Raymond K., Apichai Tuanyok, William S. Probert, Leo Kenefic, Amy J. Vogler, David C. Bruce, Christine Munk, et al. 2007. "Yersinia Pestis Evolution on a Small

Timescale: Comparison of Whole Genome Sequences from North America.” *PloS One* 2 (8): e770.
Ayalon, Yaron. 2015. *Natural Disasters in the Ottoman Empire: Plague, Famine, and Other Misfortunes*. Cambridge: Cambridge University Press.
Baczkowski, C., and Danuta Turkowska, eds. 2000. *Joannis Dlugossii Annales Seu Cronicae Incliti Regni Poloniae. Liber 11, 1413-1430*. Warsaw: Wydaw. Naukowe PWN.
Baetsen, W. A., and S. Baetsen. 2020. “Fysisch-Antropologisch Onderzoek.” In *Wat de Nieuwe Sint Jansbeek Boven Water Bracht: Dood En Leven in Het Arnhemse Verleden; Archeologisch Onderzoek Sint Jansbeek Te Arnhem – Deel 2: Menselijk Botmateriaal, Bioarcheologisch Onderzoek En Grafgebruiken. RAAP Report 4476-2.*, edited by W. A. Baetsen and G. Zielman, 355–598. Weesp: RAAP Archeologisch Adviesbureau B.V.
Baetsen, W. A., and C. L. Scheib. 2020. “DNA-Onderzoek.” In *Wat de Nieuwe Sint Jansbeek Boven Water Bracht: Dood En Leven in Het Arnhemse Verleden; Archeologisch Onderzoek Sint Jansbeek Te Arnhem – Deel 2: Menselijk Botmateriaal, Bioarcheologisch Onderzoek En Grafgebruiken. RAAP Report 4476-2.*, edited by W. A. Baetsen and G. Zielman, 600–611. Weesp: RAAP Archeologisch Adviesbureau B.V.
Baetsen, W. A., and G. Zielman, eds. 2020. *Wat de Nieuwe Sint Jansbeek Boven Water Bracht: Dood En Leven in Het Arnhemse Verleden; Archeologisch Onderzoek Sint Jansbeek Te Arnhem – Deel 2: Menselijk Botmateriaal, Bioarcheologisch Onderzoek En Grafgebruiken. RAAP Report 4476-2*. Weesp: RAAP Archeologisch Adviesbureau B.V.
Baltzarek, Franz, and Johanne Pradel. 1973. *Dir Städte Vorarlbergs*. Vienna: Brüder Hollinek.
Bazala, Vladimir. 1962. “Calendarium Pestis (II).” *Acta Historica Medicinae, Pharmaciae, Veterinae* 2: 51–61, 72–87.
Benedict, Carol. 1996. “Bubonic Plague in Nineteenth-Century China.” In . Stanford: Stanford University Press.
Berning, A. 1987-9. “Die Pest in Kaufbeuren.” *Kaufbeurer Geschichtsblätter* 11: 106–11.
*Beschreibung Des Oberamts Schorndorf*. 1851. Stuttgart: Müller’s Verlagshandlung.
Bethmann, L. C., ed. 1866. “Annales Reatini.” In *Monumenta Germaniae Historica, Scriptores 19*, 267–68. Hannover: Impensis Bibliopolii Hahniani.
Białecki, Tadeusz, and Edward Rymar, eds. 2005. *Thomas Kantzow, Pomerania. Kronika Pomorska Z XVI Wieku*. Szczecin: Uniwersytet Szczeciński.
Binder, Paul. 1983. “Epidemiile de Ciumă Din Transilvania Secolului Al XVI-Lea (1511-1603).” In *Momente Din Trecutul Medicinii: Studii, Note şi Documente*, edited by Gheorghe V. Brătescu, 99–111. Bucharest: Editura Medicală.
Biraben, Jean-Noël. 1975. *Les Hommes et La Peste En France et Dans Les Pays Européens et Méditerranéens*. Paris: Mouton.
Bisgaard, Lars. 2009. “Danish Plague Patterns, 1360-1500.” In *Living with the Black Death*, edited by Lars Bisgaard and Leif Søndergaard, 85–111, Tables. Odense: University Press of Southern Denmark.
Boiko, A. L., and A. V. Dedyul’kin. 2013. “Kreposti I Poseleniya XVII-XIX Vv. (p. 34-6).” In *Arkheologiya Nizhnego Dona*, edited by A. V. Kiyashko, Yuzhnyi Federal’nyi Universitet:34–36. Rostov-on-Don.
Bolton, J. L. 2013. “Looking for Yersinia Pestis: Scientists, Historians and the Black Death.” In *The Fifteenth Century XII: Society in an Age of Plague*, edited by L. Clark and C. Rawcliffe, 15–38. Woodbridge: Boydell & Brewer.
Borisenkov, Ye P., and V. M. Pasetskii. 1983. *Ekstremalnye Prirodnye Yavleniya v Russkikh Letopisyakh XI-XVII vv*. Leningrad: Gidrometeoizdat.
Bos, Kirsten I., Alexander Herbig, J. Sahl, N. Waglechner, M. Fourment, S. A. Forrest, J. Klunk, et al. 2016. “Eighteenth Century Yersinia Pestis Genomes Reveal the Long-Term Persistence of an Historical Plague Focus.” *eLife* 5: e12994.
Bos, Kirsten I., Kelly M. Harkins, Alexander Herbig, Mireia Coscolla, Nico Weber, Iñaki Comas, Stephen A. Forrest, et al. 2014. “Pre-Columbian Mycobacterial Genomes Reveal Seals as a Source of New World Human Tuberculosis.” *Nature* 514 (7523): 494–97.

Bos, Kirsten I., Verena J. Schuenemann, G. Brian Golding, Hernán A. Burbano, Nicholas Waglechner, Brian K. Coombes, Joseph B. McPhee, et al. 2011. “A Draft Genome of Yersinia Pestis from Victims of the Black Death.” *Nature* 478 (7370): 506–10.
Boubaker, Sadok. 1995. “La Peste Dans Les Pays Du Maghreb: Attitudes Face Au Fléau et Impacts Sur Les Activités Commerciales (XVIème-XVIIIème Siècles).” *Revue D’histoire Maghrébine* 79-80: 311–41.
Boyens, Kathrin. 2004. “Die Krise in Der Krise. Die Masnahmen Hamburgs Während Der Letzten Pest 1712–1714.” In *Die Leidige Seuche: Pest-Fälle in Der Frühen Neuzeit*, edited by Otto Ulbricht. Cologne: Böhlau.
Bramanti, Barbara, Katharine R. Dean, Lars Walløe, and Nils Chr Stenseth. 2019. “The Third Plague Pandemic in Europe.” *Proceedings of the Royal Society B: Biological Sciences* 286 (1901): 20182429–20182429.
Bramanti, Barbara, Yarong Wu, Ruifu Yang, Yujun Cui, and Nils Chr Stenseth. 2021. “Assessing the Origins of the European Plagues Following the Black Death: A Synthesis of Genomic, Historical, and Ecological Information.” *Proceedings of the National Academy of Sciences of the United States of America* 118 (36). https://doi.org/10.1073/pnas.2101940118.
Brázdil, Rudolf, Ladislava Řezníčková, Hubert Valášek, Andrea Kiss, and Oldřich Kotyza. 2013. “Past Locust Outbreaks in the Czech Lands: Do They Indicate Particular Climatic Patterns?” *Theoretical and Applied Climatology* 116. https://doi.org/10.1007/s00704-013-0950-9.
Buganov, V. I., and B. A. Rybakov, eds. 2002. *Novgorodskaya Karamzinskaya Letopis’*. Polnoye Sobraniye Russkikh Letopisei 42. St Petersburg: Dmitrii Bulanin.
Bunge, Friedrich Georg von, ed. 1857. *Liv-, Est- Und Kurländisches Urkundenbuch, Band 3. 1368-1393*. Reval: Heinrich Laakmann.
Büntgen, Ulf, Willy Tegel, Kurt Nicolussi, Michael McCormick, David Frank, Valerie Trouet, Jed Kaplan, et al. 2011. “2500 Years of European Climate Variability and Human Susceptibility.” *Science* 331: 578–82.
Büntgen, Ulf, Otmar Urban, Paul Krusic, Michal Rybníček, Tomas Kolar, Tomáš Kyncl, Alexander Ač, et al. 2021. “Recent European Drought Extremes beyond Common Era Background Variability.” *Nature Geoscience* 14: 1–7.
Burchardt, Jarosław, Roman K. Meissner, and Dorota Burchardt. 2009. “Oddech śmierci Zaraza Dżumy W Wielkopolsce I W Poznaniu W Pierwszej Połowie XVIII Wieku.” *Nowiny Lekarskie: Organ Wydzialu Lekarskiego Towarzystwa Przyjaciol Nauk Posnanskiego* 78 (1): 79–84.
Burton, John Hill, ed. 1878. *The Register of the Privy Council of Scotland, Vol. 2, A.D. 1569-1578*. Edinburgh: H.M. General Getister House.
Bychkov, A. F., and K. N. Bestuzhev-Ryumin, eds. 1889. *Letopisnyi Sbornik, Imenuemyi Letopis’yu Avraamki*. Polnoye Sobraniye Russkikh Letopisei 16. St Petersburg.
*Calendar of State Papers, Foreign: Elizabeth. Volume 19, August 1584-August 1585*. 1916. London: Longman, Roberts, & Green.
Camenisch, Chantal. 2015. *Endlose Kälte. Witterungsverlauf Und Getreidepreise in Den Burgundischen Niederlanden Im 15. Jahrhundert*. Basel: Schwabe AG.
Camenisch, Chantal, Rudolf Brázdil, Andrea Kiss, Christian Pfister, Oliver Wetter, Christian Rohr, Antonio Contino, and Dag Retsö. 2020. “Extreme Heat and Drought in 1473 and Their Impacts in Europe in the Context of the Early 1470s.” *Regional Environmental Change* 20. https://doi.org/10.1007/s10113-020-01601-0.
Carmichael, Ann G. 2014. “Plague Persistence in Western Europe: A Hypothesis.” In , 157–92. Kalamazoo, Bradford.
Cassese, L. 1941. “La «Chronica Civitatis Aquile» Di Alessandro de Ritiis.” *Archivio Storico per Le Province Napoletane N.s.* 27: 151–261.
Cessford, Craig, Christiana L. Scheib, Meriam Guellil, Marcel Keller, Craig Alexander, Sarah A. Inskip, and John E. Robb. 2021. “Beyond Plague Pits: Using Genetics to Identify Responses to Plague in Medieval Cambridgeshire.” *European Journal of Archaeology*, 1–23.

Chain, Patrick S. G., Ping Hu, Stephanie A. Malfatti, Lyndsay Radnedge, Frank Larimer, Lisa M. Vergez, Patricia Worsham, May C. Chu, and Gary L. Andersen. 2006. “Complete Genome Sequence of Yersinia Pestis Strains Antiqua and Nepal516: Evidence of Gene Reduction in an Emerging Pathogen.” *Journal of Bacteriology* 188 (12): 4453–63.
Chain, P. S. G., E. Carniel, F. W. Larimer, J. Lamerdin, P. O. Stoutland, W. M. Regala, A. M. Georgescu, et al. 2004. “Insights into the Evolution of Yersinia Pestis through Whole-Genome Comparison with Yersinia Pseudotuberculosis.” *Proceedings of the National Academy of Sciences of the United States of America* 101 (38): 13826–31.
Charewiczowa, Łucja. 1930. *Klęski Zaraz W Dawnym Lwowie*. Lwów: Książnica-Atlas.
Christensen, Peter. 2003. “‘In These Perilous Times’: Plague and Plague Policies in Early Modem Denmark.” *Medical History* 47: 413–50.
“Chroniken von Closener Und Koeningshoven.” 1845. In *Code Historique et Diplomatique de La Ville de Strasbourg*. Vol. 1. Strasbourg: G. Silbermann.
Clarke, Jack Alden. 1966. *Huguenot Warrior: The Life and Times of Henri de Rohan, 1579-1638*. The Hague: Springer Science.
Cobban, Alan B. 1969. *The King’s Hall Within the University of Cambridge in the Later Middle Ages*. Cambridge: Cambridge University Press.
Cohn, Samuel K. 2002. *The Black Death Transformed: Disease and Culture in Early Renaissance Europe*. London: Arnold.
Cook, Edward, Richard Seager, Yochanan Kushnir, Keith Briffa, Ulf Büntgen, David Frank, Paul Krusic, et al. 2015. “Old World Megadroughts and Pluvials during the Common Era.” *Science Advances* 1: 1–9.
Cooper, Charles Henry. 1842. *Annals of Cambridge*. Cambridge: Warwick and Co.
Cui, Yujun, Chang Yu, Haihong Han, Dongfang Li, Yanjun Li, Thibaut Jombart, Lucy Weinert, et al. 2013. “Historical Variations in Mutation Rate in an Epidemic Pathogen, Yersinia Pestis.” *Proceedings of the National Academy of Sciences of the United States of America* 110. https://doi.org/10.1073/pnas.1205750110.
Cummins, Neil, Morgan Kelly, and Cormac Ó Gráda. 2016. “Living Standards and Plague in London, 1560-1665.” *The Economic History Review* 69 (1): 3–34.
Curtis, Daniel R., and Joris Roosen. 2017. “The Sex-selective Impact of the Black Death and Recurring Plagues in the Southern Netherlands, 1349–1450.” *American Journal of Physical Anthropology* 164 (2): 246–59.
Dai, Ruixia, Baiqing Wei, Haoming Xiong, Xiaoyan Yang, Yao Peng, Jian He, Juan Jin, et al. 2018. “Human Plague Associated with Tibetan Sheep Originates in Marmots.” *PLoS Neglected Tropical Diseases* 12: e0006635.
Dalitz, Stefan, Gisela Grupe, and Bettina Jungklaus. 2012. “Das Kleinste Massengrab Brandenburgs. Drei Tote Aus Dem Dreißigjährigen Krieg Auf Der Dominsel Der Stadt Brandenburg an Der Havel.” *Jahresbericht Des Historischen Vereins Brandenburg (Havel)* 21: 60–72.
*Debar Śepatayim: An Ottoman Hebrew Chronicle from the Crimea (1683-1730). Written by Krymchak Rabbi David Lekhno*. 2021. Boston: Academic Studies Press.
Dedyul’kin, A. V. 2010. “Arkheologia Pozdnikh Periodov Istorii (XVIII Vek) Na Nizhnem Donu.” *Izvestiya Vuzov. Severo-Kavkazskii Region. Obshchestvennye Nauki*, no. 1: 51–55.
De Groot, A. H. 1978. *The Ottoman Empire and the Dutch Republic. A History of the Earliest Diplomatic Relations, 1610-1630*. Leiden: Nederlands Historisch-Archaeologisch Instituut.
Delachenal, R., ed. 1916. *Les Grandes Chroniques de France. Chronique Des Règnes de Jean II et de Charles*. Paris: La Société de l’histoire de France.
Deng, Wen, Valerie Burland, Guy Plunkett 3rd, Adam Boutin, George F. Mayhew, Paul Liss, Nicole T. Perna, et al. 2002. “Genome Sequence of Yersinia Pestis KIM.” *Journal of Bacteriology* 184 (16): 4601–11.
*Der Feind in Der Stadt. Vom Umgang Mit Seuchen in Augsburg, München Und Nürnberg. Eine Ausstellung Der Bayerischen Archivschule Der Generaldirektion Der Staatlichen Archive Bayerns*. 2016. Munich: Generaldirektion der Staatlichen Archive Bayerns.

Derkach, V. A., and S. Ye Shun'gina. 2018. "Rezul'taty Arkheologicheskoi Razvedki v Krasnogorodskom Rayone Pskovskoi Oblasti v 2016 G." In *Arkheologiya I Istoriya Pskova I Pskovskoi Zemli. Yezhegodnik Seminara Imeni Akademika V.V. Sedova. Vypusk 33. Materialy 63-Go Zasedaniya (18-20 Aprelya 2017 G.)*, edited by Lopatin, N.V. and Salmina, Ye. V., 127–31. Moscow: Institut Arkheologii RAN.
Devaux, Christian A. 2013. "Small Oversights That Led to the Great Plague of Marseille (1720–1723): Lessons from the Past." *Infection, Genetics and Evolution: Journal of Molecular Epidemiology and Evolutionary Genetics in Infectious Diseases* 14: 169–85.
*Die Chroniken Der Fränkischen Städte. Nürnberg. Erster Band*. 1862. Leipzig: S. Hirzel.
*Die Chroniken Der Fränkischen Städte. Nürnberg. Vierter Band*. 1872. Leipzig: S. Hirzel.
*Die Chroniken Der Mittelrheinischen Städte. Mainz, Zweiter Band*. 1882. Leipzig: Verlag von S. Hirzel.
*Die Chroniken Der Schwäbischen Städte. Augsburg. Dritter Band*. 1892. Leipzig: S. Hirzel.
*Die Chroniken Der Schwäbischen Städte. Augsburg. Erster Band*. 1865. Leipzig: S. Hirzel.
*Die Chroniken Der Schwäbischen Städte. Augsburg, Vierter Band*. 1894. Leipzig: S. Hirzel.
Dierauer, Johannes, ed. 1900. *Chronik Der Stadt Zürich. Mit Fortsetzungen*. Basel: Adolf Geering.
*Die Recesse Und Andere Akten Der Hansetage von 1256-1430. Band VII*. 1893. Leipzig: Duncker & Humblot.
Ditchburn, David. 1990. "Cargoes and Commodities: Aberdeen's Trade with Scandinavia and the Baltic, C.1302-1542." *Northern Studies: The Journal of the Scottish Society for Northern Studies* 27: 12–22.
"Dopolneniya K Nikonovskoi Letopisi." 1904-1906. In *Polnoye Sobraniye Russkikh Letopisei 13*, 303–408. St Petersburg: Tipografiya I.N. Skokrokhodova.
Eckert, Edward A. 1978. "Boundary Formation and Diffusion of Plague: Swiss Epidemics from 1562 to 1669." *Annales de Démographie Historique*, 49–80.
———. 1996. *The Structure of Plagues and Pestilences in Early Modern Europe: Central Europe, 1560-1640*. Basel: Karger.
Emler, Josef, ed. 1884. "Kronika Beneše Z Weitmile." In *Fontes Rerum Bohemicarum 4*, 459–548. Prague: Nákladem Nadání Františka Palackého.
Eppinger, Mark, Zhaobiao Guo, Yinong Sebastian, Yajun Song, Luther E. Lindler, Ruifu Yang, and Jacques Ravel. 2009. "Draft Genome Sequences of Yersinia Pestis Isolates from Natural Foci of Endemic Plague in China." *Journal of Bacteriology* 191 (24): 7628–29.
Epstein, M. 1908. *The Early History of the Levant Company*. London: George Routledge & Sons Limited.
Erdel, Andreas, Hans Eugen Specker, and H-J Winckelmann. 1986. "Ulm Und Die Pest. Maßnahmen Zur Verhütung Der Pest in Der Freien Reichsstadt Ulm Im 17. Und 18. Jahrhundert." *Apothekerjournal* 9: 94–97.
Eroshenko, Galina A., Nikita Yu Nosov, Yaroslav M. Krasnov, Yevgeny G. Oglodin, Lyubov M. Kukleva, Natalia P. Guseva, Alexander A. Kuznetsov, Sabyrzhan T. Abdikarimov, Aigul K. Dzhaparova, and Vladimir V. Kutyrev. 2017. "Yersinia Pestis Strains of Ancient Phylogenetic Branch 0.ANT Are Widely Spread in the High-Mountain Plague Foci of Kyrgyzstan." *PloS One* 12 (10): e0187230.
Fancy, Nahyan, and Monica H. Green. 2021. "Plague and the Fall of Baghdad (1258)." *Medical History* 65 (2): 157–77.
Farrow, M. A. 1943-1945. *Index to Wills Proved in the Consistory Court of Norwich : And Now Preserved in the District Probate Registry at Norwich*. Norwich: Norfolk Record Society.
Fellows Yates, James A., Thiseas C. Lamnidis, Maxime Borry, Aida Andrades Valtueña, Zandra Fagernäs, Stephen Clayton, Maxime U. Garcia, Judith Neukamm, and Alexander Peltzer. 2021. "Reproducible, Portable, and Efficient Ancient Genome Reconstruction with Nf-Core/eager." *PeerJ* 9 (March): e10947.
Frandsen, Karl-Erik. 2010. *The Last Plague in the Baltic Region, 1709-1713*. Copenhagen: Museum Tusculanum Press.

Fries, Lorenz. 1994. *Chronik Der Bischöfe von Würzburg 742–1495. Band II (1127-1376)*. Edited by Christoph Bauer, Udo Beireis, Thomas Heiler, Georg Salzer, and Peter Süß. Würzburg: Ferdinand Schöningh.
Fuhrmann, P. Mathia. 1738. *Alt- Und Neues Wien, Oder Dieser Kayserlich- Und Ertz-Lands-Fürstlichen Residentz-Stadt Chronologisch- Und Historische Beschreibung*. Vienna: Joh. Baptist Prasser.
Fyodorov, V. N. 1960. "The Question of the Existence of Natural Foci of Plague in Europe in the Past." *Journal of Hygiene, Epidemiology, Microbiology, and Immunology* 4: 135–41.
"GA 1551, Inv.nr. 4187-0004 (Gelders Archief 1551 Topografisch-Historische Atlas Gelderland, Inventarisnummer 4187-0004). Aernhem, 1649. Plattegrond van Arnhem Op Een Schaal van 1:3000 Door Joan Blaeu." n.d. Accessed July 6, 2021. https://www.geldersarchief.nl/bronnen/archieven?mivast=37&mizig=284&miadt=37&miaet=1&micode=1551&minr=43436320&miview=ldt.
Gahlbeck, Christian, Madlena Mahling, Klaus Neitmann, and Matthias Thumse, eds. 2020. *Liv-, Est- Und Kurländisches Urkundenbuch, Band 13. 1472-1479*. Cologne: Böhlau.
Gapp, Johann. 1986. "Die Pest Und Die Sebastianiverehrung in Telfs." *Tiroler Heimat* 61: 11–19.
Gapurov, Sh A., and Kh S. Umkhayev. 2018. "Problemy Etnogeneza I Drevnei Istorii Chechentsev." In *Etnogenez I Etnicheskaya Istoriya Narodov Kavkaza. Materialy I Mezhdunarodnogo Nakhskogo Nauchnogo Kongressa, G. Groznyi. 11-12 Sentyabrya 2018 G.*, 8–18. Groznyi: Akademiya Nauk Chechenskoi Respubliki.
Garcia, Emilio, Patrick Chain, Patricia Worsham, Scott W. Bearden, Stephanie Malfatti, Dorothy Lang, Frank Larimer, and Luther Lindler. 2007. "Pestoides F, an Atypical Yersinia Pestis Strain from the Former Soviet Union." In *The Genus Yersinia: From Genomics to Function*, edited by Robert D. Perry and Jacqueline D. Fetherston, 603:17–22. New York, NY: Springer New York.
Gardiner, J., ed. 1880. *Three Fifteenth-Century Chronicles: With Historical Memoranda by John Stowe, the Antiquary, and Contemporary Notes of Occurrences Written by Him in the Reign of Queen Elizabeth*. London: Camden Society.
Garrelt, C., and I. Wiechmann. 2003. "Detection of Yersinia Pestis DNA in Early and Late Medieval Bavarian Burials." *Documenta Archaeobiologiae* 1: 247–54.
Gerhards, Guntis. 2011. "Epidēmijas Viduslaiku Un Jauno Laiku Rīgā." *Latvijas Vēstures Institūta Žurnāls* 4: 37–65.
Giedroyć, Franciszek. 1899. *Mór W Polsce W Wiekach Ubiegłych: Zarys Historyczny*. Warsaw: L. Szkaradziński.
Giffin, Karen, Aditya Kumar Lankapalli, Susanna Sabin, Maria Spyrou, Cosimo Posth, Justina Kozakaitė, Ronny Friedrich, et al. 2020. "A Treponemal Genome from an Historic Plague Victim Supports a Recent Emergence of Yaws and Its Presence in 15 Century Europe." *Scientific Reports* 10. https://doi.org/10.1038/s41598-020-66012-x.
Glaser, Rüdiger. 2001. *Klimageschichte Mitteleuropas: 1000 Jahre Wetter, Klima, Katastrophen*. Darmstadt: Wissenschaftliche Buchgesellschaft.
Goncharova, S. M., M. Yu Goncharov, M. V. Yermolayenko, A. N. Maslovskii, and A. P. Minayev. 2014. "Arkheologicheskiye Issledovaniya v Gorode Azove v 2012 Godu." *Istoriko-Arkheologicheskiye Issledovaniya v Azove I Na Nizhnem Donu* 28: 142–62.
Gooskens, Frans. 1987. "Pestepidemieën in Breda Tijdens de Middeleeuwen (1382-1535)." *Jaarboek van de Geschied- En Oudheidkundige Kring van Stad En Land van Breda "De Oranjeboom"* 39: 18–54.
Goryushkina, L. P. 2021. "Morovye Povetriya, Karantinnyye Mery I Tserkov' v XVI Stoletii." In *Rus' Pod Udarom Morovykh Povetrii*, edited by A. V. Pimenova, 34–61. Moscow: Universitet Dmitriya Pozharskogo.
Gottfried, R. S. 1978. *Epidemic Disease in Fifteenth-Century England: The Medical Response and the Demographic Consequences*. New Brunswick, N.J.: Rutgers University Press.
Grainger, I., D. Hawkins, L. Cowal, and R. Mikulski. 2008. *The Black Death Cemetery, East Smithfield, London*. London: Museum of London Archaeology Service.

Green, Monica H. 2018. “Putting Africa on the Black Death Map: Narratives from Genetics and History.” *Afriques*, no. 09. https://doi.org/10.4000/afriques.2125.
———. 2020. “The Four Black Deaths.” *The American Historical Review* 125 (5): 1601–31.
Guellil, Meriam, Oliver Kersten, Amine Namouchi, Stefania Luciani, Isolina Marota, Caroline Arcini, Elisabeth Iregren, et al. 2020. “A Genomic and Historical Synthesis of Plague in 18th Century Eurasia.” *Proceedings of the National Academy of Sciences* 117 (45): 1–8.
Gutsmiedl-Schümann, Doris, Bernd Päffgen, Heiner Schwarzberg, Marcel Keller, Andreas Rott, and Michaela Harbeck. 2018. “Digging up the Plague: A Diachronic Comparison of aDNA Confirmed Plague Burials and Associated Burial Customs in Germany.” *Praehistorische Zeitschrift* 92: 405–27.
Harper, Kyle. 2017. *The Fate of Rome: Climate, Disease, and the End of an Empire*. Princeton: Princeton University Press.
———. 2023. “The First Plague Pandemic in Italy: The Written Evidence.” *Speculum* 98 (2): 369–420.
Hartmann, S. 1973. *Reval Im Nordischen Krieg. Quellen Und Studien Zur Baltischen Geschichte*. Bonn.
Hartmann, Stefan, ed. 1991. *Herzog Albrecht von Preußen Und Das Bistum Ermland, Regesten, Vol. 2 (1550-1568)*. Cologne: Böhlau.
———. , ed. 2005. *Herzog Albrecht von Preussen Und Livland (1551-1557): Regesten Aus Dem Herzoglichen Briefarchiv Und Den Ostpreussischen Folianten*. Cologne: Böhlau.
———. , ed. 2008. *Herzog Albrecht von Preussen Und Livland (1565-70). Regesten Aus Dem Herzoglichen Briefarchiv*. Cologne: Böhlau.
Heywood, Colin. 2007. “Sickness and Death in an Ill Climate: The Detention of the Blackham Galley at Izmir 1697-8.” In *Ottoman Izmir: Studies in Honour of Alexander H. de Groot*, edited by Maurits H. Van den Boogert, 39–52. Leiden: Nederlands Instituut voor het Nabije Oosten.
Hildebrand, Hermann, ed. 1889. *Liv-, Est- Und Kurländisches Urkundenbuch, Band 9. 1436-1443*. Riga: Heinrich Laakmann.
Horanin, Mariusz. 2019. “Die Pest in Augsburg Um 1500. Die Soziale Konstruktion Einer Krankheit.” PhD, Georg-August-Universität Göttinge.
Hrabak, Bogumil. 1989. “Talasi Kuge Na Bosanskohercegovačkom Upravnom Prostoru 1463 – 1800.” *Acta Historica Medicinae, Stomatologiae, Pharmaciae, Medicinae Veterinae* 29 (1): 19–36.
Hye, Franz-Heinz. 1980. *Die Städte Tirols, 1. Teil: Bundesland Tirol*. Vienna: Verlag der Österreichischen Akademie der Wissenschaften.
Ibs, Jürgen Hartwig. 1994. *Die Pest in Schleswig-Holstein von 1350 Bis 1547/48: Eine Sozialgeschichtliche Studie über Eine Wiederkehrende Katastrophe*. Frankfurt am Main: Peter Lang.
Ilmoni, Immanuel. 1846-9. *Bidrag till Nordens Sjukdoms-Historia*. Helsingfors: J. Simelii arfvingar.
Iorga, Nicolae. 1970. *Istoria Armatei Româneşti*. Bucharest: Editura militară.
*Ipat'yevskaya Letopis'*. 1843. Vol. 2. Polnoye Sobraniye Russkikh Letopisei. St Petersburg: Tipografiya Eduard Pratsa.
Israel, Jonathan I. 1989. *Dutch Primacy in World Trade, 1585-1740*. Oxford: Oxford University Press.
Jankrift, Kay Peter. 2020. *Im Angesicht Der “Pestilenz”. Seuchen in Westfälischen Und Rheinischen Städten (1349–1600)*. Stuttgart: Franz Steiner Verlag.
Jennings, Ronald C. 1993. *Christians and Muslims in Ottoman Cyprus and the Mediterranean World, 1571- 1640*. New York: New York University Press.
Johansen, Paul, and Heinz von zur Mühlen. 1973. *Deutsch Und Undeutsch Im Mittelalterlichen Und Frühneuzeitlichen Reval*. Cologne: Böhlau.
Jordan, P. 1880. “Geschichte Der Pest in Estland Im Jahre 1710.” *St. Petersburger Kalender* 1880: 66–82.
Kalabukhov, N. I. 1961. “The Structures and Changes of the Natural Foci of Plague.” *Journal of Microbiology, Epidemiology and Immunobiology. Zhurnal Mikrobiologii, Epidemiologii*

*I Immunobiologii* 32: 877–83.
Kal'yu, V. V. 2004. *Krasnyi Gorodets*. Krasnogorodsk.
Kanold, Johann. 1721. *Einiger Marsilianischen Medicorum in Frantzösischer Sprache Ausgefertigte, Und Ins Teutsche üebersetzte Sendschreiben von Der Pest in Marsilien*. Leipzig: Philipp Wilhelm Stock.
Karpacz, Emilia. 2012. "„Opłakane Czasy" – Epidemia Dżumy W Krakowie W Latach 1707–1710. Przyczynek Do Badań Nad Upadkiem Królewskiego Miasta." *Folia Historica Cracoviensia* 18: 239–56.
Karpiński, Andrzej. 2000. *W Walce Z Niewidzialnym Wrogiem : Epidemie Chor'ob Zaka'znych W Rzeczypospolitej W XVI-XVIII Wieku I Ich Następstwa Demograficzne, Społeczno-Ekonomiczne I Polityczne*. Warsaw: Neriton : Instytut Historii PAN.
Kehrberg, Augustin. 1724. *Augustini Kehrberges Erleuterter Historisch-Chronologischer Abriß, Der Stadt Königsberg in Der Neu-Marck*. Berlin: Gottfried Gedicken.
Khaydarov, T. F. 2018. *Epokha "Chernoi Smerti" v Zolotoi Orde I Prilegayushchikh Regionakh, Konets XIII–Pervaya Polovina XV vv*. Kazan: Institut Istorii im. Sh. Mardzhani.
———. 2021. "Rol' Ordynsko-Russkoi Politii I Krupnykh Epidemicheskikh Vspyshek Chumy v Istorii Tatar' in Zolotoordynskoye Naslediye. Materialy Mezhdunarodnoi Nauchnoi Konferentsii 'Transformatsiya Politiko-Etnicheskoi Karty Vostochnoi Yevropy: Velikaya Vengriya, Volzhskaya Bulgariya I Obrazovaniye Zolotoi Ordy." In *Sbornik State*, edited by I. M. Mirgaleyev, 190–226. Kazan: Institut Istorii im. Sh. Mardzhani.
Khoroshkevich, A. L., S. V. Polekhov, V. A. Voronin, A. I. Grusha, A. A. Zhlutko, E. R. Skvairs, and A. G. Tyul'pin, eds. 2015. *Polotskie Gramoty XIII-Nachala XVI Veka*. Moscow: Universitet Dmitriya Pozharskogo.
Kimya, Osman. 2021. "17. Yüzyılda Osmanlı Devleti'nde Meydana Gelen Salgın Hastalıklar, Bunların Neden Olduğu Toplumsal Sarsıntılar ve Göçler." *Tarih Okulu Dergisi* 14: 2382–2403.
Kislichkina, Angelina A., Aleksandr G. Bogun, Lidiya A. Kadnikova, Nadezhda V. Maiskaya, Mikhail E. Platonov, Nikolai V. Anisimov, Elena V. Galkina, Svetlana V. Dentovskaya, and Andrey P. Anisimov. 2015. "Nineteen Whole-Genome Assemblies of Yersinia Pestis Subsp. Microtus, Including Representatives of Biovars Caucasica, Talassica, Hissarica, Altaica, Xilingolensis, and Ulegeica." *Genome Announcements* 3 (6): e01342–15.
Kislichkina, Angelina A., Aleksandr G. Bogun, Lidiya A. Kadnikova, Nadezhda V. Maiskaya, Viktor I. Solomentsev, Mikhail E. Platonov, Svetlana V. Dentovskaya, and Andrey P. Anisimov. 2017. "Eight Whole-Genome Assemblies of Yersinia Pestis Subsp. Microtus Bv. Caucasica Isolated from the Common Vole (Microtus Arvalis) Plague Focus in Dagestan, Russia." *Genome announcementsGenome* 5 (34): 5–6.
Kislichkina, Angelina A., Alexandr G. Bogun, Lidiya A. Kadnikova, Nadezhda V. Maiskaya, Viktor I. Solomentsev, Svetlana V. Dentovskaya, Sergey V. Balakhonov, and Andrey P. Anisimov. 2018. "Nine Whole-Genome Assemblies of Yersinia Pestis Subsp. Microtus Bv. Altaica Strains Isolated from the Altai Mountain Natural Plague Focus (No. 36) in Russia." *Genome Announcements* 6 (3): 1–2.
Kislichkina, Angelina A., Alexandr G. Bogun, Lidiya A. Kadnikova, Nadezhda V. Maiskaya, Viktor I. Solomentsev, Angelika A. Sizova, Svetlana V. Dentovskaya, Sergey V. Balakhonov, and Andrey P. Anisimov. 2018. "Six Whole-Genome Assemblies of Yersinia Pestis Subsp. Microtus Bv. Ulegeica (Phylogroup 0.PE5) Strains Isolated from Mongolian Natural Plague Foci." *Genome Announcements* 6 (25): 5–6.
Kiss, Andrea. 2017. "Droughts and Low Water Levels in Late Medieval Hungary II: 1361, 1439, 1443-4, 1455, 1473, 1480, 1482(?), 1502-3, 1506: Documentary Versus Tree-Ring (OWDA) Evidence." *Journal of Environmental Geography* 10: 43–56.
Kizik, Edmund. 2012. "Zarazy W Gdańsku Od XIV Do Połowy XVIII Wieku. Epidemie Oraz Liczba Ofiar W świetle Przekazów Nowożytnych Oraz Badaczy Współczesnych." In *Dżuma, Ospa, Cholera. W Trzechsetną Rocznicę Wielkiej Epidemii W Gdańsku I Na Ziemiach Rzeczypospolitej W Latach 1708–1711*, edited by Edmund Kizik. Gdańsk: Muzeum Historyczne Miasta Gdańska.

Kolia, Ioanna. 1990. “Athanasios Ieromonachos O Eks Agrafon (1656-1719). I Epistolografia Tou.” *Mesaionika Kai Nea Ellinika* 3: 215–54.

Kompaniets, O. V. 2022. “Nadzvichaini Yavyshcha Pryrody Ta Ikh Vplyv Na Sotsial’no-Ekonomichnyi Rozvytok Kryms'kogo Khanstva U XVII St.” *Zaporizhzhia Historical Review* 6: 23–31.

Koppmann, Karl, ed. 1884. *Die Chroniken Der Niedersächsischen Städte. Lübeck, Erster Band*. Leipzig: S. Hirzel.

Koretskii, V. I. 1980. “Solovetskii Letopisets Kontsa XVI v.” In *Letopisi I Khroniki*, 223–43. Moscow: Nauka.

Kotenev, Ye S., V. M. Dubyanskii, A. S. Volynkina, A. A. Zaitsev, A. N. Kulichenko, and S. L. Kravtsova. 2016. “Istoriya Epidemii Chumy Na Severnom Kavkaze I Sovremennyi Epidemicheskii Potentsial Ochagov Chumy.” *Meditsinskii Vestnik Severnogo Kavkaza* 11 (4): 612–16.

Krämer, Daniel. 2014. “Bevölkerung Und Wegenetz: Leben in Abgeschiedenheit.” In *Geschichte Des Kantons Nidwalden. Von Der Urzeit Bis 1850. Band 1*, 160–81. Stans: Historischer Verein Nidwalden.

Krämer, Sigrid. 1972. *Die Sogenannte Weihenstephaner Chronik. Text Und Untersuchung*. München: Arbeo-Gesellschaft.

Kriiska, Aivar. 1990. “Aruanne Arheoloogilistest Väljakaevamistest Lehmja-Pildiküla Asulakohal Ja Matmispaigal (Jüri Khk.) 27. Augustist Kuni 13. Septembrini 1990. a.” *(Manuscript of Field Report in Archaeological Research Collection at University of Tallinn)*.

———. 1994. “Tapa Tapa Tragonisi, Teisa Tragonisi Tuleb Veel Küll.” *Horisont*, 31–33.

———. 2017. “Kriiska, A. 1991. Der Siedlungsplatz Und Die Grabstätte Zu Lehmja-Pildiküla. Eesti Teaduste Akadeemia Toimetised. Humanitaar- Ja Sotsiaalteadused, 4, 392−396,” August.

*Kronika Polska, Litewska, Zmodzka I Wszystkiej Rusi Macieja Stryjkowskiego*. 1846. Warsaw: Nakład Gustawa Leona Glücksberga.

Kutyrev, Vladimir V., Galina A. Eroshenko, Vladimir L. Motin, Nikita Y. Nosov, Jaroslav M. Krasnov, Lyubov M. Kukleva, Konstantin A. Nikiforov, et al. 2018. “Phylogeny and Classification of Yersinia Pestis Through the Lens of Strains From the Plague Foci of Commonwealth of Independent States.” *Frontiers in Microbiology* 9 (May): 1–11.

Ladan, Rudolph. 2012. *Gezondheidszorg in Leiden in de Late Middeleeuwen*. Hilversum: Verloren.

Later, Christian. 2010. “Merowingerzeitliche Tuffplattengräber Und Frühmittelalterliche Kirchenbauten – Zu Den Anfängen Der Ehemaligen Pfarrkirche St. Benedikt in Starnberg.” *Bericht Der Bayerischen Bodendenkmalpflege* 52: 373–402.

Lavrov, L. I. 1984. “Stikhiinyye Bedstviya Na Severnom Kavkaze Do XIX v.” *Kavkazskii Etnograficheskii Sbornik* 8: 65–71.

Leader, Damian Riehl. 1988. *A History of the University of Cambridge. Volume 1: The University to 1546*. Cambridge: Cambridge University Press.

Leidinger, Georg, ed. 1915. *Veit Arnpeck: Sämtliche Chroniken*. Munich: Rieger’sche Universitäts-Buchhandlung.

———. , ed. 1918. “Chronica de Ducibus Bavariae.” In *Monumenta Germaniae Historica, Scriptores Rerum Germanicarum 19*, 139–75. Hanover: Impensis Bibliopolii Hahniani.

*Letopisnyi Sbornik Imenuyemyi Patriarsheyu Ili Nikonovskoyu Letopis’yu*. 1885. Polnoye Sobraniye Russkikh Letopisei 11. St Petersburg: Tipografiya Ministrestva Vnutrennikh Del.

Lewicki, Anatolii, ed. 1894. *Monumenta Medii Aevi Historica Res Gestas Poloniae Illustrantia, Tomus XIV*. Kraków: Sumptibus Academiae Litterarum Cracoviensis.

Lorenz, P. 1868-9. “Historisch-Medicinische Skizzen Aus Graubünden.” *Jahresberichte Der Naturforschenden Gesellschaft Graubünden* 14: 3–108.

Lythe, S. G. E. 1955. “Scottish Trade with the Baltic, 1550-1650.” In *Economic Essays in Commemoration of the Dundee School of Economics, 1931-1955*, edited by J. K. Eastham, 63–84. London: William Culross & Son.

Magyary-Kossa, Gyula. 1931. *Magyar Orvosi Emlékek. Értekezések a Magyar Orvostörténelem Köréből. III. Kötet. Adattár 1000-Től 1700-Ig*. Budapest: Magyar Orvosi Könyvkiadó Társulat.
———. 1940. *Magyar Orvosi Emlékek. Értekezések a Magyar Orvostörténelem Köréből. IV Kötet. Adattár 1700-Től 1800-Ig*. Budapest: Magyar Orvosi Könyvkiadó Társulat.
Mahling, Madlena, Klaus Neitmann, and Matthias Thumse, eds. 2018. *Liv-, Est- Und Kurländisches Urkundenbuch, Band 13. 1472-1479*. Cologne: Böhlau.
Maissen, F. 1971. "Die Letzte Pestepidemie in Der Eidgenossenschaft Und Ihre Folgen Für Graubünden 1665-1668." *Bündner Monatsblatt: Zeitschrift Für Bündner Geschichte, Landeskunde Und Baukultur* 11/12: 213–37.
Majander, Kerttu, Saskia Pfrengle, Arthur Kocher, Judith Neukamm, Louis du Plessis, Marta Pla-Díaz, Natasha Arora, et al. 2020. "Ancient Bacterial Genomes Reveal a High Diversity of Treponema Pallidum Strains in Early Modern Europe." *Current Biology: CB* 30 (19): 3788–3803.e10.
Majorossy, Judit. 2012. "'I Wish My Body to Hallowed Ground': Testamentary Orders of the Burghers of Late Medieval Pressburg about Their Own Burial." In *On Old Age. Approaching Death in Antiquity and the Middle Ages*, edited by Christian Krötzl and Katariina Mustakallio. Turnhout: Crepols.
Malewicz, Małgorzata Hanna. 1980. *Zjawiska Przyrodnicze W Relacjach Dziejopisarzy Polskiego średniowiecza*. Wroclaw: Zakład Narodowy im. Ossolińskich--Wydawn. Polskiej Akademii Nauk.
Malve, Martin, and Andres Tvauri. 2022. "The 1710 Plague Burial Ground on the Outskirts of Tallinn." *Archaeological Fieldwork in Estonia* 2021: 235–248.
Malve, Malve, and Heiki Valk. 2021. "14th-Century Cemetery at Otepää, Southern Estonia." *Archaeological Fieldwork in Estonia* 2020: 143–150.
Manolova-Nikolova, Nadya. 2004. *Chumavite Vremena (1700-1850)*. Sofia: IF-94.
Manyshev, S. B. 2015. "Iz Istorii Epidemii v Dagestane v XVIII v." In *Lavrovskii Sbornik. Material XXXVIII I XXXIX Sredneaziatsko-Kavkazskikh Chtenii 2014-2015 Gg. Etnologiya, Istoriya, Arkheologiya, Kul'turologiya*, edited by Yu Yu Karpov and M. Ye Rezvan, 95–99. St Petersburg: MAE RAN.
Masson, David, ed. 1880. *The Register of the Privy Council of Scotland, Vol. 3, A.D. 1578-1585*. Edinburgh: H.M. General Getister House.
Masson, Paul. 1896. *Histoire Du Commerce Français Dans Le Levant Au XVIIe Siècle*. Paris: Librarie Hachette & C.
*Materialy Po Istorii Russko-Gruzinskikh Otnoshenii Vtoroi Poloviny XVIII Veka. Chast' III*. 1988. Tbilisi.
Maur, Eduard. 1989. "Příspěvek K Demografické Problematice Předhusitských Čech (1348–1419)." *Acta Universitatis Carolinae. Philosophica et Historica* 1: 7–71.
*Mémoires et Lettres Sur La Guerre de La Valteline de Henri Duc de Rohan*. 1758. Paris: Chez Vincent, Imprimeur-Libraire.
Mencke, Johann Burchard, ed. 1728. *Achilles Pirminius Gasser, Annales Augstburgenses*. Vol. 1. Scriptores Rerum Germanicarum. Leipzig: Johann Christian Martin.
Mertens, Thom. 1999. "Rondom Het Sterfbed van Lubbert Ten Busch. De Moderne Devoten En de Pest Te Deventer in 1398." In *De Pest in de Nederlanden: Medisch-Historische Beschouwingen 650 Jaar Na de Zwarte Dood*, 141–158. Brussels: Koninklijke Academie voor Geneeskunde van België.
Mertzios, Konstantinos D. 1969-1970. "Eidiseis Peri Tis Stereas Ellados Ek Ton Archeion Tis Venetias, 1690-1736." *Epetiris Etaireias Stereoelladikon Meleton* 2: 339–436.
Mironov, N. P. 1958. "The Past Existence of Natural Foci of Plague in the Steppes of Southern Europe." *Journal of Microbiology, Epidemiology and Immunobiology. Zhurnal Mikrobiologii, Epidemiologii I Immunobiologii* 29: 1193–98.
Molvygin, Andreas. 1969. "Über Die Münz- Und Geldgeschichte Estlands Vom Beginn Der Einheimischen Münzprägung Bis Zum II. Viertel Des 15. Jahrhunderts." *Nordisk Numismatisk Årsskrift*, 37–65.
Morelli, Giovanna, Yajun Song, Camila J. Mazzoni, Mark Eppinger, Philippe Roumagnac,

David M. Wagner, Mirjam Feldkamp, et al. 2010. “Yersinia Pestis Genome Sequencing Identifies Patterns of Global Phylogenetic Diversity.” *Nature Genetics* 42 (12): 1140–43.
Morozova, Irina, Artem Kasianov, Sergey Bruskin, Judith Neukamm, Martyna Molak, Elena Batieva, Aleksandra Pudło, Frank Ruhli, and Verena Schuenemann. 2020. “New Ancient Eastern European Yersinia Pestis Genomes Illuminate the Dispersal of Plague in Europe.” *Philosophical Transactions of the Royal Society of London. Series B, Biological Sciences* 375: 20190569.
Motylewicz, Jerzy. 1993. *Miasta Ziemi Przemyskiej I Sanockiej W Drugiej Połowie XVII I W XVIII Wieku*. Przemyśl: Południowo-Wschodni Instytut Nauk. w Przemyślu.
Możejko, Beata. 2012. “Zarazy W Późnośredniowiecznym Gdańsku.” In *Dżuma, Ospa, Cholera. W Trzechsetną Rocznicę Wielkiej Epidemii W Gdańsku I Na Ziemiach Rzeczypospolitej W Latach 1708–1711*, edited by Edmund Kizik, 43–61. Gdańsk: Muzeum Historyczne Miasta Gdańska.
Müller, Konrad M. 2006. *Das “Grosse Sterben” Im Allgäu: Pest Und Andere Seuchen in Mittelalter Und Früher Neuzeit*. Vol. 2004–5. Memminger Geschichtsblätter. Memmingen.
Müllner, Johannes. 1984. *Die Annalen Der Reichsstadt Nürnberg von 1623, Teil II, Von 1351-1469*. Edited by Gerhard HIrschmann. Niremberg: Im Selbstverlag des Stadtrats Nürnberg.
———. 2003. *Die Annalen Der Reichsstadt Nürnberg von 1623, Teil III: 1470-1544*. Edited by Gerhard HIrschmann. Niremberg: Im Selbstverlag des Stadtrats Nürnberg.
Munro, John H. 2007. “South German Silver, European Textiles, and Venetian Trade with the Levant and Ottoman Empire, C. 1370 to C. 1720: A Non-Mercantilist Approach to the Balance of Payments Problem.” In *Relazioni Economiche Tra Europa E Mondo Islamico, Secoli XIII - XVIII/ Europe’s Economic Relations with the Islamic World, 13th - 18th Centuries*, edited by Simonetta Cavaciocchi, 907–62. Florence: Le Monnier.
Muratori, Ludovico Antonio, ed. 1734. “Vitae Romanorum Pontificum.” In *Rerum Italicarum Scriptores*. Vol. 3.2. Milano: Typographia Societatis Palatinae.
Myrdal, Janken. 2003. *Digerdöden, Pestvågor Och ödeläggelse. Ett Perspektiv På Senmedeltidens Sverige*. Stockholm: Sällskapet Runica et Mediævalia.
Mytsyuk, Oleksandr. 1938. *Narysy z sotsial'ne-hospodars'koi istorii Pidkarpats'koi Rusi, Vol. 2*. Prague: Hrdlička.
Nada Patrone, Anna Maria, and Irma Naso. 1978. *Le Epidemie Del Tardo Medioevo Nell’area Pedemontana*. Torino: Centro Studi Piemontesi.
Naloyeva, Ye D. 2015. *Kabarda v Pervoy Polovine XVIII Veka: Genezis Adygskogo Feodal’nogo Sotsiuma I Problemy Sotsial'no-Politicheskoy Istorii*. Nalchik: Pechatnyi Dvor.
Namouchi, Amine, Meriam Guellil, Oliver Kersten, Stephanie Hänsch, Claudio Ottoni, Boris Schmid, Elsa Pacciani, et al. 2018. “Integrative Approach Using Yersinia Pestis Genomes to Revisit the Historical Landscape of Plague during the Medieval Period.” *Proceedings of the National Academy of Sciences* 115: 201812865.
Napiersky, C. E. 1846. *Beitrag Zur Geschichte Des Ehemaligen Bisthums Dorpat*. Riga.
Nasonov, A., ed. 1941. *Pskovskie Letopisi, Vypusk Pervyi*. Moscow and Leningrad: Izdatel’stvo Akademii Nauk SSSR.
Nasonov, A. N., ed. 1955. *Pskovskie Letopisi, Vypusk Vtoroi*. Moscow: Izdatel’stvo Akademii Nauk SSSR.
Niinre, A. 1990. “Pildikülast Leitud Luustikest Ja Nende Panustest.” *(Manuscript of Field Report in Archaeological Research Collection at University of Tallinn)*.
Nikolov, Z., and M. Michev. 1978. “O Chumnykh Epidemiyakh Na Bolgarskoi Zemle Na Protyazhenii XV-XIX Vekov.” *Asklepii: Bolgaro-Sovetskii Ezhegodnik Istorii I Teorii Meditsiny* 4.
Noordegraaf, Leo, and Gerrit Valk. 1996. “De Gave Gods. De Pest in Holland Vanaf de Late Middeleeuwen.” In . Amsterdam: Bert Bakker.
Nothdurfter, Hans, Thomas Kersting, and Brigitte Gebauer. 2019. “Der Kirchenbau Und Die Friedhöfe.” In *St. Prokulus in Naturns*. Bozen: Athesia.

Oefele, Andreas Felix von, ed. 1763. “Anonymi Chronicon Noribergense et Locorum Vicinorum.” In *Rerum Boicarum Scriptores*, 1:322–29. Augsburg: Ignaz Adam Veith & Franz Anton Veith.
Oja, Tiiu. 1996. “Katk Põhjasõja Ajal Eestis.” In *Artiklite Kogumik Eesti Ajalooarhiivi 75.aastapäevaks*, 217–53. Tartu: Eesti Ajalooarhiiv.
Oram, Richard D. 2006. “‘It Cannot Be Decernit Quha Are Clean and Quha Are Foulle.’ Responses to Epidemic Disease in Sixteenth-and Seventeenth-Century Scotland.” *Renaissance & Reformation/Renaissance et Reforme* 30 (4): 13–39.
Panzac, Daniel. 1973. “La Peste a Smyrne Au XVIIIe Siecle.” *Annales : économies, Sociétés, Civilisations* 28 (4): 1071–93.
———. 1985. *La Peste Dans l’Empire Ottoman, 1700-1850*. Leuven: Peeters.
Papke, Hans-Ulrich. 1947. “Die Pest in Frankfurt.” PhD, Johann Wolfgang Goethe-Universität.
Parkhill, J., B. W. Wren, N. R. Thomson, R. W. Titball, M. T. Holden, M. B. Prentice, M. Sebaihia, et al. 2001. “Genome Sequence of Yersinia Pestis, the Causative Agent of Plague.” *Nature* 413 (6855): 523–27.
Pärn, A. 1987. “Mogil’nik Mõisaküla-Margu.” *Eesti NSV Teaduste Akadeemia Toimetised. Bioloogia = Izvestiia Akademii Nauk Estonskoi SSR. Biologiia* 4: 357–59.
Parts, T. 2010. “1710. Aasta Katku Ohvrid Tallinnas.” *Tuna* 3: 92–93.
Pękacka-Falkowska, Katarzyna. 2019. *Dżuma W Toruniu W Trakcie III Wojny Północnej*. Lublin: Towarzystwo Naukowe Katolickiego Uniwersytetu Lubelskiego Jana Pawła II.
Penskoi, V. V., O. N. Polukhin, S. N. Borisov, and R. A. Dmitrakov. 2021. “Vsadnik Na Belom Kone: Chuma Ivana Groznogo.” *Problemy Sotsial’noi Gigiyeny, Zdravokhraneniya I Istorii Meditsiny* 29 (1): 173–79.
Petersone-Gordina, Elina, Charlotte Roberts, Andrew R. Millard, Janet Montgomery, and Guntis Gerhards. 2018. “Dental Disease and Dietary Isotopes of Individuals from St Gertrude Church Cemetery, Riga, Latvia.” *PloS One* 13 (1): e0191757.
Petersone-Gordina, E., J. Montgomery, A. R. Millard, C. Roberts, D. R. Gröcke, and G. Gerhards. 2020. “Investigating the Dietary Life Histories and Mobility of Children Buried in St Gertrude Church Cemetery, Riga, Latvia, 15th–17th Centuries Ad.” *Archaeometry* 62 (S1): 3–18.
Petresco, G. Z. 1933. *Les Dernières épidémies de Peste Dans Les Pays Roumains Au XVIIIe et Au XIXe Siècle*. Bucharest: Tipografia Culture.
Pfaff, Karl. 1845. *Geschichte Der Stadt Stuttgart, Nach Archival-Urkunden Und Andern Bewährten Quellen. Erster Theil*. Stuttgart: C. A. Sonnewald’schen Buchhandlung.
Pieth, Friedrich. 1945. *Bündnergeschichte*. Chur: Schuler.
Pogodin, M., ed. 1837. *Pskovskaya Letopis’ Izdannaya Na Izhdivenii Obshchestva Istorii I Drevnostei Rossiiskikh, Pri Moskovskom Universitete*. Moscow: Universitetskaya Tipografiya.
Porter, Stephen. 2005. *London’s Plague Years*. Stroud: Tempus.
Pribyl, Kathleen. 2017. *Farming, Famine and Plague: The Impact of Climate in Late Medieval England*. Springer.
Prokhorov, D. A. 2016. “Posledstviya Prirodnykh Kataklizmov I Stikhiinykh Bedstvii Na Krymskom Poluostrove v Opisaniyakh Avtorov I Dokumentakh XVII-XVIII vv.” *Bosporskiye Issledovaniya* 33: 319–47.
Rahelinirina, Soanandrasana, Minoarisoa Rajerison, Sandra Telfer, Cyril Savin, Elisabeth Carniel, and Jean-Marc Duplantier. 2017. “The Asian House Shrew Suncus Murinus as a Reservoir and Source of Human Outbreaks of Plague in Madagascar.” *PLoS Neglected Tropical Diseases* 11: e0006072.
Rascovan, Nicolás, Karl-Göran Sjögren, Kristian Kristiansen, Rasmus Nielsen, Eske Willerslev, Christelle Desnues, and Simon Rasmussen. 2019. “Emergence and Spread of Basal Lineages of Yersinia Pestis during the Neolithic Decline.” *Cell* 176 (1-2): 295–305.e10.
Rayfield, Donald. 2012. *Edge of Empires: A History of Georgia*. London: Reaktion Books.
Raymond, André. 1972. “Les Grandes épidémies de Peste Au Caire Aux XVII et XVIII

Siècles.” *Bulletin D’études Orientales* 25: 203–10.
Redstone, Vincent B/. 1907. *Calendar of Pre-Reformation Wills, Testaments, Probates at Bury St Edmunds*. Ipswich: W.E. Harrison.
Restifo, Giuseppe. 2005. *I Porti Della Peste. Epidemie Mediterranee Fra Sette E Ottocento*. Messina: MESOGEA.
Risberg, Sara, ed. 2008. *Auctoritate Papae. The Church Province of Uppsala and the Apostolic Penitentiary 1410–1526*. Stockholm: National Archives of Sweden.
*Rogozhskii Letopisets*. 2000. Polnoye Sobraniye Russkikh Letopisei 15. Moscow: Yazyki Russkoi Kul’tury.
Rogudeyev, V. V. 2007. “Kompleksy I Otdel’nyye Nakhodki XVIII-XIX Vekov.” *Arkheologicheskie Zapiski* 5: 65–81.
Rorke, Martin. 2001. “Scottish Overseas Trade, 1275/86-1597.” PhD, University of Edinburgh.
Rudenko, S. A. 2013. “Mogil’niki Volzhskoi Bulgarii I Bulgarskogo Ulusa Zolotoi Ordy Kak Istochnik Po Izucheniyu Mirovozzreniya Srednevekovogo Naseleniya Volgo-Kam'ya: Voprosy Sistematizatsii.” In *Mirovozreniye Naseleniya Yuzhnoi Sibiri I Tsentral’noi Azii v Istoricheskoi Retrospektive*, edited by P. K. Dashkovskiy, 72–101. barnaul: Izdatel’stvo Altaiskogo Gosudarstvennogo Universiteta.
Russell, J. C. 1976. “The Earlier Medieval Plague in the British Isles.” *Viator* 7: 65–78.
Russow, Balthasar. 1584. *Chronica Der Provinz Lyfflandt*. Barth: Andreas Seitner.
Saag, Lehti, Margot Laneman, Liivi Varul, Martin Malve, Heiki Valk, Maria A. Razzak, Ivan G. Shirobokov, et al. 2019. “The Arrival of Siberian Ancestry Connecting the Eastern Baltic to Uralic Speakers Further East.” *Current Biology: CB* 29 (10): 1701–11.e16.
Sahm, Wilhelm. 1905. *Geschichte Der Rest in Ostpreussen*. Leipzig: Duncker & Humblot.
Schmid, Boris V., Ulf Büntgen, W. Ryan Easterday, Christian Ginzler, Lars Walløe, Barbara Bramanti, and Nils Chr Stenseth. 2015. “Climate-Driven Introduction of the Black Death and Successive Plague Reintroductions into Europe.” *Proceedings of the National Academy of Sciences of the United States of America* 112 (10): 3020–25.
Schmölzer, Hilde. 2015. *Die Pest in Wien*. Innsbruck: Haymon.
Schnurrer, Ludwig. 1987. “Die Pest in Rothenburg Im Ausgehenden Mittelalter (1472/73, 1483/84, 1484/95).” *Die Linde* 69: 21–24.
Schnyder, Franz. 1932. “Pest Und Pestverordnungen Im Alten Luzern.” *Der Geschichtsfreund. Mitteilungen Des Historischen Vereins Zentralschweiz* 87: 102–18.
Schröder, Johann Heinrich. 1854. *Om Pesten I Stockholm, 1710*. Stockholm: Hos. Joh. Beckman.
Schwalm, Jakob, ed. 1895. *Die Chronica Novella Des Hermann Korner*. Göttingen: Vandenhoeck & Ruprecht.
Schwartz, Philipp, ed. 1905. *Liv-, Est- Und Kurländisches Urkundenbuch, Band 11. 1450-1459*. Riga: J. Deubner.
Schwarz, Klaus. 1996. “Die Pest in Bremen. Epidemien Und Freier Handel in einer Deutschen Hafenstadt, 1350-1713.” Bremen: Staatsarchiv Bremen.
Seguin-Orlando, Andaine, Caroline Costedoat, Clio Sarkissian, Stéfan Tzortzis, Célia Kamel, Norbert Telmon, Love Dalén, Catherine Thèves, Michel Signoli, and Ludovic Orlando. 2021. “No Particular Genomic Features Underpin the Dramatic Economic Consequences of 17th Century Plague Epidemics in Italy.” *iScience* 24: 102383.
Seifert, Lisa, Ingrid Wiechmann, Michaela Harbeck, Astrid Thomas, Gisela Grupe, Michaela Projahn, Holger Scholz, and Julia Riehm. 2016. “Genotyping Yersinia Pestis in Historical Plague: Evidence for Long-Term Persistence of Y. Pestis in Europe from the 14th to the 17th Century.” *PloS One* 11: e0145194.
Seifert, Mathias, Christine Cooper, Marcel Keller, Meriam Guellil, and Christiana L. Scheib. 2020. “Der Pestfriedhof des 17. Jahrhunderts.” In: Archäologischer Dienst Graubünden, Lorena Burkhardt (Ed.) *Domat/Ems, Sogn Pieder: vom frühmittelalterlichen Herrenhof zum neuzeitlichen Pestfriedhof*, 2:241–259. Chur: Somedia-Buchverlag.
*Seyid-Muhammed Riza. Sem’ Planet v Izvestiyakh O Tsaryakh Tatarskikh*. 2019. Kazan: Institut Istorii im. Sh. Mardzhani.

Shrewsbury, J. F. D. 1971. *A History of Bubonic Plague in the British Isles*. Cambridge: Cambridge University Press.
Signoli, Michel. 2022. “History of the Plague of 1720-1722, in Marseille.” *La Presse Médicale* 51 (3): 104138.
Signoli, Michel, and Stéfan Tzortzis. 2018. “La Peste à Marseille et Dans Le Sud-Est de La France En 1720–1722: Les épidémies d’Orient de Retour En Europe.” *Cahiers de La Méditerranée* 96: 217–30.
Sincera, Rege. 1810. “Observations, Both Historical and Moral upon the Burning of London, September, 1666. With an Account of the Losses, and a Most Remarkable Parallel between London and Moscow, Both as to the Plague and Fire.” In *Harleian Miscellany 7*, 324–43. London: Robert Dutton.
Slack, Paul. 1985. *The Impact of the Plague in Tudor and Stuart England*. London: Routledge.
Slavin, Philip. 2021. “Out of the West: Formation of a Permanent Plague Reservoir in South-Central Germany (1349–1356) and Its Implications.” *Past & Present* 252 (1): 3–51.
———. 2022. “Reply: Out of the West — and Neither East, nor North, nor South.” *Past & Present* 256 (1): 325–60.
Sloane, Barney. 2011. *The Black Death in London*. Stroud: The HIstory Press.
Solleder, Fridolin. 1938. *München Im Mittelalter*. München: R. Oldenbourg.
Song, Yajun, Zongzhong Tong, Jin Wang, Li Wang, Zhaobiao Guo, Yanpin Han, Jianguo Zhang, et al. 2004. “Complete Genome Sequence of Yersinia Pestis Strain 91001, an Isolate Avirulent to Humans.” *DNA Research: An International Journal for Rapid Publication of Reports on Genes and Genomes* 11 (3): 179–97.
Spyrou, Maria A., Marcel Keller, Rezeda I. Tukhbatova, Christiana L. Scheib, Elizabeth A. Nelson, Aida Andrades Valtueña, Gunnar U. Neumann, et al. 2019. “Phylogeography of the Second Plague Pandemic Revealed through Analysis of Historical Yersinia Pestis Genomes.” *Nature Communications* 10 (1). https://doi.org/10.1038/s41467-019-12154-0.
Spyrou, Maria A., Lyazzat Musralina, Guido A. Gnecchi Ruscone, Arthur Kocher, Pier-Giorgio Borbone, Valeri I. Khartanovich, Alexandra Buzhilova, et al. 2022. “The Source of the Black Death in Fourteenth-Century Central Eurasia.” *Nature* 606 (7915): 718–24.
Spyrou, Maria A., Rezeda Tukhbatova, Michal Feldman, Joanna Drath, Sacha Kacki, Julia Beltrán de Heredia Bercero, Susanne Arnold, et al. 2016. “Historical Y. Pestis Genomes Reveal the European Black Death as the Source of Ancient and Modern Plague Pandemics.” *Cell Host & Microbe* 19: 874–881.
Stahleder, Helmuth, ed. 2020. *Die Protokolle Des Münchner Stadtrats 1501 Bis September 1532 (Teil II, Bände 5 Bis 10)*. Munich: Bayerisches Hauptstaatsarchiv.
Stefanov, A. T. 2011. “Arkhiv Kreposti Sv. Dimitriya Rostovskogo (Nyne - Gorod Rostov-Na-Donu).” *Donskii Vremennik* 19: 101–5.
Steinhilber, Wilhelm. 1956. *Das Gesundheitswesen Im Alten Heilbronn 1281–1871*. Heilbronn: Archiv d. Stadt.
Stenseth, Nils Chr, Yuxin Tao, Chutian Zhang, Barbara Bramanti, Ulf Büntgen, Xianbin Cong, Yujun Cui, et al. 2022. “No Evidence for Persistent Natural Plague Reservoirs in Historical and Modern Europe.” *Proceedings of the National Academy of Sciences of the United States of America* 119 (51): e2209816119.
Sturm, Patrick. 2014. *Leben Mit Dem Tod in Den Reichsstätten Esslingen, Nördlingen Und Schwäbisch Hall. Epidemien Und Deren Auswirkungen Vom Frühen 15. Bis Zum Frühen 17. Jahrhundert*. Ostfildern: Jan Thorbecke.
Sultan, Halim Geray. 2004. *Rozovyi Kust Khanov Ili Istoriya Kryma*. Translated by A. Il’mi. Simferopol: Dolya.
Suntsov, Victor. 2019. “Origin of the Plague: Prospects of Ecological - Molecular - Genetic Synthesis.” *Herald of the Russian Academy of Sciences* 89: 271–78.
———. 2021. “Genomogenesis of the Plague Bacteria Yersinia Pestis as a Process of Mosaic Evolution.” *Russian Journal of Genetics* 57: 139–51.
Suponitskii, M. V. 2004. *“Chernaya Smert’”. K Zagadka m Pandemii Chumy 1346-1351 Gg.*

Universum.
———. 2005. *Gde Skryvaetsya Chuma?* Universum.
Suponitskii, M. V., and N. S. Suponitskaya. 2006. *Ocherki Istorii Chumy*. Moscow: Vuzovskaya Kniga.
Süreyya, Mehmed. 1890. *Sicill-i Osmanî*. Istanbul: Maba'a-'i 'mire.
Susat, Julian, Joanna Bonczarowska, Elīna Pētersone-Gordina, Alexander Immel, Almut Nebel, Guntis Gerhards, and Ben Krause-Kyora. 2020. "Yersinia Pestis Strains from Latvia Show Depletion of the Pla Virulence Gene at the End of the Second Plague Pandemic." *Scientific Reports* 10: 14628.
*The Levant Voyage of the Blackham Galley (1696–1698): The Sea Journal of John Looker, Ship's Surgeon*. 2022. London: The Hakluyt Society.
*The Negotiations of Sir Thomas Roe, in His Embassy to the Ottoman Porte, from the Year 1621 to 1628 Inclusive*. 1740. London: Samuel Richardson.
Thurley, Clifford A., Dorothea Thurley, E. S. Leedham-Green, and Rosemary Rodd. 1994-1996. *Index of the Probate Records of the Consistory Court of Ely, 1449-1858*. London: British Record Society.
Tikhomirov, M. N., ed. 1965. *Novgorodskaya Vtoraya (Arkhivnaya) Letopis'*. Polnoye Sobraniye Russkikh Letopisei 30. Moscow: Nauka.
Topping, Peter. 1977. "The Post-classical Documents." In *Studies on Latin Greece A.D. 1205-1715*. London: Variorum Collected Studies Series.
Tsiamis, Costas. 2010. "Epidemic Waves during Justinian's Plague in the Byzantine Empire (6th-8th C. AD)." *Vesalius: Acta Internationales Historiae Medicinae* Suppl (December): 12–18.
Ulashchik, N. N., ed. 1980. *Letopisi Belorussko-Litovskie*. Vol. 35. Polnoye Sobraniye Russkikh Letopisei. Moscow: Nauka.
*Ustyuzhskiye I Vologodskiye Letopisi XVI-XVIII vv*. 1982. Vol. 37. Polnoye Sobraniye Russkikh Letopisei. Leningrad: Nauka.
Valk, H. 1985. "Der Dorffriedhof von Mäletjärve." *Eesti NSV Teaduste Akadeemia Toimetised. Bioloogia = Izvestiia Akademii Nauk Estonskoi SSR. Biologiia* 4: 376–79.
———. 1997. "Archaeological Investigations in Otepää and Its Surroundings in 1996." *Stilus. Eesti Arheoloogiaseltsi Väljaanne* 7: 124–29.
Van Veen, J. S. 1903. "De Pest En Hare Bestrijding in Gelderland, in Het Bijzonder Te Arnhem." *Bijdragen En Mededelingen Gelre* 6: 1–66.
Varlık, Nükhet. 2015. *Plague and Empire in the Early Modern Mediterranean World. The Ottoman Experience, 1347–1600*. Cambridge: Cambridge University Press.
———. 2022. "The Rise and Fall of a Historical Plague Reservoir. The Case of Ottoman Anatolia." In *Disease and the Environment in the Medieval and Early Modern Worlds*, edited by Lori Jones, 159–83. London: Routledge.
Vasil'yev, K. G., and A. Ye Segal. 1960. *Istoriya Epidemii v Rossii*. Moscow: Medgiz.
Vătămanu, Nicolae. 1972. "Ciuma de Dupa Războieni (1476)." In *Din Istoria Luptei Antiepidemice în România: Studii și Note*, edited by Gheorghe V. Brătescu, 35–37.
Vierdag, J. 2020. "Sint Jansbeek: Bakermat van Arnhem." In *Sint Jansbeek Te Arnhem. Levensader van Een Stad.*, edited by W. Jansen, J. Storms, and S. Weigand, 8–25. Amersfoort: Jansen & de Feijter / Het Colofon.
Von der Ropp, Goswin Freiherr, ed. 1888. *Hanserecesse von 1431-1476. Zweite Abtheilung, Fünfter Band*. Leipzig: Duncker & Humblot.
Von Planta, Peter Conradin. 2020. "Die Kirchen von Domat/Ems." In: Archäologischer Dienst Graubünden, Lorena Burkhardt (Ed.) *Domat/Ems, Sogn Pieder: vom frühmittelalterlichen Herrenhof zum neuzeitlichen Pestfriedhof*, 2:359–393. Chur: Somedia-Buchverlag.
Von Rauch, Moriz. 1913. *Urkundenbuch Der Stadt Heilbronn. Zweiter Band (1476—1500)*. Stuttgart: W. Kohlhammer.
Von Sprecher, J. A. 1942. "Die Pest in Graubünden Währen Der Kriege Und Unruhen 1628-1635." *Bündnerisches Monatsblatt : Zeitschrift Für Bündnerische Geschichte, Landes- Und Volkskunde* 1: 21–32.

Walawander, Antoni. 1932. *Kronika Klęsk Elementarnych W Polsce I W Krajach Sąsiednich W Latach 1450-1586*. Lwów: Instytut Popierania Polskiej Twórczości Naukowej.
Wattenbach, W., ed. 1851. “Annales Mellicenses.” In *Monumenta Germaniae Historica, Scriptores 9*, 480–84. Hannover: Impensis Bibliopolii Aulici Hahniani.
Weiland, L., ed. 1877. “Sächsische Weltchronik, Vierte Bairische Fortsetzung.” In *Monumenta Germaniae Historica. Deutsche Chroniken 2*, 352–84. Hannover: Hahnsche Buchhandlung.
Wolthuis, J. 2020. “Sint Jansbeek Weer Bovengronds.” In *Sint Jansbeek Te Arnhem. Levensader van Een Stad.*, edited by W. Jansen, J. Storms, and S. Weigand, 36–45. Amersfoort: Jansen & de Feijter / Het Colofon.
Xu, Lei, Leif Stige, Kyrre Kausrud, Tamara Ben Ari, Shuchun Wang, Xiye Fang, Boris Schmid, Qi-Yong Liu, Nils Chr Stenseth, and Zhibin Zhang. 2014. “Wet Climate and Transportation Routes Accelerate Spread of Human Plague.” *Proceedings. Biological Sciences / The Royal Society* 281: 20133159.
Zagorskii, V. 1897. “Chuma v Litve (istoricheskaya Zametka).” *Protokoly Imperatorskogo Vilenskago Meditsiniskago Obshchestva* 1897: 113–33.
Zapnik, Jörg. 2007. *Pest Und Krieg Im Ostseeraum: Der “Schwarze Tod” in Stralsund Während Des Großen Nordischen Krieges (1700-1721)*. Hamburg: Verlag Dr. Kovač.
Zeiller, Martin. 1646. *Topographia Hassiae et Regionum Vicinarum. Das Ist Beschreibung Der Vorne[m]bsten Stätte Und Plätze in Hessen*. Frankfurt-am-Main: Matthäus Merian.
Zhgenti, Ekaterine, Shannon L. Johnson, Karen W. Davenport, Gvantsa Chanturia, Hajnalka E. Daligault, Patrick S. Chain, and Mikeljon P. Nikolich. 2015. “Genome Assemblies for 11 Yersinia Pestis Strains Isolated in the Caucasus Region.” *Genome Announcements* 3 (5): 1030–1015.
Zielman, Gl, and W. A. Baetsen, eds. 2020. *Wat de Nieuwe Sint Jansbeek Boven Water Bracht: Dood En Leven in Het Arnhemse Verleden; Archeologisch Onderzoek Sint Jansbeek Te Arnhem – Deel 1: Sporen, Vondsten En Hun Context. RAAP Report 4476-1*. Weesp: RAAP Archeologisch Adviesbureau B.V.
Zimin, A. A. 1950. “Kratkie Letopistsy XV-XVI vv.” *Istoricheskii Arkhiv* 5: 3–39.
Zuijderduijn, Jaco. 2013. “Living La Vita Apostolica. Life Expectancy and Mortality of Nuns in Late-Medieval Holland.” *CGEH Working Paper Series*.
